# Latent neural network representations of the brain reflect broad-scale adolescent phenotypic variation

**DOI:** 10.64898/2026.04.16.718661

**Authors:** Andreas Dahl, Esten Leonardsen, Dag Alnæs, Lars T. Westlye, the Alzheimer’s Disease Neuroimaging Initiative

## Abstract

The adolescent brain is attuned to social and environmental exploration, allowing behavioral adaptation as experiences shape lasting patterns of morphological organization. Using a convolutional neural network on longitudinal structural MRI data, we assess the early part of this developmental window and derive latent brain representations reflecting patterns of structural variability linked to personal, social, and neighborhood conditions in adolescence. These representations offer a flexible framework for mapping brain-trait associations in adolescence and beyond.

## Main text

Adolescence is characterized by dynamic and nonlinear changes in brain structure and function^1^, with an accelerating reduction in gray matter during late childhood and early adolescence. This process supports the adaptation to increasing external demands, including navigating more complex social relationships, and developing greater autonomy. Structural MRI studies relate adolescent personal, familial and neighborhood contexts to brain morphology, but effects are modest and anatomical localization is inconsistent^2^.

Findings that many complex traits do not map consistently onto pre-defined brain regions have prompted calls for approaches that better capture the distributed and heterogeneous nature of brain-trait relationships^3–5^. This includes approaches that relaxes a priori assumptions of localization and instead rely on data-driven and integral procedures to characterize relevant structural and functional properties of the brain^3^. Such approaches are particular important in childhood and adolescence where the brain is subject to rapid development and adaptation, which might not be adequately captured when investigating morphological regions of interest in isolation^6,7^.

Deep learning approaches are able to derive complex representations from neuroimaging data^3^. Convolutional neural networks (CNNs) are particularly well suited as they learn spatially coherent and condensed representations directly from minimally processed MRI^8^, and recognize brain patterns associated with traits such as age^8^, sex^9^ and mental health status^10^.

Here, we expand on this work by deriving latent representations of two waves of structural MRI data from 9-13 year old children from the Adolescent Brain Cognitive Development (ABCD) Study^11^, using a Simple Fully Convolutional Network (SFCN) trained to predict multiple targets concurrently^12,13^. Multitask learning enables a richer embedding space that is not tied to any single predictive target and can plausibly capture general features of brain morphology not limited by pre-defined anatomical boundaries. We relate these embedding dimensions to the adolescent phenome spanning personal, relational and socioenvironmental domains, both cross-sectionally and across time, from childhood to early adolescence. The overall goal is to capture novel dimensions of brain morphology that explain adolescent phenotypic variance and its developmental change.

We trained and validated an SFCN multi-task model on 65,290 minimally processed MRI scans from 59,454 healthy participants from 30 different sources (Supplementary Table 1). The training set consisted of 59,171 scans from 53,335 individuals. The average age of the training fold was 53.4 (± one standard deviation of 21.3 years), and 51.8% were female. The validation fold consisted of 6119 scans from 6119 individuals (Online Methods Fig. 1B). The final full held-out test data after quality control consisted of 18,813 images of 11,497 individuals from the ABCD study, 11,103 were from the baseline session (mean age = 9.9 years), and 7710 from the 2-year follow-up (mean age = 11.9 years).

**Fig. 1.**
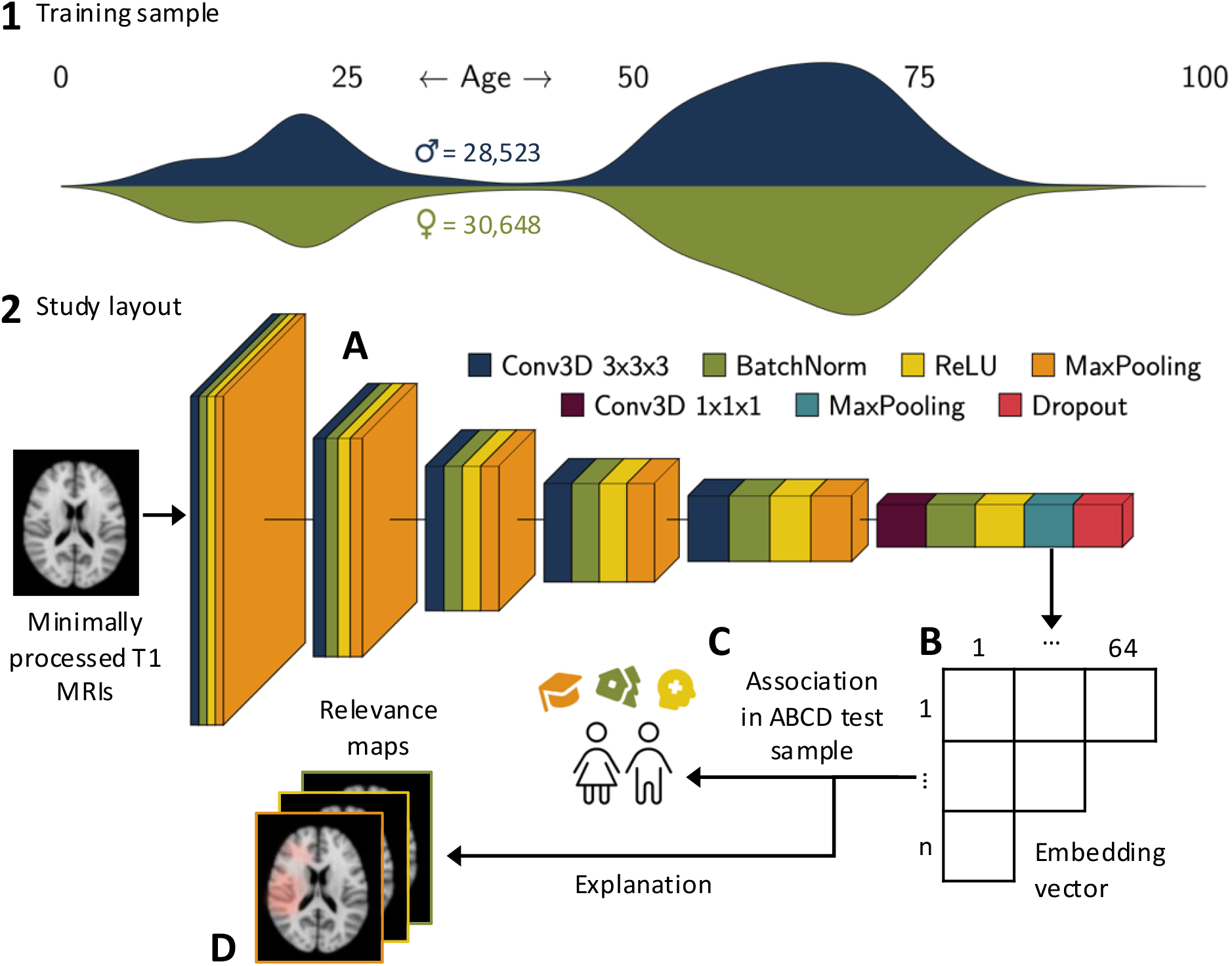
Age and sex distribution of the training sample (59,171 scans). 2A: The Simple Fully Convolutional Network architecture applied in the present study. An embedding vector is extracted from the final global max pooling layer. 2B: An n x 64 embedding matrix, where each column quantifies the extent to which the visual pattern associated with the embedding dimension is present in each MRI scan. 2C: Using linear models, each embedding dimension is linked to a large selection of traits in the hold-out ABCD test sample. 2D: For each individual, session, and embedding dimension, relevance maps are extracted for 1000 random participant using layer wise relevance propagation. An average map is used as a visual guide for which input features contributed to each dimension.

The model was trained to simultaneously predict age, sex, handedness, BMI, fluid intelligence and neuroticism. The encoder of the model consisted of five repeated blocks of 3-dimensional convolutions, batch normalization, a rectified linear unit (ReLU) activation and 3-dimensional max pooling layer, followed by a similar block designed to reduce the bottleneck dimensionality to a predefined 64 dimensions. Applying this encoder to an image produces an embedding vector that summarizes how strongly each learned high-level visual pattern is expressed in the individual brain scan. As four dimensions consistently yielded zero values across all input images, the effective dimensionality of the learned latent space was reduced from 64 to 60 dimensions. Each dimension was treated as an independent variable in the subsequent statistical analysis. As a visual guide for understanding the anatomical basis of the dimensions, we constructed relevance maps for each image for each dimension using layer-wise relevance propagation (LRP)^14^. The relevance maps indicate regions of the brain that most strongly contribute to an individual’s value for a dimension from a given timepoint (baseline or two-year follow-up)^15^.

To obtain a broad understanding of how the derived dimensions associate with the adolescent phenome, we investigated the association between each dimension in the embedding space and 38 variables, of which 26 had partially or fully available follow-up data (Supplementary Table 2). The included phenotypes represented three disjoint domains. A *personal* domain, including internal states, abilities and physical characteristics, a *relational* domain, capturing aspects of the adolescent’s social environment, and a *socioenvironmental* domain, reflecting characteristics of the adolescent’s immediate surroundings.

First, we assessed the intercorrelation between dimensions, the changes over time, and the between-subject correlation between baseline and the 2-year follow-up. For the main analysis, we employed a mass univariate approach for both sessions, where each dimension was linked to each outcome using linear regression, extracting the marginal effects from models with sex and age as nuisance variables. We also employed longitudinal linear mixed models, specifically focusing on the variance explained uniquely by the interaction between dimensions and time. As a basis for comparison with the extracted dimensions, we obtained measurements of cortical thickness, surface area, and volume for the four major brain lobes, and repeated the same analyses with surface metrics as predictors.

Although the focus of the modelling process was to create a diverse feature space, age (MAE = 2.49; Fig. 2B) and sex (AUC = 0.83) was predicted accurately in the test set (Extended Data Figure 3). Handedness (AUC = 0.52, BMI (MAE = 3.21) and fluid intelligence (*r* = 0.10) showed poor predictability. Fig. 2C shows that a diverse feature space was achieved, with an average absolute intercorrelation between embedding dimensions of *r* = 0.25 in the test set at baseline, substantially lower than previous single-task models^12^. The mean between-subject correlation between dimensions representing baseline and 2-year follow-up dimensions was *r* = 0.43 (range 0.25-0.62, Fig. 2D), and the mean intraclass correlation was 0.39 (range 0.24-0.57), indicating fair between- and within-subject stability across time, albeit lower than the corresponding *r* and ICC for surfaced-based metrics in the same participants (Extended Data Figure 4 and Supplementary Table 4 and 5). Paired-sampled t-tests revealed strong developmental changes in the embedding weights with significant differences in observed in 58 dimensions (t-scores ranging from −45.8 to 42.0, Fig. 2E, Supplemental Table 5).

**Fig. 2.**
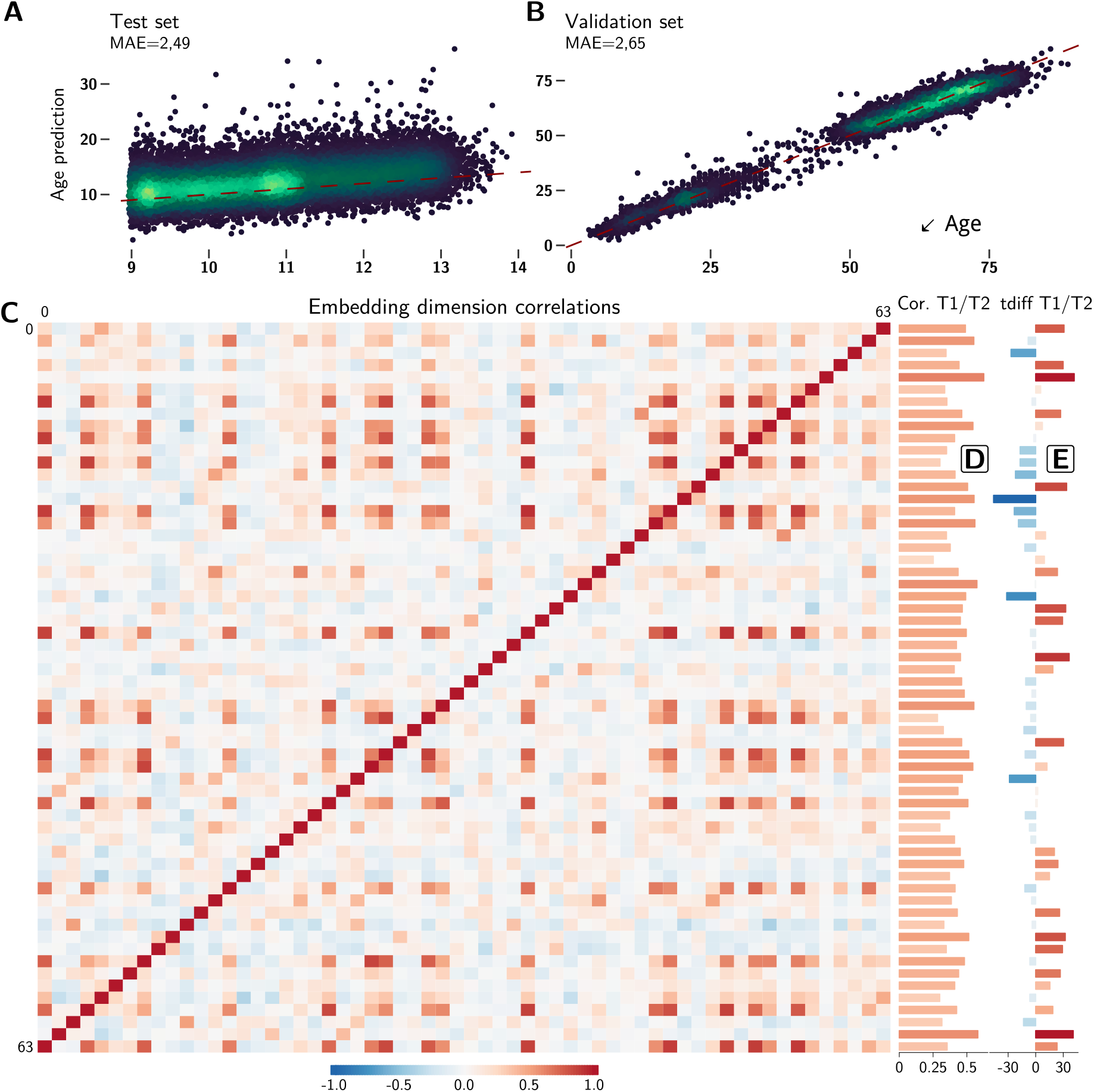
A: Age prediction MAE for the test set (both sessions) and the validation set (B). Dashed red lines indicates the identity line. C: Average intercorrelation of the 60 embedding dimensions in the test fold at baseline (zero activation embeddings 28, 51, 57 and 59 removed). D: Between-subject correlations of test set embedding dimensions from baseline to the two-year follow-up. E: T-values from paired-samples t-tests testing the difference in dimension weights between baseline and 2-year follow-up.

The main analysis revealed that embedding dimensions captured phenotypic variance across personal, social and socioenvironmental domains (Fig. 3A). As expected, given that BMI and fluid intelligence represented two of the six predictive targets in the model training, most dimensions linked significantly to anthropometrics and cognition at baseline and follow-up (Extended Data Figure 5). However, multiple dimensions also significantly traced phenotypes not directly familiar to the model, including digital device use, both parents’ age at conception, and local area deprivation. Some dimensions were distinctively associated with smaller clusters of related phenotypes (Fig. 3D). Longitudinal analyses revealed that the associations were largely stable across time, with dimension by time interactions emerging only for anthropometrics and parental involvement in the home and school environment, characteristics that change significantly in the transition to early adolescence (Fig. 3B). Comparing the embedding dimensions and surface metric with the largest correlation with each trait revealed that surface metrics more closely linked to cognition, and embedding dimensions to socioenvironmental variables (Fig. 3C).

**Fig. 3.**
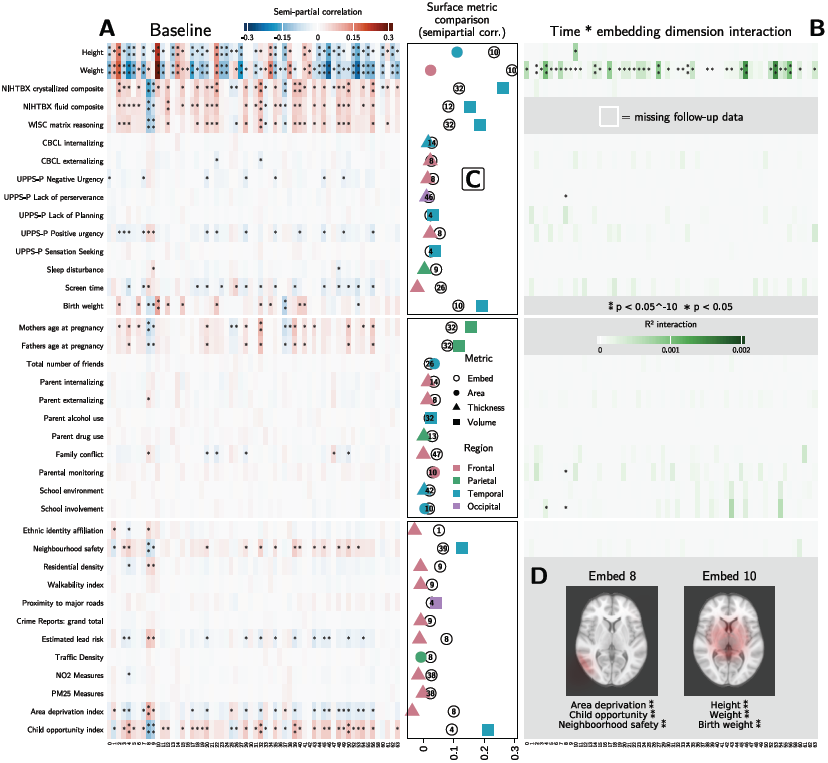
A: Semi-partial correlations between embedding dimensions and predictors from a linear model with age and sex included. B: The partial explained variance of the embedding dimension by time interaction of linear mixed models. C: For each trait, a comparison of the dimension and the surface metric with the highest semi-partial correlation. D: Relevance maps for two selected dimensions.

These findings demonstrate that a multitask CNN trained on minimally processed MRI scans capture distinct and trait-relevant brain morphological characteristics. The associations extended beyond the trained targets and encompassed clusters of social and socioenvironmental traits, most prominently composite markers of neighborhood deprivation and opportunity. Associations between the developing brain and neighborhood characteristics are well established^18^, likely attributable to constellations of material and social conditions. For area deprivation, the finding that the most strongly correlated dimension was considerably more sensitive than the strongest surface measure indicates that cumulative neighborhood effects are better captured by distributed structural configurations than by mean differences within cortical regions.

Mapping brain–trait associations during key transitions such as adolescence provides a window into how distributed brain morphology encodes emerging patterns of adaptation across behavioral domains. We have developed a framework for deriving a set of embedding vectors with a dimensionality comparable to common brain atlases, in the present case applied to a child and adolescent cohort. Our approach may be further enhanced by focally investigating the importance of design choices such as the size of the latent space (e.g. varying dimensionality), modifying targets during model training, combining different MRI modalities, and a further exploration of explainability methods^13,19^. Doing so may establish a method that complements surface- and volume-based approaches by providing flexible, data-driven representations that are sensitive to neurodevelopment and refine our understanding of how adolescent experiences become embedded in brain structure and how complex, bidirectional associations between the adolescent phenome and the brain unfold.

## Methods

### Data - Training and validation

For the present study, we trained and validated an Simple Fully Convolutional Network (SFCN) multi-task model^20^, on a set of 65,290 MRI scans from 59,454 healthy participants. Attributions and ethical approvals for each included source is given in Supplementary Table 1. The training fold consisted of 59,171 scans from 53 335 individuals, stratified by source, scanner ID, sex and age. The average age of the training fold was 53.4 (± one standard deviation of 21.3 years; Extended Data Figure 2A) The age range was 3.0 to 97.4 years and 51.8% were female.

The validation fold consisted of 6119 scans from 6119 individuals. When a participant was randomly assigned to the validation fold, all scans belonging to that person were removed from the training fold, ensuring no overlap between folds. Additionally, only one randomly selected scan for each person was retained in the validation fold. The age and sex distribution was constructed to approximate that of the training fold (average age=55.1±20.2 years, range 3.2 to 89.7 years, 51.9% females; Extended Data Figure 2B).

For all folds, we obtained, when accessible, data on age, sex, handedness, BMI, fluid intelligence and neuroticism, constituting the six predictive targets of the model. Age and BMI were included as continuous variables without any modification. Sex and handedness were coded as binary variables (female = 0, male = 1; right-handed = 0, left-handed = 1), with mixed/ambidextrous handedness set to missing. Measures of fluid intelligence and neuroticism were collected from different tests and questionnaires, before z-score standardization within each source to create a unified target measure across all sources. The training labels for the fluid intelligence prediction largely consisted of the UKBB Touch-screen Fluid intelligence test (n=30,106)^21^, and the training labels for the neuroticism prediction consisted primarily of the UKBB Derived Neuroticism score (n=27,543)^22^.

### Hold-out test data

The final full held-out test data consisted of 18,813 images of 11,497 individuals from the ABCD study, obtained from the ABCD Fast-track Imaging Data Release on the 18^th^ of March 2025. Of these images, 11,103 were from the baseline session (mean age = 9.9 ± 0.6 years), and 7710 were from the 2-year follow-up (mean age = 11.9 ± 0.6 years). A full breakdown of key participant characteristics at baseline is given for the test data in Extended Data Figure 1, as well as a comparison on key characteristics between included participants and participants excluded due to missing data or failed imaging quality control (QC; see below). Overall, we noted no signs of bias in inclusion, except for a minor trend towards excluded participants having more highly educated caregivers and having a somewhat higher household income.

The initial downloaded material consisted of 25,189 images from 11,781 individuals. Of these, 4806 images were removed due to not having any available QC data, failing QC, or having a later run from the same session passing QC. A further 1466 images were removed as they were from the 4-year follow-up session. We considered this subsample too small to include in the present study. Lastly, 50 images were removed as they lacked scanner information, and 54 images were removed due to not having matching surface-based data for that session (see *statistical analysis*).

All other data for the test fold was obtained from ABCD Curated Data Release 6.0 (doi.org/10.82525/jy7n-g441). The review and approval of the ABCD research protocol was handled by a central Institutional Review Board at the University of California, San Diego^23^. Informed consent was given by parents or guardians, and assent was given by children before participation. Our access to ABCD research data is given by Data Use Certification 22719. The current project is approved by the Norwegian Regional Committee for Medical and Health Research Ethics (REC; #2019/943).

### Image preprocessing and quality control

All images were preprocessed with FastSurfer 2.0.1^24^, using a previously described minimal preprocessing approach^12^. This consisted of aligning images to RAS (Right, Anterior, Superior) orientation and creating a binary brainmask. The RAS-aligned images were then multiplied with the binary brainmask, ensuring that only brain tissue is retained in the final image. Lastly, to reduce compute demands, all images were cropped to a 224x192x224 cube.

For all images in the training- and validation folds, we relied on the brief volume-based QC check included in FastSurfer, which identifies gross segmentation errors by comparing the total segmentation volume against a set threshold. Images with a total segmentation volume below this threshold were discarded from further analysis.

For the ABCD test data, we relied on a stepwise approach based on raw QC findings provided by ABCD (available in the ABCD 6.0 table *mr_y_qc_raw_smri_t1*). First, we checked whether a participant for a given session had any images passing the following three criteria:

- Data center received correct number of files (*mr_y_qc_raw_smri_t1_r0*_comp_indicator*).
- Scans passed protocol compliance check (*mr_y_qc_raw_smri_t1_r0*_protcom_indicator*).
- Scan passed manual quality control (*mr_y_qc_raw_smri_t1_r0*_indicator*).

If a participant for a given session had multiple runs passing all three criteria, we selected the last available run.

Kraft et al.^19^ demonstrated significant scanner effects in the ABCD study in atlas-based estimates of morphology. Thus, we employed the RELIEF method^13^ on all MRI data, which mitigates inter-scanner bias by partitioning and removing both scanner-specific means and latent scanner-specific variations (Supplementary Figure 1).

### Model architecture

Full details of the model architecture and training strategy are given in Leonardsen et al.^12^ Briefly, our approach is based on the SFCN-architecture by Gong et al.^14^ and Peng et al.^26^, consisting of five repeated blocks made up of a 3-dimensional convolution (3x3x3 filters), batch normalization, a rectified linear unit (ReLU) activation, and a 3-dimensional max pooling layer (2x2x2 patches). This is followed by a down-sampling block consisting of a 3-dimensional convolution (1x1x1 filters), batch normalization and a ReLU-activation, which is again followed by a global max pooling (GMP) layer. The final prediction head consists of six output neurons corresponding to the six predictive tasks, four of which have a linear activation (age, BMI, fluid intelligence, and neuroticism), and two having a sigmoid activation, allowing binary predictions (sex and handedness). During training, model loss was calculated as a weighted average across all six outputs. We fit a single model instance with a weight decay of 10^−3^, based on results from Leonardsen et al. ^27^. The model was fit with an Adam optimizer and an initial learning rate of 10^−3^, with a step-based reduction by a factor of 10 at epochs 30 and 50. Final model weights were obtained from the epoch with the lowest overall loss, out of a maximum of 60 epochs. As loss had to be averaged across six predictive targets of different scales, we employed a custom weighting scheme ensuring that the six losses contributed approximately equally to the total composite loss^12^.

### Model evaluation

Model performance was evaluated by calculating performance metrics for each predictive target in both the validation fold and the ABCD test fold, except for neuroticism, for which we could not identify suitable data in the test set. Predictions for age and BMI were evaluated by calculating the mean absolute error (MAE) between the predicted and true scores. Binary sex- and handedness-predictions were evaluated by calculating the area under the receiver operating characteristic curve (AUC). For fluid intelligence, we calculated the correlation *r* between the predicted and true scores.

### Embedding extraction

In the second to last layer of the model, each image is represented by a 64-dimensional embedding vector. Each dimension represents the maximum value for each embedded map across the three spatial dimensions (height, width and depth). To illustrate, consider a single embedding map *M*(*i, j, k*) with dimensions W × H × D where i ∈ [1, *W*], j ∈ [1, *H*], and *k* ∈ [1, *D*]. The global max for this embedding map is max_i,j,k_ M(i, j, k), reducing the dimensionality of the image from 3D to 1D. Each dimension represents a high-level feature of the brain, and each image is encoded as a point in this space. As four dimensions consistently yielded zero values across all input images, the dimensionality of the learned latent space was reduced to 60, with zero activation embeddings 28, 51, 57 and 59 removed.

### Embedding dimension relevance maps

To better understand and visualize the anatomical basis of each embedding, we constructed relevance maps using layer-wise relevance propagation (LRP)^29^, based on the approach outlined in Leonardsen et al.^30^. In their simplest form, the relevance maps indicate regions of the brain that most strongly contribute to a specific embedding value for a given input image. To obtain the relevance map *R* for an image *X* corresponding to a given embedding value *F*, the relevance is propagated backwards through the network from the GMP-layer to the input layer. *R* is a three-dimensional volume where each voxel *r*_*i,j,k*_ ∈ *R* with positions *i, j, k* corresponds to the position of an input voxel *x*_*i,j,k*_ ∈ *X*, meaning that the spatial relation between the input and the relevance map is preserved. With the assumption that some embeddings might not be strongly relevant for every input, we used the ‘epsilon rule’ described in Bach et al. (2015) for all layers, which avoids issues of dividing by very small numbers (e.g. neurons that are near zero and irrelevant for the explanation). The relevance *R* for a single neuron *n* in a single layer is given as:

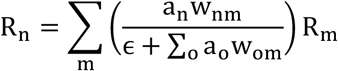

Where a_n_ is the activation of neuron *n, w*_*nm*._ is the weight of the connection between *n* and another neuron *m* in the subsequent layer, ∑_*o*_ *a*_*o*_ *w*_*om*._ is the sum of the influence of all neurons connected to *m*, and *R*. is the relevance propagation through neuron *m*.

For each image, the process outlined above creates one relevance map for each embedding. To obtain average maps per embedding, we performed the following three-step process. First, each input image was non-linearly transformed to the 1 mm MNI152 template using FNIRT from FSL version 6.0.3 ^13^. Next, each relevance map was mapped to their corresponding image input by applying the transformation matrix from the MNI-152 transformation to the relevance map for that participant. Lastly, in MNI space we obtained the average relevance for each embedding by calculating the mean of all relevance maps for that embedding. The average relevance maps were visualized by superimposing them on the MNI152 template, using the slice where the highest absolute relevance was found (Supplementary Figure 2). Before visualization, each relevance map was divided by the highest absolute relevance of that relevance map.

### Outcome measures

An exhaustive list of ABCD outcome variables is given in Supplementary Table 2. For consistency in our later modelling approach, we only included outcomes that we considered it appropriate to treat as continuous. This included ratio-, interval- and ordinal scale variables with at least ten possible unique values. For outlier removal, we adopted a relaxed approach and removed only scores for each variable that were beyond four absolute deviations from the median^14^. Upon further manual review, we additionally set body weight below 40 lbs. or above 350 lbs., as well as standing height below 30″ or above 80”, to missing, with the assumption that these values constituted data entry errors.

## Statistical analysis

We first investigated the model space and calculated the pair-wise mean absolute Pearson correlation coefficient between all embedding dimensions in the test fold, using data from the baseline session. To determine their relative between and within-subject stability across age, for each dimension we additionally calculated the Pearson correlation coefficient, the intraclass correlation (ICC), and performed paired-sample t-tests between the baseline session and the two-year follow-up.

For the cross-sectional part of the main analysis, for each session we fit 60 separate linear models for each outcome, with each model using a different dimension as the predictor variable. All models also included age and sex as covariates. The linear formula for each model can be expressed as:

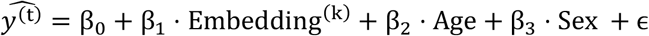

where ŷ_*l*_ is the estimated value for the outcome t, where *t* ∈ 1, …, 38 at baseline and *t* ∈ 1, …, 26 at the two-year follow-up, and Embedding_*k*_ represents one of the 60 embedding dimensions (k = 1, …, 60).

From here, for each model we extracted the semipartial correlation between the dimension and the outcome variable, which for a single model can be written as

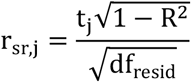

Where *t* is the t-value of predictor j, *R*^2^ is for the full model and df_resid_ is *n* − *p* − 1, with *p* being the number of predictors in the model. Reported p-values are obtained from the t statistic of the embedding dimension predictor. All p-values were false discovery rate adjusted for the total number of models for each part of the analysis (n = 2280 for the baseline cross-sectional analysis and n = 1560 for the two-year follow-up and longitudinal analyses).

For the longitudinal part of the main analysis, we fit 60 separate linear mixed models for each outcome, with each model using a different embedding as the predictor variable.

Additionally, each model included time (baseline session or the two-year follow-up), age, sex, and a time by embedding dimension interaction term. The full models can be written as:

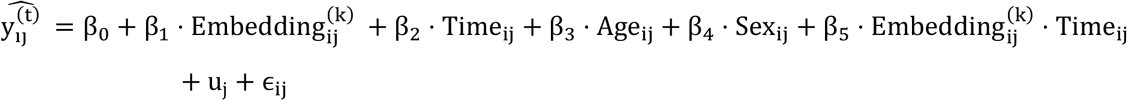

where 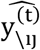 is the estimated value for the *i*-th observation of participant *j* for outcome *t*, where *t* ∈ 1, …, 26, Embedding_*k*_ represents one of the 60 embedding dimensions k, and u_j_ is the random intercept of participant *j*, with a mean of 0 and a variance of 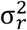.

From here, for each model we extracted the semipartial *R*^2^ of the interaction term, which amounts to the variance explained uniquely by the interaction between each dimension and the session ID (baseline or the two-year follow-up). This procedure is described in detail in Edwards et al.^15^. P-values were obtained using Wald F-tests of the fixed effect of the interaction term, implemented in the R package *r2glmm* (10.32614/CRAN.package.r2glmm).

To establish a basis for comparison with the extracted embedding dimensions, we collected measurements of cortical thickness, surface area, and volume for the frontal, occipital, temporal, and parietal lobes for both hemispheres. Measures of cortical morphology were generated by the ABCD consortium using a fuzzy clustering approach developed by Chen et al.^16^, provided by the ABCD consortium and found in the ABCD 6.0 tables *mr_y_smri_thk_fzy, mr_y_smri_vol_fzy*, and *mr_y_smri_area_fzy*. We applied the same RELIEF harmonization to this data, to keep the two approaches as similar as possible. In the final analysis, each cortical measure served as a predictor within the same model framework as previously described, substituting the Embedding_*k*_term in the linear equation.

To evaluate which approach accounted for the greatest amount of variation in the included outcomes, we conducted a comparison of embedding dimensions and cortical morphology measures for each outcome, comparing the embedding dimension with the highest semipartial correlation with the outcome, and the cortical measure with the highest semipartial correlation with the outcome. A descriptive comparison is given in Figure 3C (baseline) and Extended Data Figure 5 (two-year follow-up), and model comparisons are given in Supplementary Table 6.

## Acknowledgements

The data used in this study was compiled from several sources, each with their own acknowledgements that can be found in Supplementary Table 1

The work was funded by the Norwegian Research Council (grant numbers 324499 and 300767), NordForsk (grant number 164218), and the European Research Council (grant number 802998). All analyses were performed on the Services for Sensitive Data (TSD), University of Oslo, Norway, with resources from UNINETT Sigma2, the National Infrastructure for High-Performance Computing and Data Storage in Norway.

## Data availability

The data incorporated in this study was compiled from a variety of sources. Requests for the underlying data need to be placed with the principal investigators for each individual study. An overview of these data sources and their origins can be found in Supplementary Table 1.

## Code availability

The code and weights for the pretrained model and various interfaces for interacting with it is available on https://github.com/estenhl/pyment-public,

