## Supplementary Material for "Latent neural network representations of the brain reflect broad-scale adolescent phenotypic variation"

Extended Data Figure 1: ABCD test fold demographics at baseline

| | | Excluded (N=765) | Included (N=11,103) | Eff.size difference |
|---|---|---|---|---|
| **Age** | | | | 0.117[a] |
| | Mean (SD) | 9.9 (0.6) | 10.0 (0.6) | |
| | Median (IQR) | 9.9 (1.1) | 9.9 (1.1) | |
| | Range | 9.0 - 11.0 | 8.3 - 11.3 | |
| **Highest caregiver education** | | | | 0.097[b] |
| | Up to high school (No diploma) | 62 (8.1%) | 531 (4.8%) | |
| | High school diploma/GED | 97 (12.7%) | 1,035 (9.3%) | |
| | Some college | 242 (31.6%) | 2,832 (25.5%) | |
| | Bachelor's degree | 173 (22.6%) | 2,840 (25.6%) | |
| | Graduate school or professional degree | 190 (24.8%) | 3,852 (34.7%) | |
| | Missing | 1 (0.1%) | 13 (0.1%) | |
| **Yearly household income** | | | | 0.066[b] |
| | < 25k | 146 (19.1%) | 1,488 (13.4%) | |
| | 25k to 50k | 127 (16.6%) | 1,461 (13.2%) | |
| | 50k to 75k | 102 (13.3%) | 1,396 (12.6%) | |
| | 75k to 100k | 86 (11.2%) | 1,484 (13.4%) | |
| | 100k to 200k | 167 (21.8%) | 3,144 (28.3%) | |
| | > 200k | 53 (6.9%) | 1,197 (10.8%) | |
| | Don't know | 42 (5.5%) | 469 (4.2%) | |
| | Decline to answer | 42 (5.5%) | 462 (4.2%) | |
| | Missing | | 2 (0.0%) | |
| **Race / Ethnicity** | | | | 0.007[b] |
| | Hispanic | 172 (22.5%) | 2,273 (20.5%) | |
| | White | 287 (37.5%) | 5,898 (53.1%) | |
| | Black | 208 (27.2%) | 1,604 (14.4%) | |
| | Asian | 13 (1.7%) | 216 (1.9%) | |
| | Other | 85 (11.1%) | 1,110 (10.0%) | |
| | Missing | | 2 (0.0%) | |
| **Sex** | | | | 0.07[b] |
| | Male | 409 (53.5%) | 5,781 (52.1%) | |
| | Female | 356 (46.5%) | 5,322 (47.9%) | |

[a] Cohen's d, unequal variances assumed

[b] Cramer's V (df=1)

**Extended Data Figure 1:** Comparison of included and excluded participants on key demographics. Age ABCD name = *ab_p_demo_age*, education = *ab_g_dyn__cohort_edu__cgs*, income = *ab_g_dyn__cohort_income__hhold__6lvl*, race / ethnicity = *ab_g_stc__cohort_ethnrace__leg*, sex = *ab_g_stc__cohort_sex*.

Extended Data Figure 2: Age and sex distributions of the training and validation fold

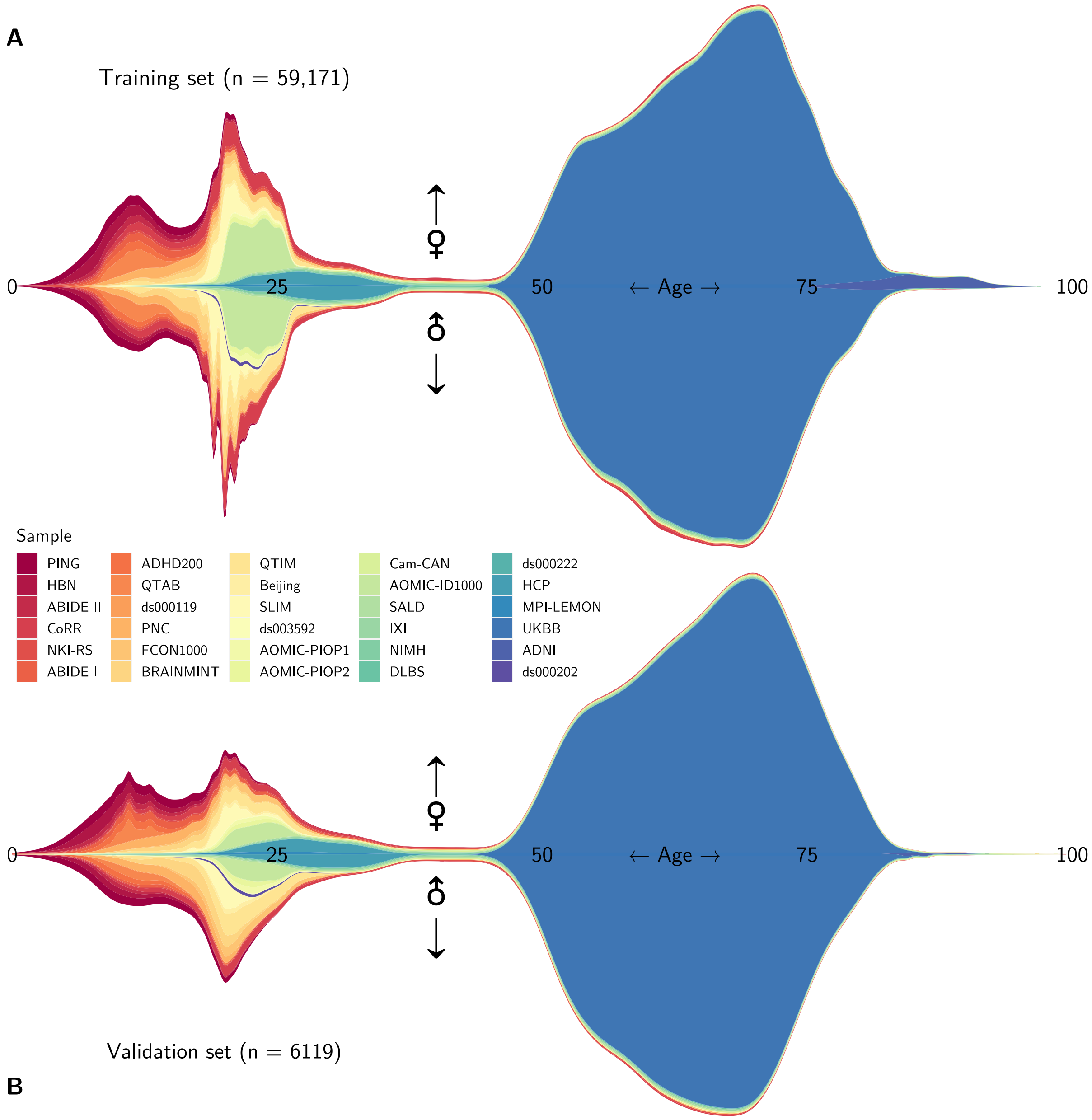

**Extended Data Figure 2:** A: Age and sex distribution of the training fold B: Age and sex distribution of the validation fold.

Extended Data Figure 3: Model training summary.

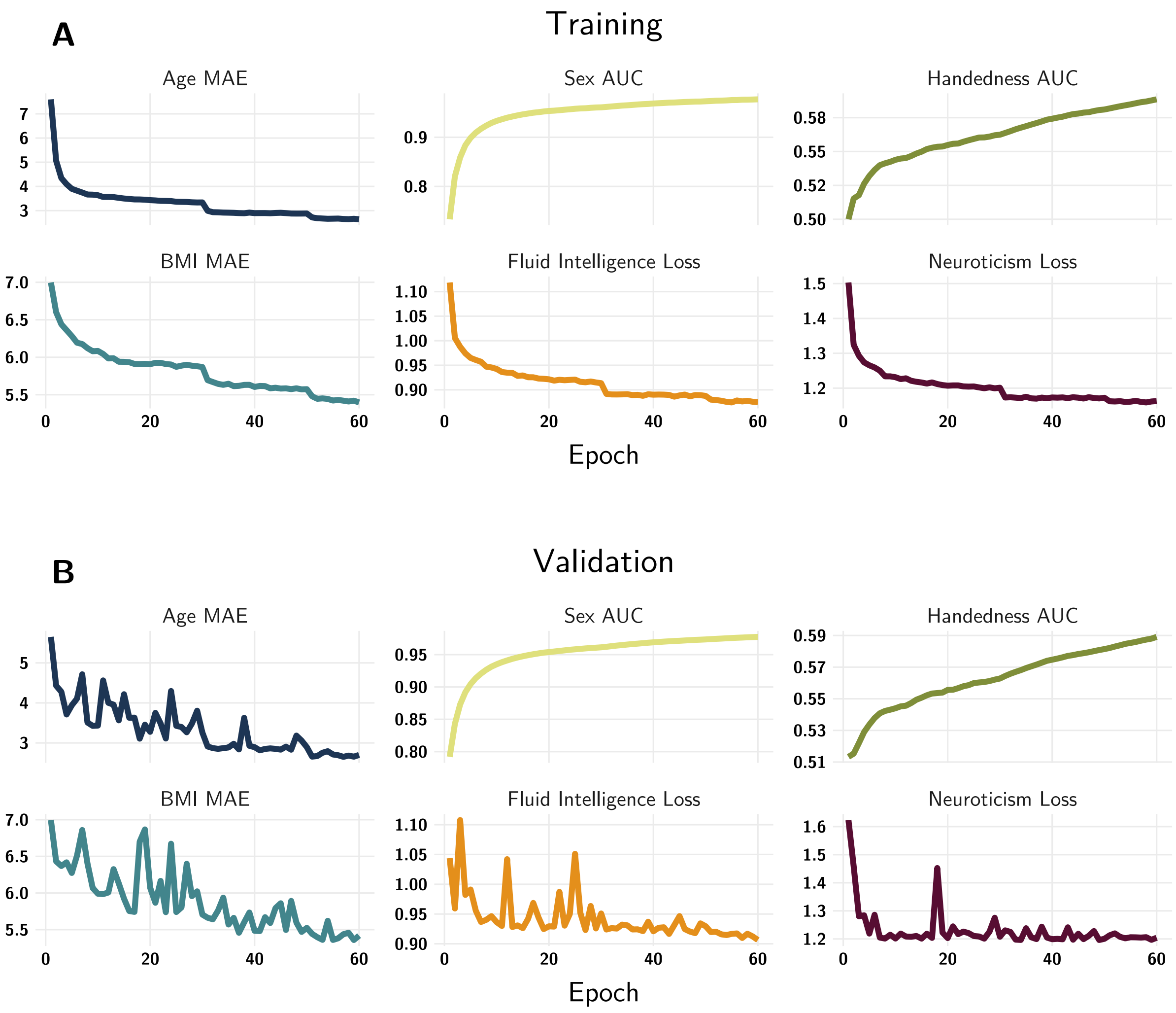

**C**

| | Age MAE | Sex AUC | Handedness AUC | BMI MAE | Neuroticism $r$ | Fluid Intelligence $r$ |
|---|---|---|---|---|---|---|
| Validation fold | 2.65 | 0.98 | 0.59 | 5.44 | 0.17 | 0.24 |
| Test fold | 2.58 | 0.83 | 0.52 | 3.21 | NA | 0.10 |

**Extended Data Figure 3:** A: Training and (B) validation performance across the different multi-task targets. Note that the Y-axis is not consistent across targets. C: Summary table of performance across targets in the validation fold and test fold, where $r$ is the correlation between the actual and the predicted values. Data on neuroticism was unavailable in the test set.

Extended Data Figure 4: Embedding dimension correlations between baseline and two-year follow-up

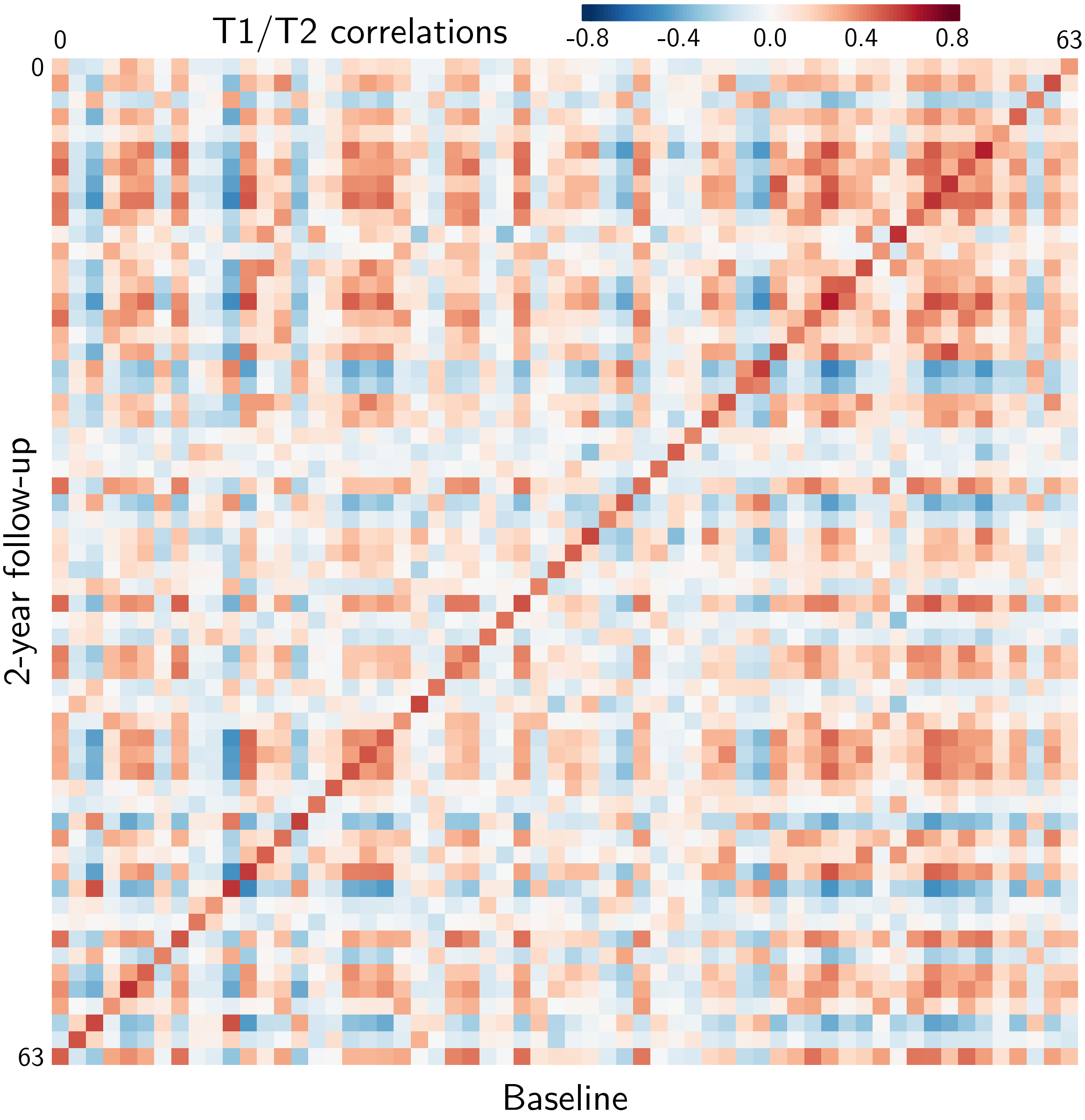

**Extended Data Figure 4:** Matrix of correlations between the weights of the embedding dimensions at baseline and at the two-year follow-up.

Extended Data Figure 5: Two-year follow-up results.

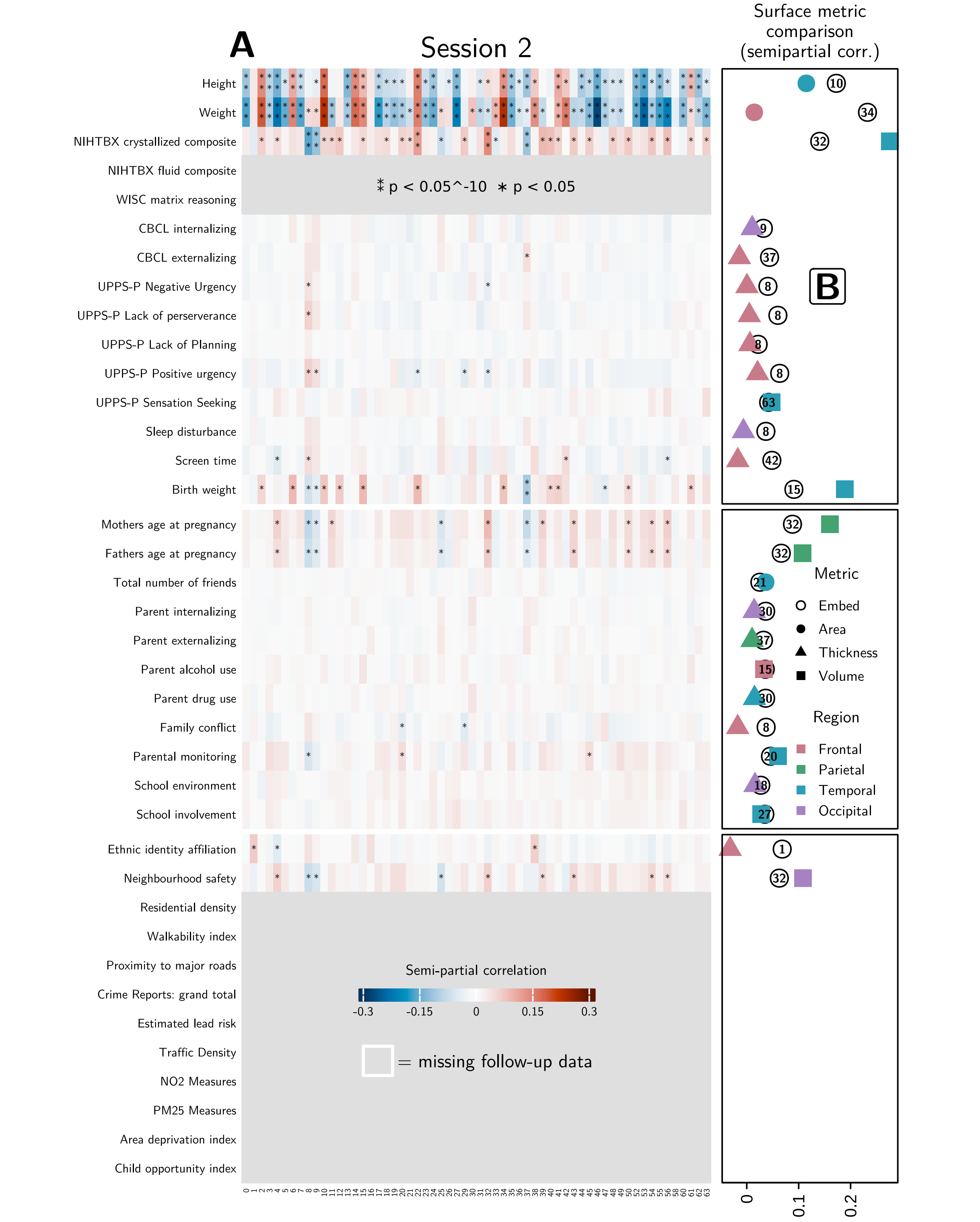

**Extended Data Figure 5:** Outcomes of cross-sectional analysis for the two-year follow-up session. A: Semi-partial correlations between embedding dimensions and outcomes, from linear models with age and sex as covariates. B: For each trait, a comparison of the embedding dimension and the surface metric with the highest semi-partial correlation.

Supplementary Table 1: Origins and acknowledgements for the data sources used in the study

| Name | Shorthand | Source | Acknowledgements | References |
| --- | --- | --- | --- | --- |
| Adolescent Brain Cognitive Development | ABCD | https://abcdstudy.org/ | Data used in the preparation of this article were obtained from the Adolescent Brain Cognitive DevelopmentSM (ABCD) Study (https://abcdstudy.org), held in the NIMH Data Archive (NDA). This is a multisite, longitudinal study designed to recruit more than 10,000 children age 9–10 and follow them over 10 years into early adulthood. The ABCD Study is supported by the National Institutes of Health and additional federal partners under award numbers U01DA041048, U01DA050989, U01DA051016, U01DA041022, U01DA051018, U01DA051037, U01DA050987, U01DA041174, U01DA041106, U01DA041117, U01DA041028, U01DA041134, U01DA050988, U01DA051039, U01DA041156, U01DA041025, U01DA041120, U01DA051038, U01DA041148, U01DA041093, U01DA041089, U24DA041123, U24DA041147. A full list of supporters is available at https://abcdstudy.org/federal-partners.html. A listing of participating sites and a complete listing of the study investigators can be found at https://abcdstudy.org/consortium_members/. ABCD consortium investigators designed and implemented the study and/or provided data but did not necessarily participate in the analysis or writing of this report. This manuscript reflects the views of the authors and may not reflect the opinions or views of the NIH or ABCD consortium investigators. Our access to ABCD research data is given by Data Use Certification 22719. | Casey et al., 2018 |
| Autism Brain Imaging Data Exchange I | ABIDE-I | https://fcon_1000.projects.nitrc.org/indi/abide/abide_I.html | Primary support for the work by Adriana Di Martino was provided by the NIMH (K23MH087770) and the Leon Levy Foundation. Primary support for the work by Michael P. Milham and the INDI team was provided by gifts from Joseph P. Healy and the Stavros Niarchos Foundation to the Child Mind Institute, as well as by an NIMH award to MPM (R03MH096321). | Di Martino et al., 2014 |
| Autism Brain Imaging Data Exchange II | ABIDE-II | https://fcon_1000.projects.nitrc.org/indi/abide/abide_II.html | Primary support for the work by Adriana Di Martino and her team was provided by the National Institute of Mental Health (NIMH 5R21MH107045). Primary support for the work by Michael P. Milham and his team was provided by the National Institute of Mental Health (NIMH 5R21MH107045); Nathan S. Kline Institute of Psychiatric Research. Additional support was provided by gifts from Joseph P. Healey, Phyllis Green and Randolph Cowen to the Child Mind Institute. | Di Martino et al., 2017 |
| ADHD-200 | ADHD200 | https://fcon_1000.projects.nitrc.org/indi/adhd200/ | F. Xavier Castellanos, David Kennedy, Michael Milham, and Stewart Mostofsky are acknowledged for their roles in the ADHD-200 project (text truncated in original extraction; see original source for full acknowledgements). | Brown et al., 2012; Milham et al., 2012 |

| | | | | |
|---|---|---|---|---|
| Beijing Normal University Enhanced Sample | Beijing | https://fcon_1000.projects.nitrc.org/indi/retro/BeijingEnhanced.html | Financial support for the data used in this project was provided by a grant from the National Natural Science Foundation of China: 30770594 and a grant from the National High Technology Program of China (863): 2008AA02Z405. | Tian et al., 2011; Yan and Zang, 2010 |
| Brain and minds in transition | BRAINMINT | https://www.sv.uio.no/psi/english/research/projects/brainmint/ | Accessed with approval from the Regional Committee for Medical Research Ethics South East Norway (REC, application number: 2019/943). Funded by the European Union (ERC, BRAINMINT, 802998). | |
| Cambridge Centre for Ageing and Neuroscience dataset | Cam-CAN | https://cam-can.mrc-cbu.cam.ac.uk/dataset/ | Data used in the preparation of this work were obtained from the CamCAN repository (available at http://www.mrc-cbu.cam.ac.uk/datasets/camcan/). Data collection and sharing for this project were provided by the Cambridge Centre for Ageing and Neuroscience (CamCAN). CamCAN funding was provided by the UK Biotechnology and Biological Sciences Research Council (grant number BB/H008217/1), together with support from the UK Medical Research Council and University of Cambridge, UK. | Shafto et al., 2014; Taylor et al., 2017 |
| Consortium for Reliability and Reproducibility | CoRR | https://fcon_1000.projects.nitrc.org/indi/CoRR/html/index.html | The National Institute on Drug Abuse and the National Natural Science Foundation of China (NSFC) have been instrumental in the CoRR collaboration, providing the necessary funding and manpower to build the foundation of the project along with the Child Mind Institute, the Institute of Psychology, Chinese Academy of Sciences and the Nathan Kline Institute. | Zuo et al., 2014 |
| Dallas Lifespan Brain Study | DLBS | https://fcon_1000.projects.nitrc.org/indi/retro/dlbs.html | We would like to thank the following individuals and research bodies for their continuing support of the study: Our scientific and support staff, who conduct the day-to-day operations and provide the long-term support necessary to keep the study running: Paula Abercrombie, Bela Bhatia, Gerard Bischof, Ph.D., Micaela Chan, Xi Chen, Arielle Click, Mark Diaz-Arrastia, Aaron Dostson, Linda Dubose, Patrick Evans, Victor Faner, Michelle Farrell, Blair Flicker, Jacqueline Gauer, Cassandra Hatt, Andy Hebrank, Marci Horn, Richard Innis, Caroline Janeway, Debby Kirchhevel, Mitchell Meltzer, April Norambuena, Heekyeong Park, Ph.D., Allison Parker, Jenny Rieck, Ph.D., Melissa Rundle, Ph.D., Prasanna Tamil, Nicole Tehrani, Erin Wooden. The Center for Vital Longevity, the University of Texas at Dallas, and the University of Texas Southwestern Medical Center, for sponsoring and providing the support and facilities needed to conduct the study. The National Institutes of Health and Aging, for their continuing financial and scientific support. AVID Radiopharmaceuticals, for providing the ligand used in the PET imaging procedure. The Aging Mind Foundation and the Alzheimer's Association, for providing additional funding for this research. Finally, the Dallas Lifespan Brain Study could not have been accomplished without the help of our | H. Lu et al., 2011 |

| | | | | |
|---|---|---|---|---|
| | | | research participants. We appreciate their continued interest and participation in the study! | |
| 1000 Functional Connectomes | FCON1000 | https://fcon_1000.projects.nitrc.org/fcpClassic/FcpTable.html | Collected at 33 independent sites by J.J. Pekar, S.H. Mostofsky, S. Colcombe, Y.F. Zang, D. Marguiles, R.L. Buckner, M.J. Low, B. Rypma, D.J. Madden, A.C. Evans, S.A.R.B. Rombouts, A. Villringer, S.J. Li, C. Sorg, V. Riedel, B. Biswal, M. Hampson, M.P. Milham, F.X. Castellanos, P. Williamson, M. Hoptman, V.J. Kiviniemi, J. Veijola, S.M. Smith, C. Mackay, M. Greicius, G. Siegle, K. McMahon, B. Schlaggar, S. Petersen, C.P. Lin, H.S. Mayberg, C.S. Monk, R.D. Seidler, S.J. Peltier. | |
| Healthy Brain Network | HBN | https://data.healthybrainnetwork.org/ | We thank the Communications, Development, Finance, and Human Resource teams at the Child Mind Institute (past and present) for their endless support, as well as the CMI Executive Team and the Child Mind Institute Scientific Research Council for their guidance and critical feedback in the planning of the Healthy Brain Network; Judith Gardner and Bernard Karmel for assisting in the recruitment efforts; Tammy Vanderwaal and Uri Hasson for their consultation in the selection of movies for the natural viewing paradigms; Simon Kelly for his assistance with devising the EEG battery; Megan Horton for advising us to add the collection of baby teeth; Antonio Convit for information regarding assessments of body composition; Stan Colcombe for information regarding fitness assessments; and Michael Michaelides for advising us to add hair samples for metals. Additionally, we would like to thank Joan Kaufman and Ken Kobak for providing access to the newly developed computerized KSADS and Ted Satterthwaite for helpful comments on the manuscript during its preparation. We also acknowledge and thank Staten Island Borough President James Oddo, Staten Island Health and Wellness Director Dr. Ginny Mantello, and New York State Senator Andrew J. Lanza and his team (specifically William Matarazzo and Anthony Reinhart) for their guidance in developing strong partnerships throughout Staten Island, and their continued support of the project. We would also like to express our sincere gratitude to the mental health organizations, service providers, and clinicians across Staten Island, and NYC at large, who continue to work with our staff and refer participants to the project. Our sincere gratitude is extended to the participants and their families for their contributions to this project. The Healthy Brain Network and its collaborative initiatives are supported by philanthropic contributions from the following individuals, foundations and organizations: Margaret Bilotti; Brooklyn Nets; Agapi and Bruce Burkard; James Chang; Phyllis Green and Randolph Cowen; Grieve Family Fund; Susan Miller and Byron Grote; Sarah and Geo Gund; George Hall; Jonathan M. Harris Family Foundation; Joseph P. Healey; The Hearst Foundations; Eve and Ross Jaffe; Howard & Irene Levine Family Foundation; Rachael and | Alexander et al., 2017 |

| | | | | |
|---|---|---|---|---|
| | | | Marshall Levine; George and Nitzia Logothetis; Christine and Richard Mack; Julie Minsko; Valerie Mnuchin; Morgan Stanley Foundation; Amy and John Phelan; Roberts Family Foundation; Jim and Linda Robinson Foundation, Inc.; Linda and Richard Schaps; Zibby Schwarzman; Abigail Pogrebin and David Shapiro; Stavros Niarchos Foundation; Preethi Krishna and Ram Sundaram; Amy and John Weinberg; Donors to the 2013 Child Advocacy Award Dinner Auction; Donors to the 2012 Brant Art Auction. | |
| Human Connectome Project | HCP | https://www.humanconnectome.org/ | Data were provided in part by the Human Connectome Project, MGH-USC Consortium (Principal Investigators: Bruce R. Rosen, Arthur W. Toga and Van Wedeen; U01MH093765) funded by the NIH Blueprint Initiative for Neuroscience Research grant; the National Institutes of Health grant P41EB015896; and the Instrumentation Grants S10RR023043, 1S10RR023401, 1S10RR019307. | Van Essen et al., 2013 |
| Amsterdam Open MRI Collection ID1000 | ID1000 | https://openneuro.org/datasets/ds003097/versions/1.2.1 | We thank all research assistants and students who helped collecting the data of the three projects, Jasper Wijnen and Marco Teunisse for advice and guidance with respect to anonymization and GDPR-related concerns, and Jos Bloemers, Sennay Ghebeab, Adriaan Tuiten, Joram van Driel, Christian Olivers, Ilja Sligte, Sara Jahfari, Guido van Wingen, and Suzanne Oosterwijk for help with designing the paradigms, Marcus Spaan for technical support, and Franklin Feingold and Joe Wexler for help with uploading the datasets to OpenNeuro. | Snoek et al., 2021 |
| Information eXtraction from Images | IXI | https://brain-development.org/ixi-dataset/ | | (no explicit citation given in extracted text) |
| Max Planck Institut Leipzig Mind-Brain-Body Dataset | MPI-LEMON | https://fcon_1000.projects.nitrc.org/indi/retro/MPI_LEMON.html | We thank all participants who volunteered to participate in our study. Moreover, we thank Elizabeth Kelly for proofreading the manuscript and Heike Schmidt-Duderstedt for editing tables and figures. | Babayan et al., 2019; Mendes et al., 2019 |
| National Institute of Mental Health Healthy Research Volunteer dataset | NIMH | https://openneuro.org/datasets/ds005752/versions/2.1.0 | We thank the NIMH Office of the Clinical Director, the outpatient behavioral health clinic and NMR center for providing support for the data collection. This work utilized the computational resources of the NIH HPC Biowulf cluster (http://hpc.nih.gov). We thank Sil van der Woerd for graciously allowing us to use his film as a behavioral task. In addition, we thank the subjects who generously contributed their data to this project. | Nugent et al., 2022 |
| Enhanced Nathan Kline Institute Rockland Sample | NKI-RS | https://fcon_1000.projects.nitrc.org/indi/enhanced/ | We would like to thank Lawrence Maayan for his key role in the design and execution of the pilot NKI-RS. | Nooner et al., 2012; Tobe et al., 2022 |

| | | | | |
|---|---|---|---|---|
| Pediatric Imaging, Neurocognition, and Genetics | PING | https://chd.ucsd.edu/research/ping-study.html | Data used in the preparation of this article were obtained from the Pediatric Imaging, Neurocognition and Genetics (PING) Study database (www.chd.ucsd.edu/research/ping-study.html, now shared through the NIMH Data Archive (NDA)). PING was a multisite, cross-sectional study that recruited more than 1,700 participants aged 3 to 20 years. The study was supported by award number RC2DA029475 from the National Institute on Drug Abuse, with additional support for data sharing provided by the Eunice Kennedy Shriver National Institute of Child Health & Human Development under award number R01HD061414. A list of participating sites and study investigators can be found at https://ping-dataportal.ucsd.edu/sharing/Authors10222012.pdf. PING investigators designed and implemented the study and/or provided data but did not necessarily participate in analysis or writing of this report. This publication is solely the responsibility of the authors and does not necessarily represent the views of the National Institutes of Health or PING investigators. | Jernigan et al., 2016 |
| Amsterdam Open MRI Collection PIOP1 | PIOP1 | https://openneuro.org/datasets/ds002785/versions/2.0.0 | We thank all research assistants and students who helped collecting the data of the three projects, Jasper Wijnen and Marco Teunisse for advice and guidance with respect to anonymization and GDPR-related concerns, and Jos Bloemers, Sennay Ghebeab, Adriaan Tuiten, Joram van Driel, Christian Olivers, Ilja Sligte, Sara Jahfari, Guido van Wingen, and Suzanne Oosterwijk for help with designing the paradigms, Marcus Spaan for technical support, and Franklin Feingold and Joe Wexler for help with uploading the datasets to OpenNeuro. | Snoek et al., 2021 |
| Amsterdam Open MRI Collection PIOP2 | PIOP2 | https://openneuro.org/datasets/ds002790/versions/2.0.0 | We thank all research assistants and students who helped collecting the data of the three projects, Jasper Wijnen and Marco Teunisse for advice and guidance with respect to anonymization and GDPR-related concerns, and Jos Bloemers, Sennay Ghebeab, Adriaan Tuiten, Joram van Driel, Christian Olivers, Ilja Sligte, Sara Jahfari, Guido van Wingen, and Suzanne Oosterwijk for help with designing the paradigms, Marcus Spaan for technical support, and Franklin Feingold and Joe Wexler for help with uploading the datasets to OpenNeuro. | Snoek et al., 2021 |
| Philadelphia Neurodevelopmental Cohort | PNC | https://www.med.upenn.edu/bbl/philadelphianeurodevelopmentalcohort.html | Support for the collection of the data sets was provided by grant RC2MH089983 awarded to R. Gur and RC2MH089924 awarded to H. Hakonarson. | Satterthwaite et al., 2016; Satterthwaite et al., 2014 |
| Queensland Twin Adolescent Brain | QTAB | https://openneuro.org/datasets/ds004146/versions/1.0.4 | We are forever grateful to the twins and their families for their willingness to participate in our studies. We thank Liza van Eijk, Victoria O'Callaghan, Islay Davies, Ethan Campi, Kimberley Huang, Eleanor Roga, Michael Day, Aiman Al-Najjar, Zoie Nott, Tom Shaw, Nicole Atcheson, and Sarah Daniel for data acquisition. We thank Naomi Wray for funding the collection of metabolic | Strike et al., 2023 |

| | | | | |
|---|---|---|---|---|
| | | | samples, including detailed dietary data, Ian Hickie and Kathleen Merikangas for funding and support of actigraphy data, and Sarah Medland and ENIGMA GWAS for funding genotyping. Special thanks to Anjali Henders, Leanne Wallace, Lorelle Nunn and the many laboratory assistants at the Human Studies Unit (part of the Program in Complex Trait Genomics based at the Institute of Molecular Bioscience, University of Queensland) for the processing and storage of biological samples. Thanks also to Julie Henry for helpful discussion of social cognition measures. The QTAB project was funded by the National Health and Medical Research Council (NHMRC), Australia (Project Grant ID: 1078756 to MJW), the Queensland Brain Institute, University of Queensland, and with the assistance of resources from the Centre for Advanced Imaging and the Queensland Cyber Infrastructure Foundation, University of Queensland. We acknowledge the Queensland Twin Registry (QTwin) (https://www.qimrberghofer.edu.au/study/queensland-twin-registry-study) for generously sharing database information for recruitment. Recruitment was further facilitated through access to Twins Research Australia, a national resource supported by a Centre of Research Excellence Grant (ID: 1079102) from the NHMRC. Lastly, we thank the many researchers worldwide for providing access to their assessments. | |
| Queensland Twin IMaging | QTIM | https://openneuro.org/datasets/ds004169/versions/1.0.6 | We are forever grateful to the twins and siblings for their willingness to participate in our studies. We thank Marlene Grace and Ann Eldridge for participant recruitment; Kerrie McAloney for study co-ordination; Kori Johnson, Aaron Quiggle, Natalie Garden, Matthew Meredith, Peter Hobden, Kate Borg, Aiman Al Najjar and Anita Burns for data acquisition; David Butler and Daniel Park for IT support. | Strike et al., 2019 |
| Southwest University Adult Lifespan Dataset | SALD | https://fcon_1000.projects.nitrc.org/indi/retro/sald.html | This data repository was supported by the National Natural Science Foundation of China (31470981; 31571137; 31500885), National Outstanding Young People Plan, the Program for the Top Young Talents by Chongqing, the Fundamental Research Funds for the Central Universities (SWU1509383, SWU1509451, SWU1609177), Natural Science Foundation of Chongqing (cstc2015jcyjA10106), Fok Ying Tung Education Foundation (151023), General Financial Grant from the China Postdoctoral Science Foundation (2015M572423, 2015M580767), Special Funds from the Chongqing Postdoctoral Science Foundation (Xm2015037, Xm2016044), Key research for Humanities and Social Sciences of Ministry of Education (14JJD880009). | Wei et al., 2018 |
| Southwest University Longitudinal | SLIM | https://fcon_1000.projects.nitrc.org/indi/ | We are grateful to all graduate students who contributed their time and wisdom to this data repository, including but not limited to Xue Du, | Y. Wang et al., 2014 |

| | | | | |
|---|---|---|---|---|
| Imaging Multimodal dataset | | retro/southwestuni_qiu_index.html | Kangcheng Wang, Jiangzhou Sun, Qunling Chen and Wei Liu. We are also thankful for assistance from Michael Milham and David O'Connor (Child Mind Institute) in constructing the webpage. This data repository was supported by: The National Natural Science Foundation of China (31271087; 31470981; 31571137; 31500885), National Outstanding Young People Plan, the Program for the Top Young Talents by Chongqing, the Fundamental Research Funds for the Central Universities (SWU1509383, SWU1509451), Natural Science Foundation of Chongqing (cstc2015jcyjA10106), Fok Ying Tung Education Foundation (151023), General Financial Grant from the China Postdoctoral Science Foundation (2015M572423, 2015M580767), Special Funds from the Chongqing Postdoctoral Science Foundation (Xm2015037), Key research for Humanities and Social Sciences of Ministry of Education (14JJD880009). | |
| Parkinson's Disease Datasets (Tao-Wu) | Tao-Wu | https://fcon_1000.projects.nitrc.org/indi/retro/parkinsons.html | This work was partially supported by the NEUROCON project (84/2012), financed by UEFISCDI. | Badea et al., 2017 |
| UK Biobank | UKBB | https://www.ukbiobank.ac.uk/ | This research has been conducted using the UK Biobank Resource (access code 27412). | Sudlow et al., 2015 |
| OpenNeuro dataset ds000119 | ds000119 | https://openneuro.org/datasets/ds000119/versions/00001 | We thank Mark McAvoy and Abraham Snyder for support and development of functional data analysis procedures. Enami Yasui provided assistance with data collection. National Institutes of Mental Health (NIMH RO1 MH067924). | Velanova et al., 2008 |
| OpenNeuro dataset ds000171 | ds000171 | https://openneuro.org/datasets/ds000171/versions/00001 | The authors wish to acknowledge Trisha Patrician and Natalie Stroupe for their assistance with screening of participants, and Allan Schmitt and Franklin Hunsinger for their role in collecting the MR data. | Lepping et al., 2016 |
| OpenNeuro dataset ds000202 | ds000202 | https://openneuro.org/datasets/ds000202/versions/00001 | | Van Schuerbeek et al., 2016 |
| OpenNeuro dataset ds000222 | ds000222 | https://openneuro.org/datasets/ds000222/versions/1.0.1 | | |

Supplementary Table 2: Included outcome variables

| Description | ABCD element name | ABCD table name | Category | n baseline | n follow-up |
|---|---|---|---|---|---|
| Height | ph_y_anthr__height_mean | ph_y_anthr | Personal | 11,085 | 7,698 |
| Weight | ph_y_anthr__weight_mean | ph_y_anthr | Personal | 11,063 | 7,495 |
| NIHTBX crystallized composite | nc_y_nihtb__comp__cryst__uncor_score | nc_y_nihtb | Personal | 10,931 | 0 |
| NIHTBX fluid composite | nc_y_nihtb__comp__fluid__uncor_score | nc_y_nihtb | Personal | 10,882 | 0 |
| WISC matrix reasoning | nc_y_wisc__raw_score | nc_y_wisc | Personal | 10,801 | 0 |
| CBCL internalizing | mh_p_cbcl__synd__int_sum | mh_p_cbcl | Personal | 10,831 | 7,506 |
| CBCL externalizing | mh_p_cbcl__synd__ext_sum | mh_p_cbcl | Personal | 10,233 | 7,209 |
| UPPS-P Negative Urgency | mh_y_upps__nurg_sum | mh_y_upps | Personal | 11,084 | 7,699 |
| UPPS-P Lack of perserverance | mh_y_upps__pers_sum | mh_y_upps | Personal | 10,827 | 7,553 |
| UPPS-P Lack of Planning | mh_y_upps__plan_sum | mh_y_upps | Personal | 10,815 | 7,555 |
| UPPS-P Positive urgency | mh_y_upps__purg_sum | mh_y_upps | Personal | 11,085 | 7,699 |
| UPPS-P Sensation Seeking | mh_y_upps__sens_sum | mh_y_upps | Personal | 11,085 | 7,699 |
| Sleep disturbance | ph_p_sds | ph_p_sds | Personal | 10,808 | 7,522 |

| Description | ABCD element name | ABCD table name | Category | n baseline | n follow-up |
|---|---|---|---|---|---|
| Screen time | nt_p_yst__wkdy/wknd__hr_001 | nt_p_yst | Personal | 10,824 | 7,360 |
| Birth weight | ph_p_dhx_birthweight | ph_p_dhx | Relational | 10,808 | 7,168 |
| Mothers age at pregnancy | ph_p_dhx_003__01*† | ph_p_dhx | Relational | 10,856 | 7,181 |
| Fathers age at pregnancy | ph_p_dhx_004__01 | ph_p_dhx | Relational | 10,496 | 7,181 |
| Total number of friends | mh_y_resil_sum | mh_y_resil | Relational | 10,424 | 7,196 |
| Parent internalizing | mh_p_asr__synd__int_sum | mh_p_asr | Relational | 10,726 | 7,429 |
| Parent externalizing | mh_p_asr__synd__ext_sum | mh_p_asr | Relational | 10,865 | 7,577 |
| Parent alcohol use | mh_p_asr__drunk_001 | mh_p_asr | Relational | 10,692 | 6,766 |
| Parent drug use | mh_p_asr__drg_001 | mh_p_asr | Relational | 10,802 | 7,234 |
| Family conflict | fc_y_fes__confl_mean | fc_y_fes | Relational | 10,978 | 7,641 |
| Parental monitoring | fc_y_pm_mean | fc_y_pm | Relational | 10,401 | 7,395 |
| School environment | fc_y_srpf__env_mean | fc_y_srpf | Relational | 11,060 | 7,685 |
| School involvement | fc_y_srpf__involv_mean | fc_y_srpf | Relational | 11,081 | 7,696 |
| Ethnic identity affiliation | fc_p_meim_mean | fc_p_meim | Socioenvironmental | 10,446 | 7,630 |
| Neighbourhood safety | fc_p_nsc__ns_mean | fc_p_nsc | Socioenvironmental | 11,058 | 7,664 |

| Description | ABCD element name | ABCD table name | Category | n baseline | n follow-up |
|---|---|---|---|---:|---:|
| Residential density | le_l_densbld__addr1_density | le_l_densbld | Socioenvironmental | 10,150 | 0 |
| Walkability index | le_l_walk__addr1_idx | le_l_walk | Socioenvironmental | 10,515 | 0 |
| Proximity to major roads | le_l_roadprox__addr1_m | le_l_roadprox | Socioenvironmental | 10,137 | 0 |
| Crime Reports: grand total | le_l_crime__addr1_count | le_l_crime | Socioenvironmental | 9,755 | 0 |
| Estimated lead risk | le_l_leadrisk__addr1_idx | le_l_leadrisk | Socioenvironmental | 10,513 | 0 |
| Traffic Density | le_l_traffic__addr1__traffic_count | le_l_traffic | Socioenvironmental | 10,198 | 0 |
| NO2 Measures | le_l_no2__addr1__no2_mean__2016 | le_l_no2 | Socioenvironmental | 10,515 | 0 |
| PM25 Measures | le_l_pm25__addr1__pm25_mean__2016 | le_l_pm25 | Socioenvironmental | 10,512 | 0 |
| Area deprivation index | le_l_adi__addr1_wsum | le_l_adi | Socioenvironmental | 9,992 | 0 |
| Child opportunity index | le_l_coi__addr1__coi__total__national_score | le_l_coi | Socioenvironmental | 10,096 | 0 |

* Weighted average of nt_p_yst__wkdy__hr_001 and nt_p_yst__wknd__hr_001

† Sum of 26 item Likert scale items

#### Supplementary Table 3: Correlation between training targets

| | Age | Sex | Handedness | BMI | Fluid Intelligence | Neuroticism |
|---|---|---|---|---|---|---|
| Age | 1 | 0.02 | 0.03 | 0.15 | -0.08 | -0.1 |
| Sex | 0.02 | 1 | -0.02 | 0.11 | 0.07 | -0.15 |
| Handedness | 0.03 | -0.02 | 1 | 0.02 | 0.02 | -0.04 |
| BMI | 0.15 | 0.11 | 0.02 | 1 | -0.04 | -0.02 |
| Fluid Intelligence | -0.08 | 0.07 | 0.02 | -0.04 | 1 | -0.03 |
| Neuroticism | -0.1 | -0.15 | -0.04 | -0.02 | -0.03 | 1 |

**Supplementary table 3**: Correlation between the targets used for model training. Correlation between continuous variables were measured using the Spearman rank coefficient, between continuous and binary variables using the point biserial coefficient, and between binary variables using the Matthews coefficient.

---

#### Supplementary Table 4: Intraclass correlation (ICC), $r$ and paired samples t-test of fuzzy clustering cortical surface measures between baseline and 2-year follow-up

| | | Thickness | | | Area | | | Volume | | |
|---|---|---|---|---|---|---|---|---|---|---|
| Region | Hemi | ICC | r | t | ICC | r | t | ICC | r | t |
| Frontal | LH | 0.69 | 0.76 | 26.11 ‡ | 0.95 | 0.95 | -2.89 | 0.94 | 0.94 | 5.32 * |
| Frontal | RH | 0.68 | 0.76 | 27.32 ‡ | 0.94 | 0.94 | -2.11 | 0.93 | 0.94 | 6.39 * |
| Occipital | LH | 0.66 | 0.80 | 36.32 ‡ | 0.96 | 0.96 | 1.71 | 0.93 | 0.95 | 13.77 ‡ |
| Occipital | RH | 0.65 | 0.79 | 36.74 ‡ | 0.96 | 0.96 | 1.59 | 0.93 | 0.95 | 13.38 ‡ |
| Parietal | LH | 0.70 | 0.79 | 28.53 ‡ | 0.96 | 0.96 | 1.00 | 0.94 | 0.96 | 9.38 ‡ |
| Parietal | RH | 0.71 | 0.79 | 27.69 ‡ | 0.96 | 0.96 | 0.41 | 0.94 | 0.95 | 8.42 ‡ |
| Temporal | LH | 0.67 | 0.75 | 27.35 ‡ | 0.96 | 0.96 | -0.51 | 0.94 | 0.95 | 7.79 ‡ |
| Temporal | RH | 0.67 | 0.76 | 28.04 ‡ | 0.96 | 0.96 | 0.06 | 0.94 | 0.95 | 8.29 ‡ |

**Supplementary Table 4**: ICC were obtained by fitting the linear mixed model $y_{ij} = \beta_0 + u_{0j} + \epsilon_{ij}$ for each included surface measure and then taking $\mathrm{ICC} = \frac{\sigma^2_{\mathrm{subject}}}{\sigma^2_{\mathrm{subject}} + \sigma^2_{\mathrm{resid}}}$. $r$ is the between subject correlation between the first and second session. t is from paired samples t-tests between the first and second session (df = 7315), where ‡ = $p < 0.05^{-10}$ and * = $p < 0.05$.

Supplementary Table 5: Intraclass correlation (ICC), r and paired samples t-test of embedding dimensions between baseline and 2-year follow-up

---

**Supplementary Table 5**: ICC were obtained by fitting the linear mixed model $y_{ij} = \beta_0 + u_{0j} + \epsilon_{ij}$ for each included embedding dimension and then taking $\mathrm{ICC} = \frac{\sigma^2_{\mathrm{subject}}}{\sigma^2_{\mathrm{subject}} + \sigma^2_{\mathrm{resid}}}$. r is the between subject correlation between the first and second session. t is from paired samples t-tests between the first and second session (df = 7315), where ‡ = $p < 0.05^{-10}$ and * = $p < 0.05$.

| Dimension | ICC | r | t |
|---|---|---|---|
| 0 | 0.39 | 0.49 | 30.87 ‡ |
| 1 | 0.55 | 0.55 | -8.21 ‡ |
| 2 | 0.27 | 0.35 | -26.83 ‡ |
| 3 | 0.36 | 0.44 | 29.90 ‡ |
| 4 | 0.45 | 0.62 | 41.98 ‡ |
| 5 | 0.33 | 0.34 | 5.50 * |
| 6 | 0.35 | 0.35 | -4.02 * |
| 7 | 0.39 | 0.46 | 27.12 ‡ |
| 8 | 0.54 | 0.54 | 7.32 ‡ |
| 9 | 0.41 | 0.41 | -2.13 * |
| 10 | 0.29 | 0.35 | -16.95 ‡ |
| 11 | 0.25 | 0.30 | -16.97 ‡ |
| 12 | 0.36 | 0.41 | -22.05 ‡ |
| 13 | 0.40 | 0.51 | 33.69 ‡ |
| 14 | 0.35 | 0.55 | -45.85 ‡ |
| 15 | 0.34 | 0.41 | -23.36 ‡ |
| 16 | 0.52 | 0.56 | -18.80 ‡ |
| 17 | 0.34 | 0.35 | 10.75 ‡ |
| 18 | 0.36 | 0.38 | -11.86 ‡ |
| 19 | 0.24 | 0.25 | 9.51 ‡ |
| 20 | 0.38 | 0.43 | 23.70 ‡ |
| 21 | 0.57 | 0.57 | -0.58 |
| 22 | 0.40 | 0.49 | -31.56 ‡ |
| 23 | 0.36 | 0.47 | 32.68 ‡ |
| 24 | 0.37 | 0.45 | 29.26 ‡ |
| 25 | 0.49 | 0.50 | -5.69 * |
| 26 | 0.42 | 0.42 | -3.27 * |
| 27 | 0.33 | 0.45 | 36.47 ‡ |
| 29 | 0.37 | 0.41 | 18.75 ‡ |
| 30 | 0.44 | 0.46 | -10.92 ‡ |
| 31 | 0.47 | 0.48 | -4.11 * |
| 32 | 0.54 | 0.55 | -10.24 ‡ |
| 33 | 0.28 | 0.28 | -5.44 * |
| 34 | 0.30 | 0.33 | -12.80 ‡ |
| 35 | 0.37 | 0.46 | 30.36 ‡ |
| 36 | 0.50 | 0.51 | -11.47 ‡ |
| 37 | 0.53 | 0.54 | 12.32 ‡ |
| 38 | 0.39 | 0.47 | -28.58 ‡ |
| 39 | 0.43 | 0.43 | 2.28 * |
| 40 | 0.51 | 0.51 | 1.80 |
| 41 | 0.35 | 0.37 | -11.30 ‡ |
| 42 | 0.28 | 0.30 | -6.84 ‡ |
| 43 | 0.40 | 0.41 | -5.28 * |
| 44 | 0.41 | 0.45 | 20.39 ‡ |
| 45 | 0.42 | 0.48 | 24.24 ‡ |
| 46 | 0.34 | 0.37 | 15.03 ‡ |
| 47 | 0.40 | 0.41 | -11.82 ‡ |
| 48 | 0.38 | 0.39 | -4.45 * |
| 49 | 0.36 | 0.43 | 26.03 ‡ |
| 50 | 0.31 | 0.33 | -3.88 * |
| 52 | 0.41 | 0.51 | 32.20 ‡ |
| 53 | 0.27 | 0.35 | 29.16 ‡ |
| 54 | 0.46 | 0.48 | -6.38 * |
| 55 | 0.37 | 0.44 | 26.78 ‡ |
| 56 | 0.39 | 0.41 | 15.50 ‡ |
| 58 | 0.29 | 0.30 | -6.40 * |
| 60 | 0.39 | 0.42 | 18.68 ‡ |
| 61 | 0.30 | 0.32 | -13.25 ‡ |
| 62 | 0.43 | 0.58 | 40.90 ‡ |
| 63 | 0.31 | 0.35 | 23.28 ‡ |

Supplementary Table 6: Model comparison, baseline

| Outcome | Surface measure | Embedding number | Adjusted $R^2$ without embedding in model | Adjusted $R^2$ with embedding in model | F | p |
|---|---|---|---|---|---|---|
| Height | Area temporal LH | 10 | 0.214 | 0.258 | 653.913 | 6.629e-139 |
| Weight | Area frontal LH | 10 | 0.061 | 0.149 | 1,149.670 | 2.936e-238 |
| NIHTBX crystallized composite | Volume temporal RH | 32 | 0.138 | 0.139 | 11.849 | 1.223e-03 |
| NIHTBX fluid composite | Volume temporal RH | 12 | 0.095 | 0.098 | 29.464 | 2.211e-07 |
| WISC matrix reasoning | Volume temporal RH | 32 | 0.063 | 0.064 | 6.317 | 1.685e-02 |
| CBCL internalizing | Thickness temporal RH | 14 | 0.000 | 0.001 | 9.808 | 3.310e-03 |
| CBCL externalizing | Thickness frontal LH | 8 | 0.010 | 0.010 | 7.276 | 1.108e-02 |
| UPPS-P Negative Urgency | Thickness frontal LH | 8 | 0.008 | 0.009 | 11.226 | 1.618e-03 |
| UPPS-P Lack of perserverance | Thickness occipital RH | 46 | 0.006 | 0.006 | 3.353 | 6.894e-02 |
| UPPS-P Lack of Planning | Volume temporal RH | 4 | 0.016 | 0.017 | 3.857 | 5.380e-02 |
| UPPS-P Positive urgency | Thickness frontal LH | 8 | 0.008 | 0.011 | 32.748 | 5.111e-08 |

| Outcome | Surface measure | Embedding number | Adjusted $R^2$ without embedding in model | Adjusted $R^2$ with embedding in model | F | p |
|---|---|---|---|---|---|---|
| UPPS-P Sensation Seeking | Volume temporal LH | 4 | 0.019 | 0.019 | 3.920 | 5.335e-02 |
| Sleep disturbance | Thickness parietal LH | 9 | 0.000 | 0.002 | 19.106 | 3.648e-05 |
| Screen time | Thickness frontal LH | 26 | 0.008 | 0.012 | 35.057 | 1.791e-08 |
| Birth weight | Volume temporal RH | 10 | 0.054 | 0.058 | 40.631 | 1.212e-09 |
| Mothers age at pregnancy | Volume parietal LH | 32 | 0.025 | 0.026 | 18.152 | 5.583e-05 |
| Fathers age at pregnancy | Volume parietal RH | 32 | 0.013 | 0.015 | 17.440 | 7.572e-05 |
| Total number of friends | Area temporal LH | 26 | 0.005 | 0.005 | 6.605 | 1.547e-02 |
| Parent internalizing | Thickness frontal RH | 14 | 0.001 | 0.002 | 14.862 | 2.602e-04 |
| Parent externalizing | Thickness frontal LH | 8 | 0.001 | 0.002 | 15.447 | 2.028e-04 |
| Parent alcohol use | Volume temporal RH | 32 | 0.001 | 0.001 | 1.670 | 1.964e-01 |
| Parent drug use | Thickness parietal RH | 13 | 0.000 | 0.001 | 9.170 | 4.462e-03 |
| Family conflict | Thickness frontal LH | 47 | 0.006 | 0.008 | 21.266 | 1.281e-05 |
| Parental monitoring | Area frontal RH | 10 | 0.036 | 0.037 | 4.489 | 3.955e-02 |
| School environment | Thickness temporal RH | 42 | 0.012 | 0.012 | 4.478 | 3.955e-02 |

| Outcome | Surface measure | Embedding number | Adjusted $R^2$ without embedding in model | Adjusted $R^2$ with embedding in model | F | p |
|---|---|---|---|---|---|---|
| School involvement | Area temporal RH | 10 | 0.024 | 0.025 | 3.594 | 6.125e-02 |
| Ethnic identity affiliation | Thickness frontal LH | 1 | 0.001 | 0.003 | 25.769 | 1.352e-06 |
| Neighbourhood safety | Volume temporal RH | 39 | 0.017 | 0.017 | 8.308 | 6.534e-03 |
| Residential density | Thickness frontal LH | 9 | 0.000 | 0.003 | 30.736 | 1.279e-07 |
| Walkability index | Thickness frontal LH | 9 | 0.000 | 0.001 | 8.448 | 6.327e-03 |
| Proximity to major roads | Volume occipital RH | 4 | 0.002 | 0.002 | 4.564 | 3.955e-02 |
| Crime Reports: grand total | Thickness frontal LH | 9 | 0.000 | 0.000 | 4.546 | 3.955e-02 |
| Estimated lead risk | Thickness frontal LH | 8 | 0.001 | 0.006 | 60.609 | 7.236e-14 |
| Traffic Density | Area parietal RH | 8 | 0.000 | 0.001 | 4.968 | 3.386e-02 |
| NO2 Measures | Thickness frontal LH | 38 | 0.000 | 0.001 | 6.485 | 1.592e-02 |
| PM25 Measures | Thickness frontal LH | 38 | 0.001 | 0.002 | 5.595 | 2.447e-02 |
| Area deprivation index | Thickness frontal LH | 8 | 0.002 | 0.012 | 99.120 | 3.862e-22 |
| Child opportunity index | Volume temporal RH | 4 | 0.046 | 0.050 | 42.634 | 5.258e-10 |

**Supplementary Table 6:** For each outcome, we obtained the cortical measure the embedding with the highest semipartial correlation with the outcome, and the cortical measure with the highest semipartial correlation with the outcome. For each outcome we then fit a model $\widehat{y^{(t)}} = \beta_0 + \beta_1 \cdot \text{Cortical} + \beta_2 \cdot \text{Age} + \beta_3 \cdot \text{Sex} \ + \epsilon$ , where $Cortical$ is the cortical measure with the highest semipartial correlation with the outcome. Then we fit $\widehat{y^{(t)}} = \beta_0 + \beta_1 \cdot \text{Cortical} + \beta_2 \cdot \text{Embedding} + \beta_3 \cdot \text{Age} + \beta_4 \cdot \text{Sex} \ + \epsilon$, where $Embedding$ is the embedding with the highest semipartial correlation with the outcome, before comparing the two global models using ANOVA tests. The goal is to determine if adding the embedding effect adds to the global $R^2$.

#### Embedding dimension 1

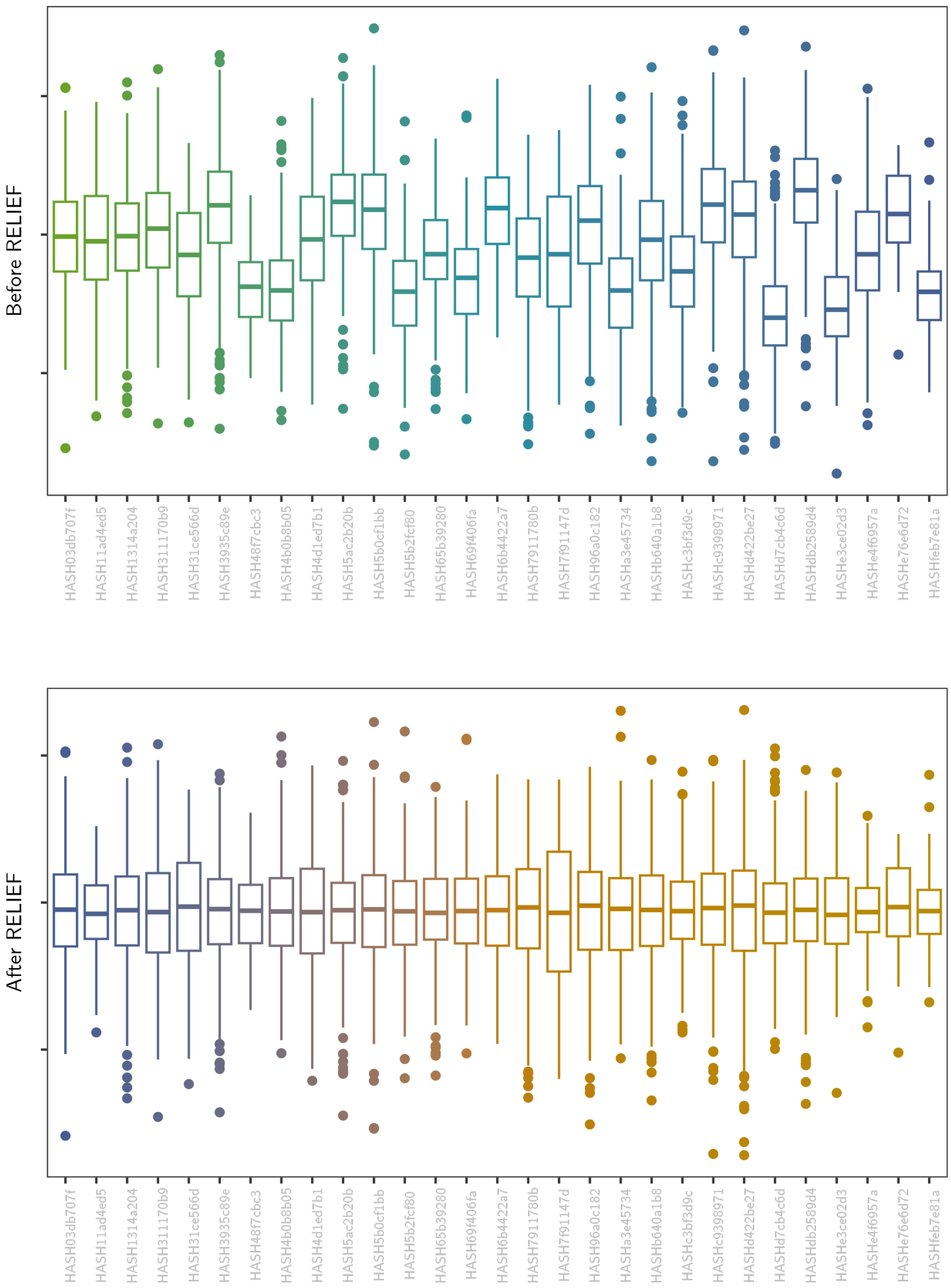

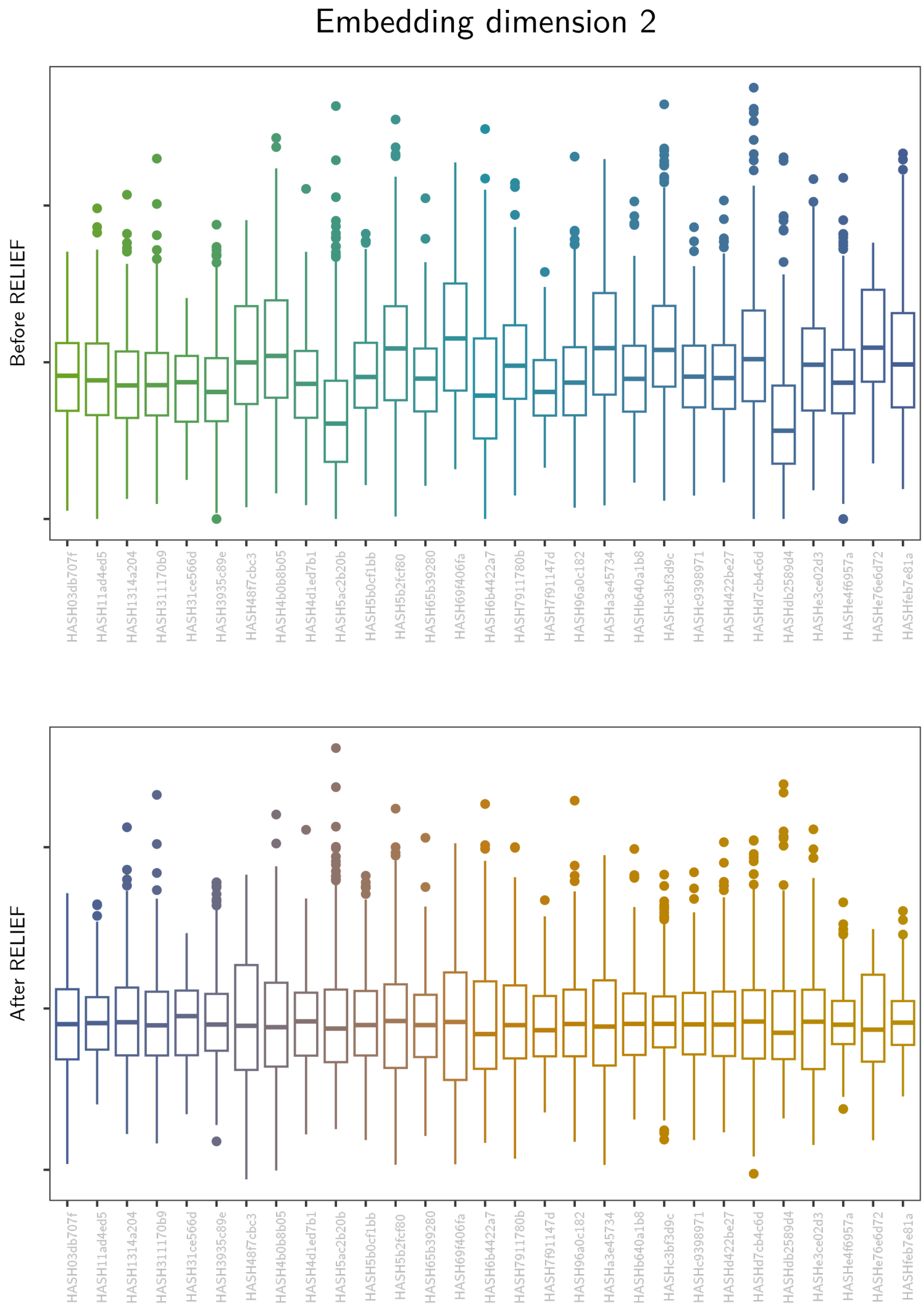

Embedding dimension 2
Before RELIEF
After RELIEF

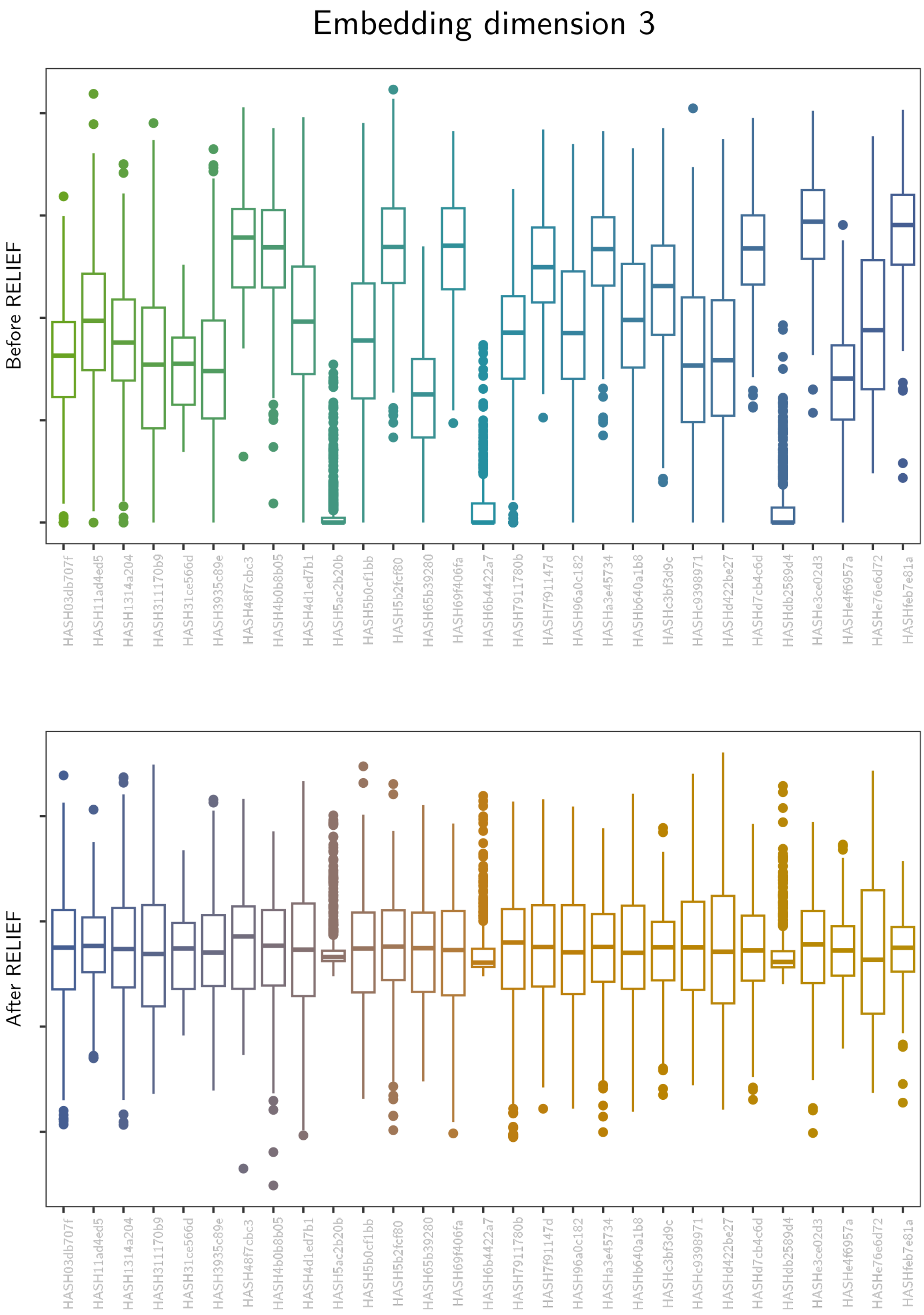
Embedding dimension 3
Before RELIEF
After RELIEF
HASH03db707f
HASH11ad4ed5
HASH1314a204
HASH311170b9
HASH31ce566d
HASH3935c89e
HASH48f7cbc3
HASH4b0b8b05
HASH4d1ed7b1
HASH5ac2b20b
HASH5b0cf1bb
HASH5b2fcf80
HASH65b39280
HASH69f406fa
HASH6b4422a7
HASH7911780b
HASH7f91147d
HASH96a0c182
HASHa3e45734
HASHb640a1b8
HASHc3bf3d9c
HASHc9398971
HASHd422be27
HASHd7cb4c6d
HASHdb2589d4
HASHe3ce02d3
HASHe4f6957a
HASHe76e6d72
HASHfeb7e81a

Embedding dimension 4

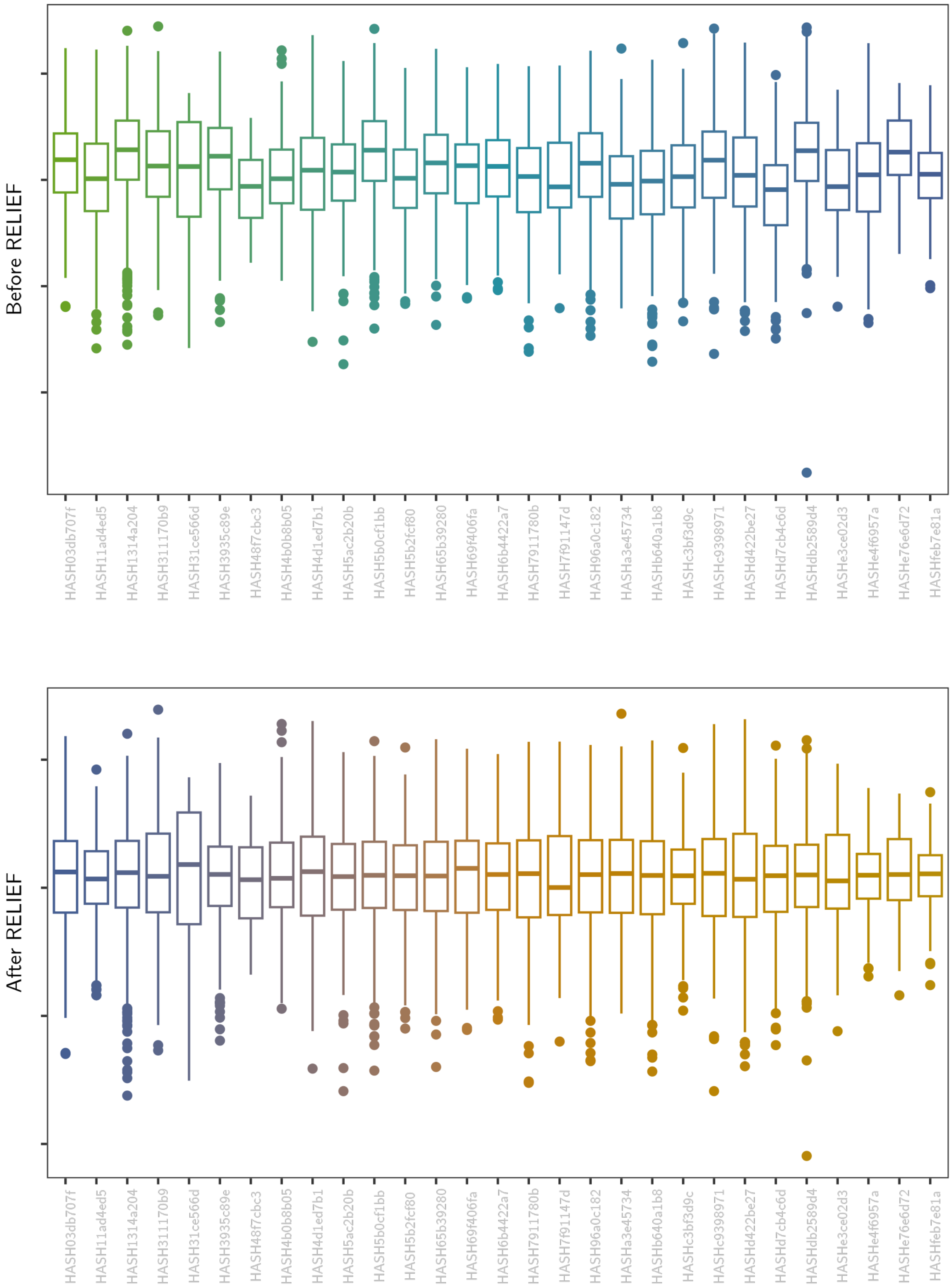

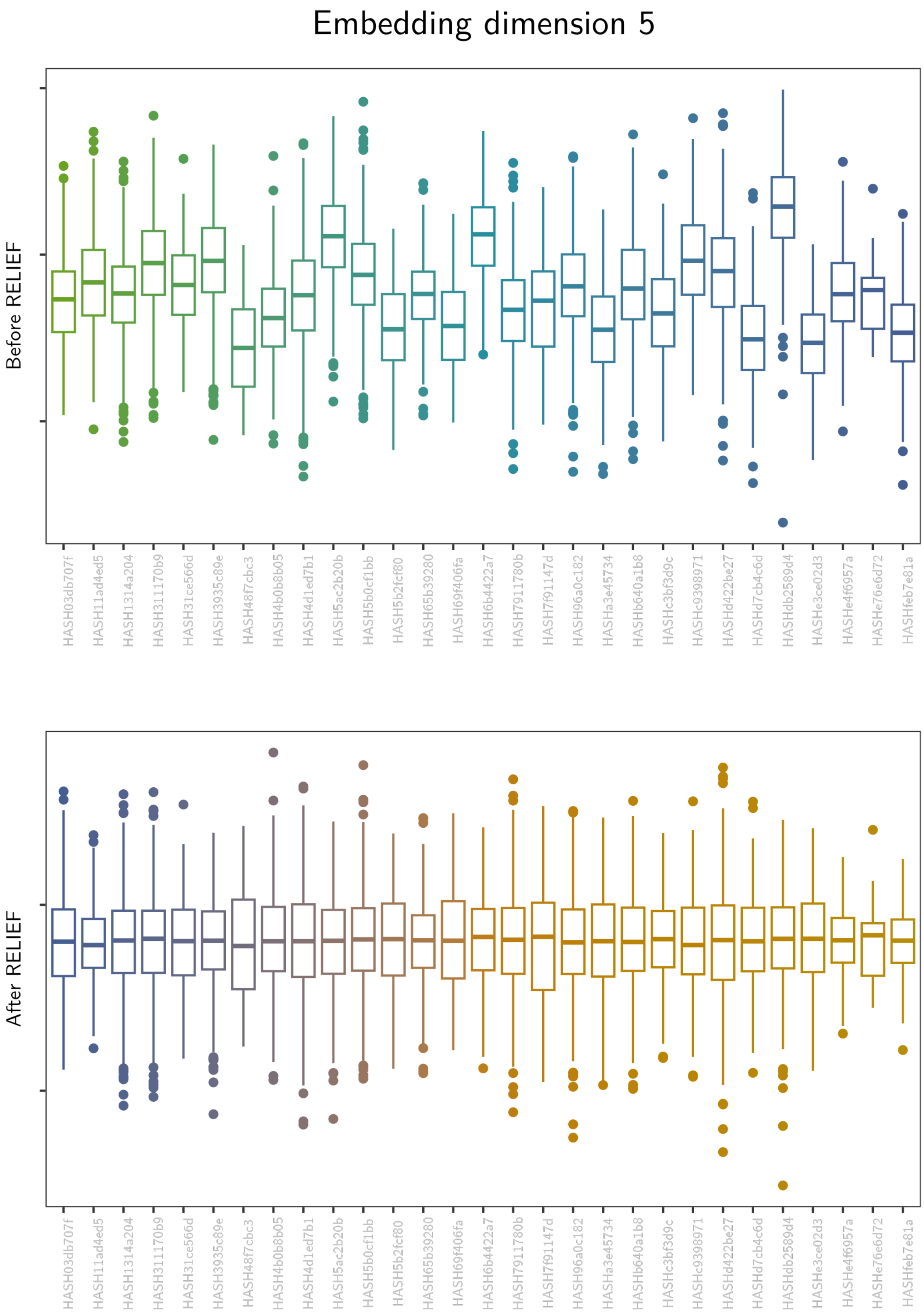

Embedding dimension 5
Before RELIEF
After RELIEF
HASH03db707f
HASH11ad4ed5
HASH1314a204
HASH311170b9
HASH31ce566d
HASH3935c89e
HASH48f7cbc3
HASH4b0b8b05
HASH4d1ed7b1
HASH5ac2b20b
HASH5b0cf1bb
HASH5b2fcf80
HASH65b39280
HASH69f406fa
HASH6b4422a7
HASH7911780b
HASH7f91147d
HASH96a0c182
HASHa3e45734
HASHb640a1b8
HASHc3bf3d9c
HASHc9398971
HASHd422be27
HASHd7cb4c6d
HASHdb2589d4
HASHe3ce02d3
HASHe4f6957a
HASHe76e6d72
HASHfeb7e81a

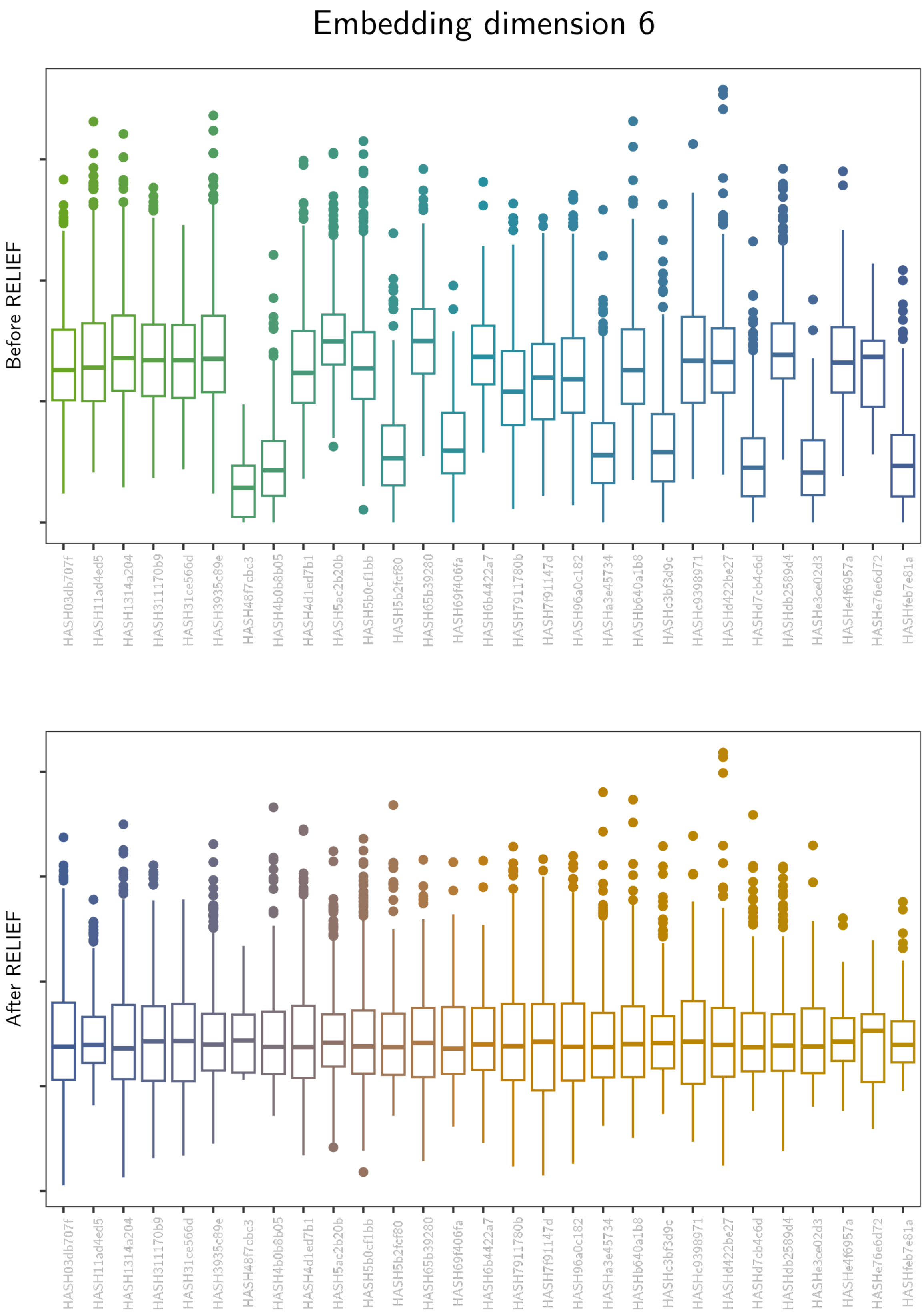
Embedding dimension 6
Before RELIEF
After RELIEF
HASH03db707f
HASH11ad4ed5
HASH1314a204
HASH311170b9
HASH31ce566d
HASH3935c89e
HASH48f7cbc3
HASH4b0b8b05
HASH4d1ed7b1
HASH5ac2b20b
HASH5b0cf1bb
HASH5b2fcf80
HASH65b39280
HASH69f406fa
HASH6b4422a7
HASH7911780b
HASH7f91147d
HASH96a0c182
HASHa3e45734
HASHb640a1b8
HASHc3bf3d9c
HASHc9398971
HASHd422be27
HASHd7cb4c6d
HASHdb2589d4
HASHe3ce02d3
HASHe4f6957a
HASHe76e6d72
HASHfeb7e81a

Embedding dimension 7

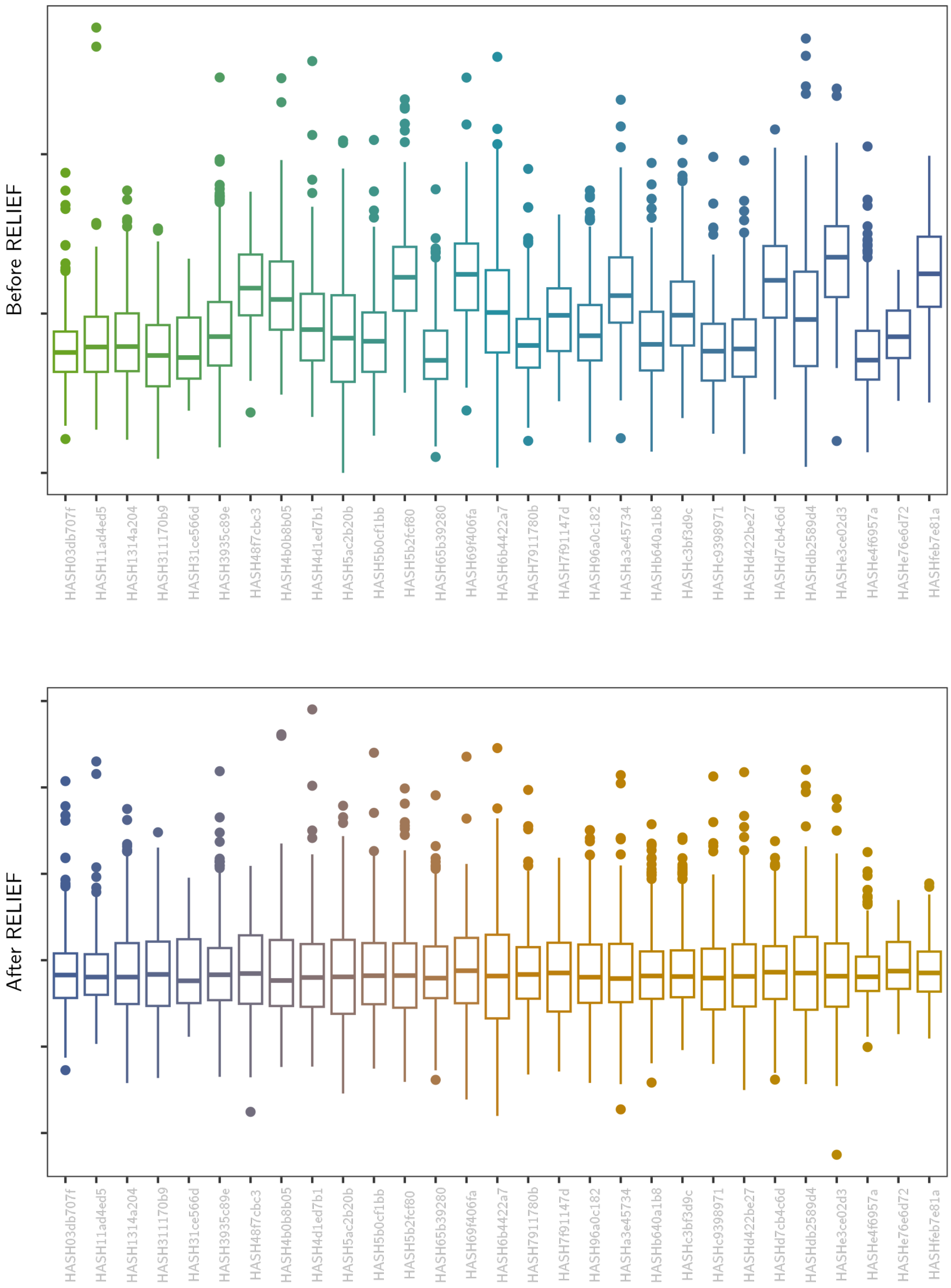

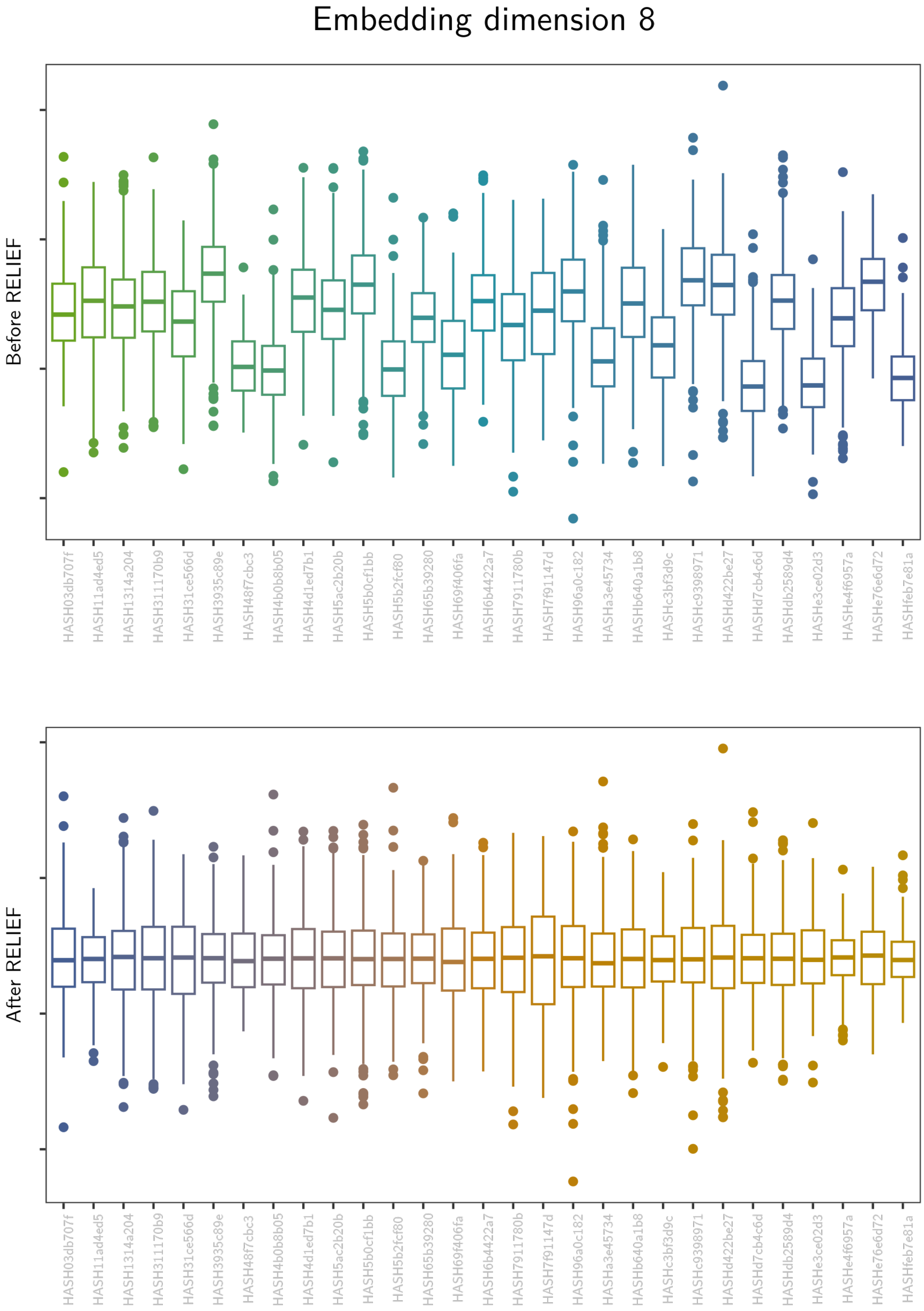
Embedding dimension 8
Before RELIEF
After RELIEF
HASH03db707f
HASH11ad4ed5
HASH1314a204
HASH311170b9
HASH31ce566d
HASH3935c89e
HASH48f7cbc3
HASH4b0b8b05
HASH4d1ed7b1
HASH5ac2b20b
HASH5b0cf1bb
HASH5b2fcf80
HASH65b39280
HASH69f406fa
HASH6b4422a7
HASH7911780b
HASH7f91147d
HASH96a0c182
HASHa3e45734
HASHb640a1b8
HASHc3bf3d9c
HASHc9398971
HASHd422be27
HASHd7cb4c6d
HASHdb2589d4
HASHe3ce02d3
HASHe4f6957a
HASHe76e6d72
HASHfeb7e81a

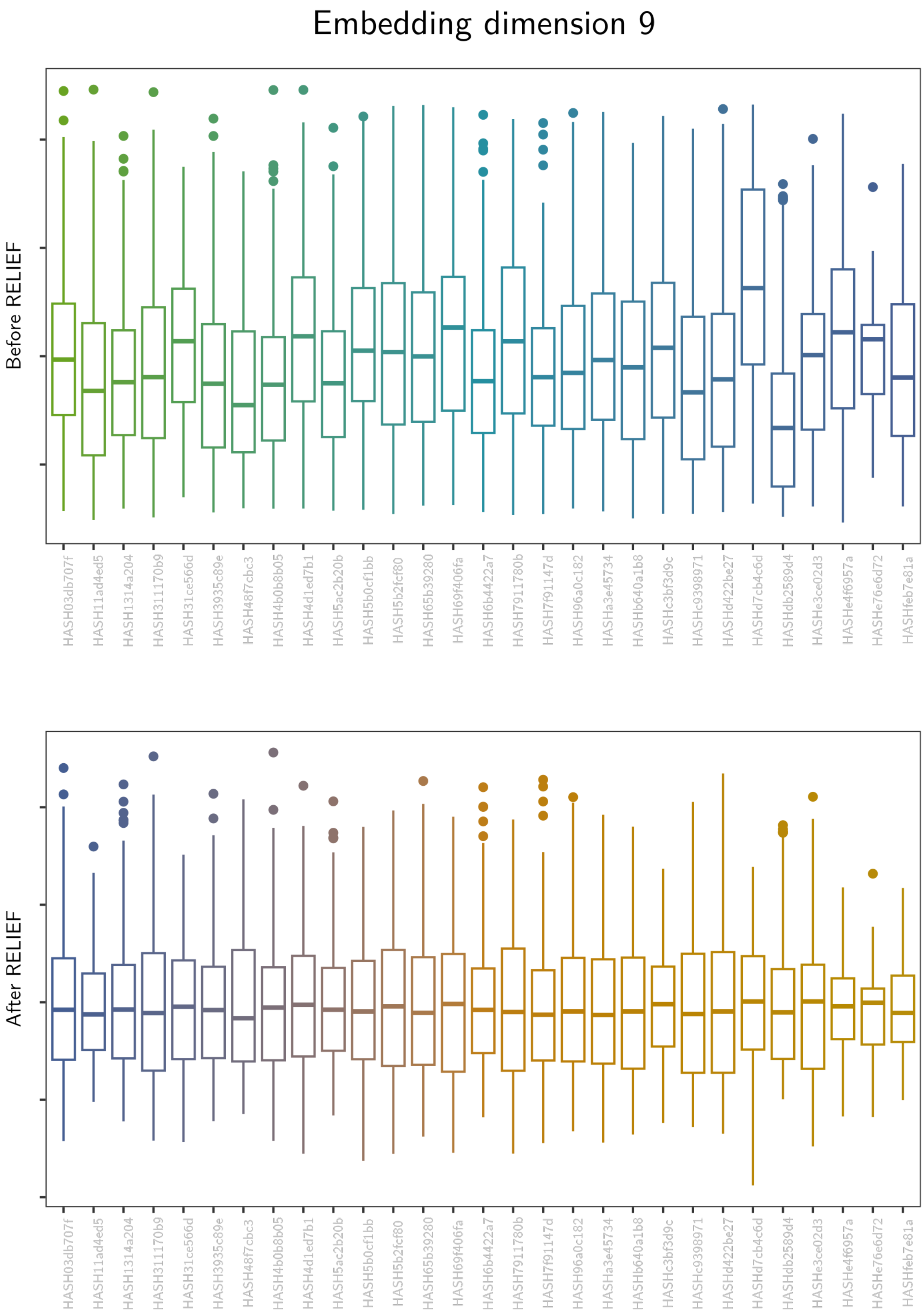

Embedding dimension 9
Before RELIEF
After RELIEF
HASH03db707f
HASH11ad4ed5
HASH1314a204
HASH311170b9
HASH31ce566d
HASH3935c89e
HASH48f7cbc3
HASH4b0b8b05
HASH4d1ed7b1
HASH5ac2b20b
HASH5b0cf1bb
HASH5b2fcf80
HASH65b39280
HASH69f406fa
HASH6b4422a7
HASH7911780b
HASH7f91147d
HASH96a0c182
HASHa3e45734
HASHb640a1b8
HASHc3bf3d9c
HASHc9398971
HASHd422be27
HASHd7cb4c6d
HASHdb2589d4
HASHe3ce02d3
HASHe4f6957a
HASHe76e6d72
HASHfeb7e81a

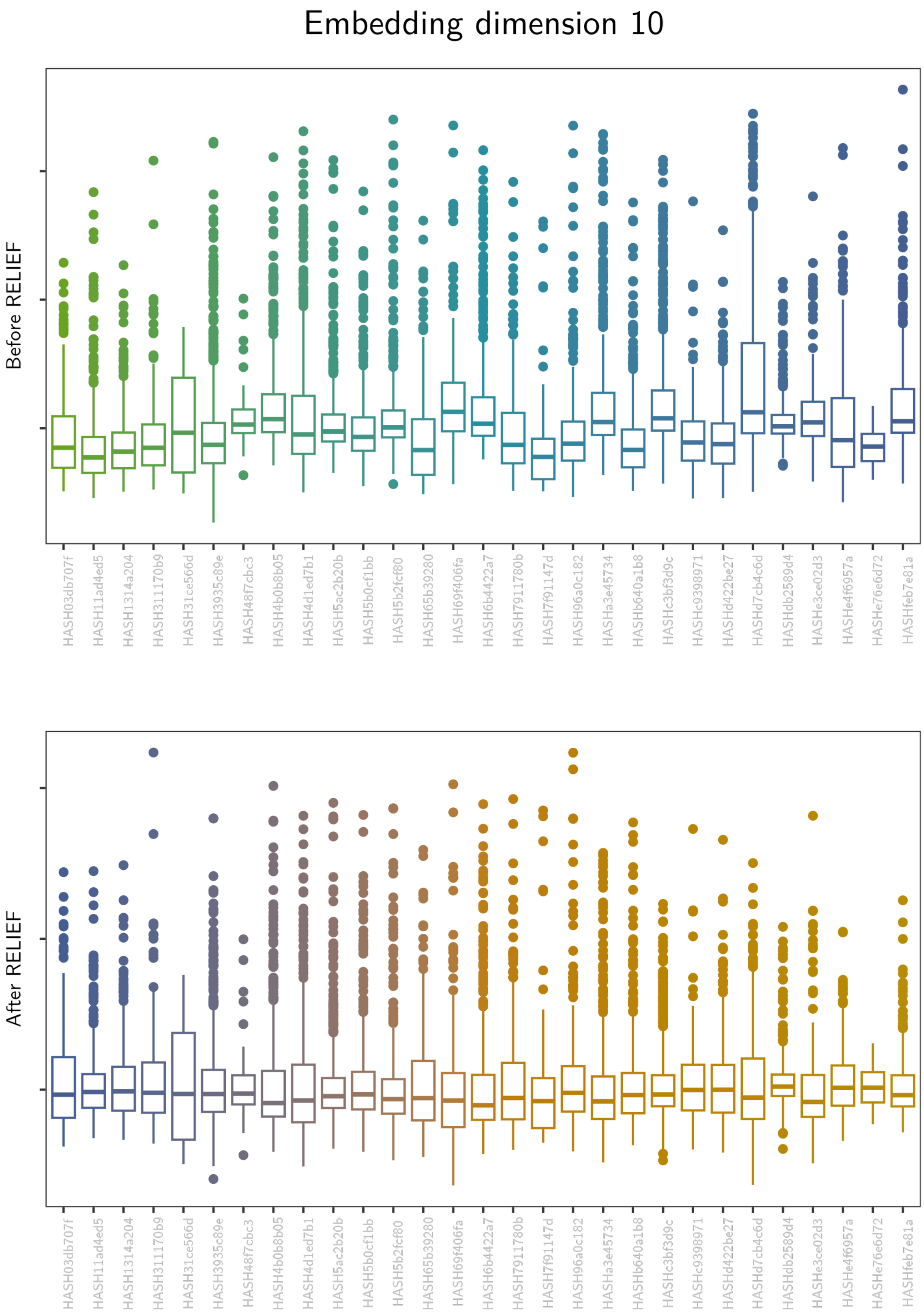
Embedding dimension 10
Before RELIEF
After RELIEF
HASH03db707f
HASH11ad4ed5
HASH1314a204
HASH311170b9
HASH31ce566d
HASH3935c89e
HASH48f7cbc3
HASH4b0b8b05
HASH4d1ed7b1
HASH5ac2b20b
HASH5b0cf1bb
HASH5b2fcf80
HASH65b39280
HASH69f406fa
HASH6b4422a7
HASH7911780b
HASH7f91147d
HASH96a0c182
HASHa3e45734
HASHb640a1b8
HASHc3bf3d9c
HASHc9398971
HASHd422be27
HASHd7cb4c6d
HASHdb2589d4
HASHe3ce02d3
HASHe4f6957a
HASHe76e6d72
HASHfeb7e81a

Embedding dimension 11

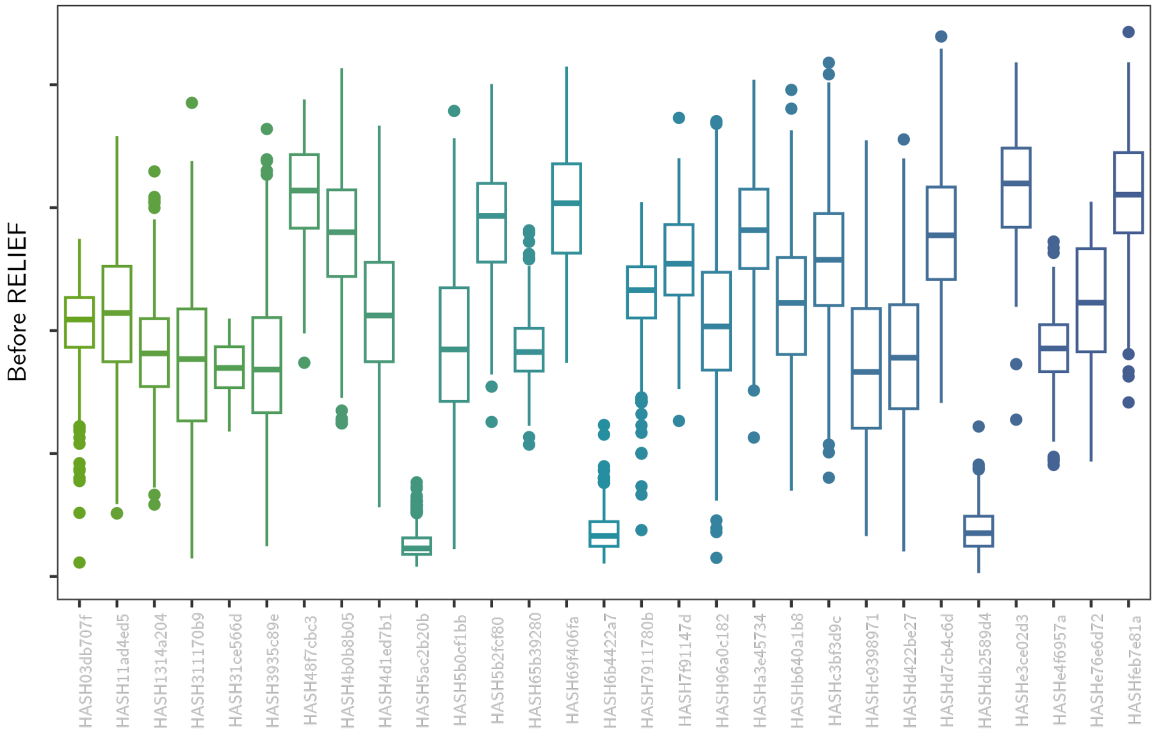

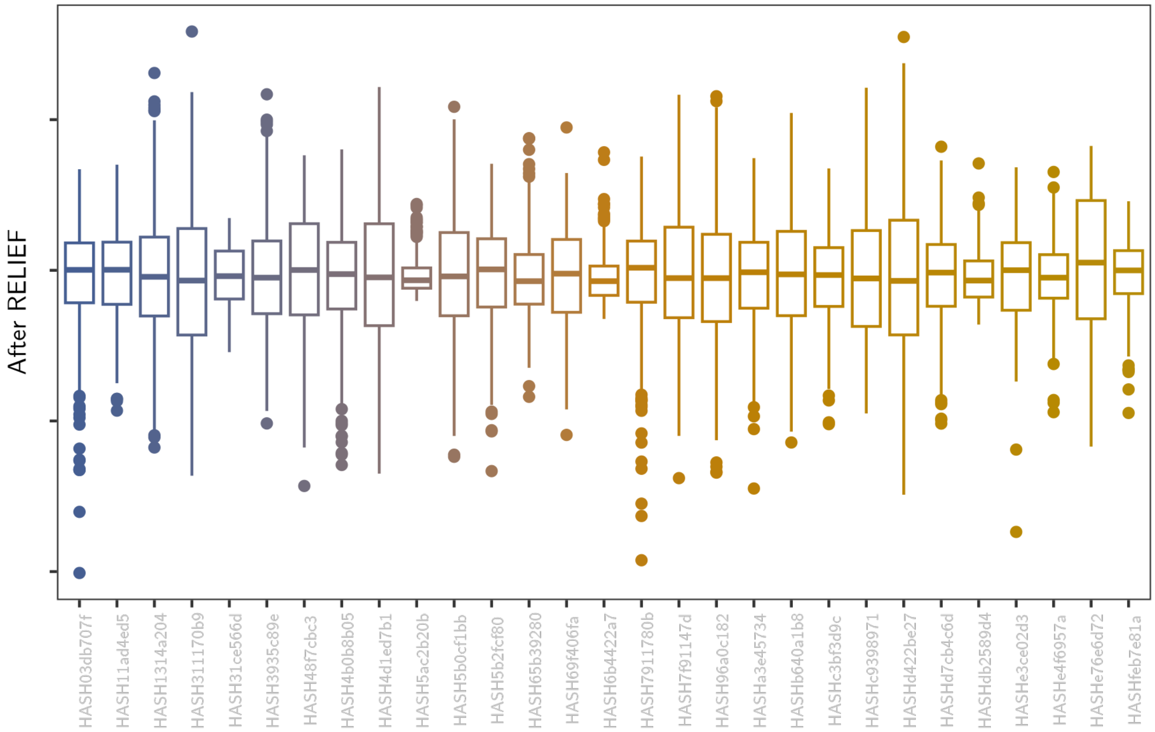

#### Embedding dimension 12

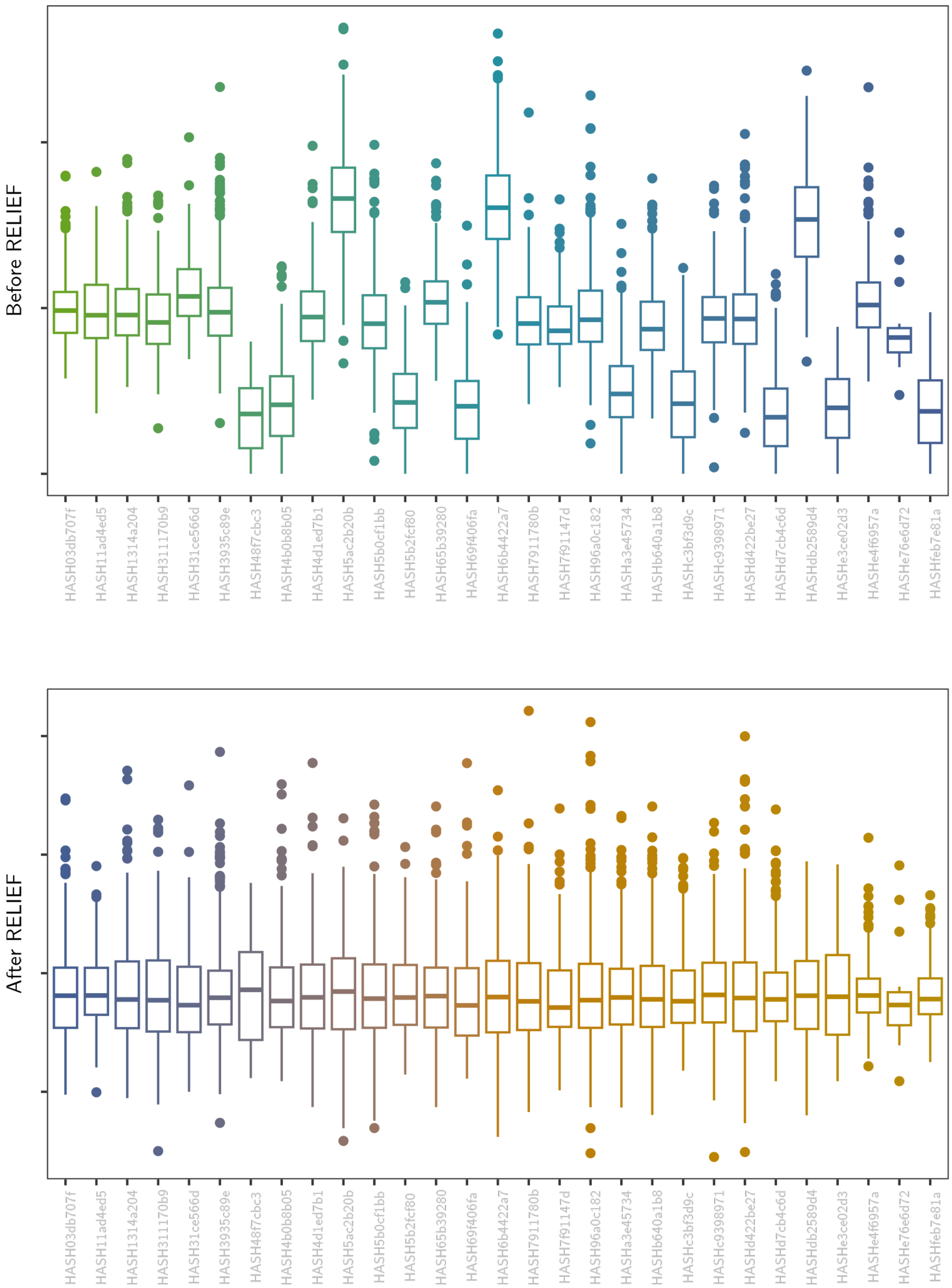

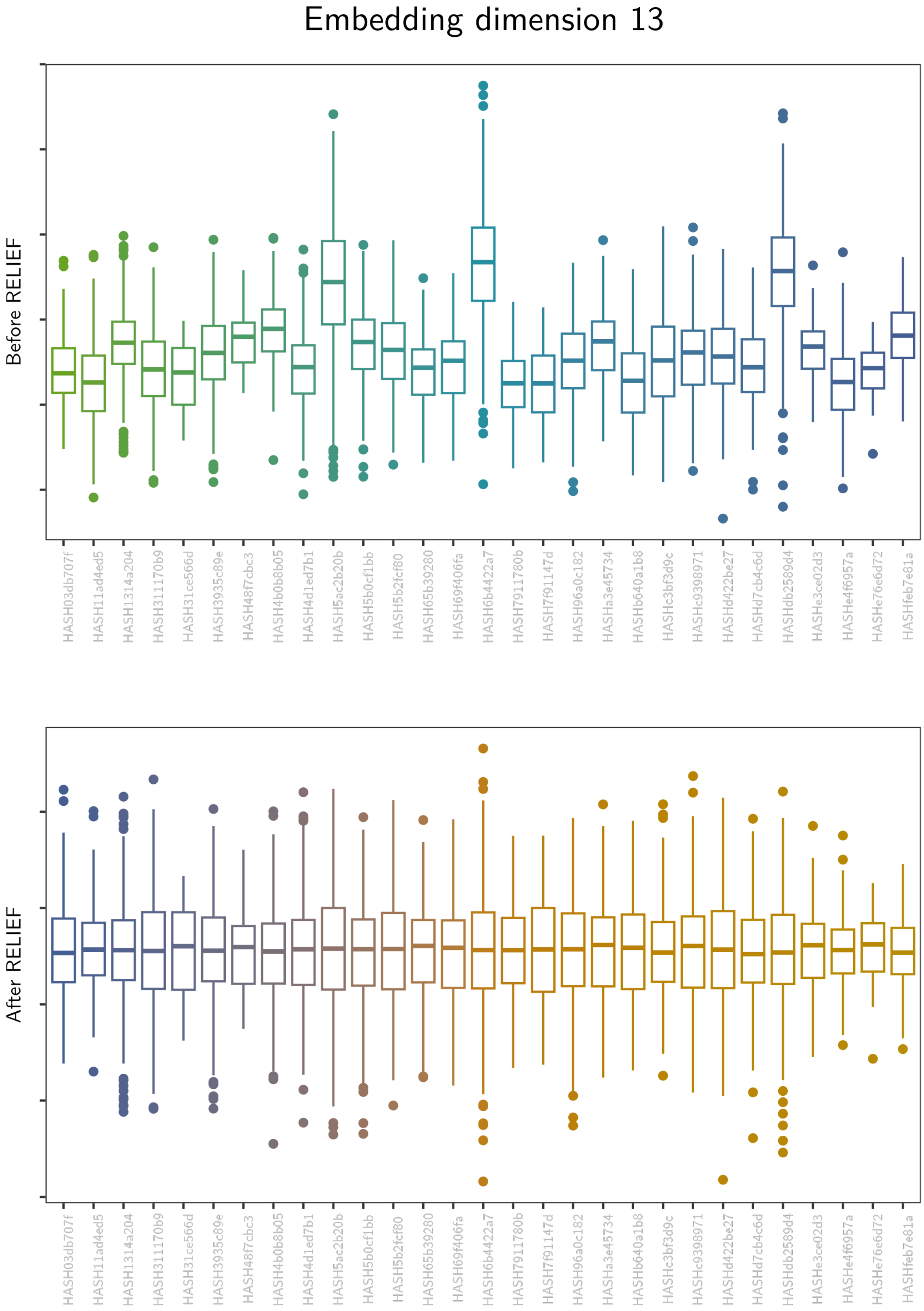
Embedding dimension 13
Before RELIEF
After RELIEF
HASH03db707f
HASH11ad4ed5
HASH1314a204
HASH311170b9
HASH31ce566d
HASH3935c89e
HASH48f7cbc3
HASH4b0b8b05
HASH4d1ed7b1
HASH5ac2b20b
HASH5b0cf1bb
HASH5b2fcf80
HASH65b39280
HASH69f406fa
HASH6b4422a7
HASH7911780b
HASH7f91147d
HASH96a0c182
HASHa3e45734
HASHb640a1b8
HASHc3bf3d9c
HASHc9398971
HASHd422be27
HASHd7cb4c6d
HASHdb2589d4
HASHe3ce02d3
HASHe4f6957a
HASHe76e6d72
HASHfeb7e81a

#### Embedding dimension 14

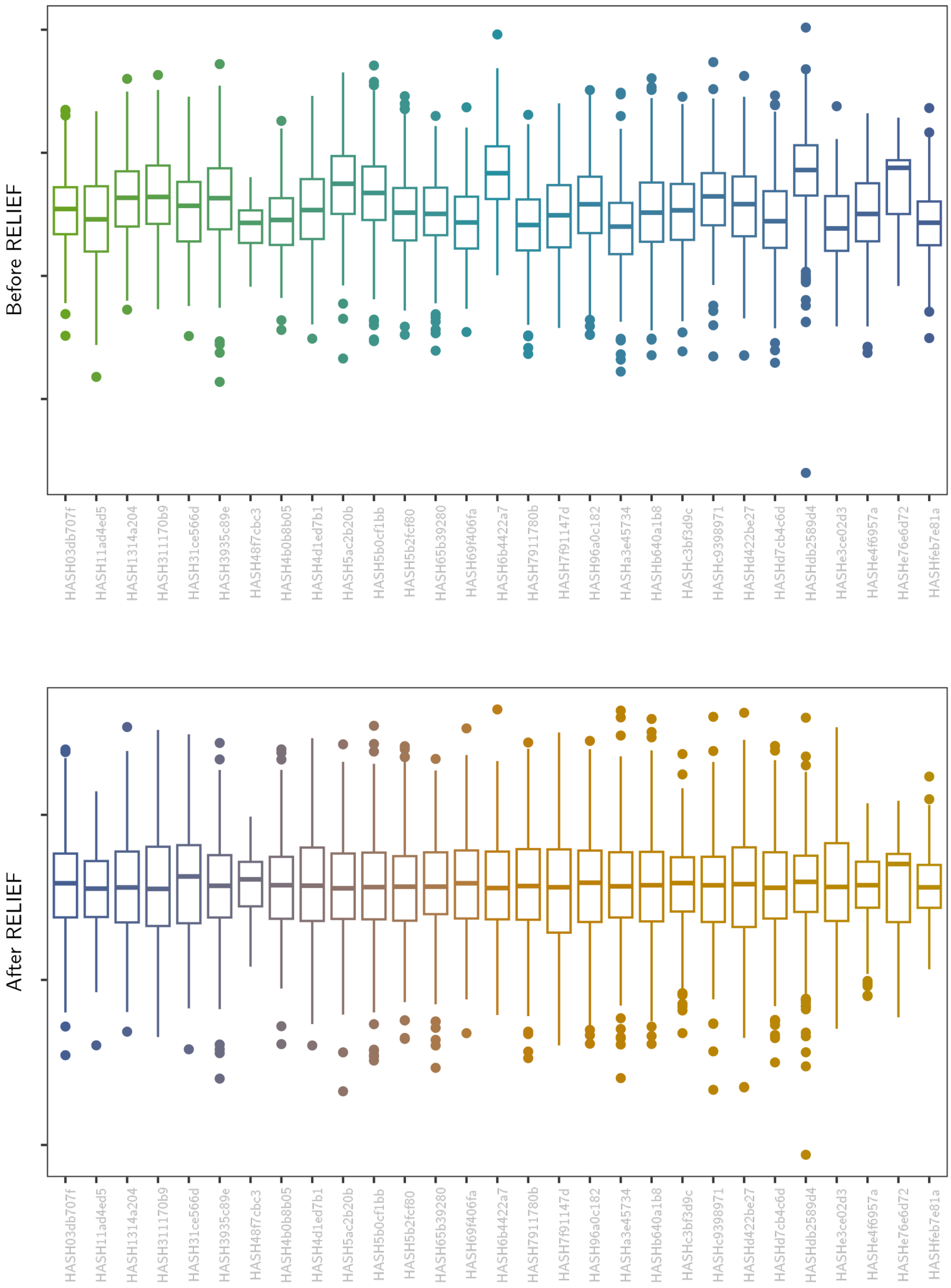

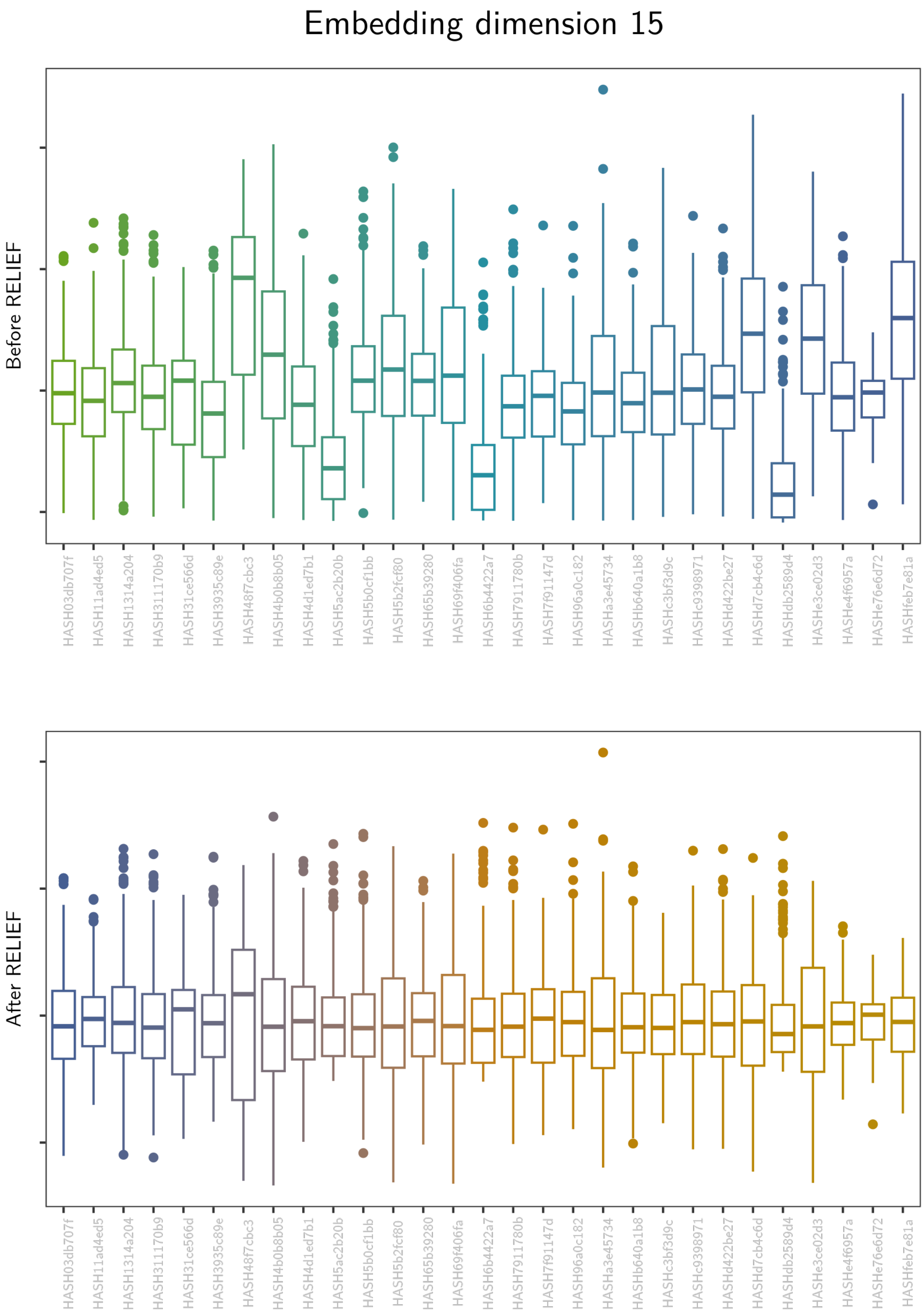
Embedding dimension 15
Before RELIEF
After RELIEF
HASH03db707f
HASH11ad4ed5
HASH1314a204
HASH311170b9
HASH31ce566d
HASH3935c89e
HASH48f7cbc3
HASH4b0b8b05
HASH4d1ed7b1
HASH5ac2b20b
HASH5b0cf1bb
HASH5b2fcf80
HASH65b39280
HASH69f406fa
HASH6b4422a7
HASH7911780b
HASH7f91147d
HASH96a0c182
HASHa3e45734
HASHb640a1b8
HASHc3bf3d9c
HASHc9398971
HASHd422be27
HASHd7cb4c6d
HASHdb2589d4
HASHe3ce02d3
HASHe4f6957a
HASHe76e6d72
HASHfeb7e81a

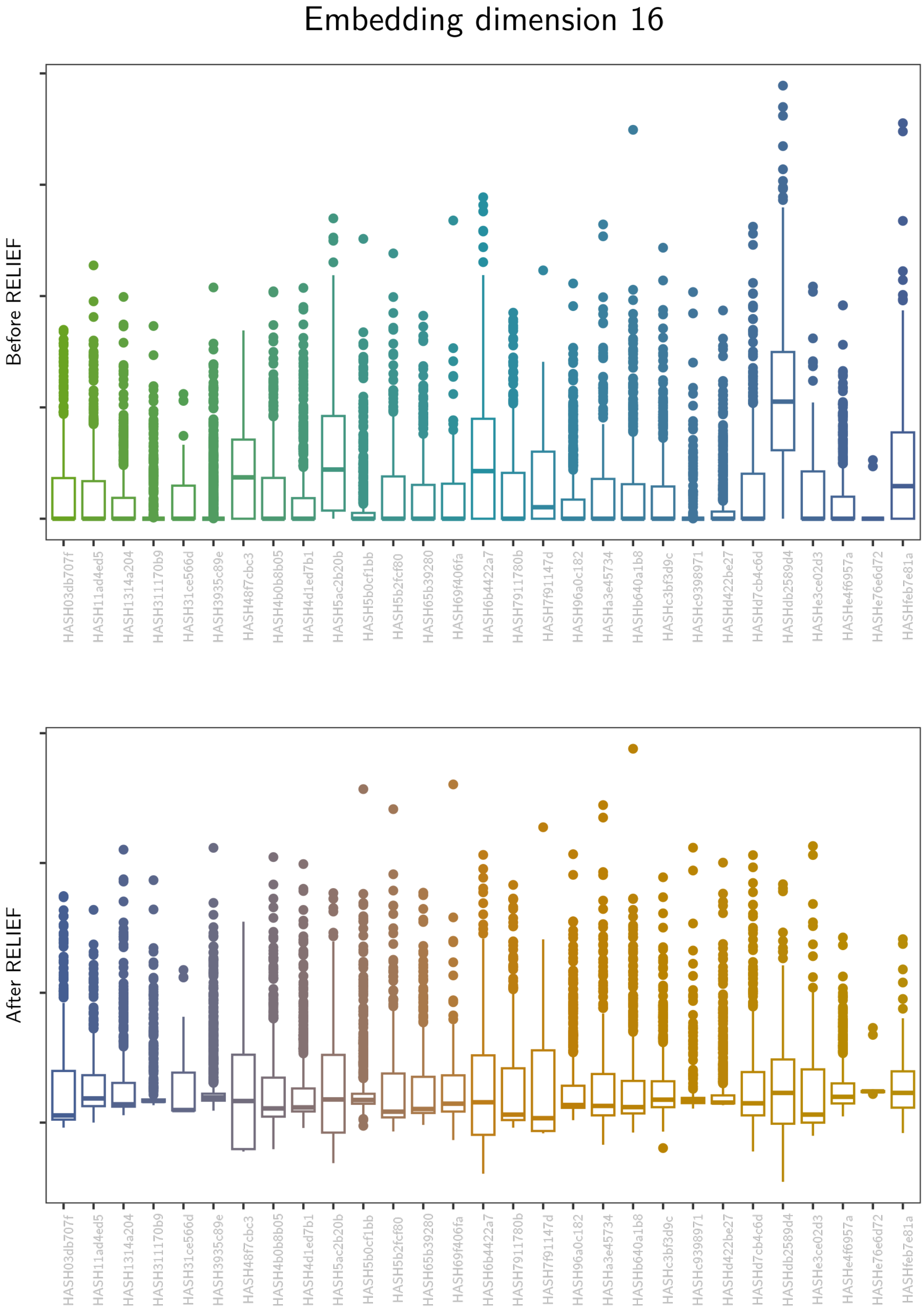
Embedding dimension 16
Before RELIEF
HASH03db707f
HASH11ad4ed5
HASH1314a204
HASH311170b9
HASH31ce566d
HASH3935c89e
HASH48f7cbc3
HASH4b0b8b05
HASH4d1ed7b1
HASH5ac2b20b
HASH5b0cf1bb
HASH5b2fcf80
HASH65b39280
HASH69f406fa
HASH6b4422a7
HASH7911780b
HASH7f91147d
HASH96a0c182
HASHa3e45734
HASHb640a1b8
HASHc3bf3d9c
HASHc9398971
HASHd422be27
HASHd7cb4c6d
HASHdb2589d4
HASHe3ce02d3
HASHe4f6957a
HASHe76e6d72
HASHfeb7e81a
After RELIEF
HASH03db707f
HASH11ad4ed5
HASH1314a204
HASH311170b9
HASH31ce566d
HASH3935c89e
HASH48f7cbc3
HASH4b0b8b05
HASH4d1ed7b1
HASH5ac2b20b
HASH5b0cf1bb
HASH5b2fcf80
HASH65b39280
HASH69f406fa
HASH6b4422a7
HASH7911780b
HASH7f91147d
HASH96a0c182
HASHa3e45734
HASHb640a1b8
HASHc3bf3d9c
HASHc9398971
HASHd422be27
HASHd7cb4c6d
HASHdb2589d4
HASHe3ce02d3
HASHe4f6957a
HASHe76e6d72
HASHfeb7e81a

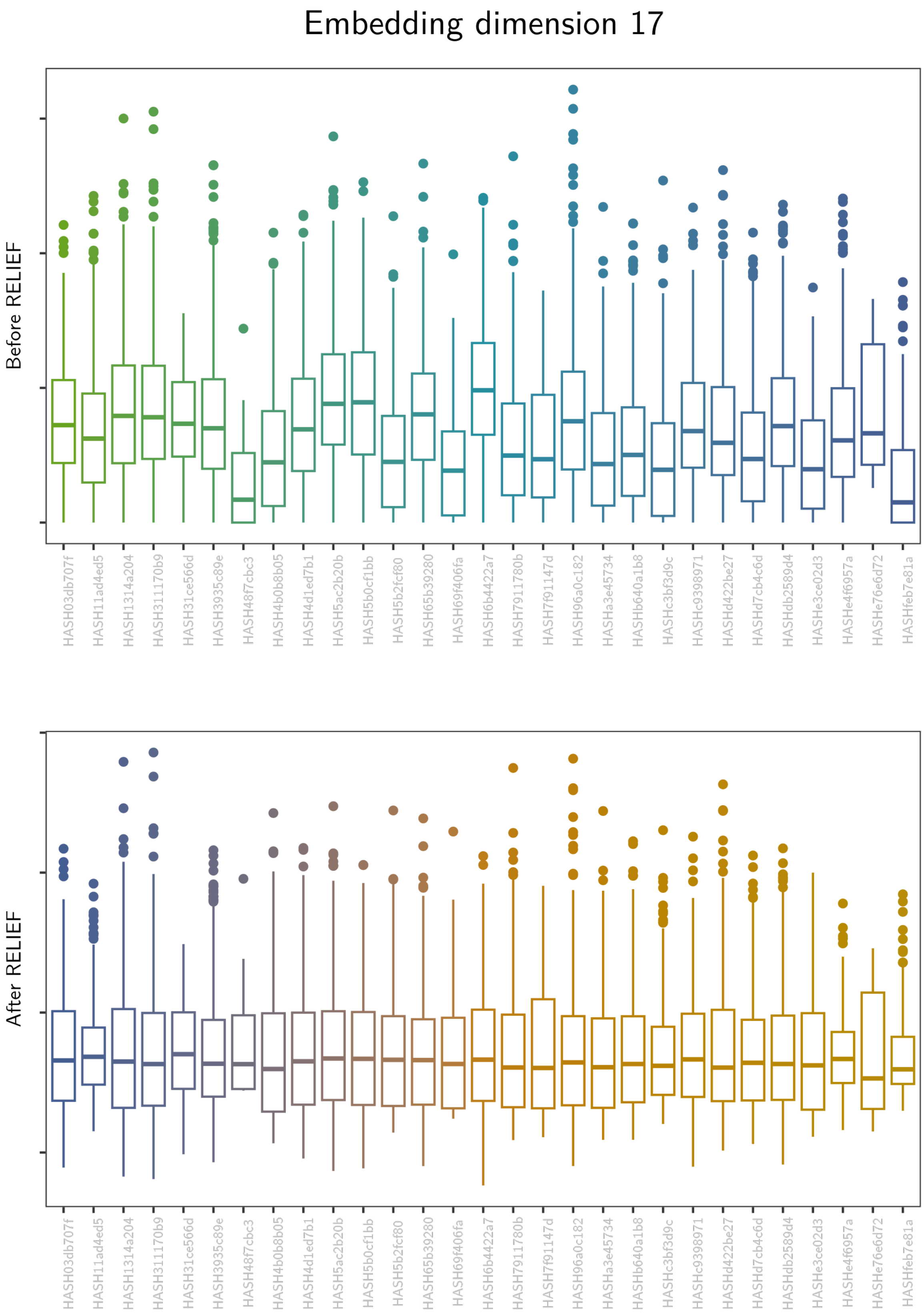
Embedding dimension 17
Before RELIEF
After RELIEF
HASH03db707f
HASH11ad4ed5
HASH1314a204
HASH311170b9
HASH31ce566d
HASH3935c89e
HASH48f7cbc3
HASH4b0b8b05
HASH4d1ed7b1
HASH5ac2b20b
HASH5b0cf1bb
HASH5b2fcf80
HASH65b39280
HASH69f406fa
HASH6b4422a7
HASH7911780b
HASH7f91147d
HASH96a0c182
HASHa3e45734
HASHb640a1b8
HASHc3bf3d9c
HASHc9398971
HASHd422be27
HASHd7cb4c6d
HASHdb2589d4
HASHe3ce02d3
HASHe4f6957a
HASHe76e6d72
HASHfeb7e81a

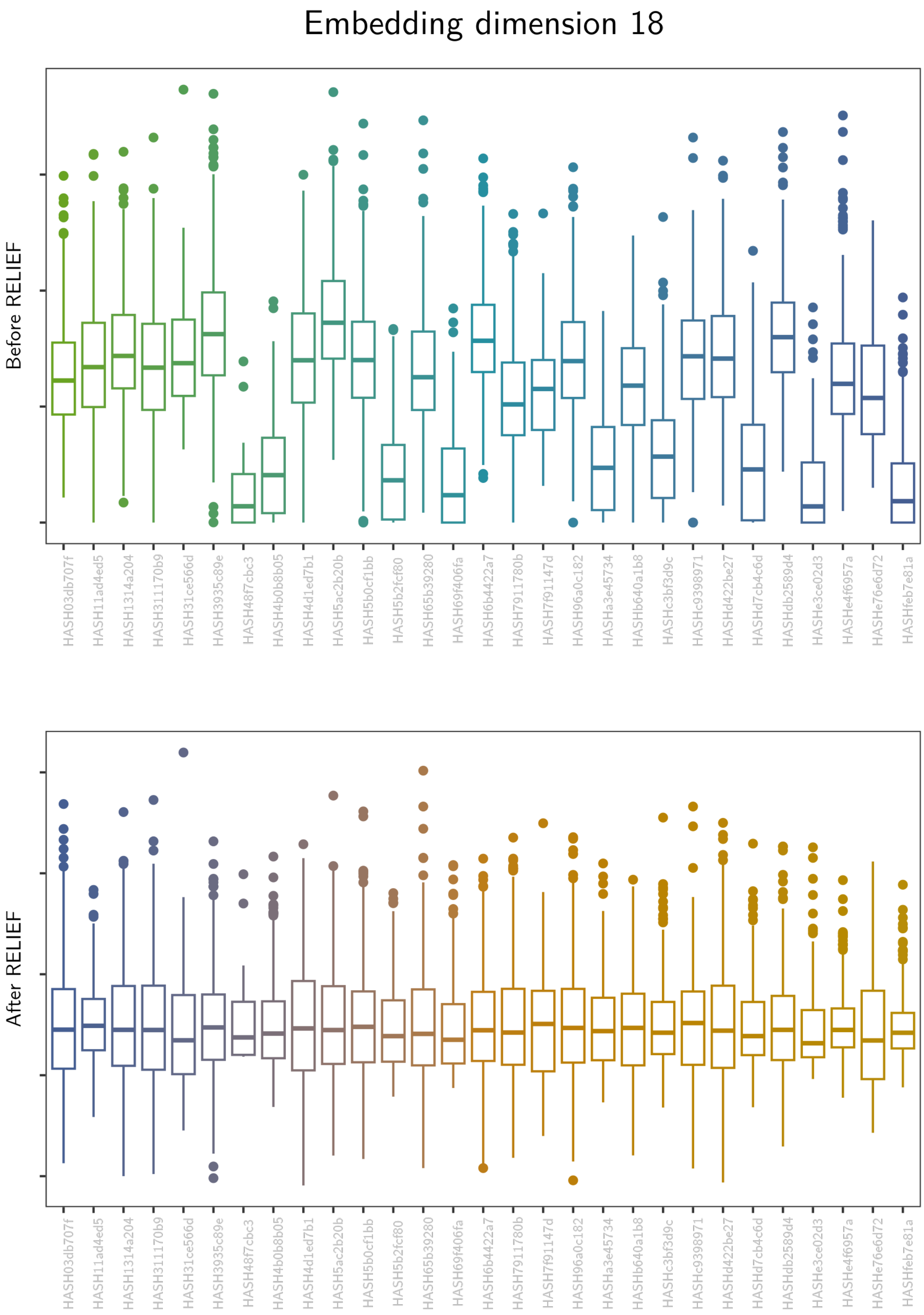
Embedding dimension 18
Before RELIEF
After RELIEF
HASH03db707f
HASH11ad4ed5
HASH1314a204
HASH311170b9
HASH31ce566d
HASH3935c89e
HASH48f7cbc3
HASH4b0b8b05
HASH4d1ed7b1
HASH5ac2b20b
HASH5b0cf1bb
HASH5b2fcf80
HASH65b39280
HASH69f406fa
HASH6b4422a7
HASH7911780b
HASH7f91147d
HASH96a0c182
HASHa3e45734
HASHb640a1b8
HASHc3bf3d9c
HASHc9398971
HASHd422be27
HASHd7cb4c6d
HASHdb2589d4
HASHe3ce02d3
HASHe4f6957a
HASHe76e6d72
HASHfeb7e81a

#### Embedding dimension 19

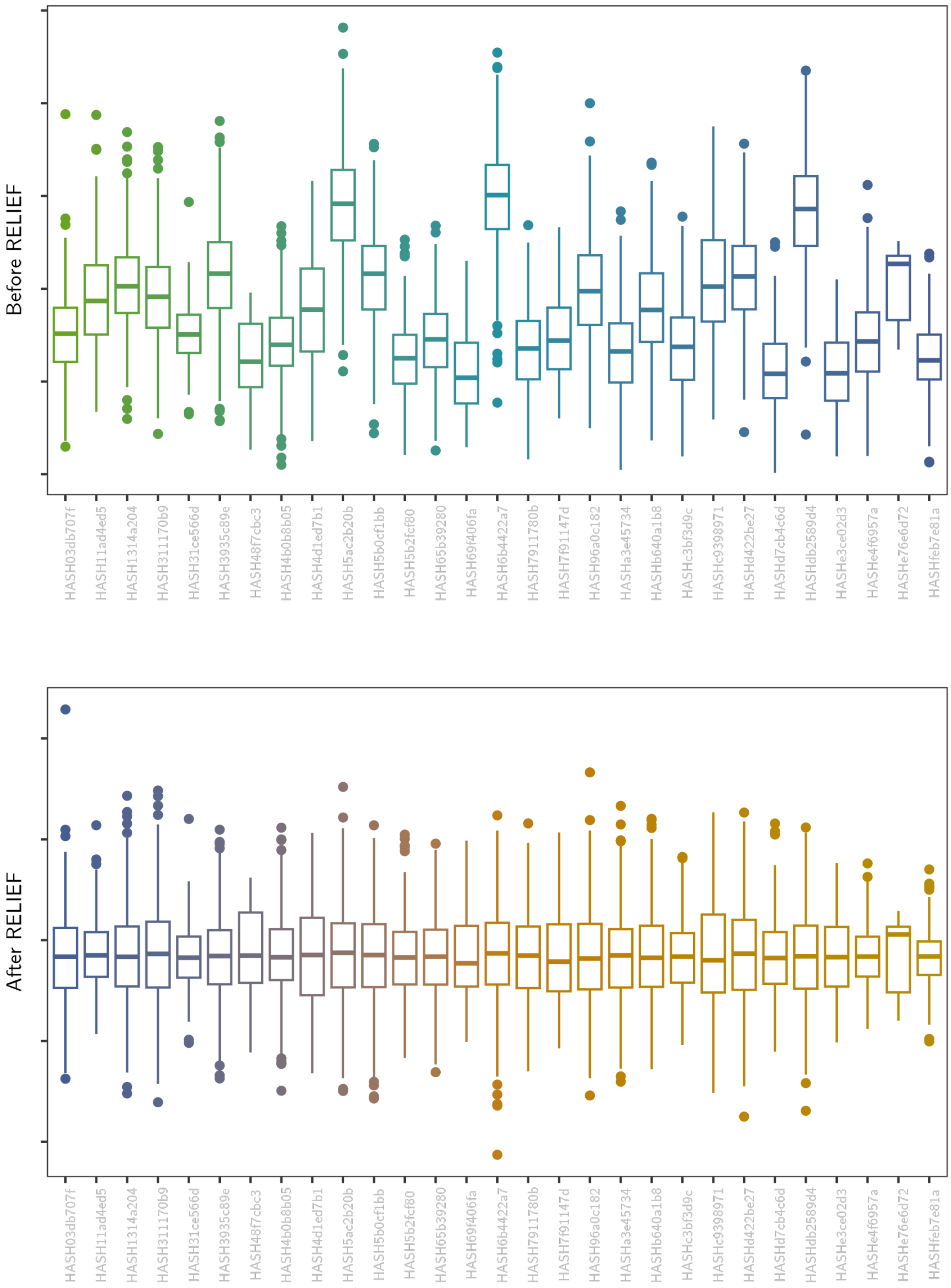

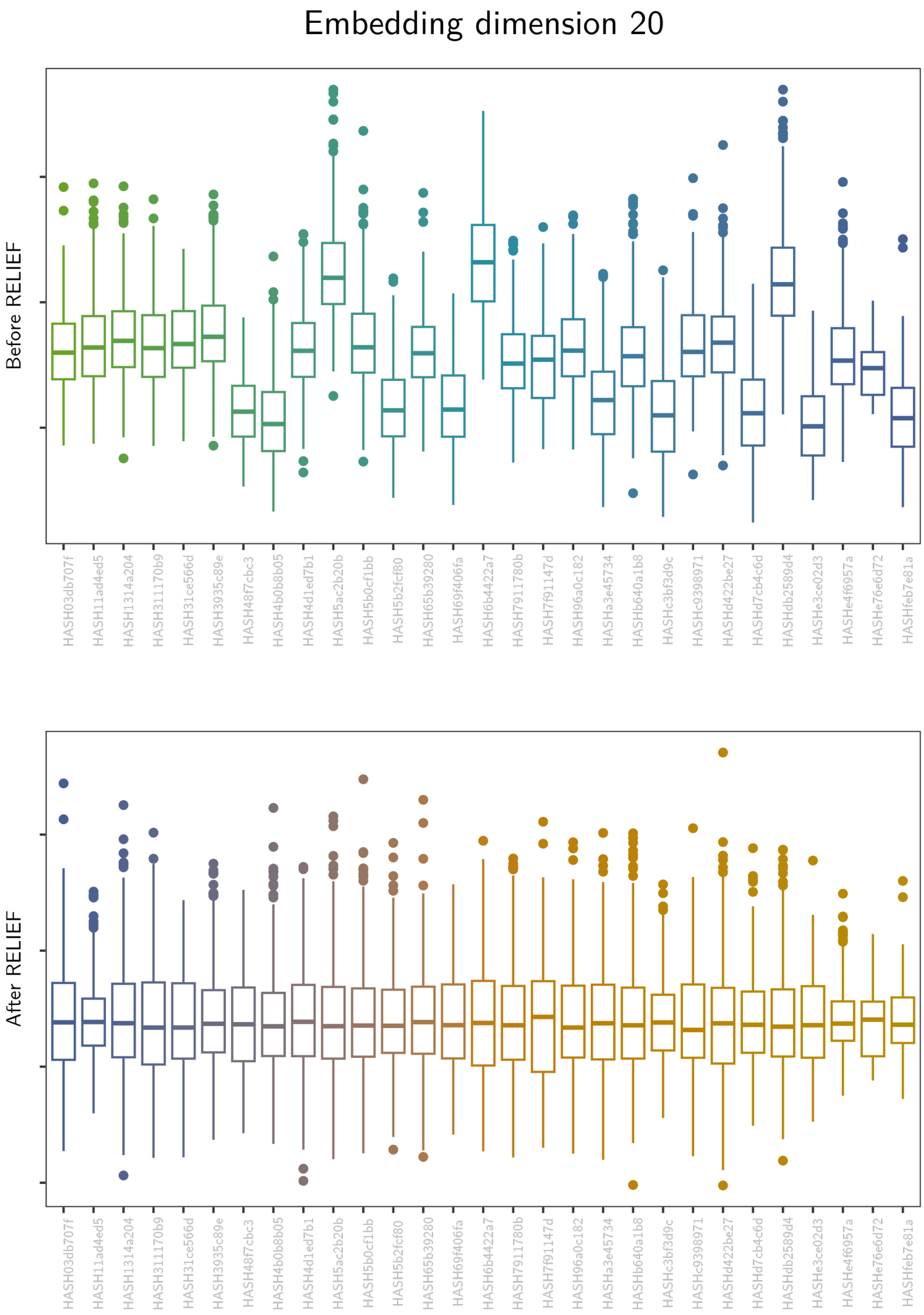
Embedding dimension 20
Before RELIEF
After RELIEF
HASH03db707f
HASH11ad4ed5
HASH1314a204
HASH311170b9
HASH31ce566d
HASH3935c89e
HASH48f7cbc3
HASH4b0b8b05
HASH4d1ed7b1
HASH5ac2b20b
HASH5b0cf1bb
HASH5b2fcf80
HASH65b39280
HASH69f406fa
HASH6b4422a7
HASH7911780b
HASH7f91147d
HASH96a0c182
HASHa3e45734
HASHb640a1b8
HASHc3bf3d9c
HASHc9398971
HASHd422be27
HASHd7cb4c6d
HASHdb2589d4
HASHe3ce02d3
HASHe4f6957a
HASHe76e6d72
HASHfeb7e81a

Embedding dimension 21

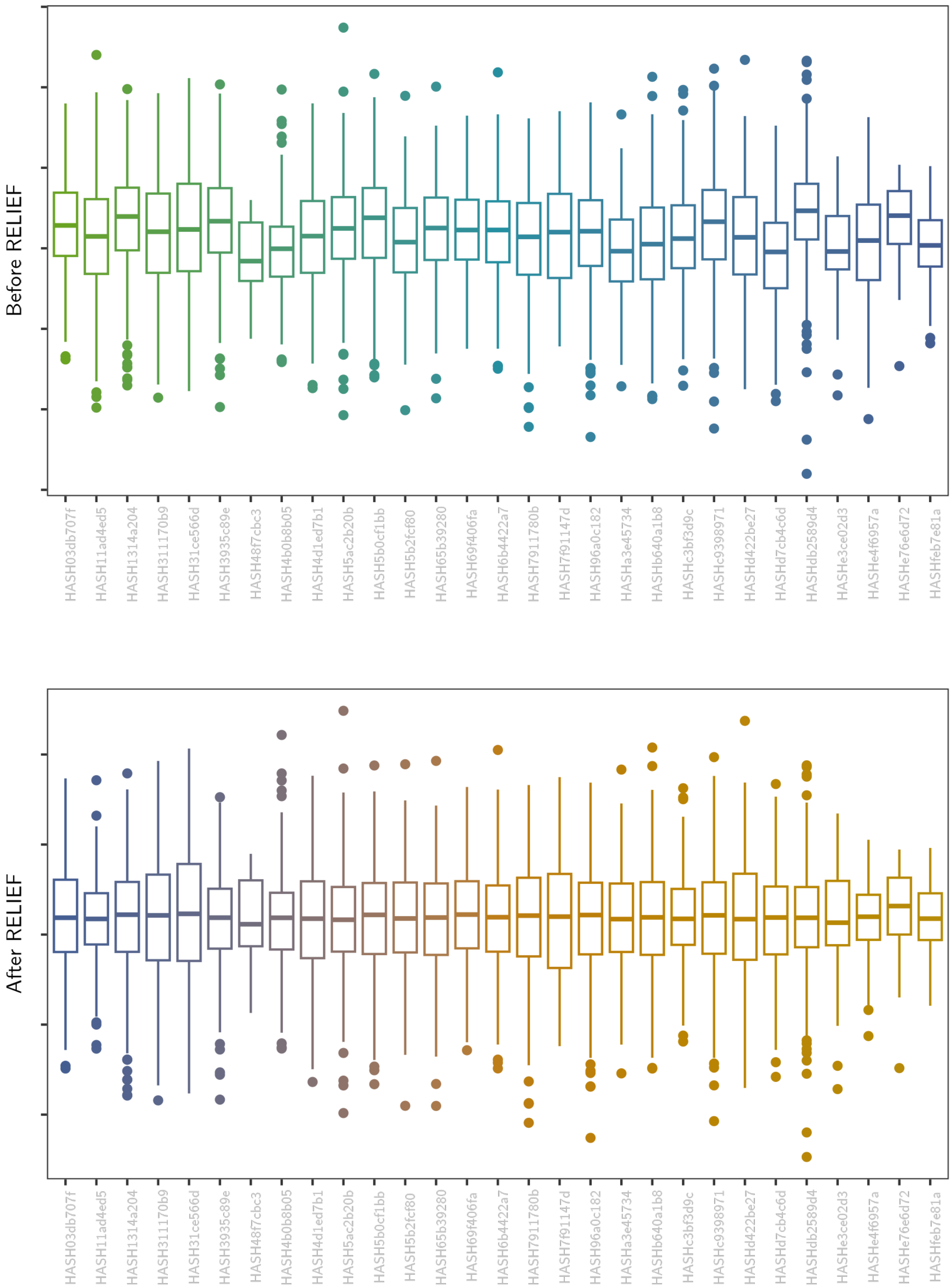

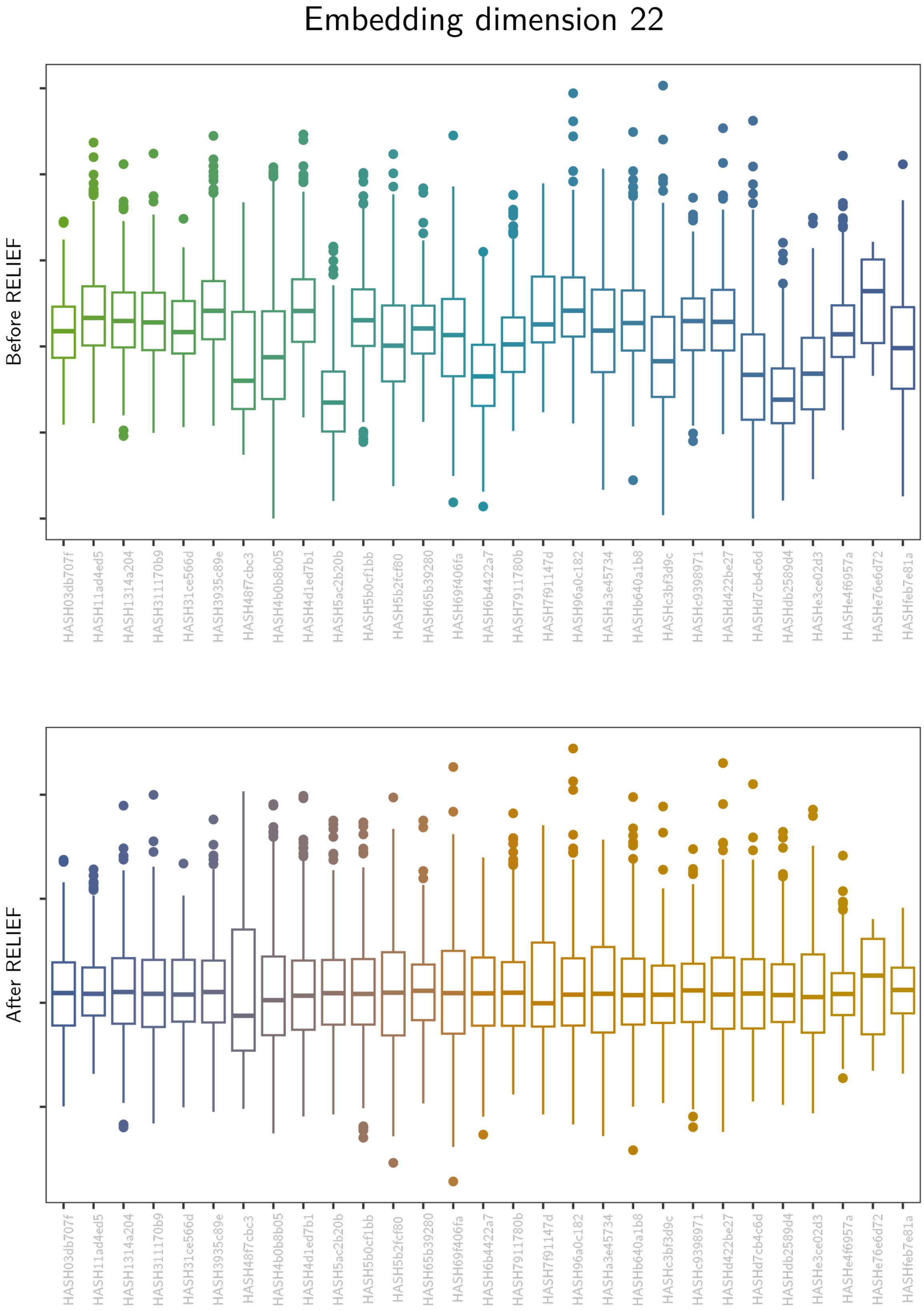
Embedding dimension 22
Before RELIEF
HASH03db707f
HASH11ad4ed5
HASH1314a204
HASH311170b9
HASH31ce566d
HASH3935c89e
HASH48f7cbc3
HASH4b0b8b05
HASH4d1ed7b1
HASH5ac2b20b
HASH5b0cf1bb
HASH5b2fcf80
HASH65b39280
HASH69f406fa
HASH6b4422a7
HASH7911780b
HASH7f91147d
HASH96a0c182
HASHa3e45734
HASHb640a1b8
HASHc3bf3d9c
HASHc9398971
HASHd422be27
HASHd7cb4c6d
HASHdb2589d4
HASHe3ce02d3
HASHe4f6957a
HASHe76e6d72
HASHfeb7e81a
After RELIEF
HASH03db707f
HASH11ad4ed5
HASH1314a204
HASH311170b9
HASH31ce566d
HASH3935c89e
HASH48f7cbc3
HASH4b0b8b05
HASH4d1ed7b1
HASH5ac2b20b
HASH5b0cf1bb
HASH5b2fcf80
HASH65b39280
HASH69f406fa
HASH6b4422a7
HASH7911780b
HASH7f91147d
HASH96a0c182
HASHa3e45734
HASHb640a1b8
HASHc3bf3d9c
HASHc9398971
HASHd422be27
HASHd7cb4c6d
HASHdb2589d4
HASHe3ce02d3
HASHe4f6957a
HASHe76e6d72
HASHfeb7e81a

#### Embedding dimension 23

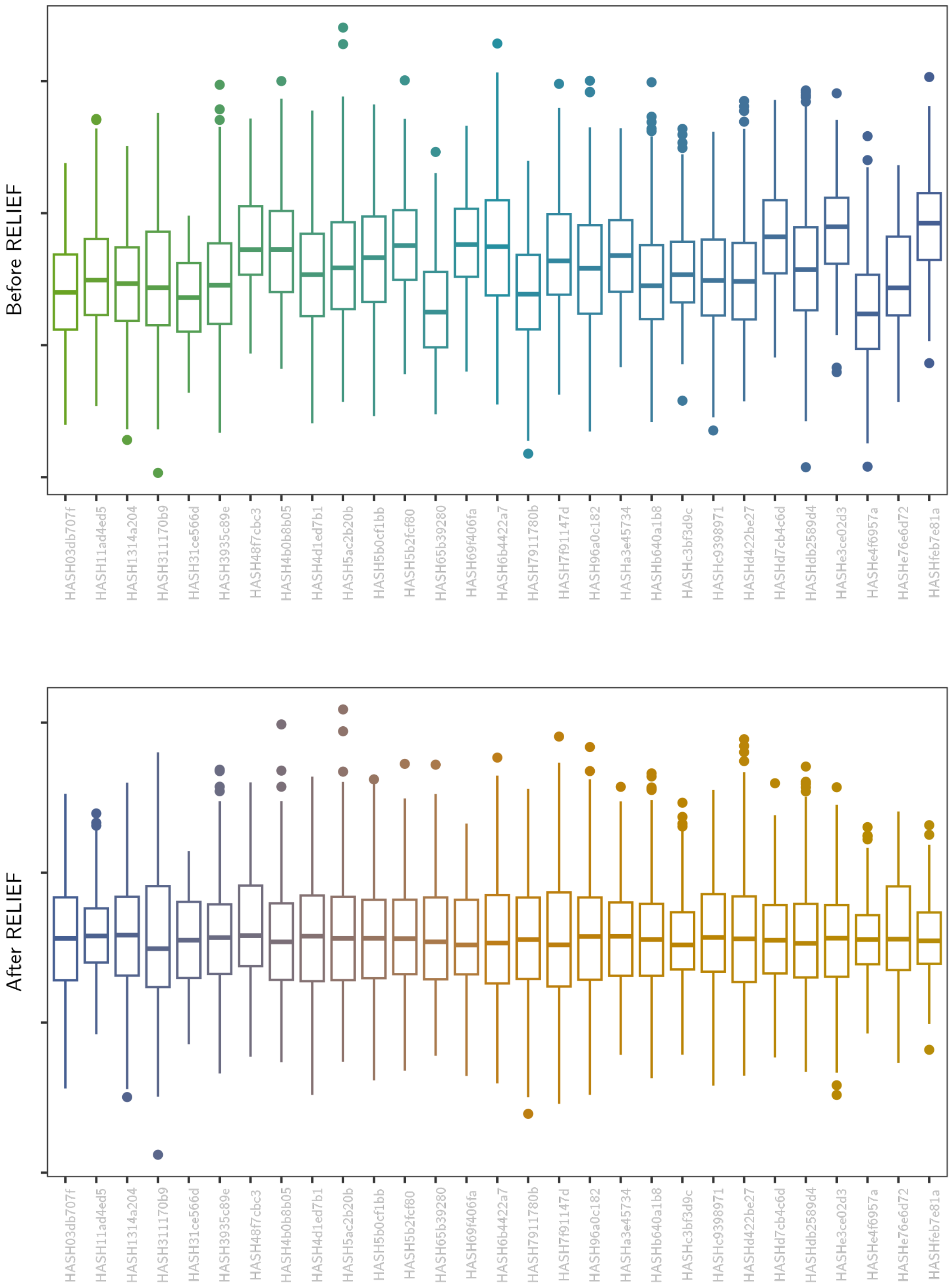

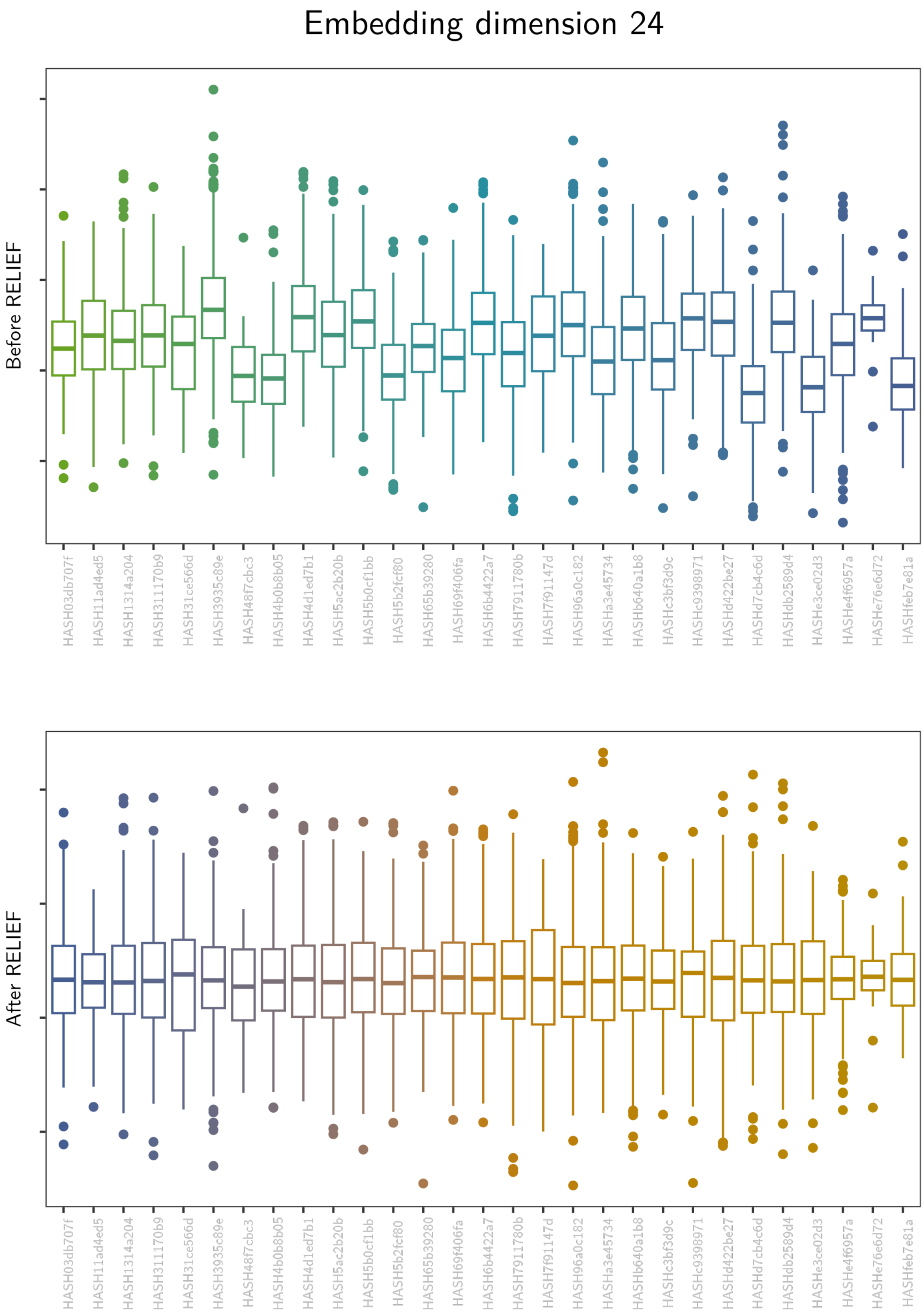
Embedding dimension 24
Before RELIEF
After RELIEF
HASH03db707f
HASH11ad4ed5
HASH1314a204
HASH311170b9
HASH31ce566d
HASH3935c89e
HASH48f7cbc3
HASH4b0b8b05
HASH4d1ed7b1
HASH5ac2b20b
HASH5b0cf1bb
HASH5b2fcf80
HASH65b39280
HASH69f406fa
HASH6b4422a7
HASH7911780b
HASH7f91147d
HASH96a0c182
HASHa3e45734
HASHb640a1b8
HASHc3bf3d9c
HASHc9398971
HASHd422be27
HASHd7cb4c6d
HASHdb2589d4
HASHe3ce02d3
HASHe4f6957a
HASHe76e6d72
HASHfeb7e81a

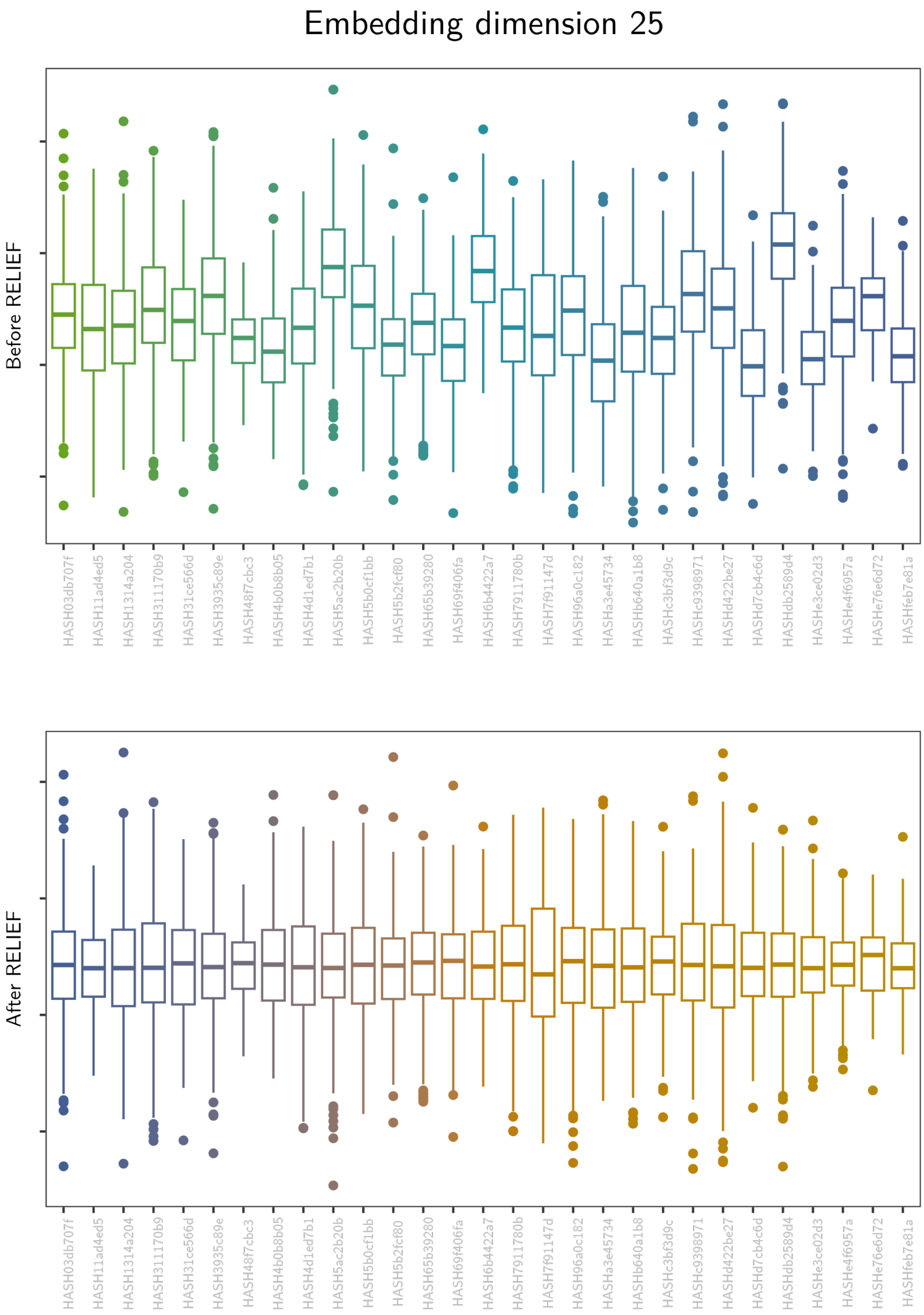
Embedding dimension 25
Before RELIEF
HASH03db707f
HASH11ad4ed5
HASH1314a204
HASH311170b9
HASH31ce566d
HASH3935c89e
HASH48f7cbc3
HASH4b0b8b05
HASH4d1ed7b1
HASH5ac2b20b
HASH5b0cf1bb
HASH5b2fcf80
HASH65b39280
HASH69f406fa
HASH6b4422a7
HASH7911780b
HASH7f91147d
HASH96a0c182
HASHa3e45734
HASHb640a1b8
HASHc3bf3d9c
HASHc9398971
HASHd422be27
HASHd7cb4c6d
HASHdb2589d4
HASHe3ce02d3
HASHe4f6957a
HASHe76e6d72
HASHfeb7e81a
After RELIEF
HASH03db707f
HASH11ad4ed5
HASH1314a204
HASH311170b9
HASH31ce566d
HASH3935c89e
HASH48f7cbc3
HASH4b0b8b05
HASH4d1ed7b1
HASH5ac2b20b
HASH5b0cf1bb
HASH5b2fcf80
HASH65b39280
HASH69f406fa
HASH6b4422a7
HASH7911780b
HASH7f91147d
HASH96a0c182
HASHa3e45734
HASHb640a1b8
HASHc3bf3d9c
HASHc9398971
HASHd422be27
HASHd7cb4c6d
HASHdb2589d4
HASHe3ce02d3
HASHe4f6957a
HASHe76e6d72
HASHfeb7e81a

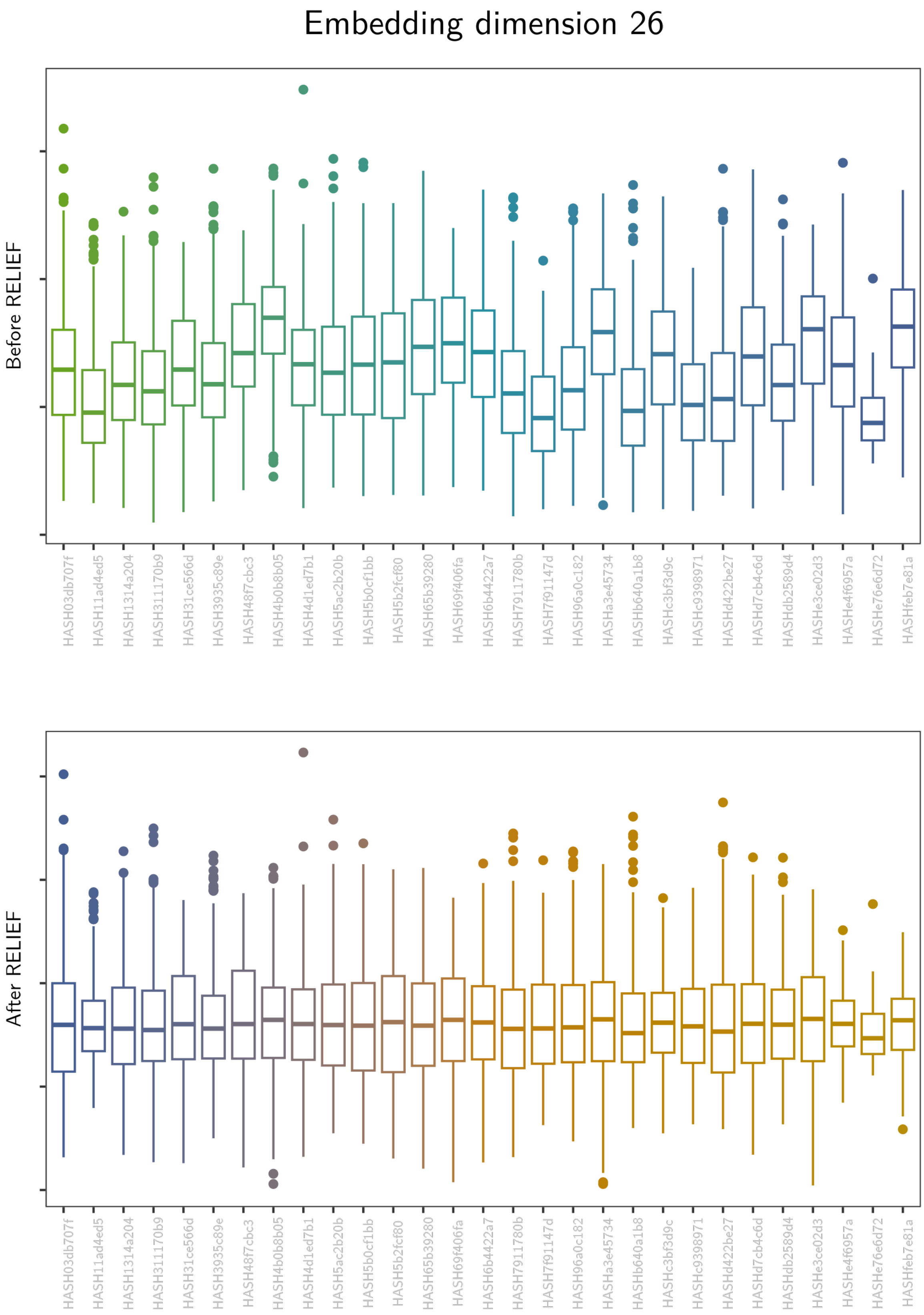

Embedding dimension 26
Before RELIEF
After RELIEF
HASH03db707f
HASH11ad4ed5
HASH1314a204
HASH311170b9
HASH31ce566d
HASH3935c89e
HASH48f7cbc3
HASH4b0b8b05
HASH4d1ed7b1
HASH5ac2b20b
HASH5b0cf1bb
HASH5b2fcf80
HASH65b39280
HASH69f406fa
HASH6b4422a7
HASH7911780b
HASH7f91147d
HASH96a0c182
HASHa3e45734
HASHb640a1b8
HASHc3bf3d9c
HASHc9398971
HASHd422be27
HASHd7cb4c6d
HASHdb2589d4
HASHe3ce02d3
HASHe4f6957a
HASHe76e6d72
HASHfeb7e81a

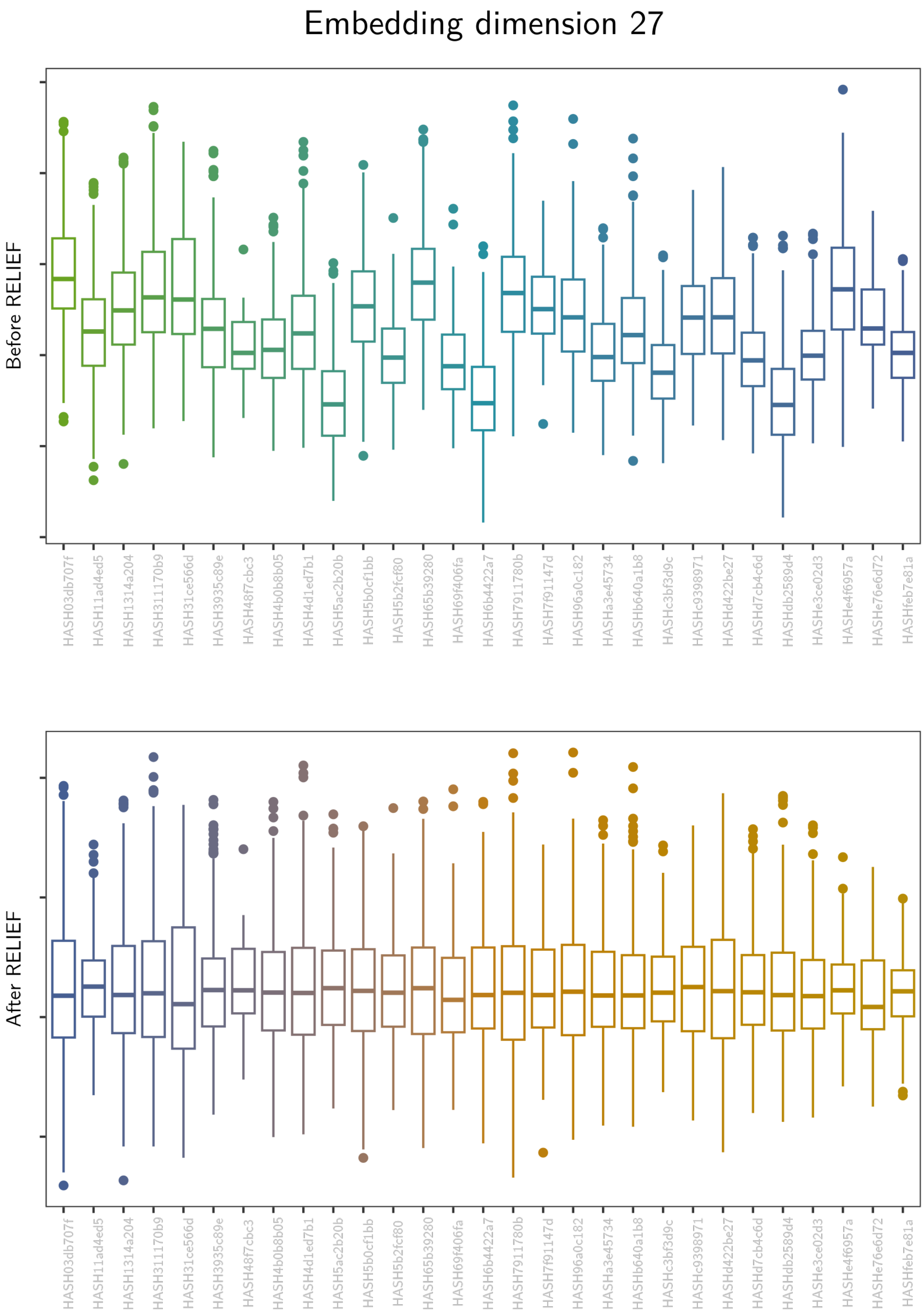
Embedding dimension 27
Before RELIEF
After RELIEF
HASH03db707f
HASH11ad4ed5
HASH1314a204
HASH311170b9
HASH31ce566d
HASH3935c89e
HASH48f7cbc3
HASH4b0b8b05
HASH4d1ed7b1
HASH5ac2b20b
HASH5b0cf1bb
HASH5b2fcf80
HASH65b39280
HASH69f406fa
HASH6b4422a7
HASH7911780b
HASH7f91147d
HASH96a0c182
HASHa3e45734
HASHb640a1b8
HASHc3bf3d9c
HASHc9398971
HASHd422be27
HASHd7cb4c6d
HASHdb2589d4
HASHe3ce02d3
HASHe4f6957a
HASHe76e6d72
HASHfeb7e81a

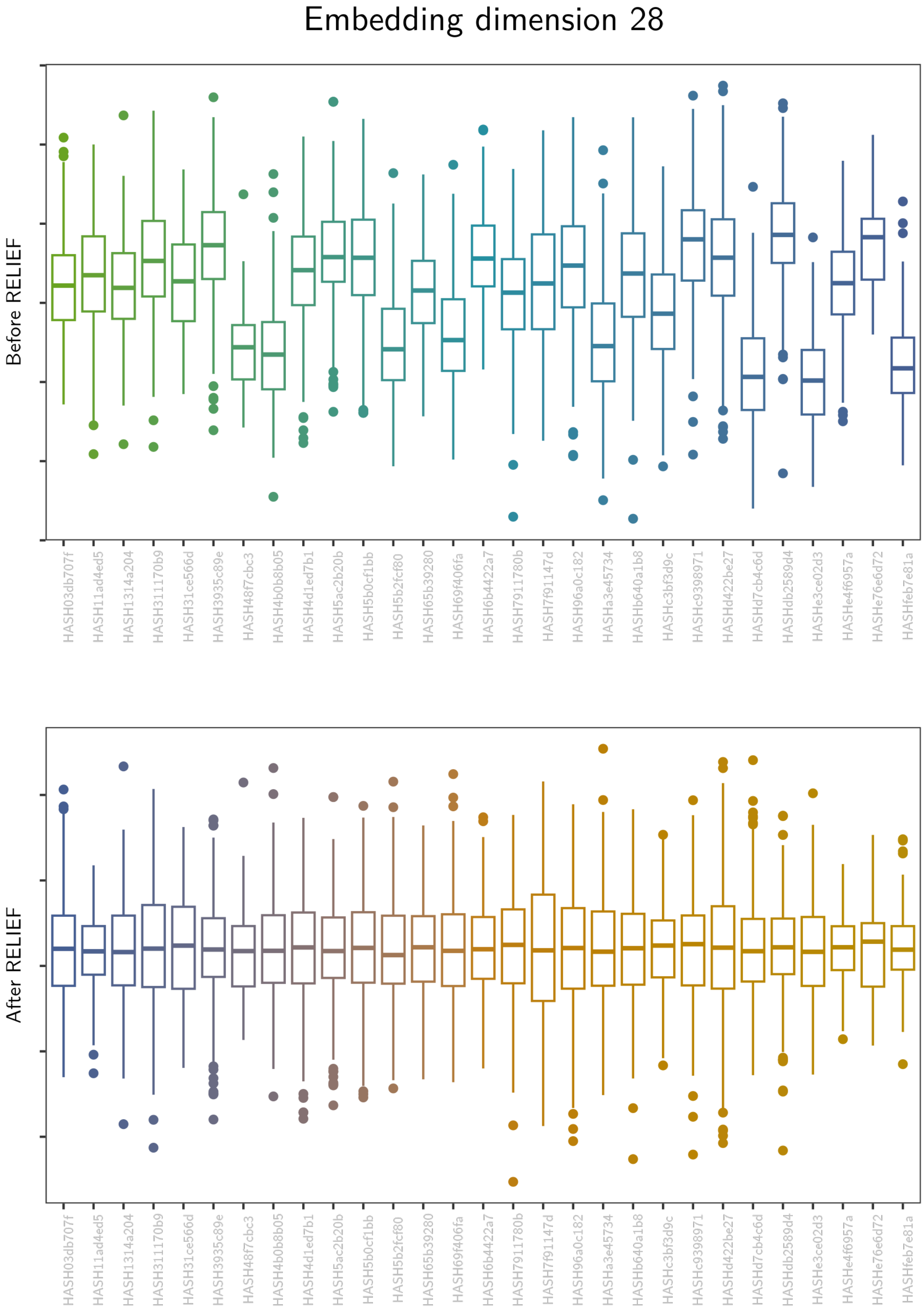
Embedding dimension 28
Before RELIEF
After RELIEF
HASH03db707f
HASH11ad4ed5
HASH1314a204
HASH311170b9
HASH31ce566d
HASH3935c89e
HASH48f7cbc3
HASH4b0b8b05
HASH4d1ed7b1
HASH5ac2b20b
HASH5b0cf1bb
HASH5b2fcf80
HASH65b39280
HASH69f406fa
HASH6b4422a7
HASH7911780b
HASH7f91147d
HASH96a0c182
HASHa3e45734
HASHb640a1b8
HASHc3bf3d9c
HASHc9398971
HASHd422be27
HASHd7cb4c6d
HASHdb2589d4
HASHe3ce02d3
HASHe4f6957a
HASHe76e6d72
HASHfeb7e81a

### Embedding dimension 29

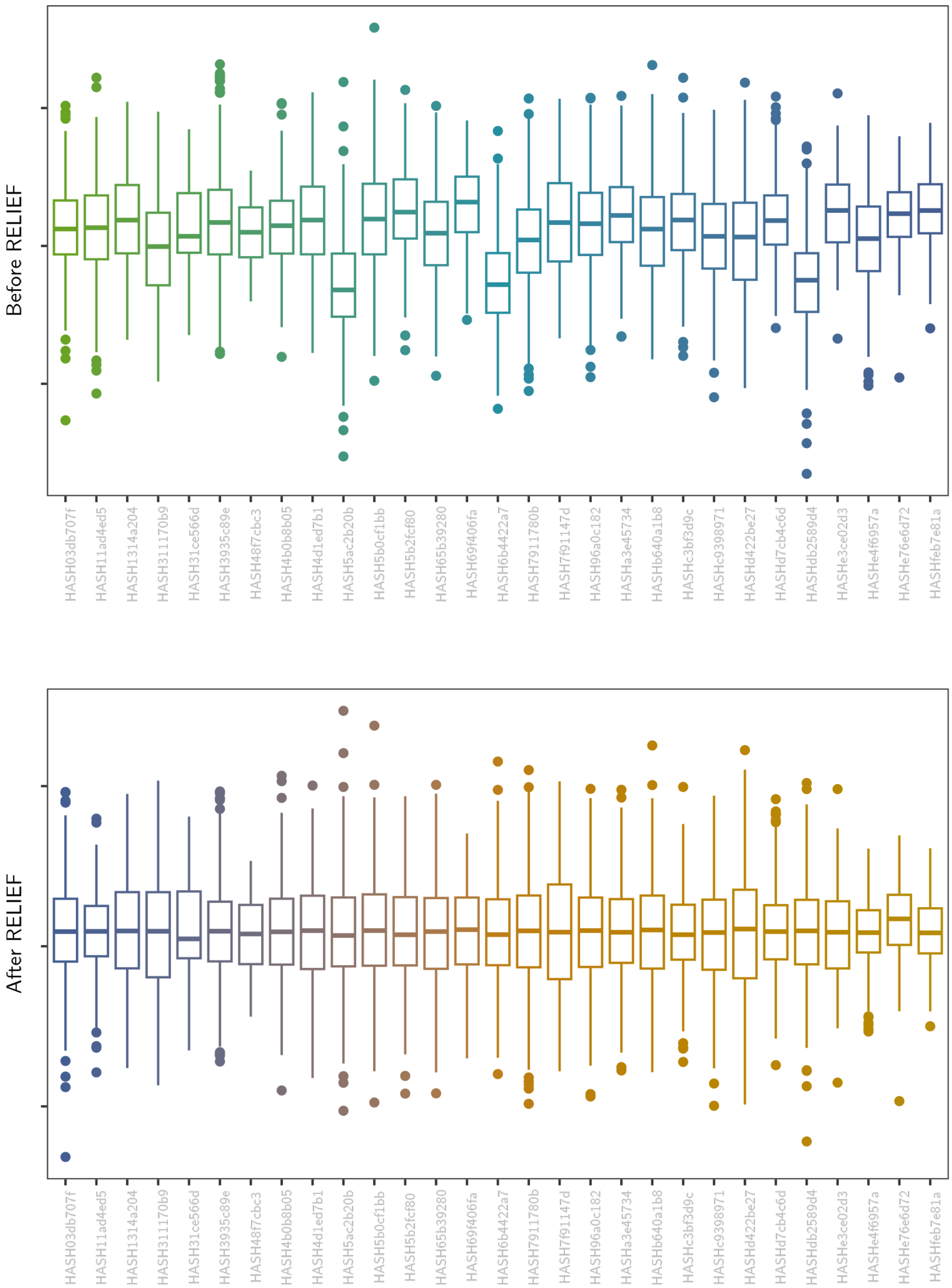

#### Embedding dimension 30

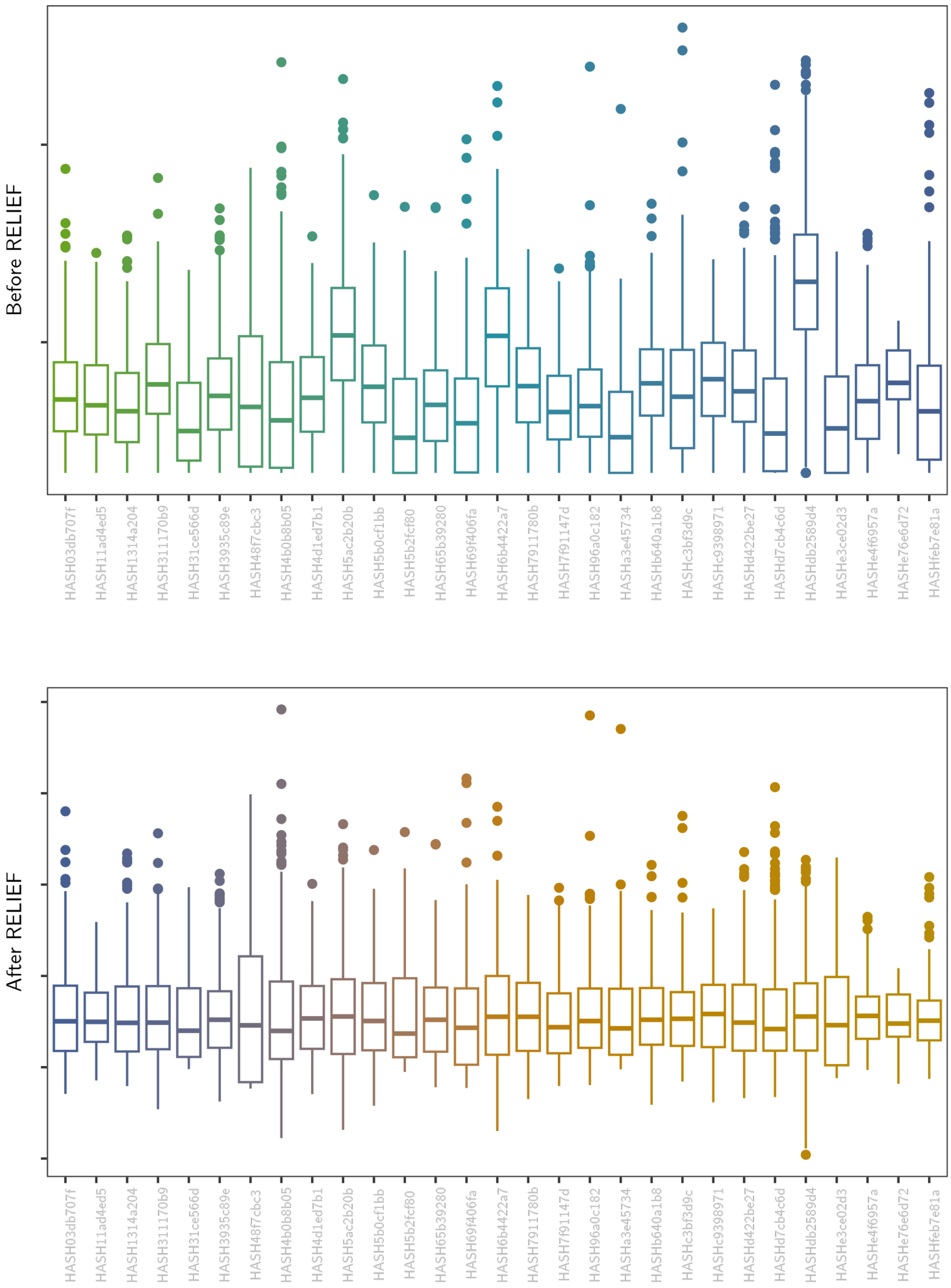

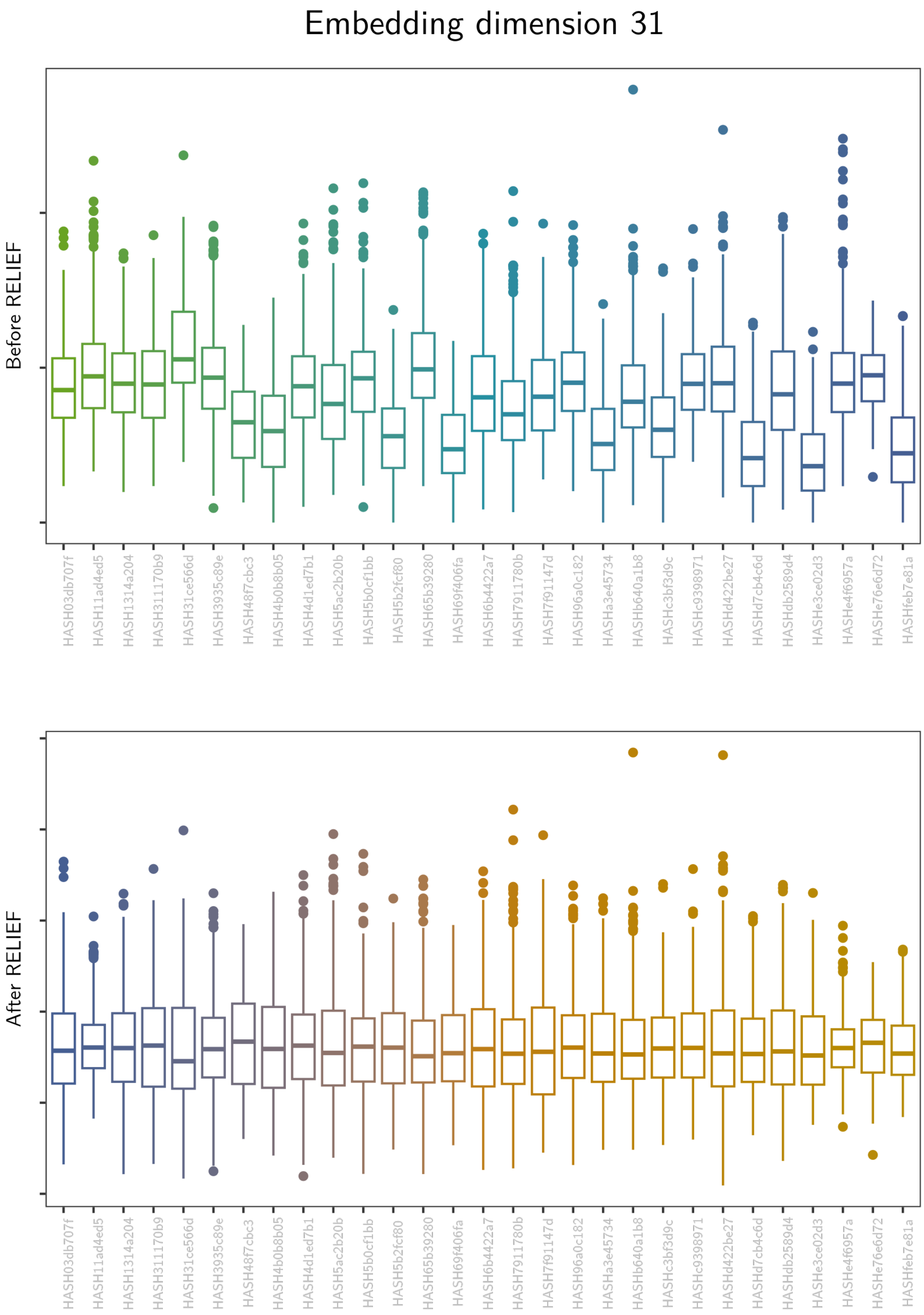
Embedding dimension 31
Before RELIEF
After RELIEF
HASH03db707f
HASH11ad4ed5
HASH1314a204
HASH311170b9
HASH31ce566d
HASH3935c89e
HASH48f7cbc3
HASH4b0b8b05
HASH4d1ed7b1
HASH5ac2b20b
HASH5b0cf1bb
HASH5b2fcf80
HASH65b39280
HASH69f406fa
HASH6b4422a7
HASH7911780b
HASH7f91147d
HASH96a0c182
HASHa3e45734
HASHb640a1b8
HASHc3bf3d9c
HASHc9398971
HASHd422be27
HASHd7cb4c6d
HASHdb2589d4
HASHe3ce02d3
HASHe4f6957a
HASHe76e6d72
HASHfeb7e81a

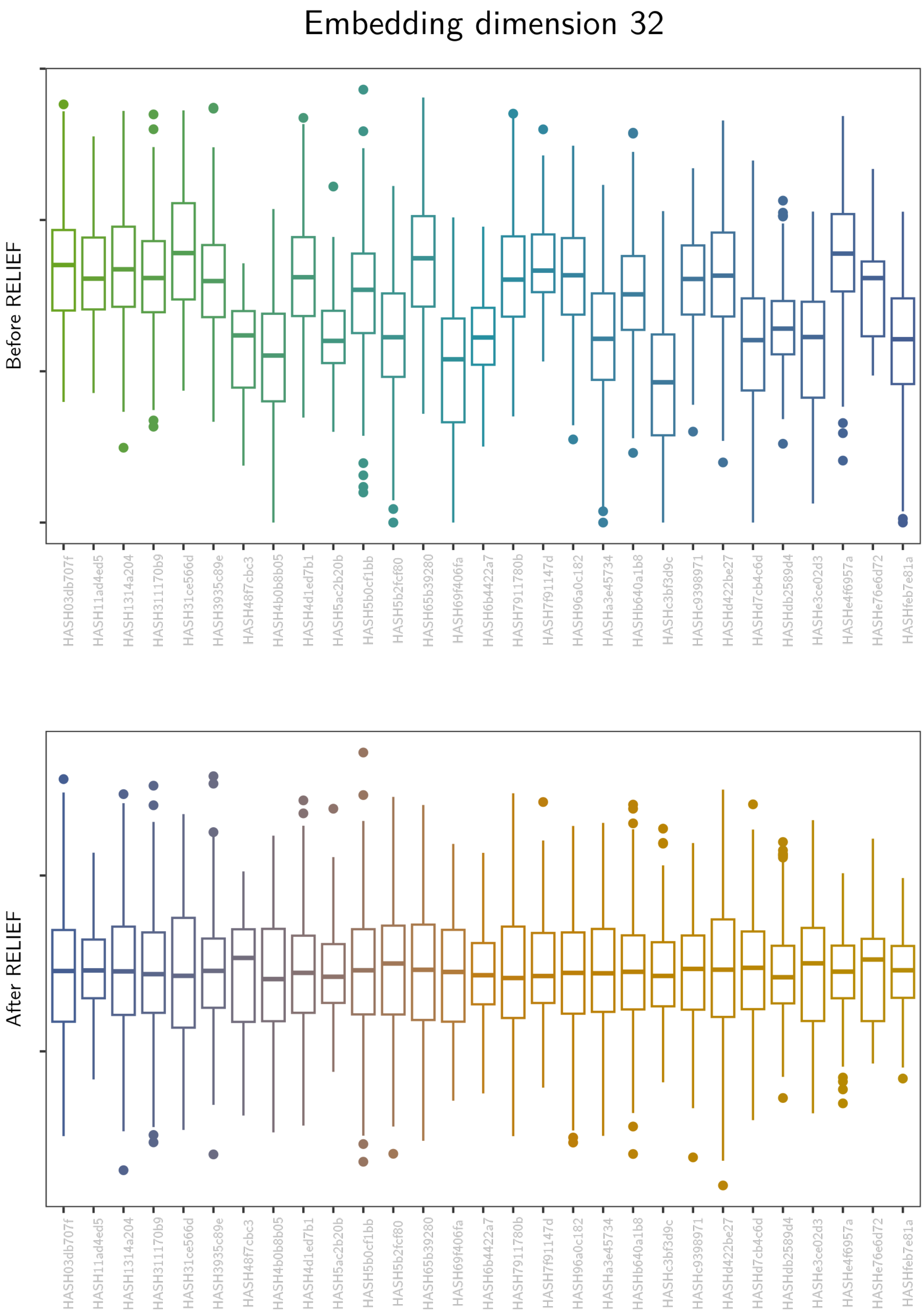
Embedding dimension 32
Before RELIEF
After RELIEF
HASH03db707f
HASH11ad4ed5
HASH1314a204
HASH311170b9
HASH31ce566d
HASH3935c89e
HASH48f7cbc3
HASH4b0b8b05
HASH4d1ed7b1
HASH5ac2b20b
HASH5b0cf1bb
HASH5b2fcf80
HASH65b39280
HASH69f406fa
HASH6b4422a7
HASH7911780b
HASH7f91147d
HASH96a0c182
HASHa3e45734
HASHb640a1b8
HASHc3bf3d9c
HASHc9398971
HASHd422be27
HASHd7cb4c6d
HASHdb2589d4
HASHe3ce02d3
HASHe4f6957a
HASHe76e6d72
HASHfeb7e81a

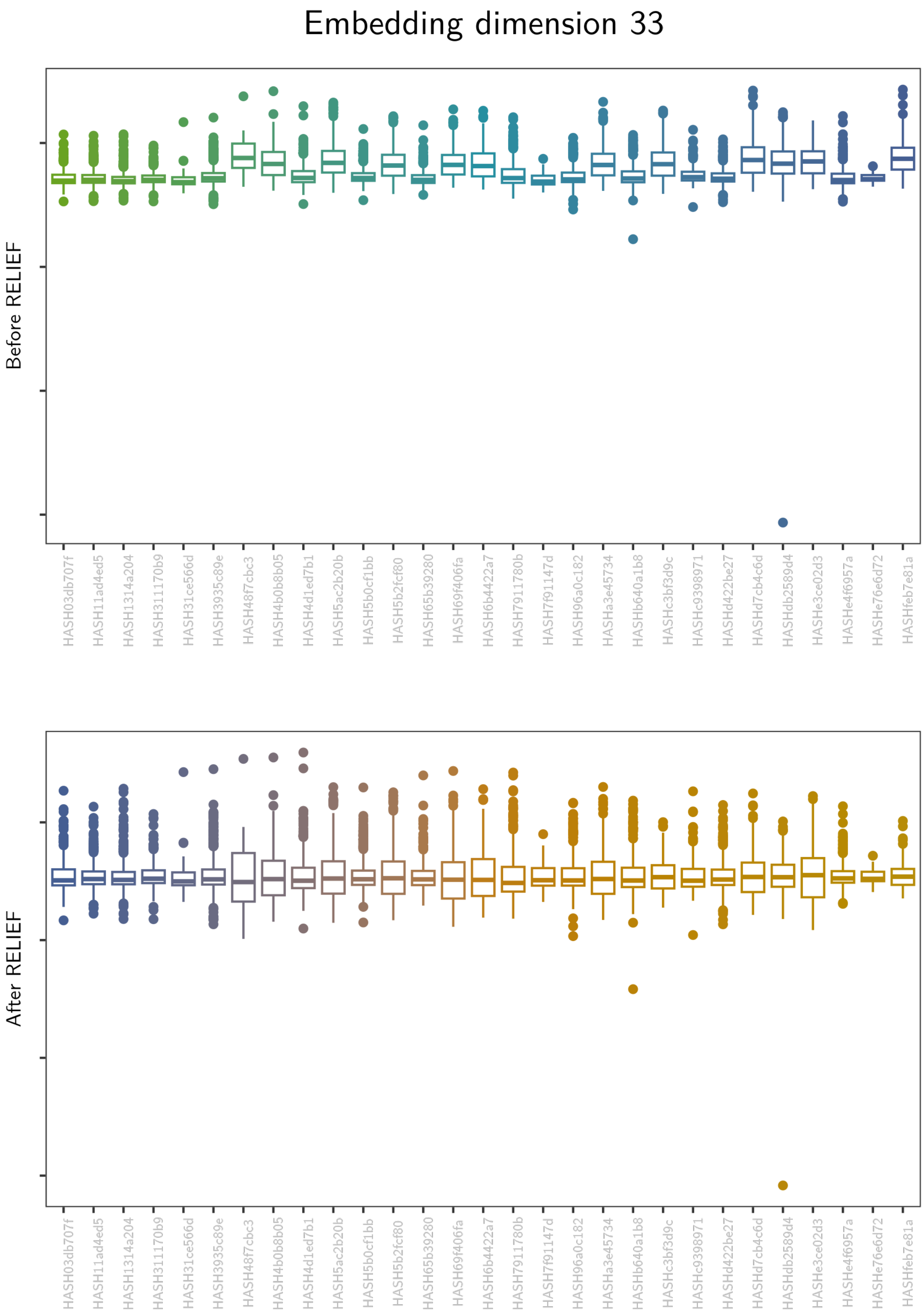
Embedding dimension 33
Before RELIEF
After RELIEF
HASH03db707f
HASH11ad4ed5
HASH1314a204
HASH311170b9
HASH31ce566d
HASH3935c89e
HASH48f7cbc3
HASH4b0b8b05
HASH4d1ed7b1
HASH5ac2b20b
HASH5b0cf1bb
HASH5b2fcf80
HASH65b39280
HASH69f406fa
HASH6b4422a7
HASH7911780b
HASH7f91147d
HASH96a0c182
HASHa3e45734
HASHb640a1b8
HASHc3bf3d9c
HASHc9398971
HASHd422be27
HASHd7cb4c6d
HASHdb2589d4
HASHe3ce02d3
HASHe4f6957a
HASHe76e6d72
HASHfeb7e81a

Embedding dimension 34

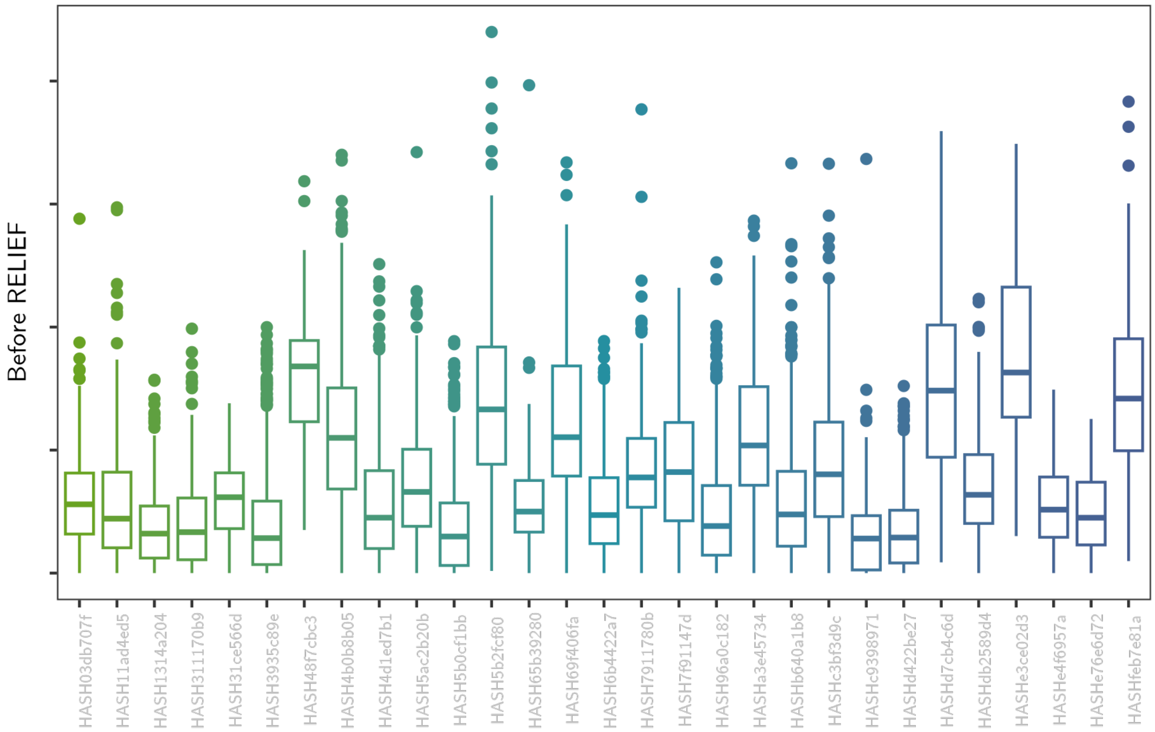

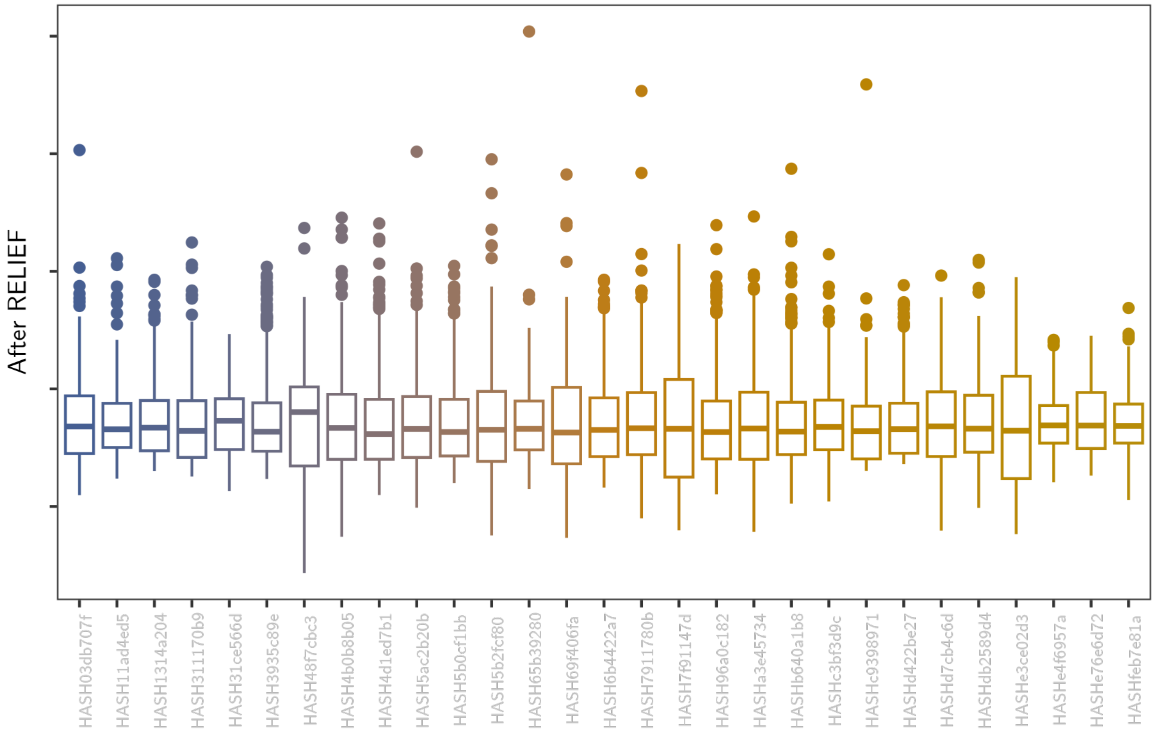

Embedding dimension 35

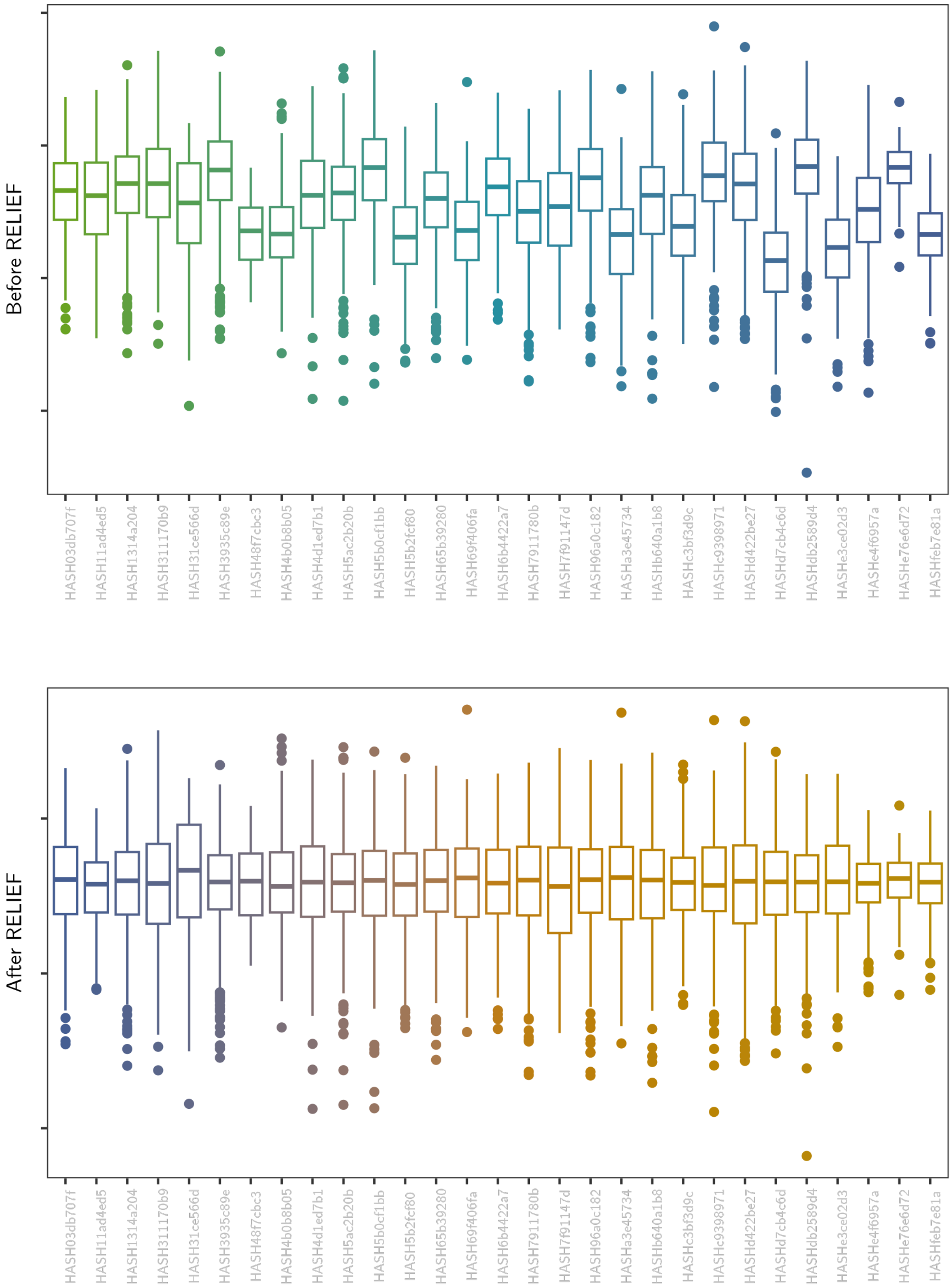

#### Embedding dimension 36

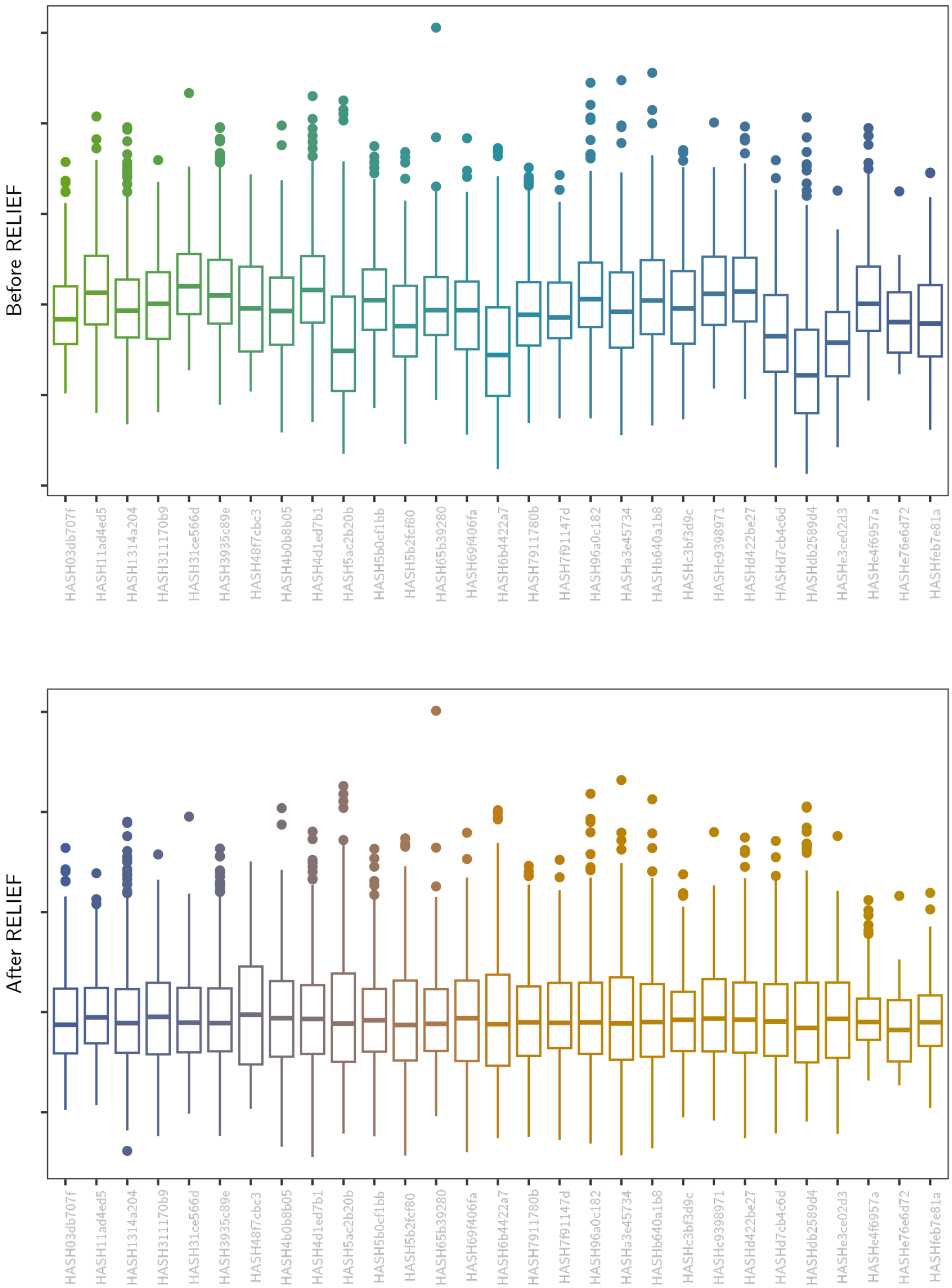

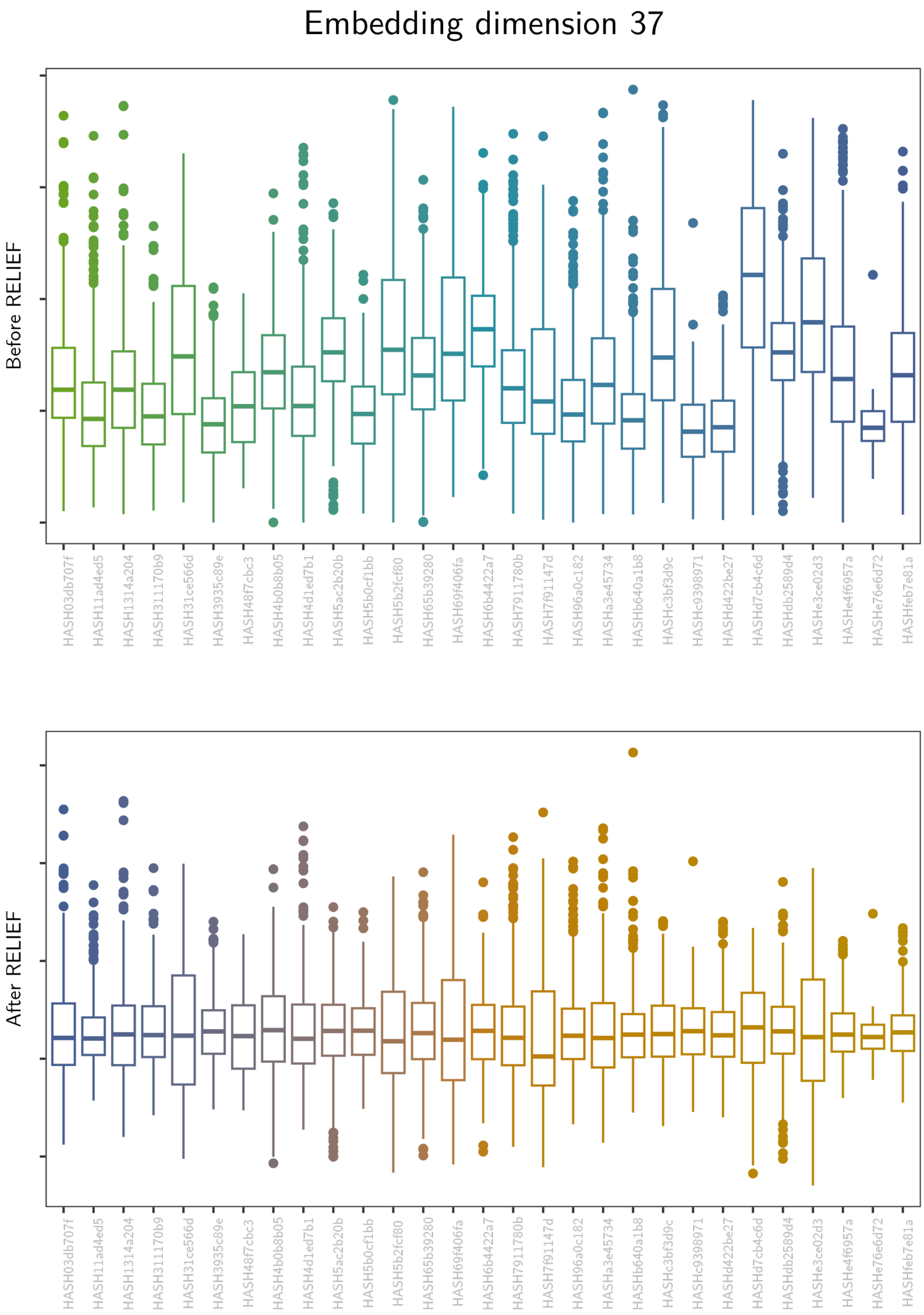
Embedding dimension 37
Before RELIEF
After RELIEF
HASH03db707f
HASH11ad4ed5
HASH1314a204
HASH311170b9
HASH31ce566d
HASH3935c89e
HASH48f7cbc3
HASH4b0b8b05
HASH4d1ed7b1
HASH5ac2b20b
HASH5b0cf1bb
HASH5b2fcf80
HASH65b39280
HASH69f406fa
HASH6b4422a7
HASH7911780b
HASH7f91147d
HASH96a0c182
HASHa3e45734
HASHb640a1b8
HASHc3bf3d9c
HASHc9398971
HASHd422be27
HASHd7cb4c6d
HASHdb2589d4
HASHe3ce02d3
HASHe4f6957a
HASHe76e6d72
HASHfeb7e81a

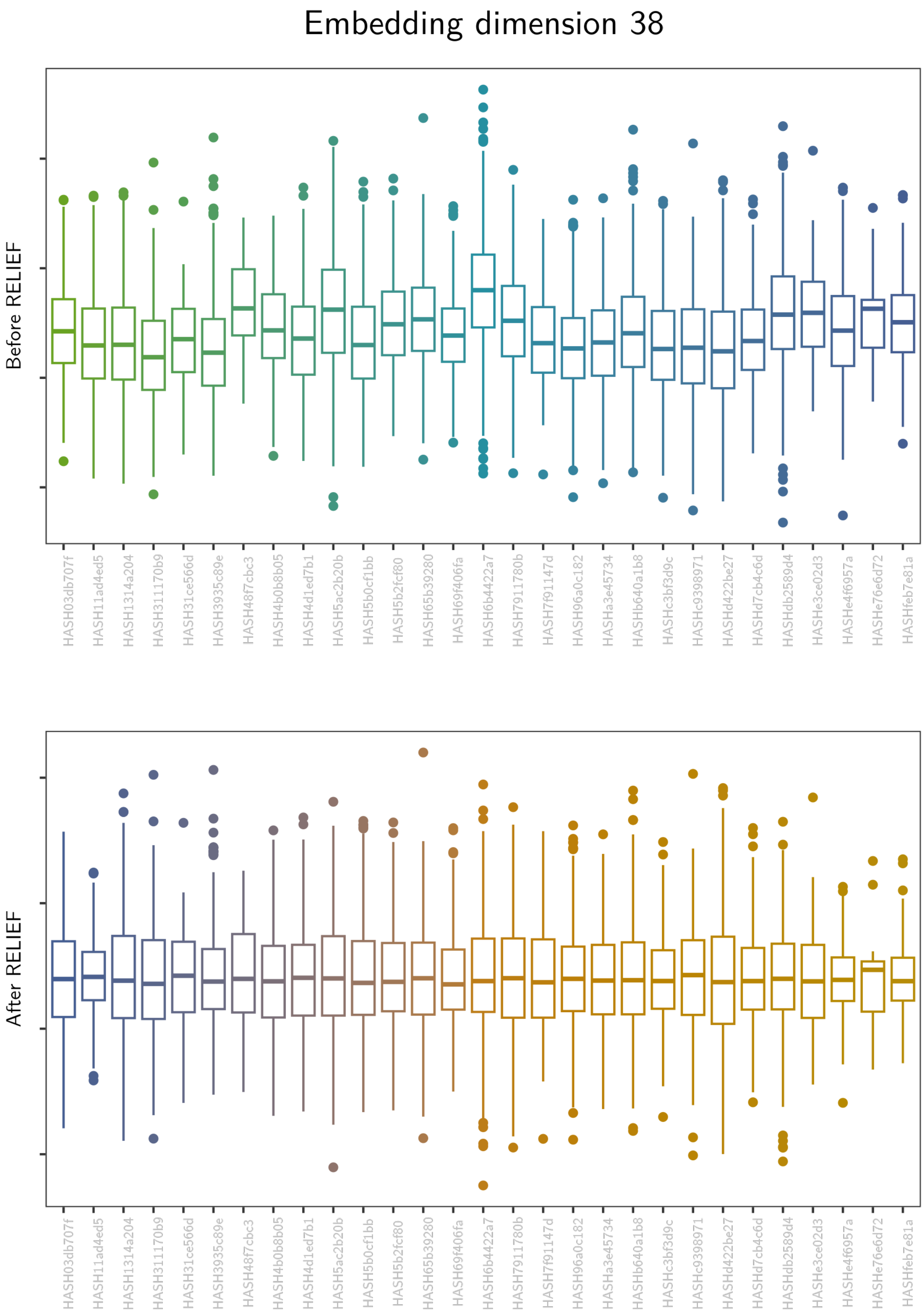
Embedding dimension 38
Before RELIEF
After RELIEF
HASH03db707f
HASH11ad4ed5
HASH1314a204
HASH311170b9
HASH31ce566d
HASH3935c89e
HASH48f7cbc3
HASH4b0b8b05
HASH4d1ed7b1
HASH5ac2b20b
HASH5b0cf1bb
HASH5b2fcf80
HASH65b39280
HASH69f406fa
HASH6b4422a7
HASH7911780b
HASH7f91147d
HASH96a0c182
HASHa3e45734
HASHb640a1b8
HASHc3bf3d9c
HASHc9398971
HASHd422be27
HASHd7cb4c6d
HASHdb2589d4
HASHe3ce02d3
HASHe4f6957a
HASHe76e6d72
HASHfeb7e81a

Embedding dimension 39

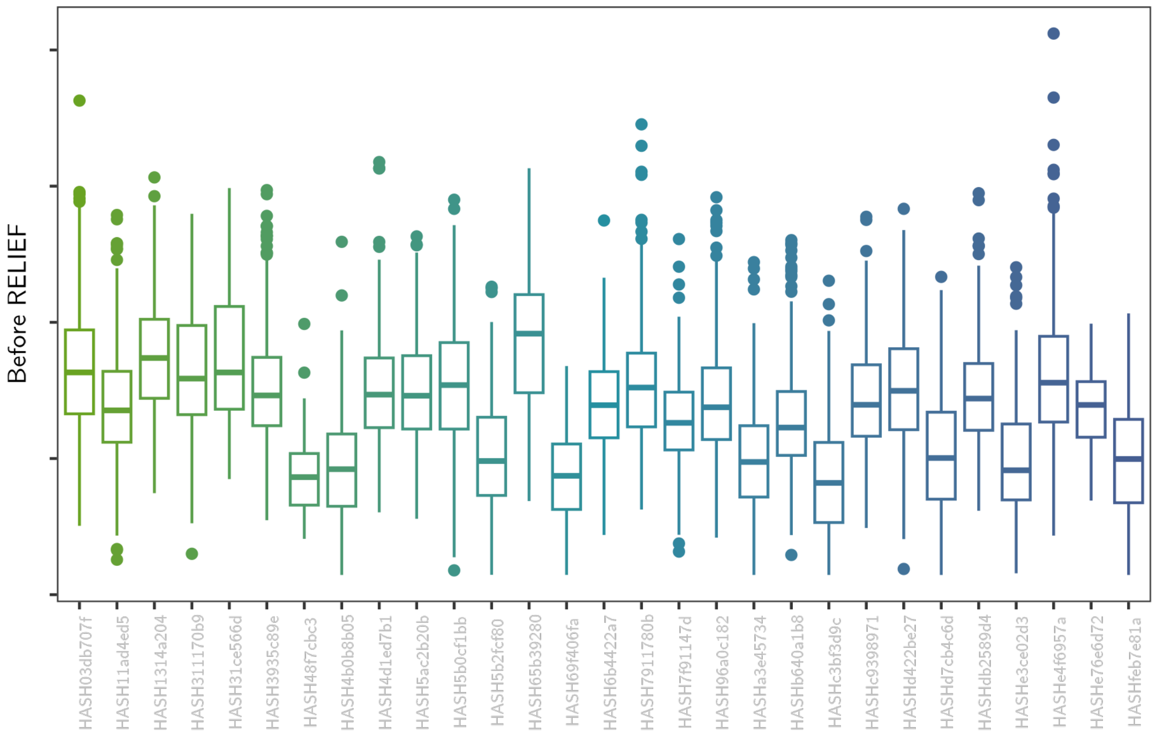

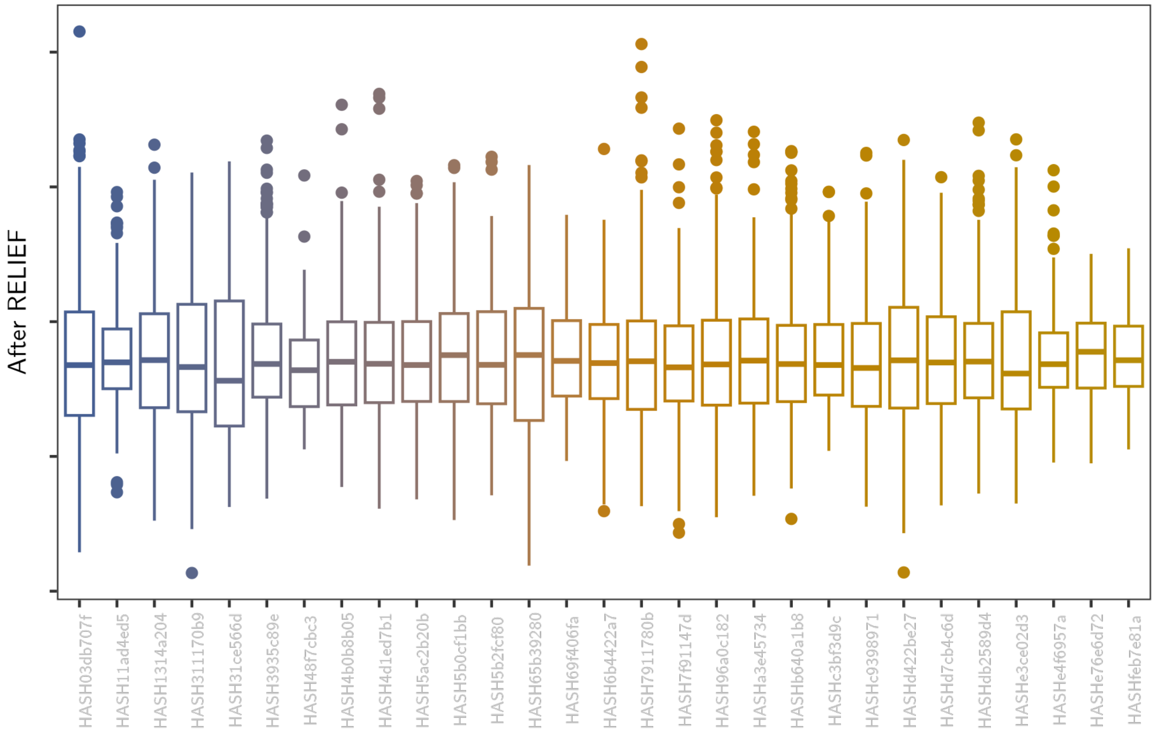

### Embedding dimension 40

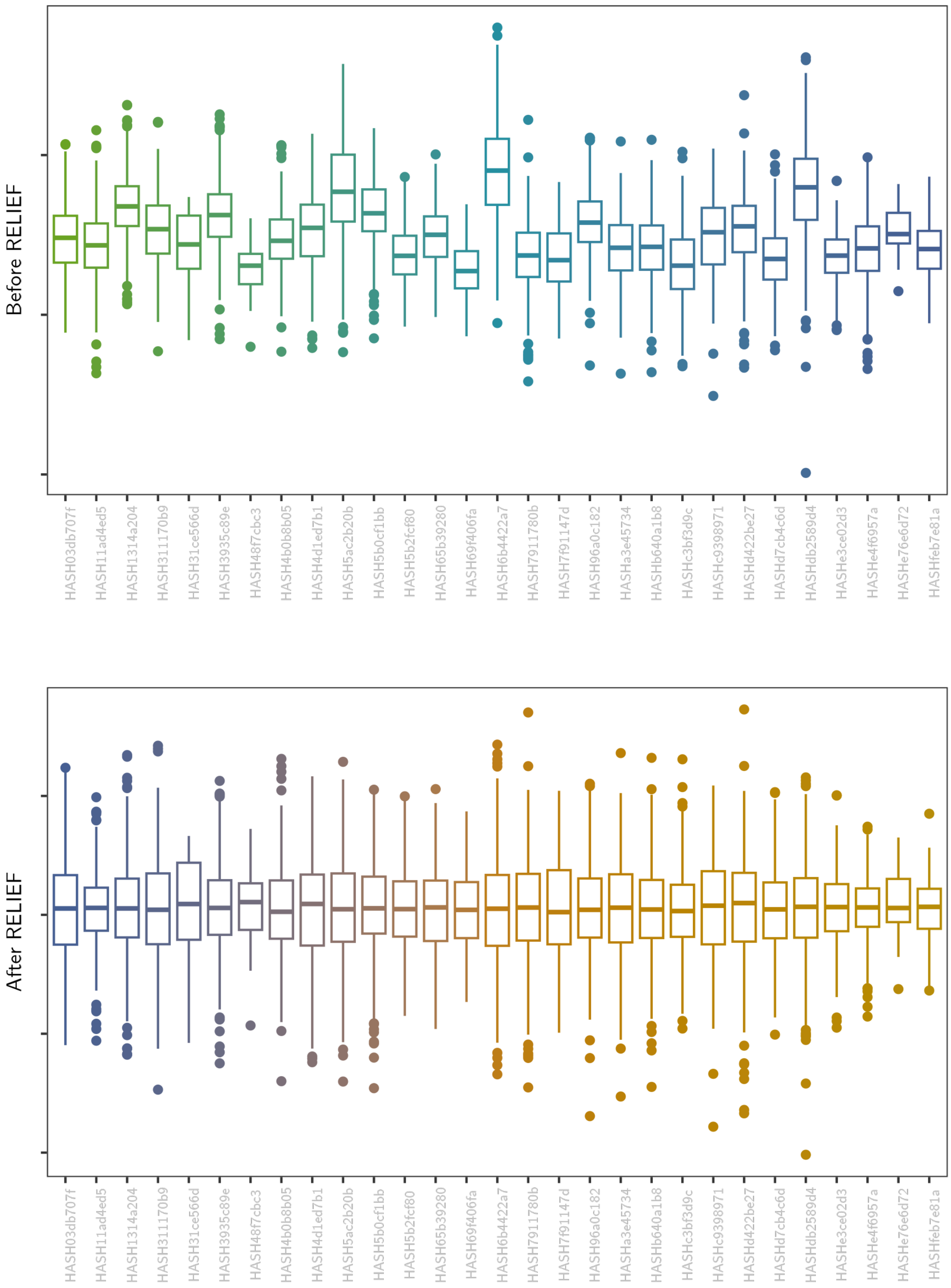

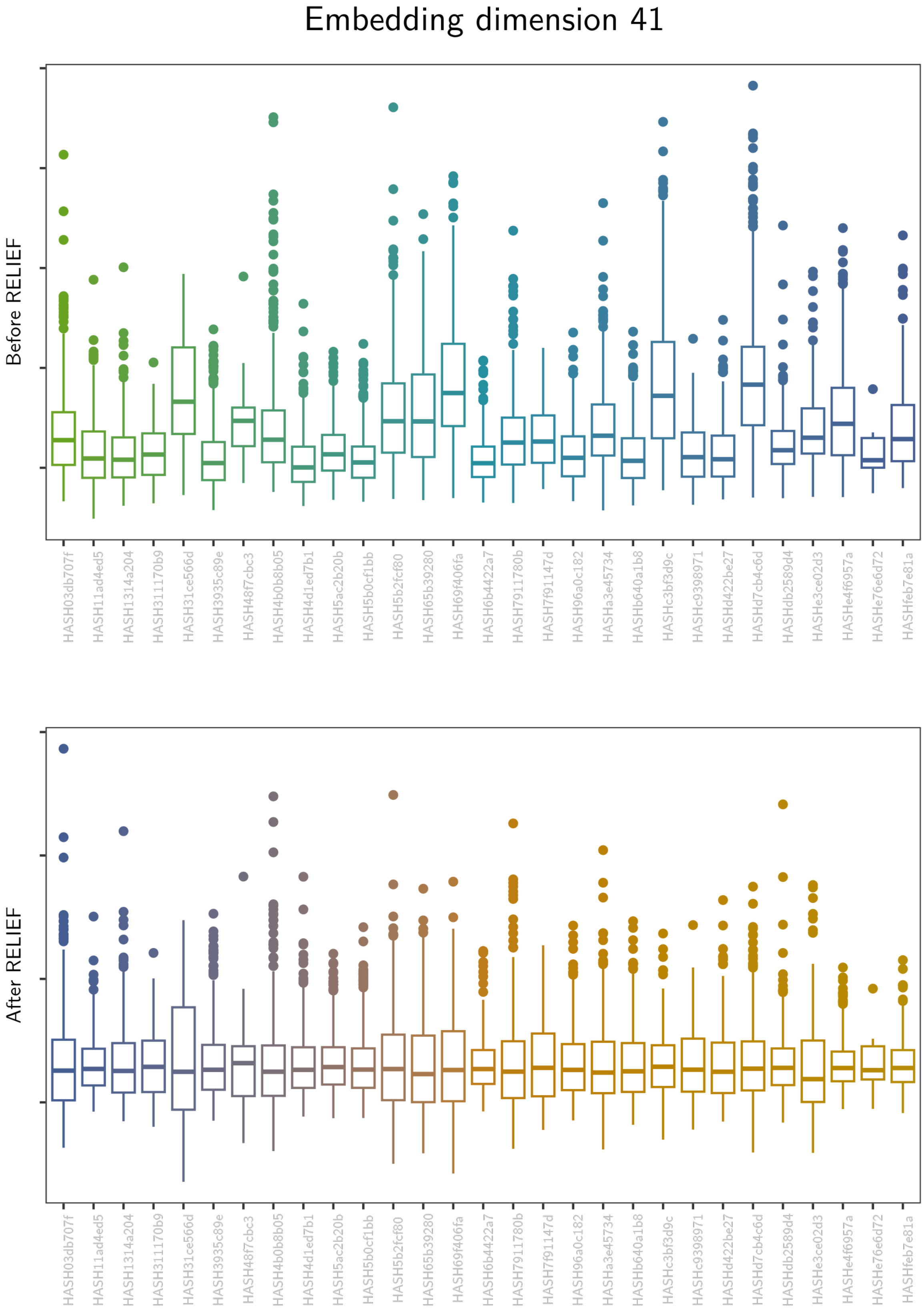

Embedding dimension 41
Before RELIEF
After RELIEF
HASH03db707f
HASH11ad4ed5
HASH1314a204
HASH311170b9
HASH31ce566d
HASH3935c89e
HASH48f7cbc3
HASH4b0b8b05
HASH4d1ed7b1
HASH5ac2b20b
HASH5b0cf1bb
HASH5b2fcf80
HASH65b39280
HASH69f406fa
HASH6b4422a7
HASH7911780b
HASH7f91147d
HASH96a0c182
HASHa3e45734
HASHb640a1b8
HASHc3bf3d9c
HASHc9398971
HASHd422be27
HASHd7cb4c6d
HASHdb2589d4
HASHe3ce02d3
HASHe4f6957a
HASHe76e6d72
HASHfeb7e81a

#### Embedding dimension 42

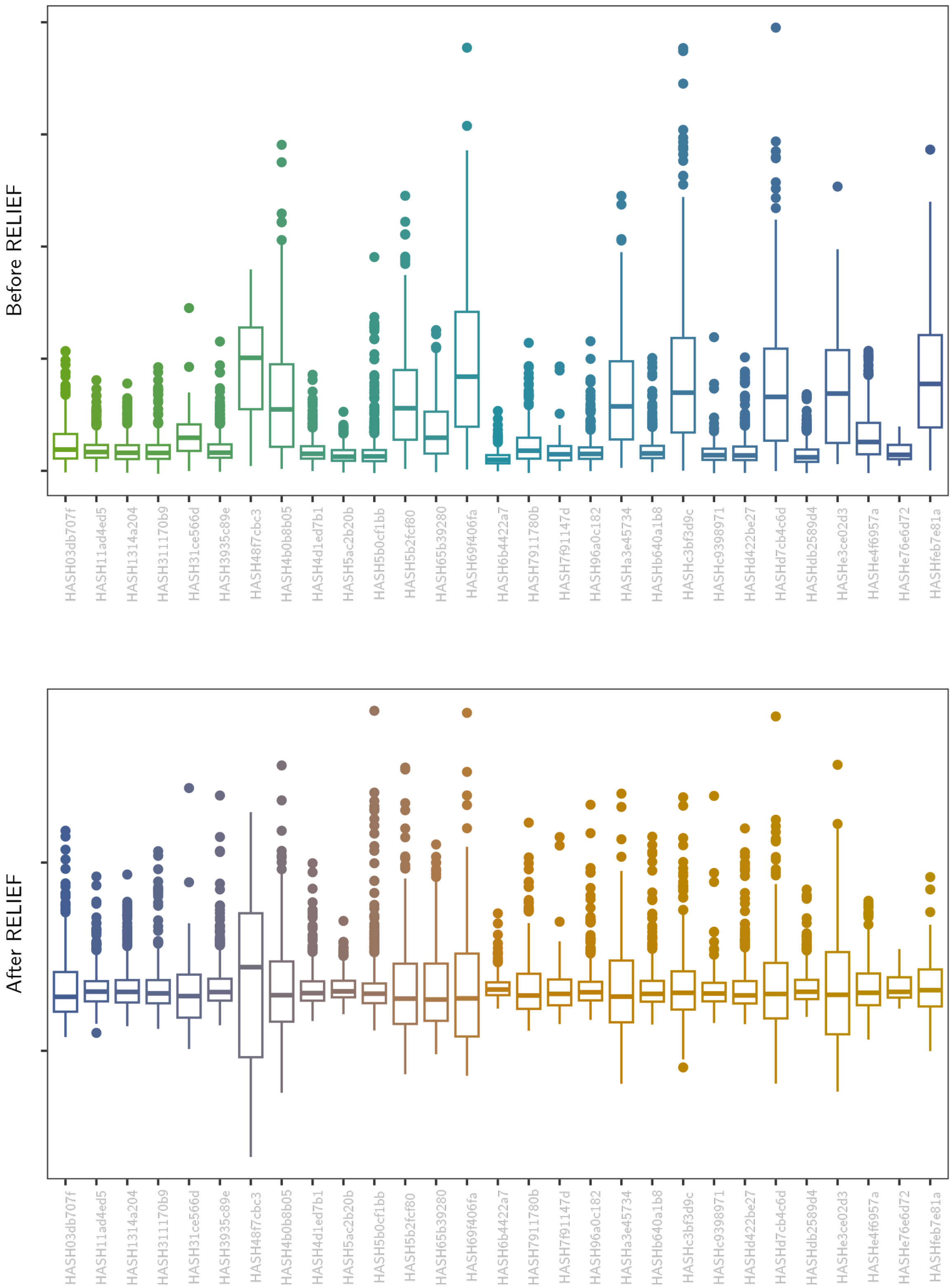

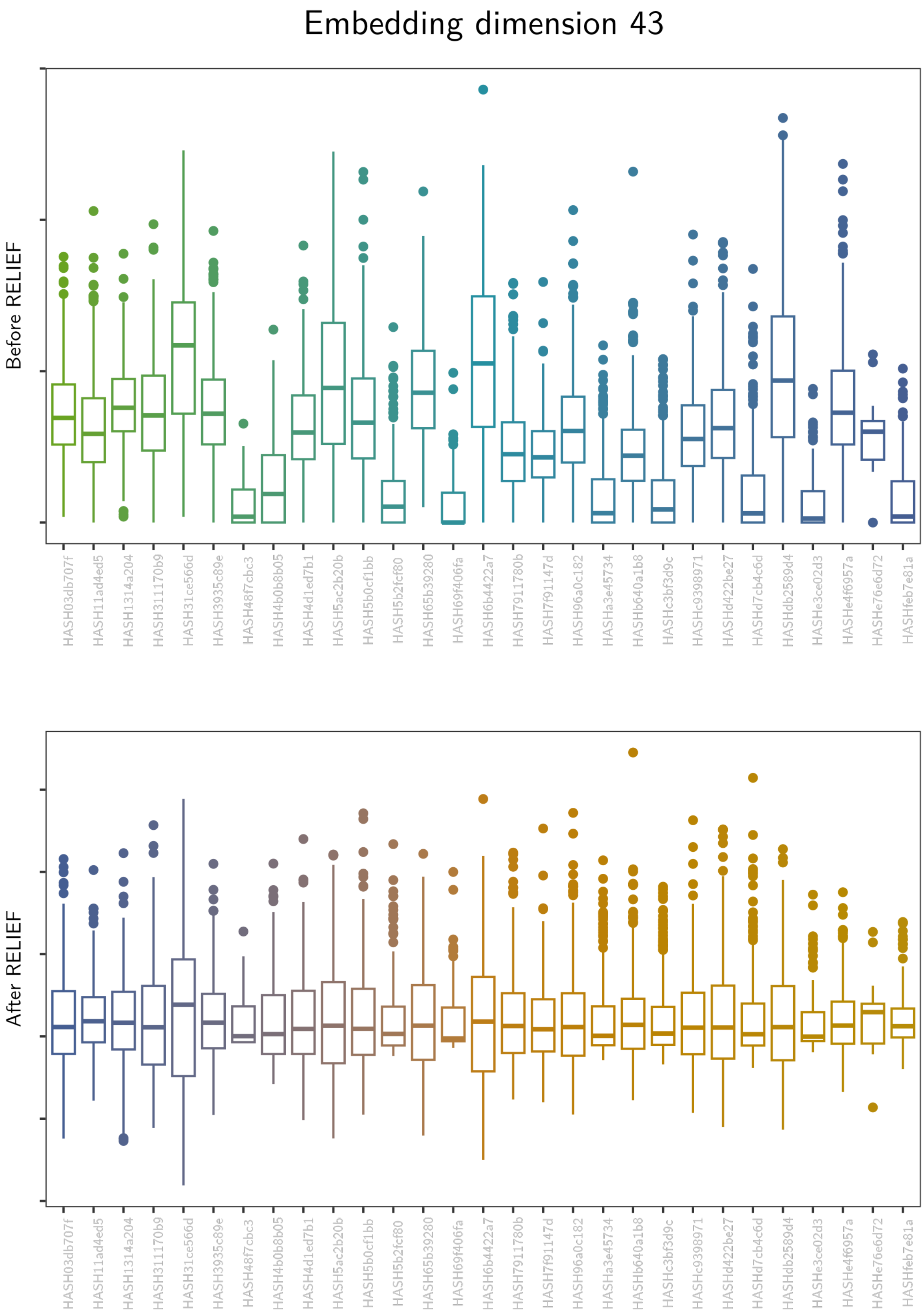
Embedding dimension 43
Before RELIEF
After RELIEF
HASH03db707f
HASH11ad4ed5
HASH1314a204
HASH311170b9
HASH31ce566d
HASH3935c89e
HASH48f7cbc3
HASH4b0b8b05
HASH4d1ed7b1
HASH5ac2b20b
HASH5b0cf1bb
HASH5b2fcf80
HASH65b39280
HASH69f406fa
HASH6b4422a7
HASH7911780b
HASH7f91147d
HASH96a0c182
HASHa3e45734
HASHb640a1b8
HASHc3bf3d9c
HASHc9398971
HASHd422be27
HASHd7cb4c6d
HASHdb2589d4
HASHe3ce02d3
HASHe4f6957a
HASHe76e6d72
HASHfeb7e81a

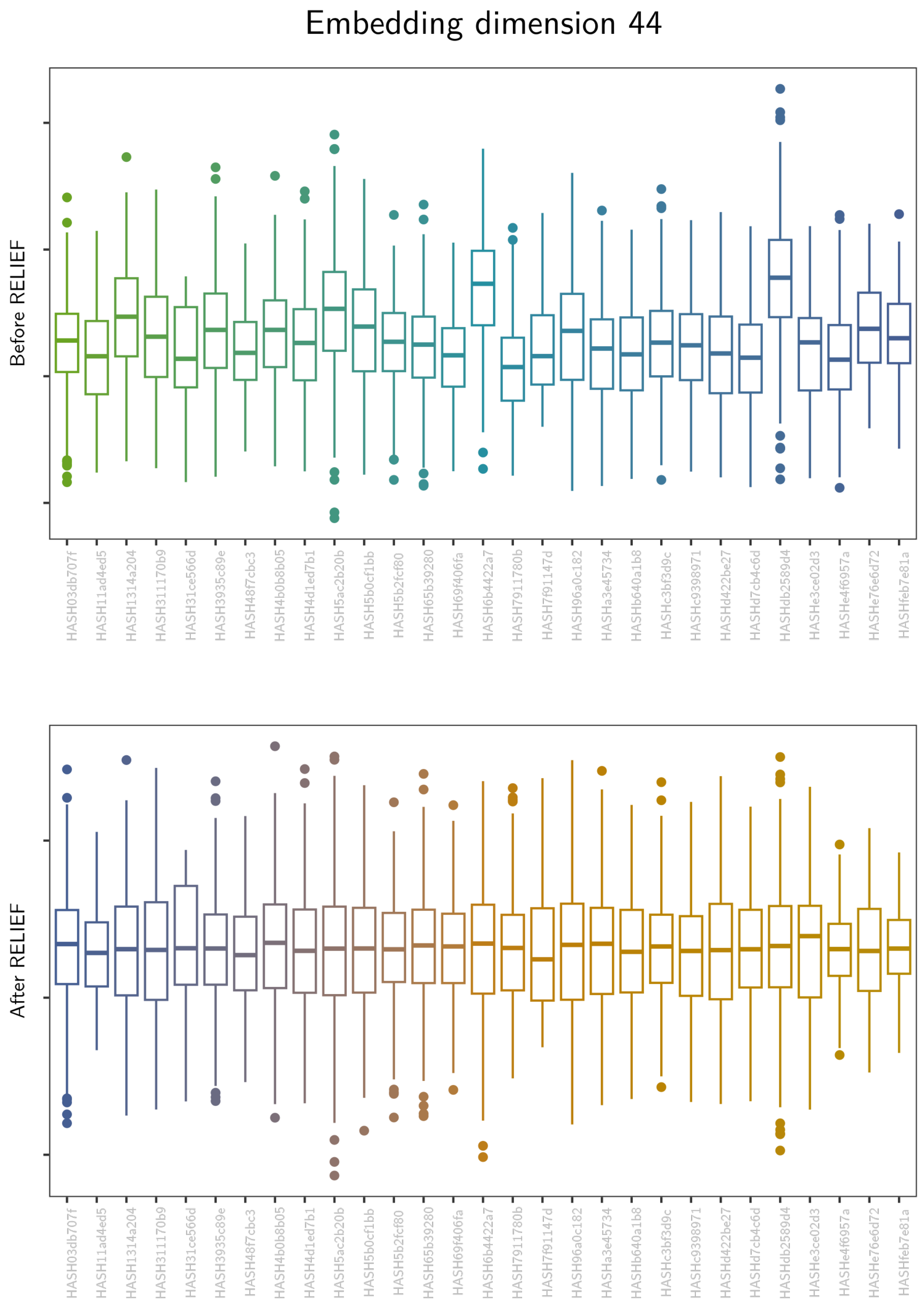
Embedding dimension 44
Before RELIEF
After RELIEF
HASH03db707f
HASH11ad4ed5
HASH1314a204
HASH311170b9
HASH31ce566d
HASH3935c89e
HASH48f7cbc3
HASH4b0b8b05
HASH4d1ed7b1
HASH5ac2b20b
HASH5b0cf1bb
HASH5b2fcf80
HASH65b39280
HASH69f406fa
HASH6b4422a7
HASH7911780b
HASH7f91147d
HASH96a0c182
HASHa3e45734
HASHb640a1b8
HASHc3bf3d9c
HASHc9398971
HASHd422be27
HASHd7cb4c6d
HASHdb2589d4
HASHe3ce02d3
HASHe4f6957a
HASHe76e6d72
HASHfeb7e81a

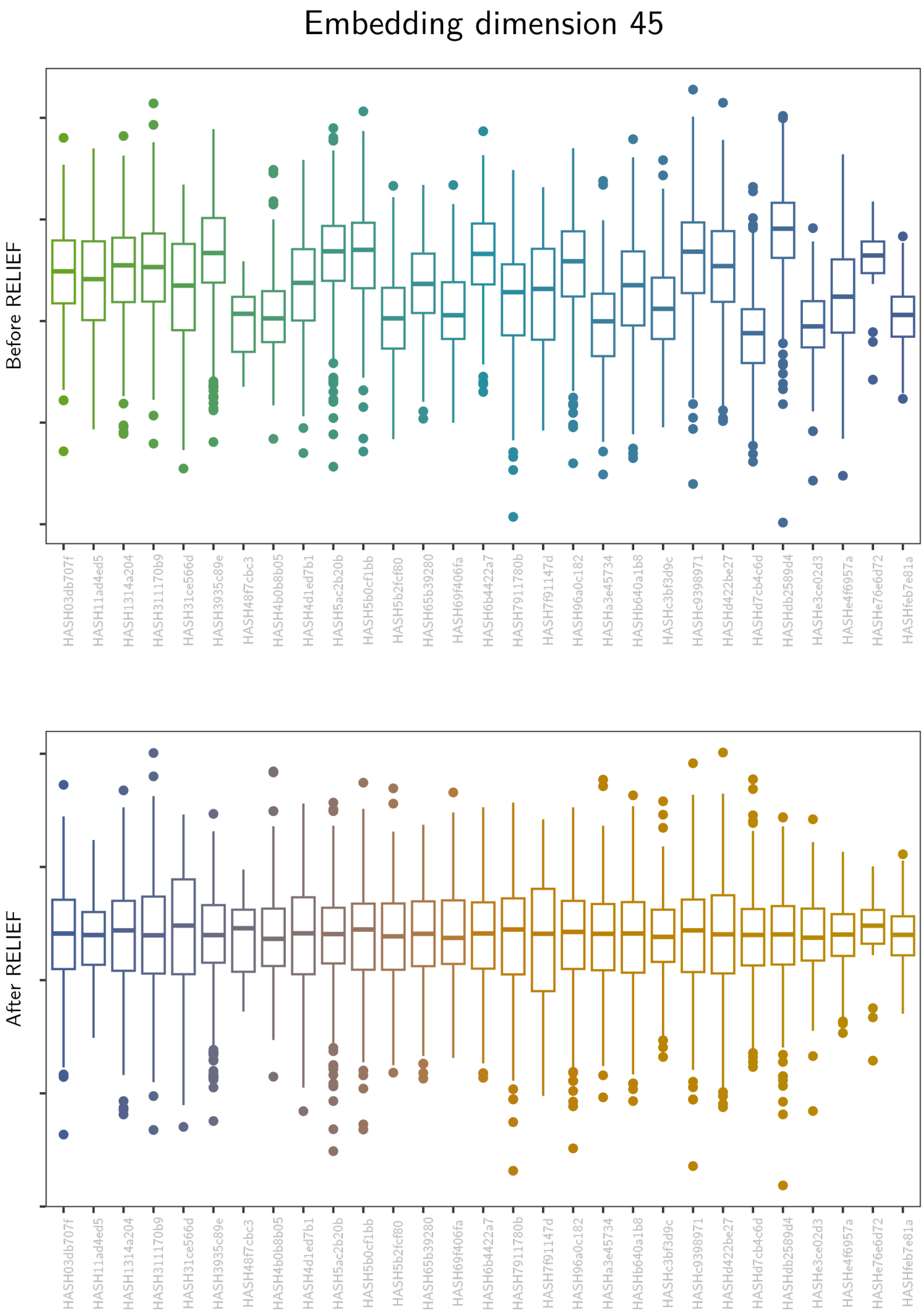
Embedding dimension 45
Before RELIEF
After RELIEF
HASH03db707f
HASH11ad4ed5
HASH1314a204
HASH311170b9
HASH31ce566d
HASH3935c89e
HASH48f7cbc3
HASH4b0b8b05
HASH4d1ed7b1
HASH5ac2b20b
HASH5b0cf1bb
HASH5b2fcf80
HASH65b39280
HASH69f406fa
HASH6b4422a7
HASH7911780b
HASH7f91147d
HASH96a0c182
HASHa3e45734
HASHb640a1b8
HASHc3bf3d9c
HASHc9398971
HASHd422be27
HASHd7cb4c6d
HASHdb2589d4
HASHe3ce02d3
HASHe4f6957a
HASHe76e6d72
HASHfeb7e81a

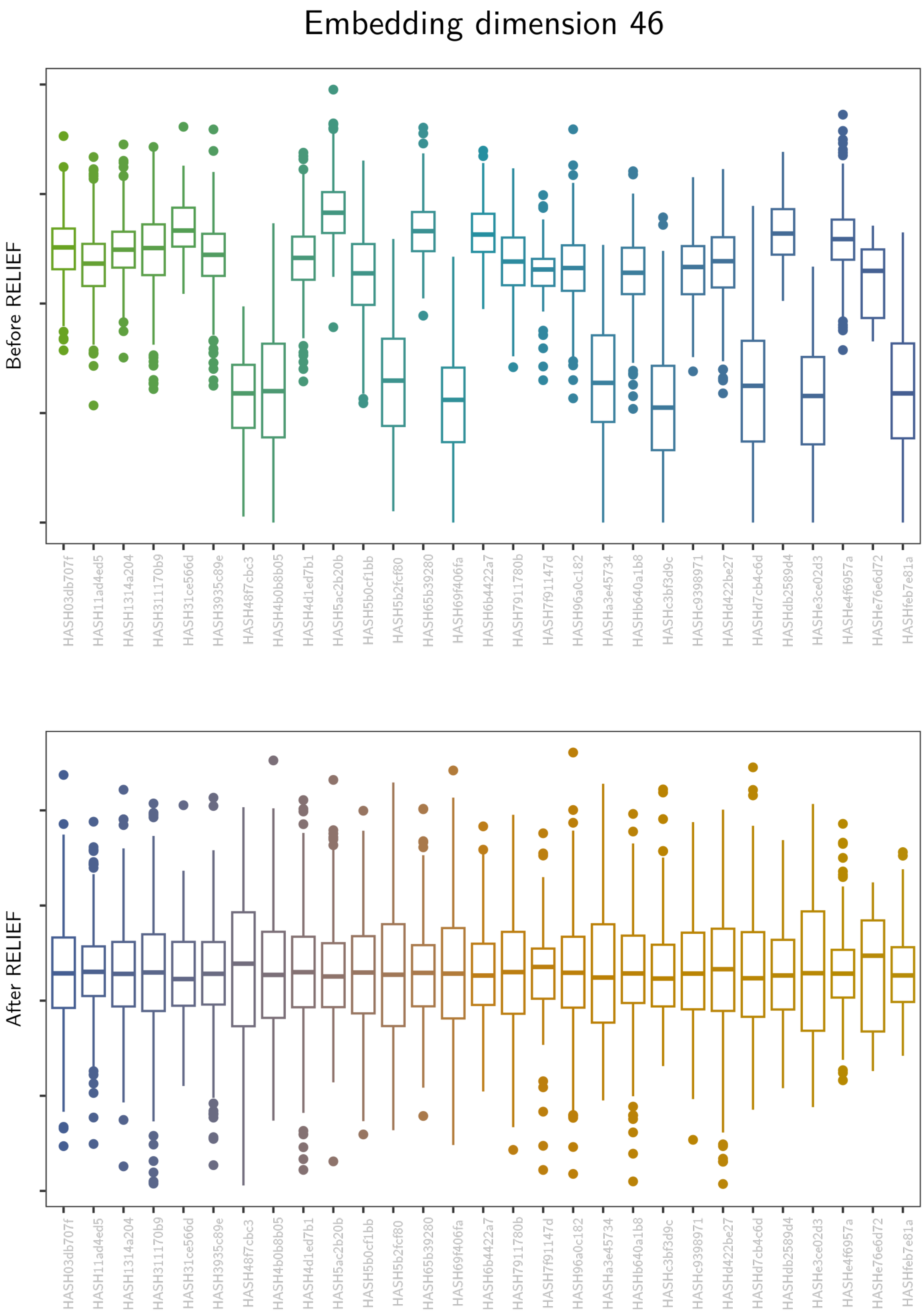
Embedding dimension 46
Before RELIEF
After RELIEF
HASH03db707f
HASH11ad4ed5
HASH1314a204
HASH311170b9
HASH31ce566d
HASH3935c89e
HASH48f7cbc3
HASH4b0b8b05
HASH4d1ed7b1
HASH5ac2b20b
HASH5b0cf1bb
HASH5b2fcf80
HASH65b39280
HASH69f406fa
HASH6b4422a7
HASH7911780b
HASH7f91147d
HASH96a0c182
HASHa3e45734
HASHb640a1b8
HASHc3bf3d9c
HASHc9398971
HASHd422be27
HASHd7cb4c6d
HASHdb2589d4
HASHe3ce02d3
HASHe4f6957a
HASHe76e6d72
HASHfeb7e81a

Embedding dimension 47

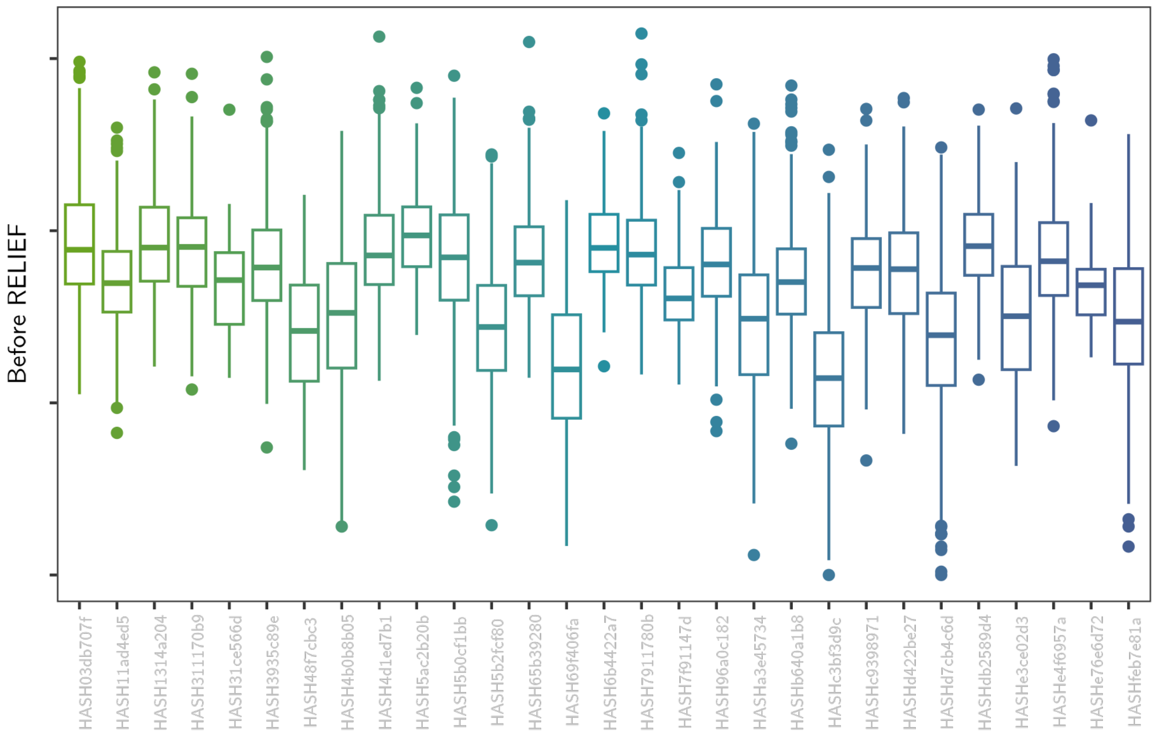

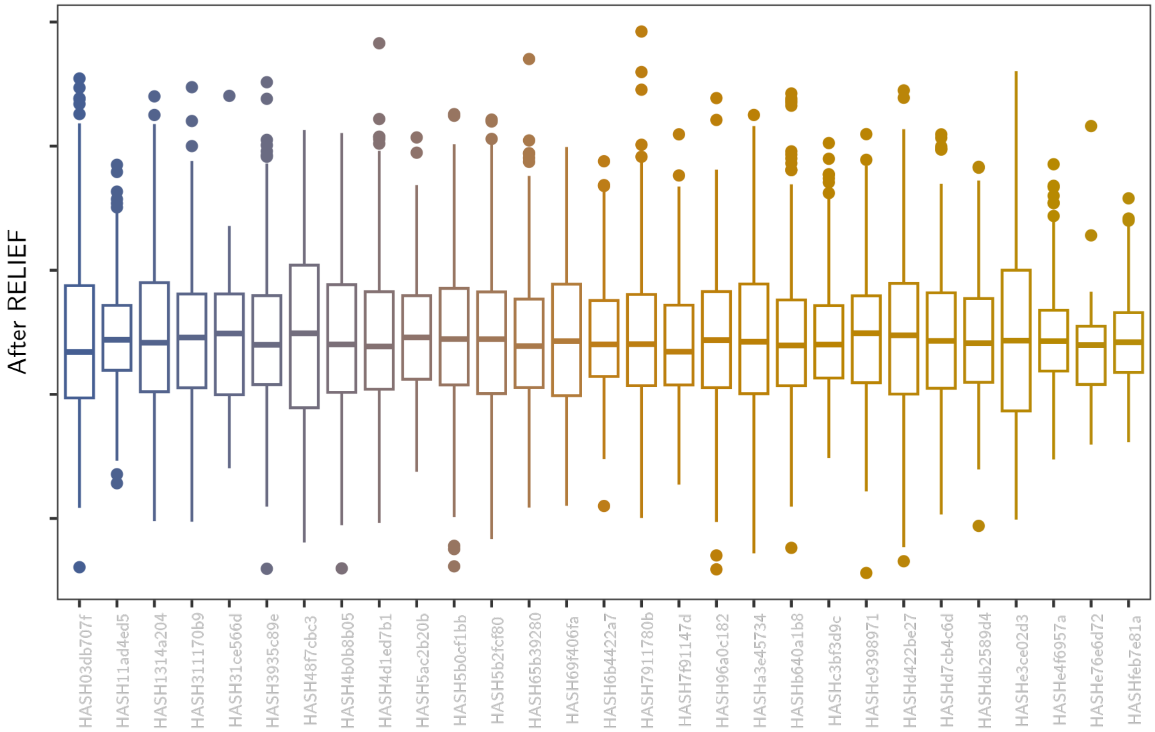

Embedding dimension 48

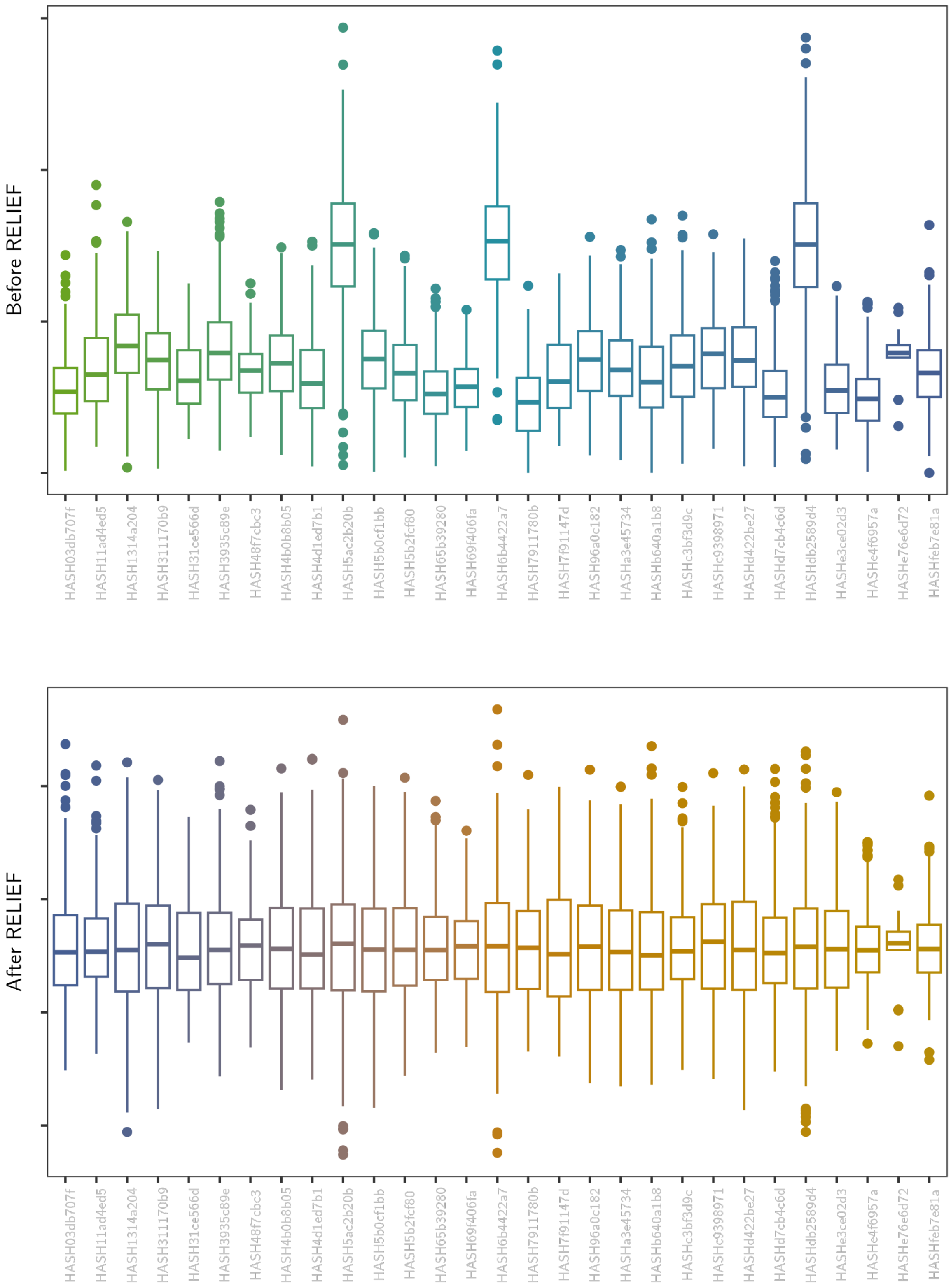

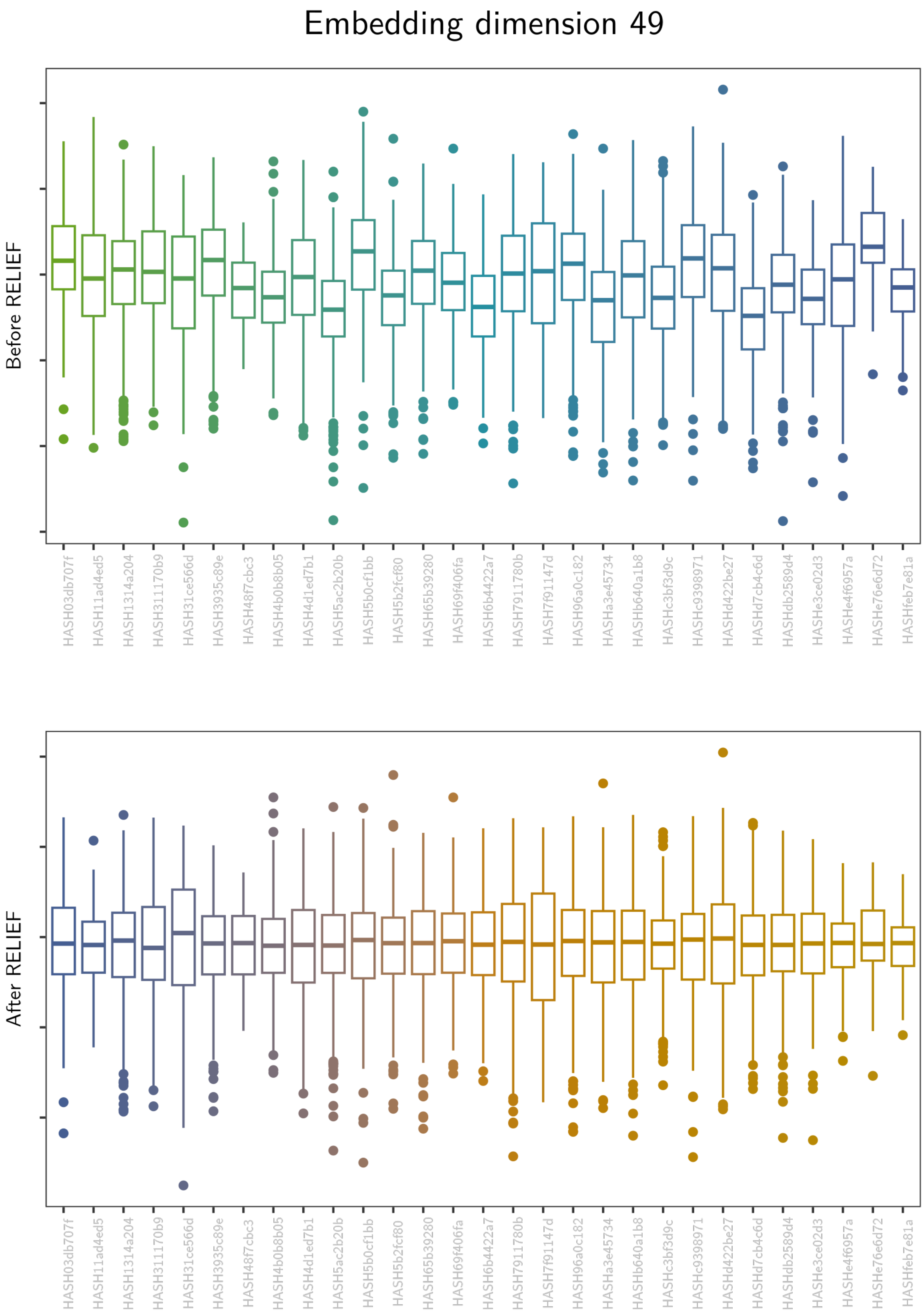

Embedding dimension 49
Before RELIEF
After RELIEF
HASH03db707f
HASH11ad4ed5
HASH1314a204
HASH311170b9
HASH31ce566d
HASH3935c89e
HASH48f7cbc3
HASH4b0b8b05
HASH4d1ed7b1
HASH5ac2b20b
HASH5b0cf1bb
HASH5b2fcf80
HASH65b39280
HASH69f406fa
HASH6b4422a7
HASH7911780b
HASH7f91147d
HASH96a0c182
HASHa3e45734
HASHb640a1b8
HASHc3bf3d9c
HASHc9398971
HASHd422be27
HASHd7cb4c6d
HASHdb2589d4
HASHe3ce02d3
HASHe4f6957a
HASHe76e6d72
HASHfeb7e81a

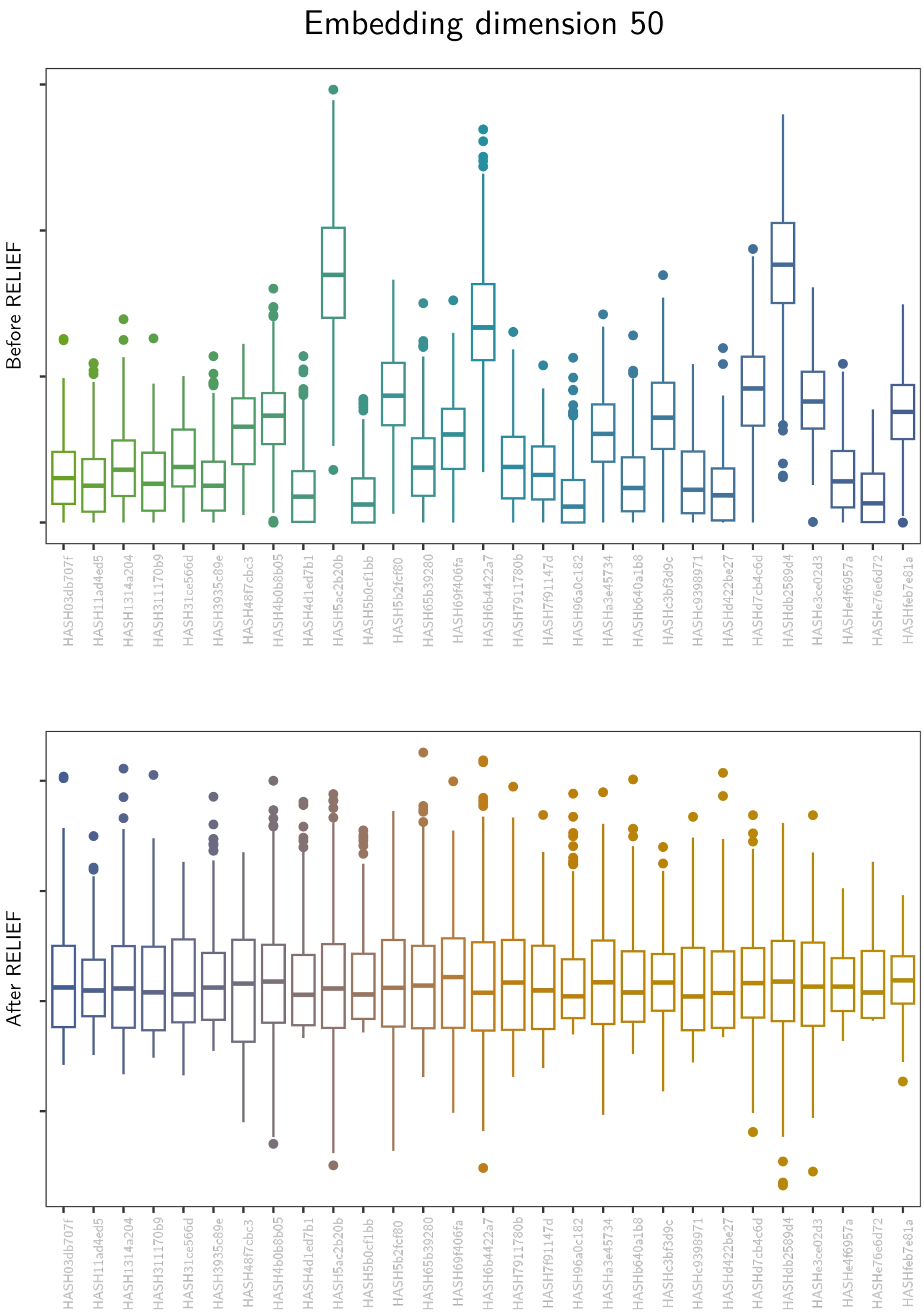
Embedding dimension 50
Before RELIEF
After RELIEF
HASH03db707f
HASH11ad4ed5
HASH1314a204
HASH311170b9
HASH31ce566d
HASH3935c89e
HASH48f7cbc3
HASH4b0b8b05
HASH4d1ed7b1
HASH5ac2b20b
HASH5b0cf1bb
HASH5b2fcf80
HASH65b39280
HASH69f406fa
HASH6b4422a7
HASH7911780b
HASH7f91147d
HASH96a0c182
HASHa3e45734
HASHb640a1b8
HASHc3bf3d9c
HASHc9398971
HASHd422be27
HASHd7cb4c6d
HASHdb2589d4
HASHe3ce02d3
HASHe4f6957a
HASHe76e6d72
HASHfeb7e81a

#### Embedding dimension 51

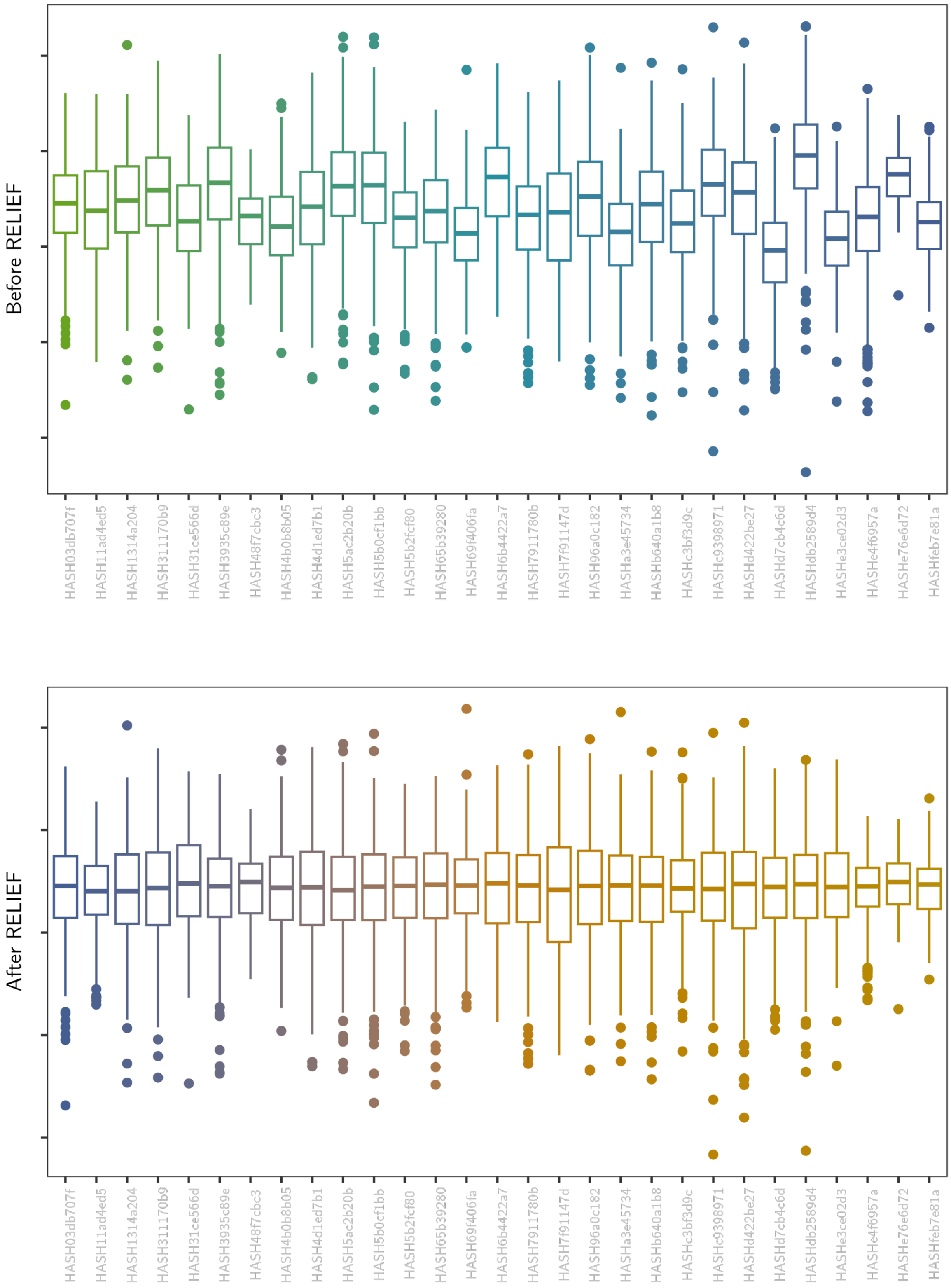

#### Embedding dimension 52

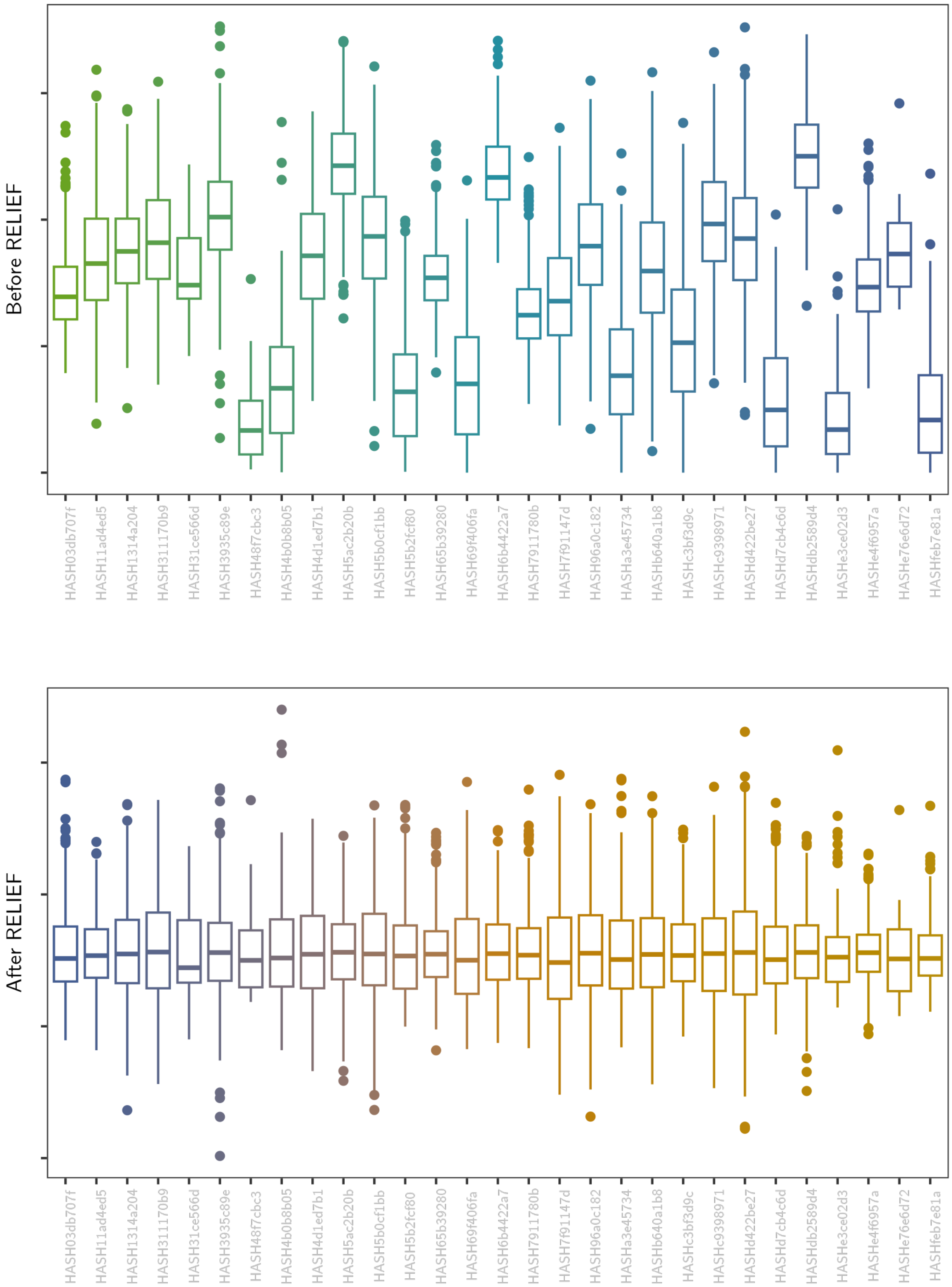

Embedding dimension 53

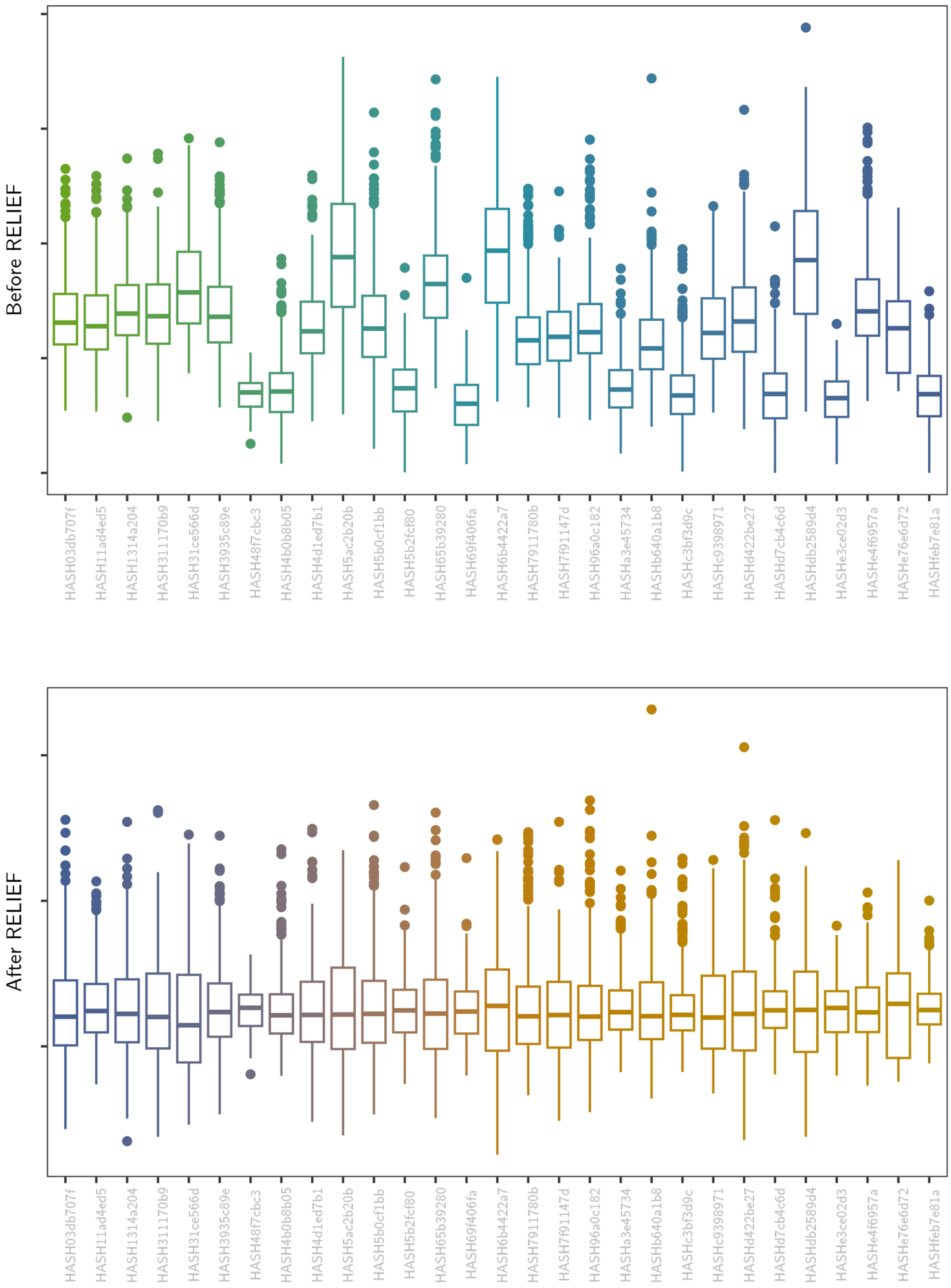

### Embedding dimension 54

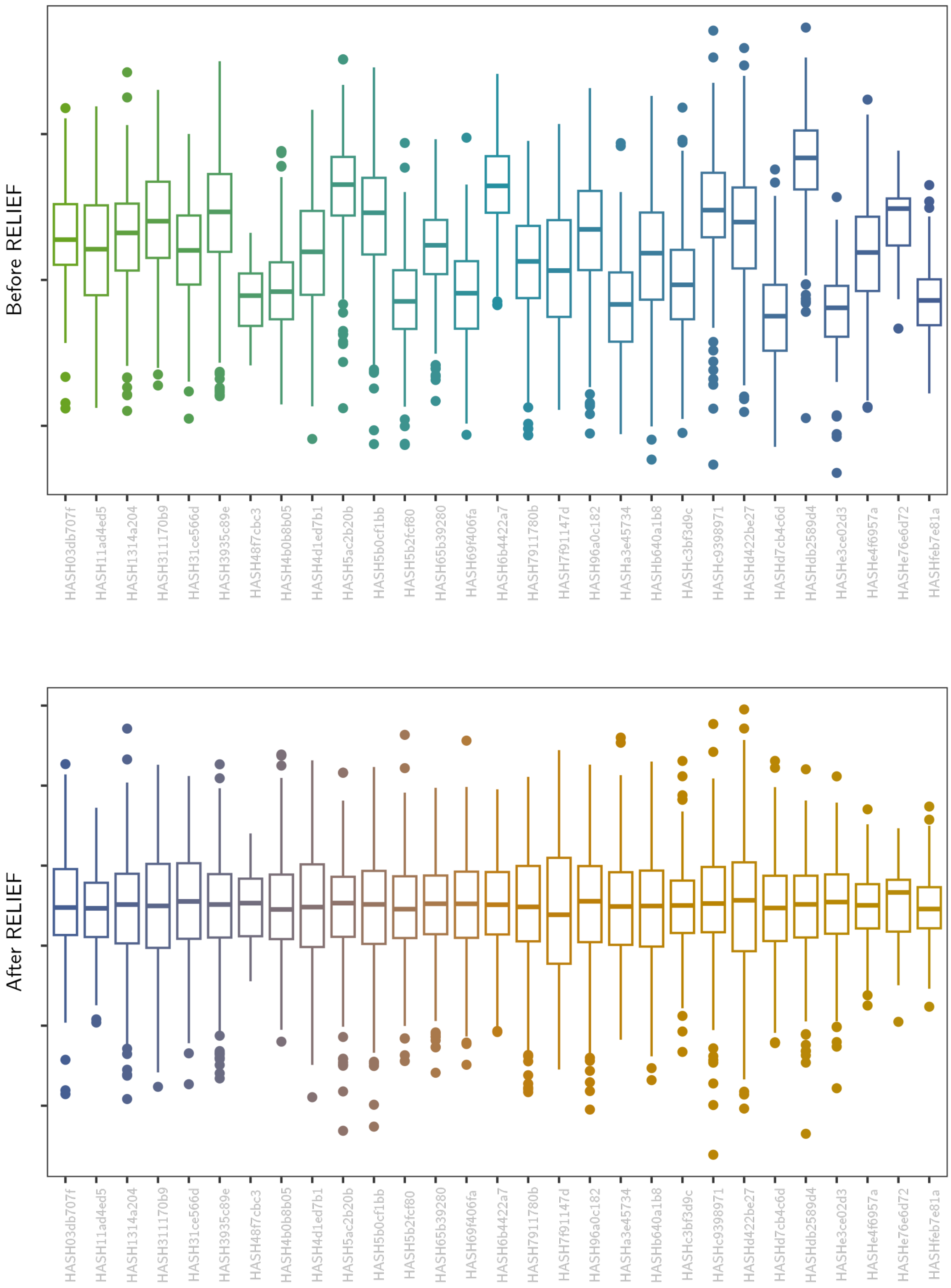

Embedding dimension 55

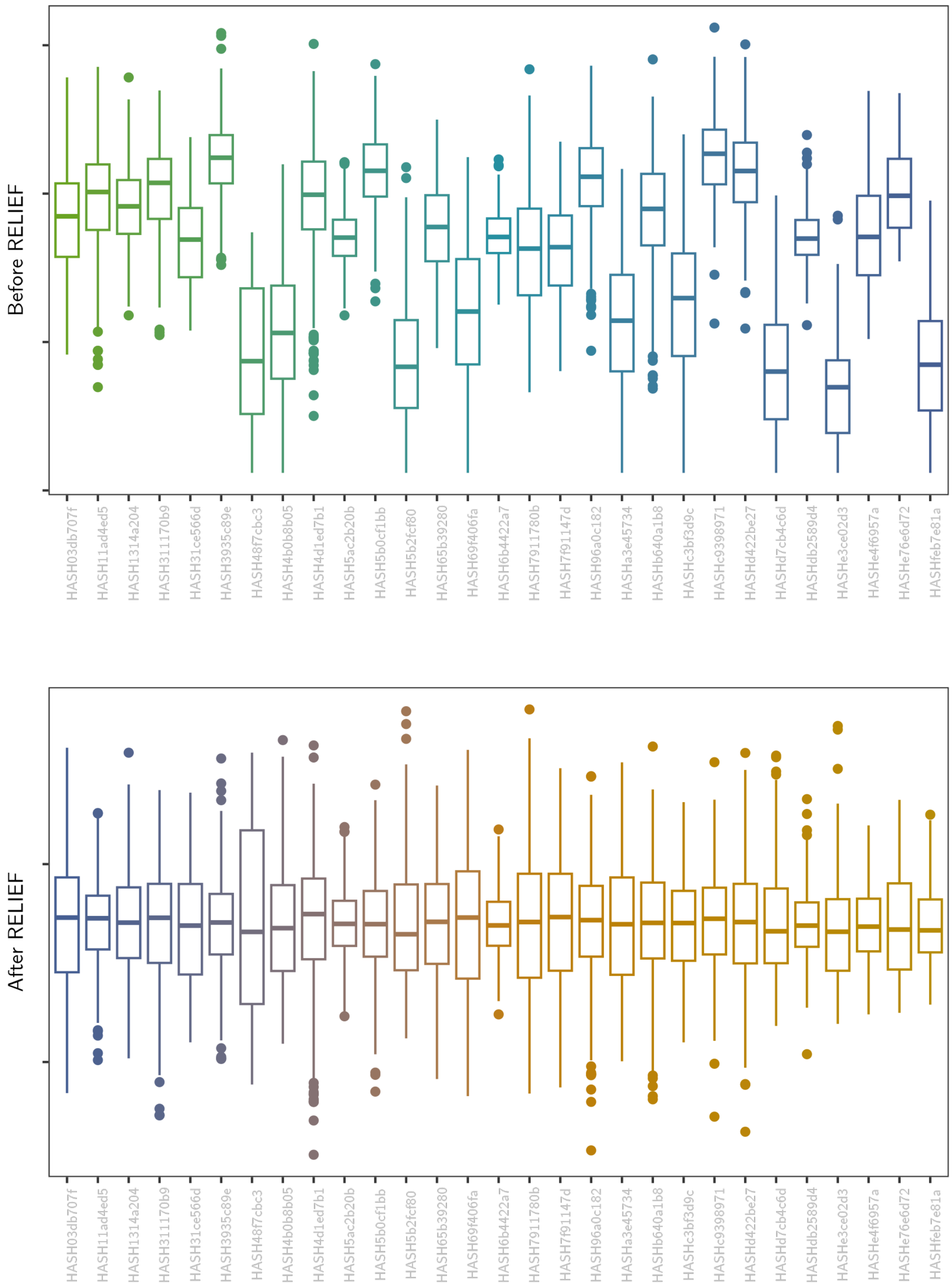

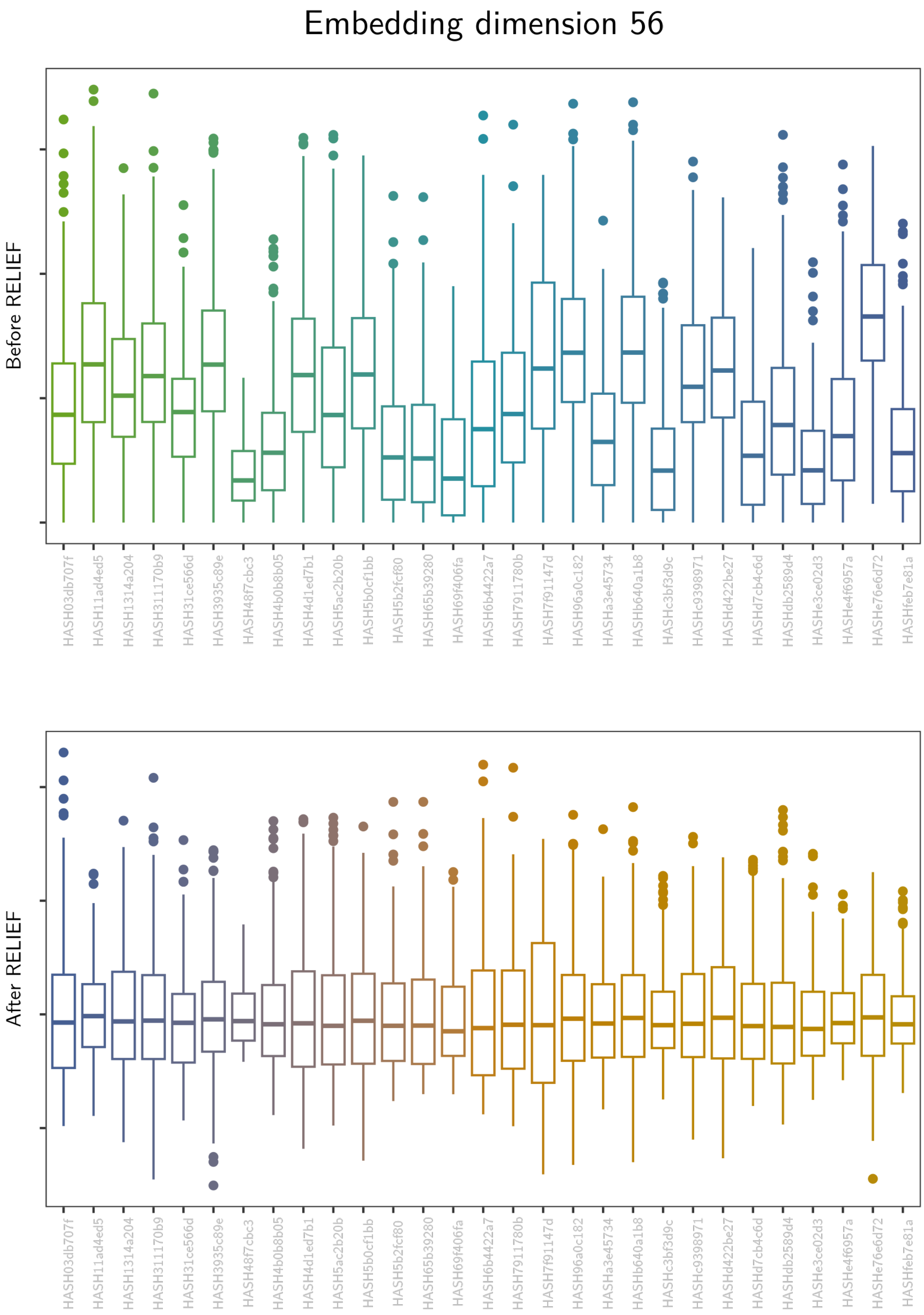

Embedding dimension 56
Before RELIEF
After RELIEF
HASH03db707f
HASH11ad4ed5
HASH1314a204
HASH311170b9
HASH31ce566d
HASH3935c89e
HASH48f7cbc3
HASH4b0b8b05
HASH4d1ed7b1
HASH5ac2b20b
HASH5b0cf1bb
HASH5b2fcf80
HASH65b39280
HASH69f406fa
HASH6b4422a7
HASH7911780b
HASH7f91147d
HASH96a0c182
HASHa3e45734
HASHb640a1b8
HASHc3bf3d9c
HASHc9398971
HASHd422be27
HASHd7cb4c6d
HASHdb2589d4
HASHe3ce02d3
HASHe4f6957a
HASHe76e6d72
HASHfeb7e81a

### Embedding dimension 57

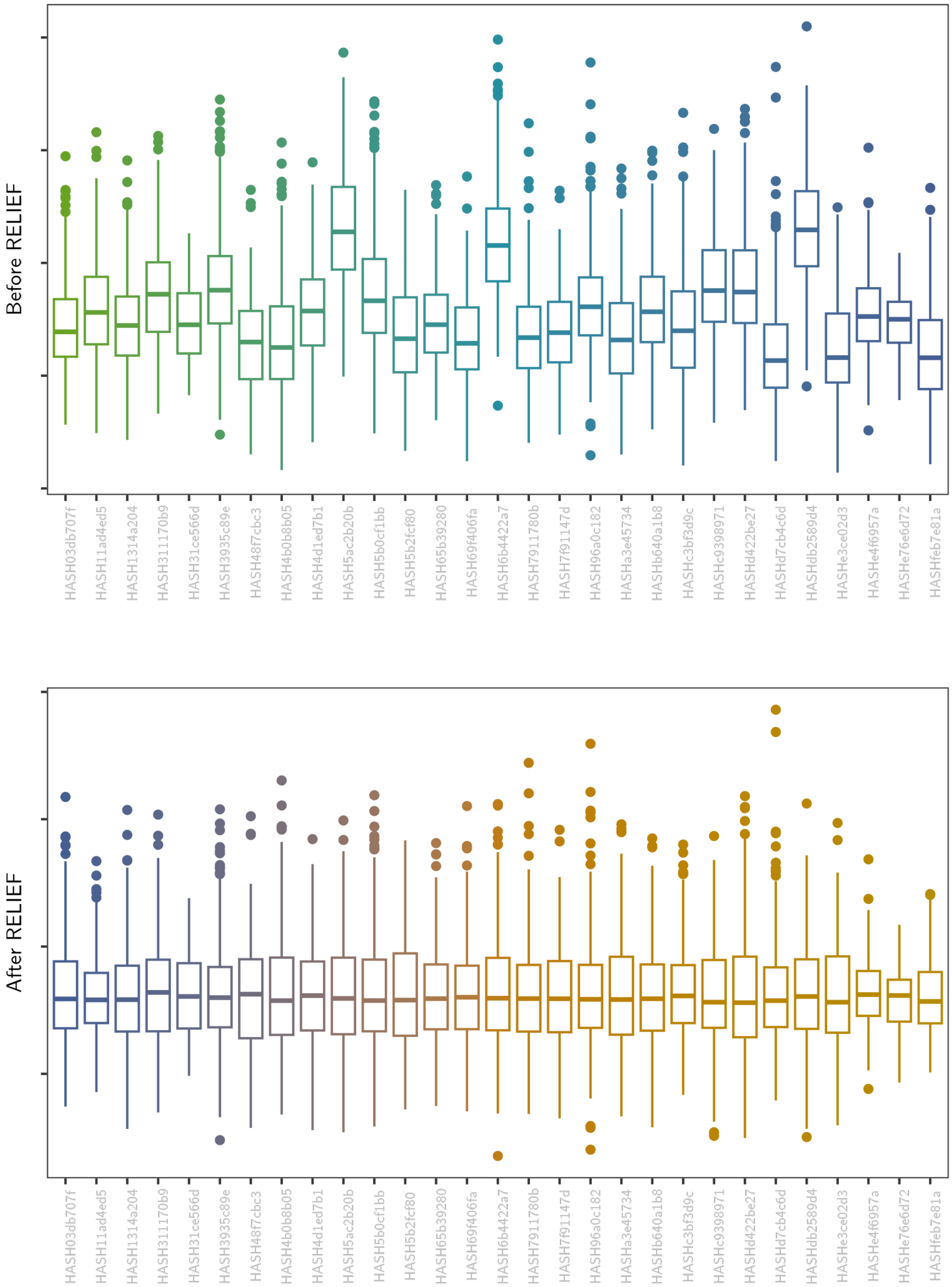

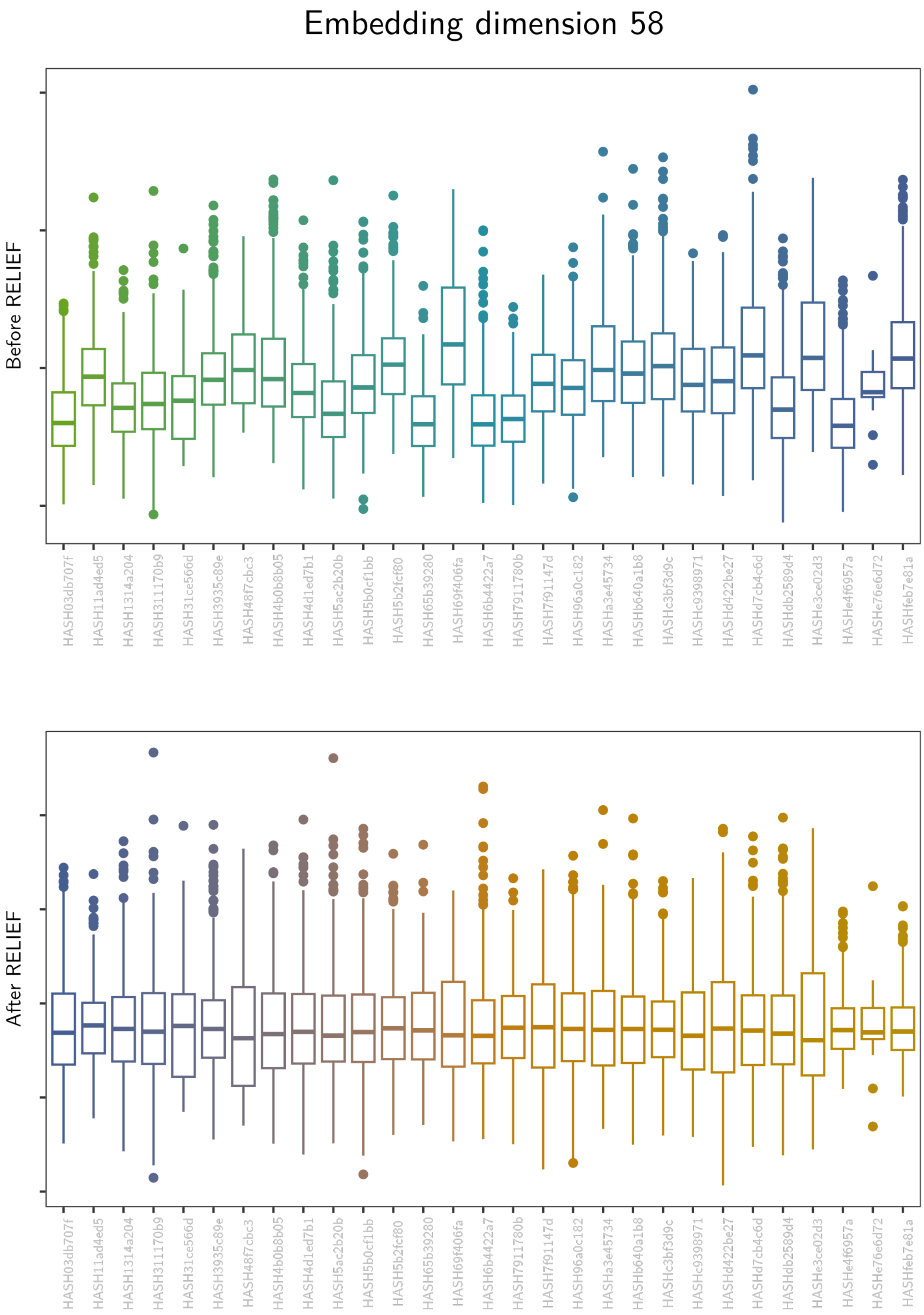
Embedding dimension 58
Before RELIEF
After RELIEF
HASH03db707f
HASH11ad4ed5
HASH1314a204
HASH311170b9
HASH31ce566d
HASH3935c89e
HASH48f7cbc3
HASH4b0b8b05
HASH4d1ed7b1
HASH5ac2b20b
HASH5b0cf1bb
HASH5b2fcf80
HASH65b39280
HASH69f406fa
HASH6b4422a7
HASH7911780b
HASH7f91147d
HASH96a0c182
HASHa3e45734
HASHb640a1b8
HASHc3bf3d9c
HASHc9398971
HASHd422be27
HASHd7cb4c6d
HASHdb2589d4
HASHe3ce02d3
HASHe4f6957a
HASHe76e6d72
HASHfeb7e81a

### Embedding dimension 59

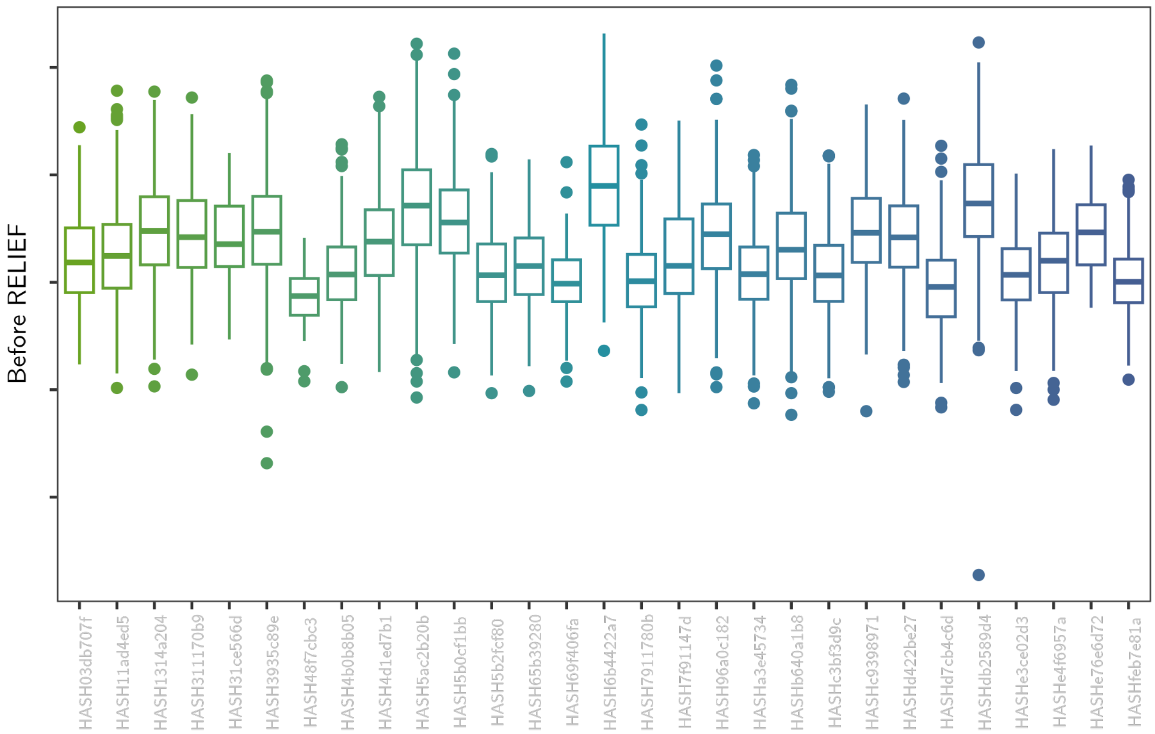

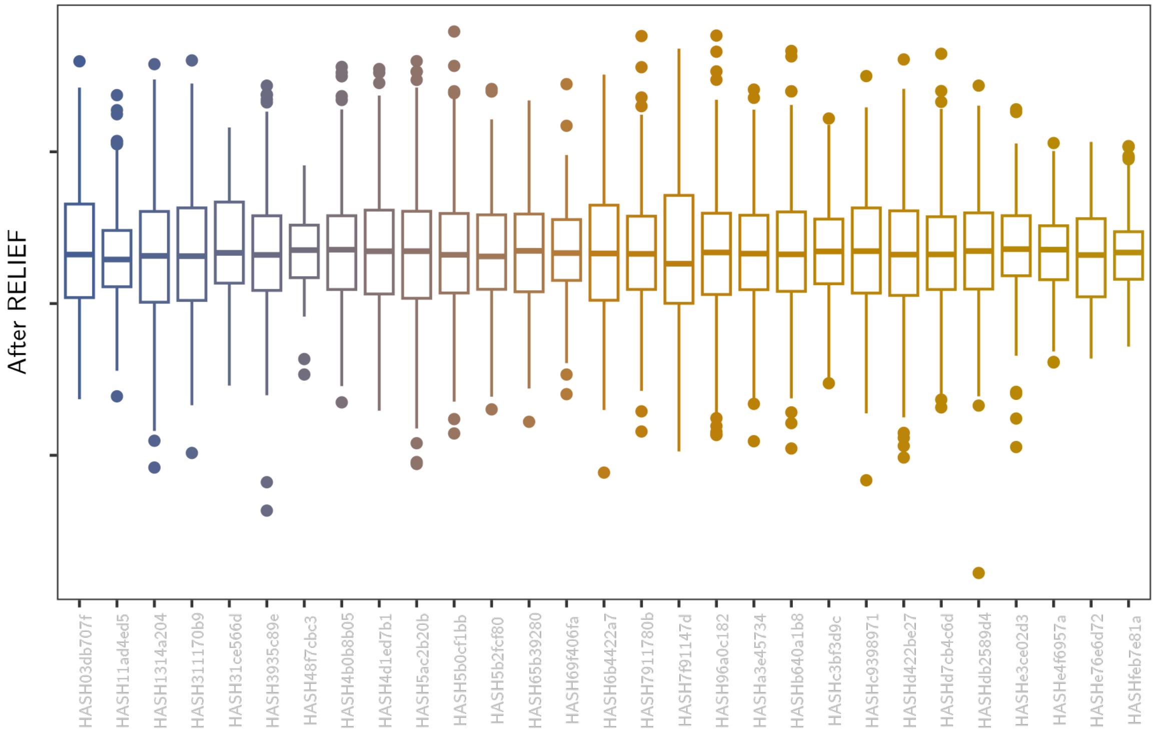

Embedding dimension 60

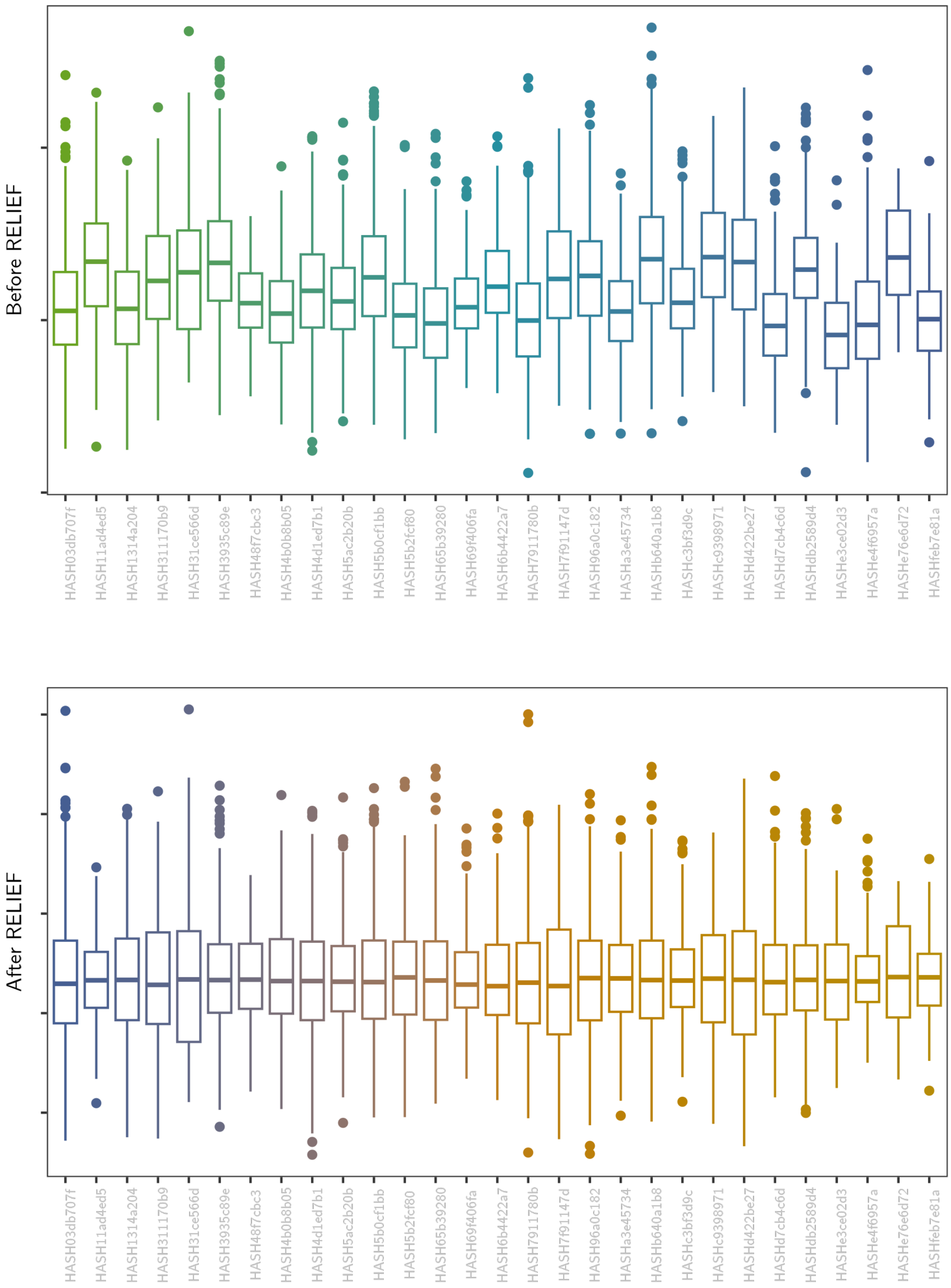

Supplementary Figure 2.1 to 2.60: Embedding dimension average explanations

Average relevance for each embedding dimension, consisting of the mean of all non-linearly FNIRT-transformed relevance maps for a given dimension for 1000 random test fold subjects. Each relevance map was divided by the highest absolute relevance of that map, before being superimposed on the MNI152 template, using the slice where the highest absolute relevance was found.

Dimension 0, Baseline

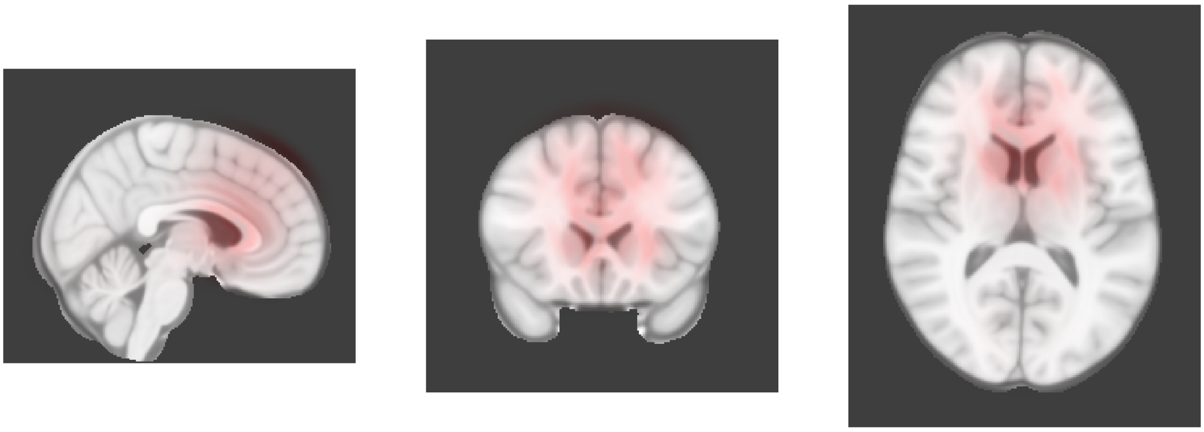

Dimension 0, 2-year follow-up

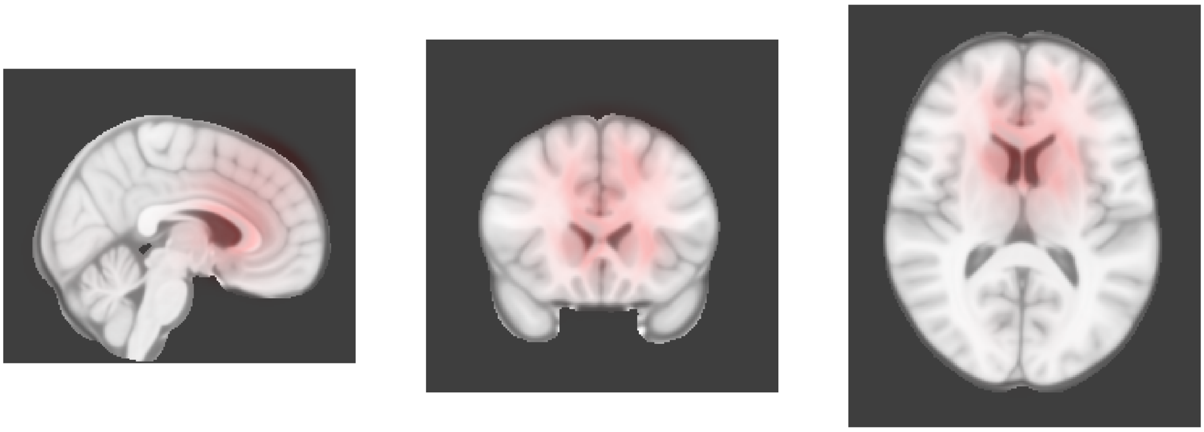

Dimension 1, Baseline

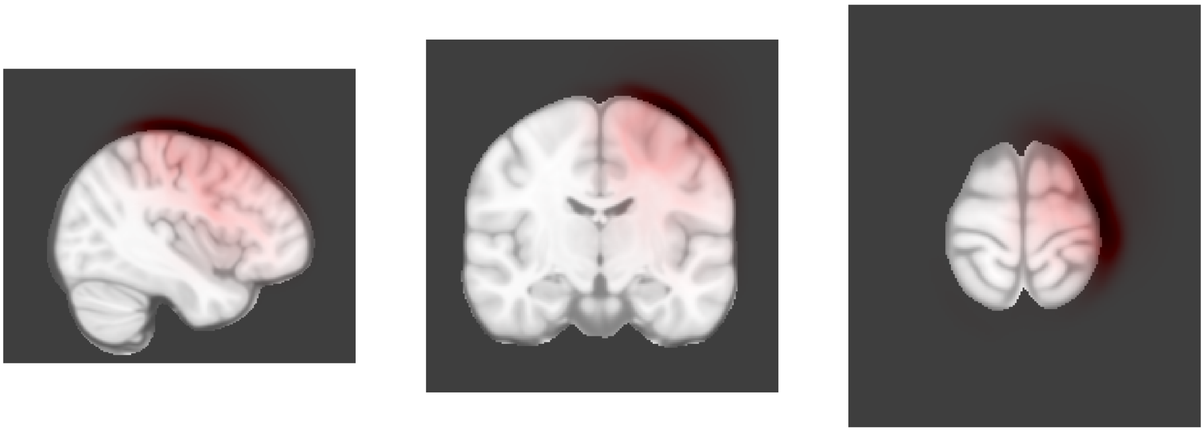

Dimension 1, 2-year follow-up

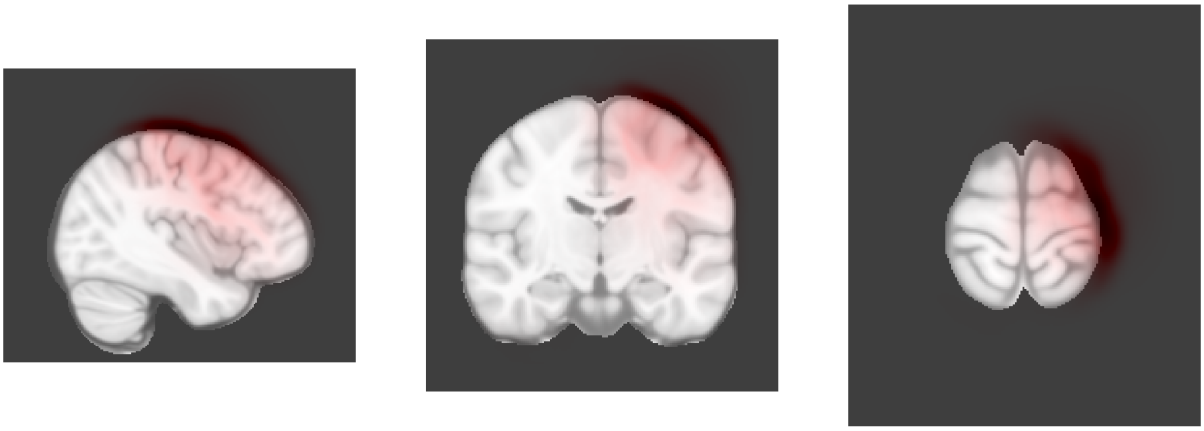

Dimension 2, Baseline

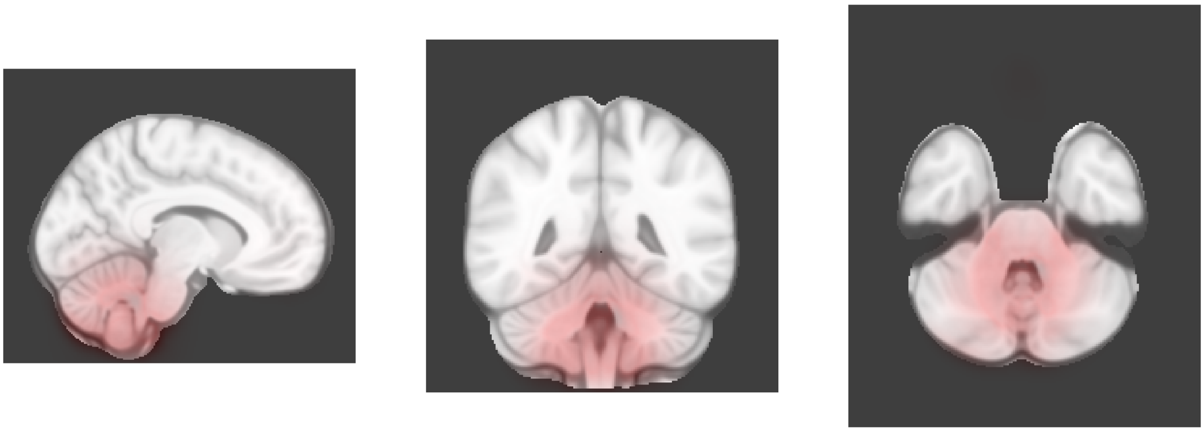

Dimension 2, 2-year follow-up

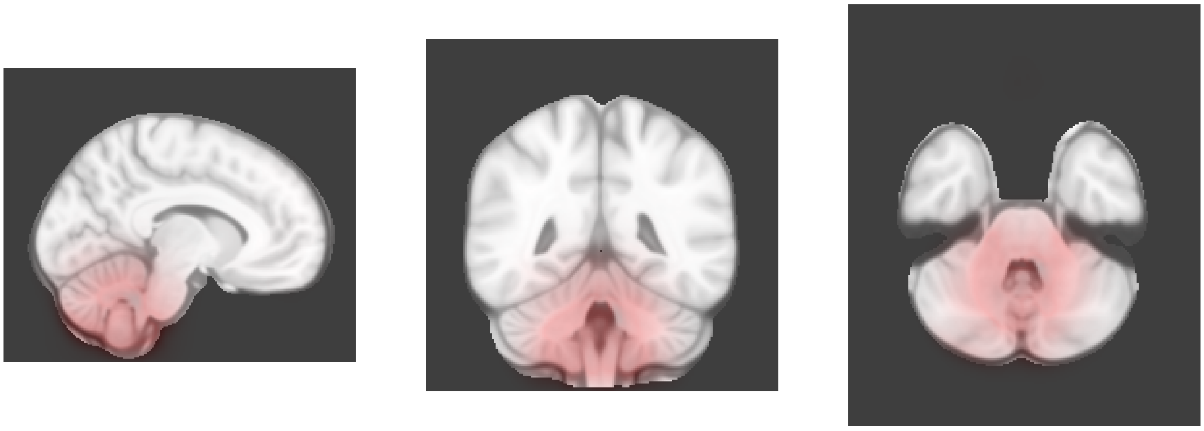

Dimension 3, Baseline

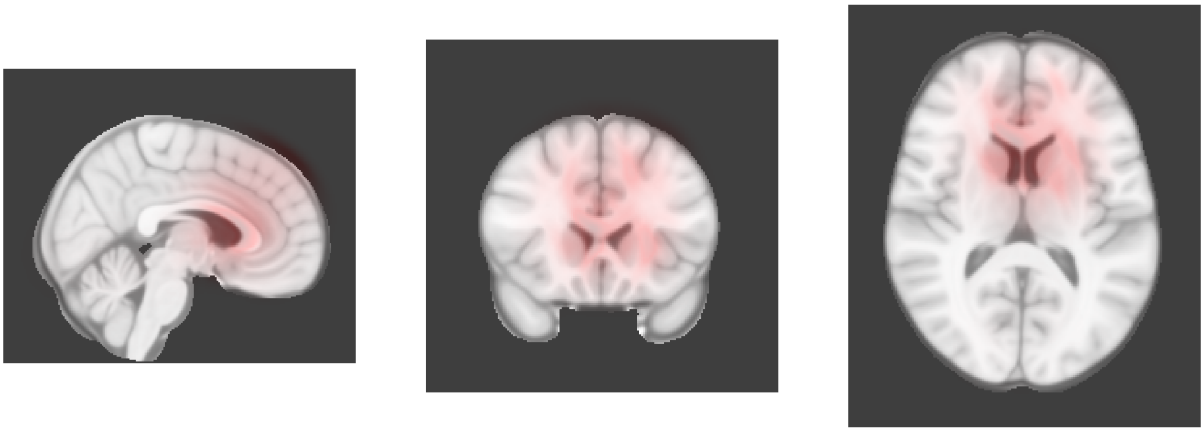

Dimension 3, 2-year follow-up

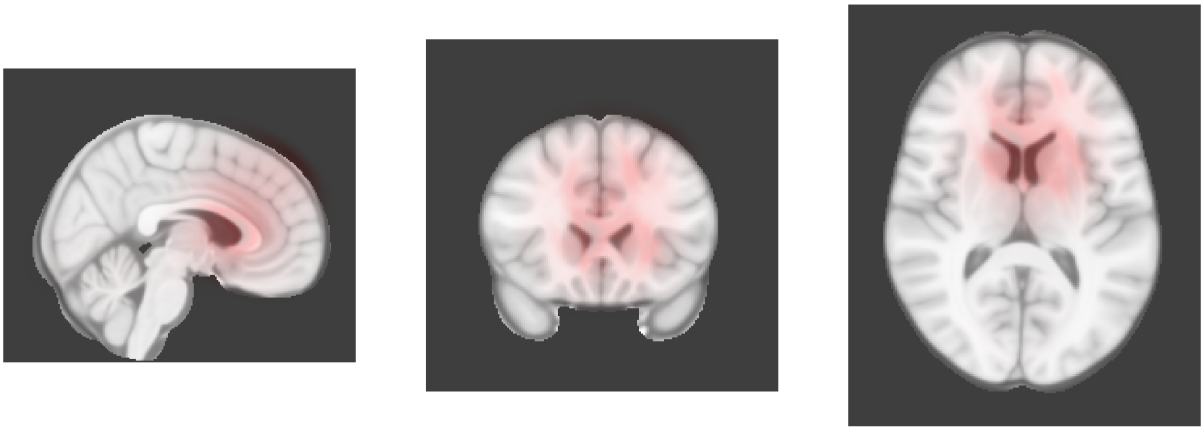

Dimension 4, Baseline

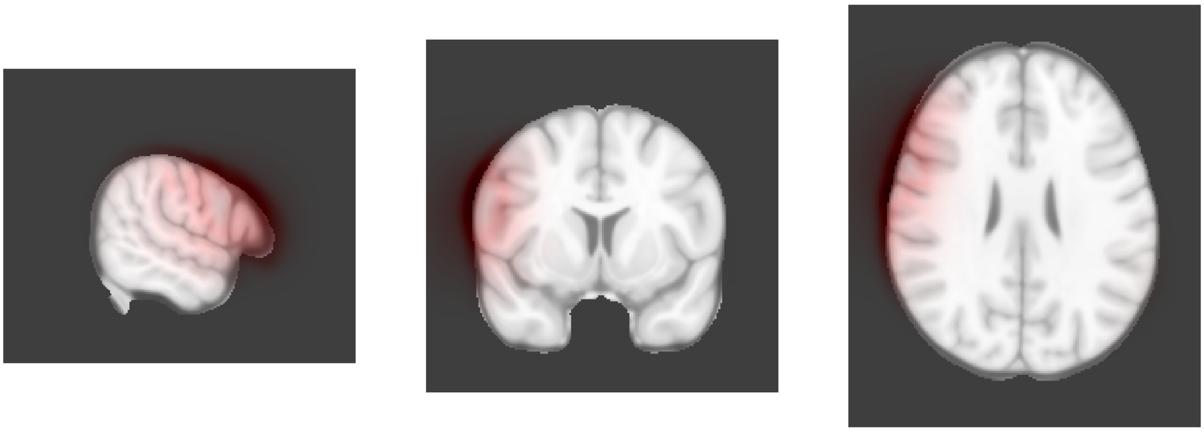

Dimension 4, 2-year follow-up

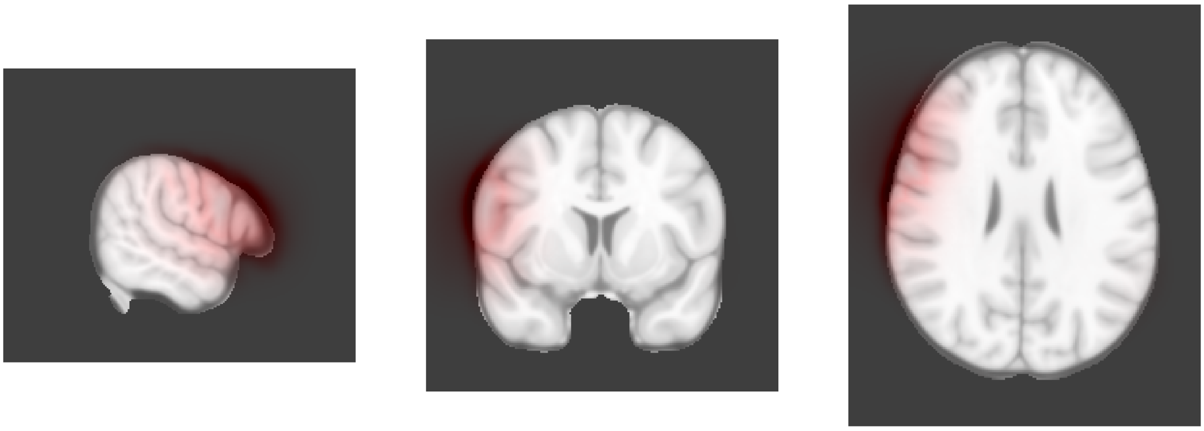

Dimension 5, Baseline

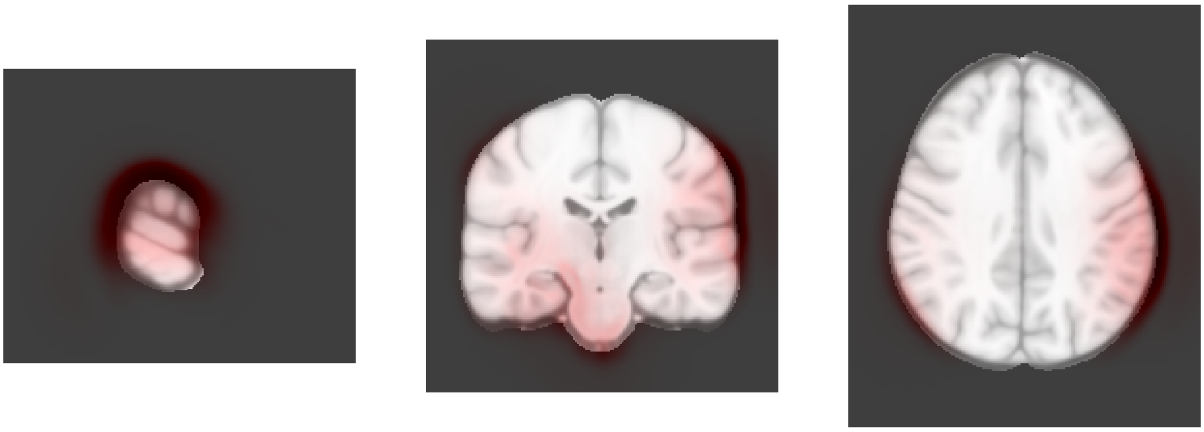

Dimension 5, 2-year follow-up

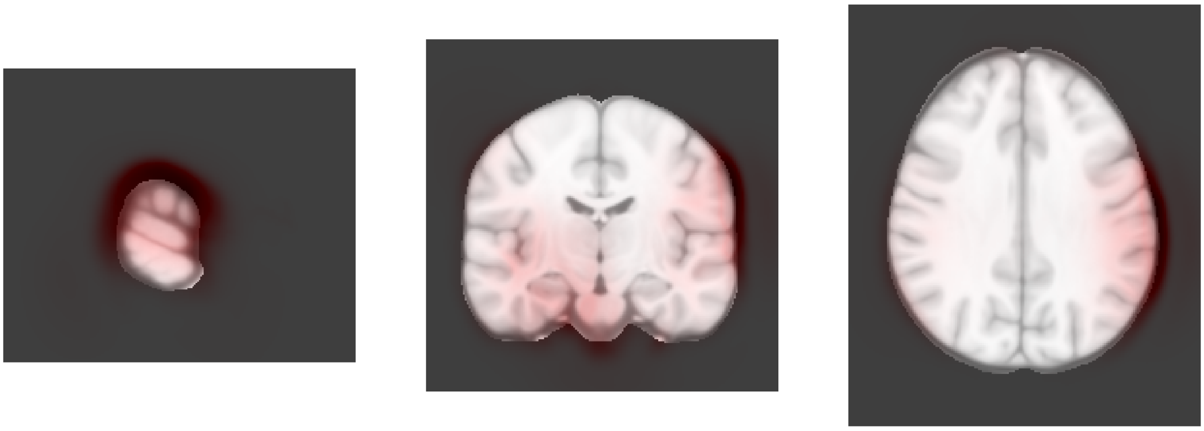

Dimension 6, Baseline

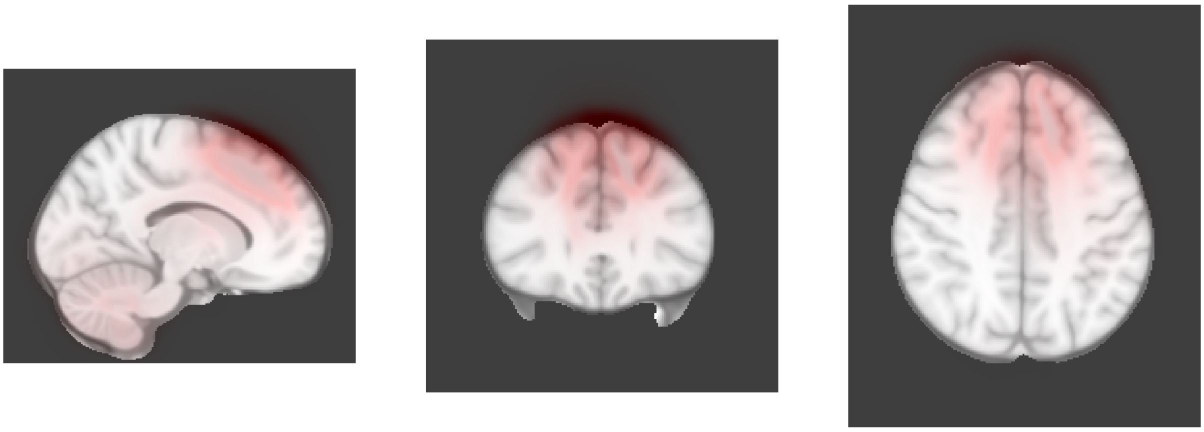

Dimension 6, 2-year follow-up

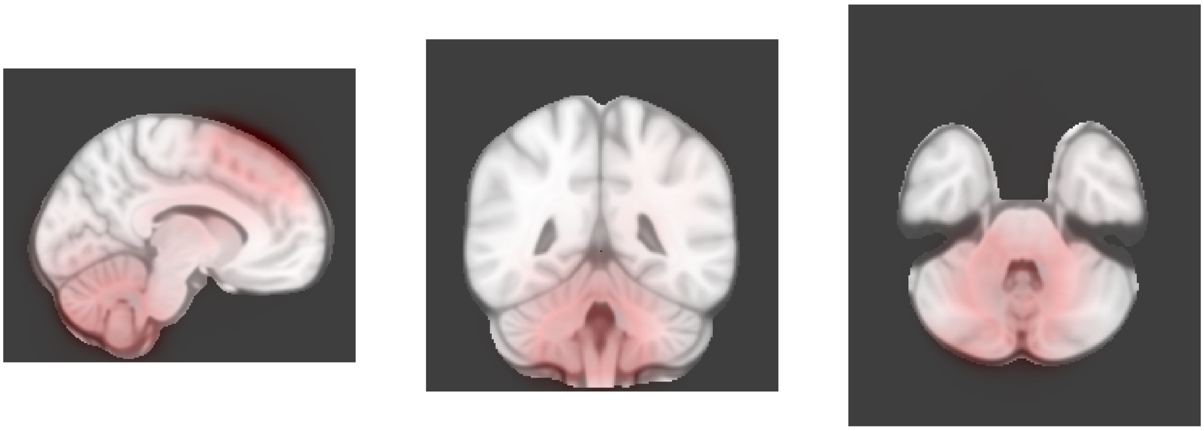

Dimension 7, Baseline

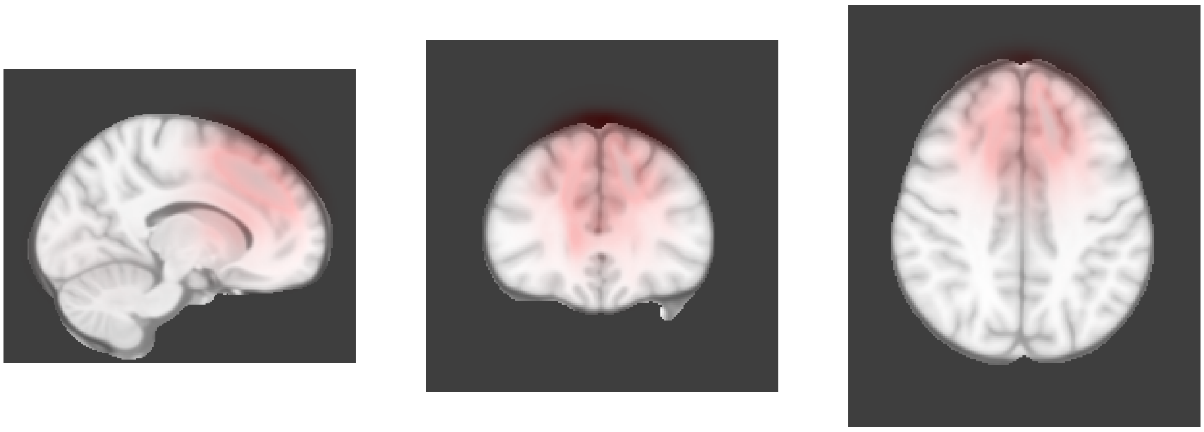

Dimension 7, 2-year follow-up

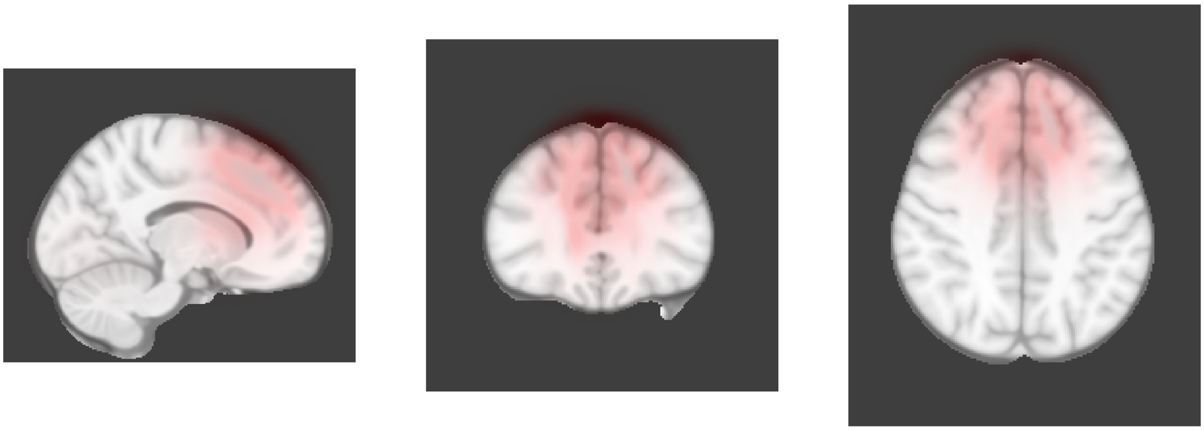

Dimension 8, Baseline

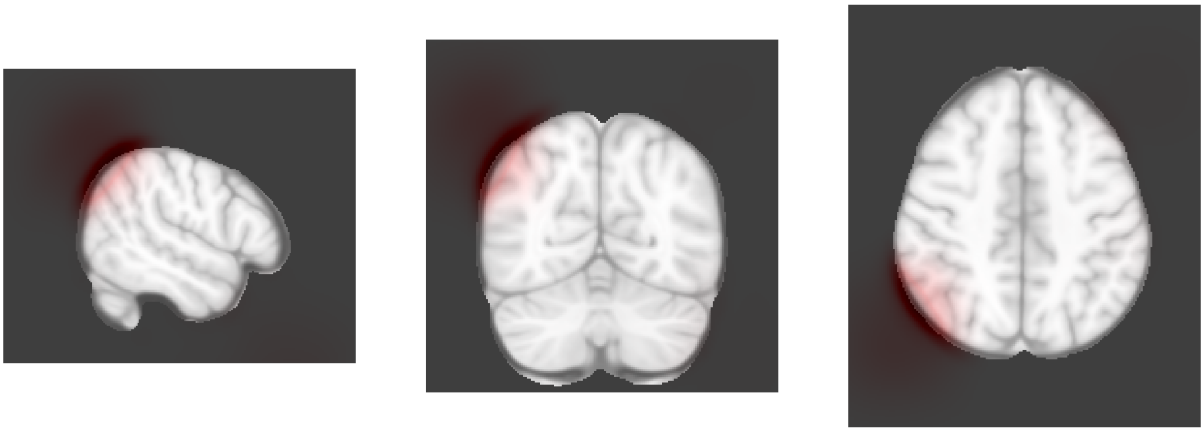

Dimension 8, 2-year follow-up

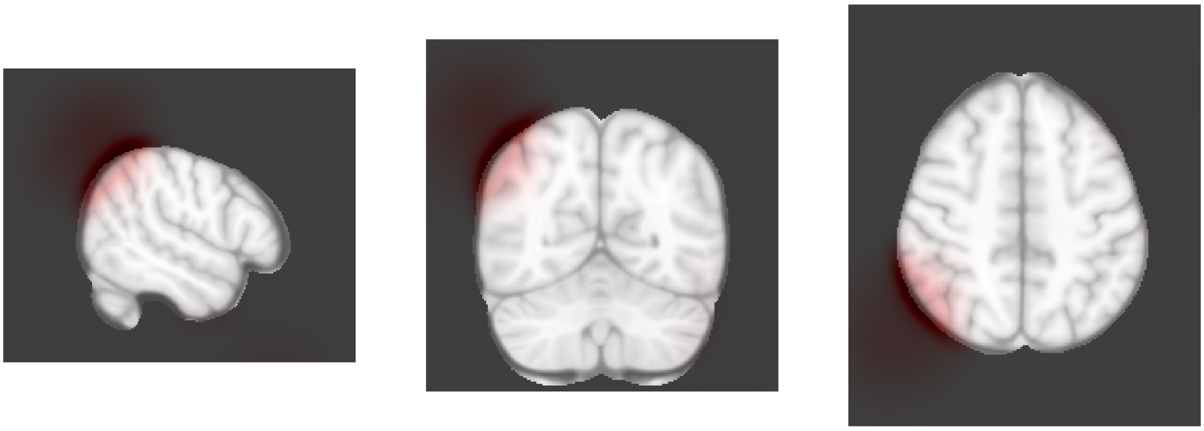

Dimension 9, Baseline

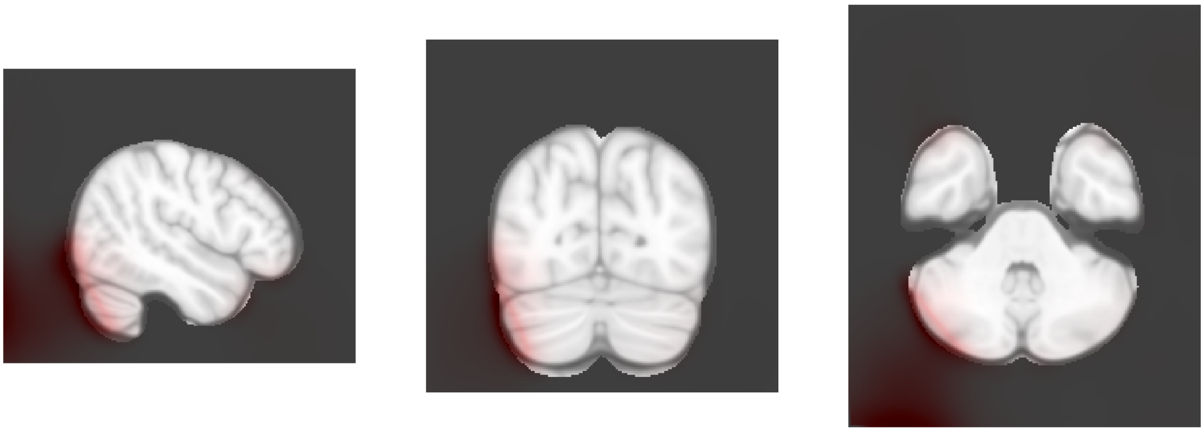

Dimension 9, 2-year follow-up

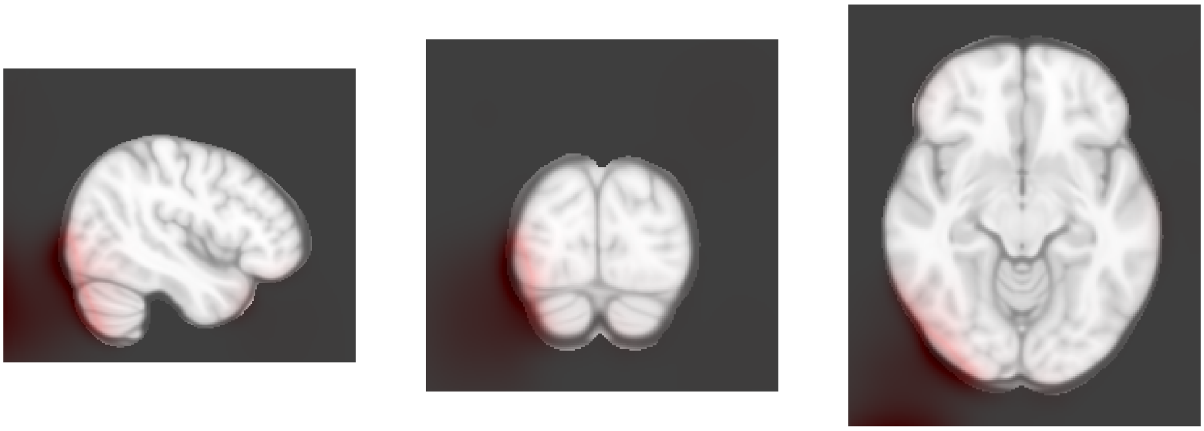

Dimension 10, Baseline

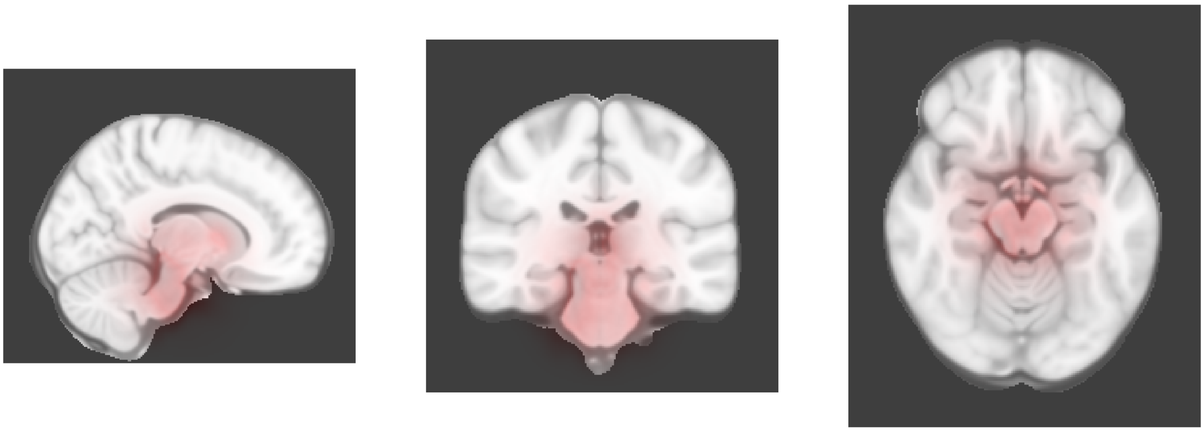

Dimension 10, 2-year follow-up

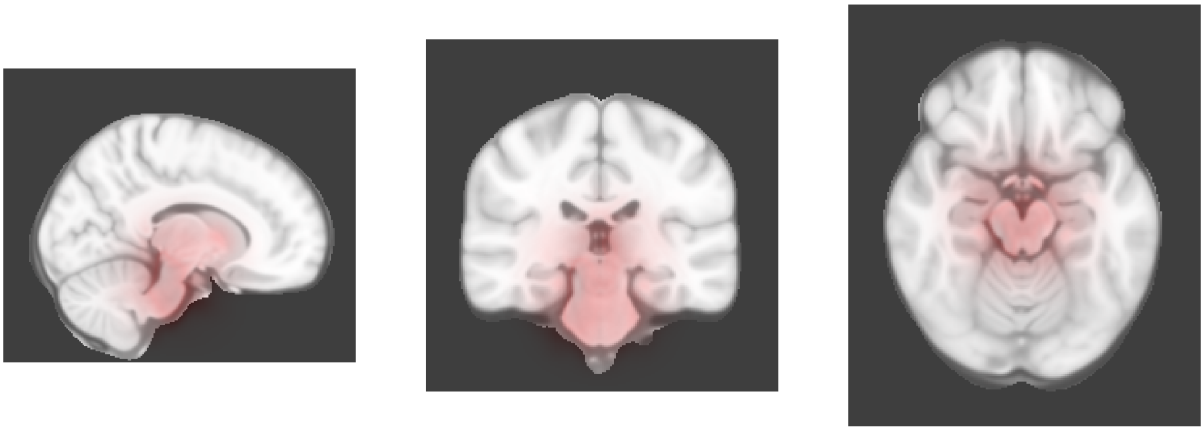

Dimension 11, Baseline

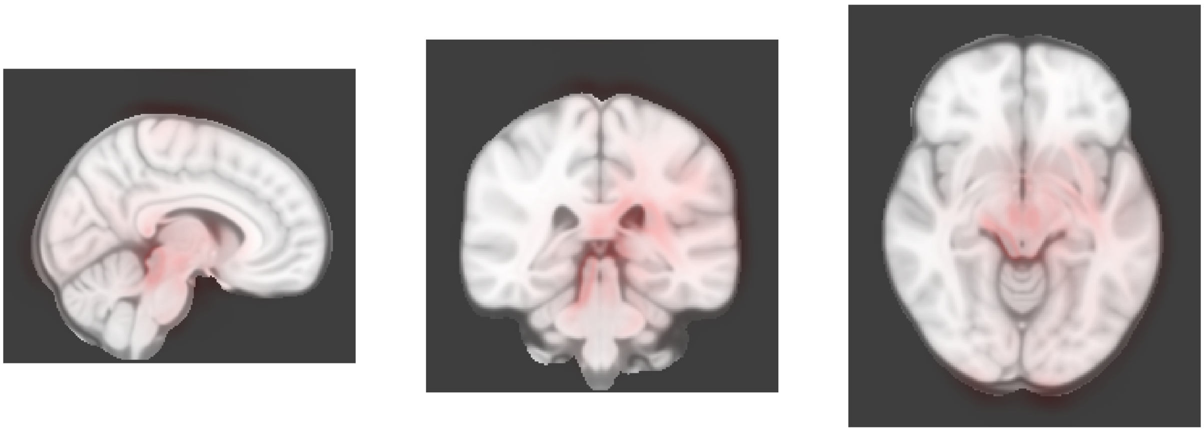

Dimension 11, 2-year follow-up

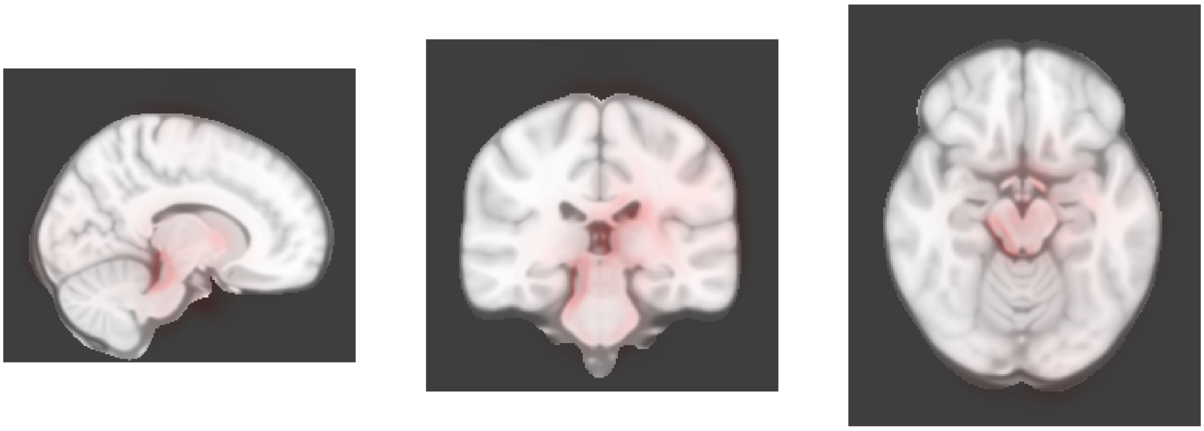

Dimension 12, Baseline

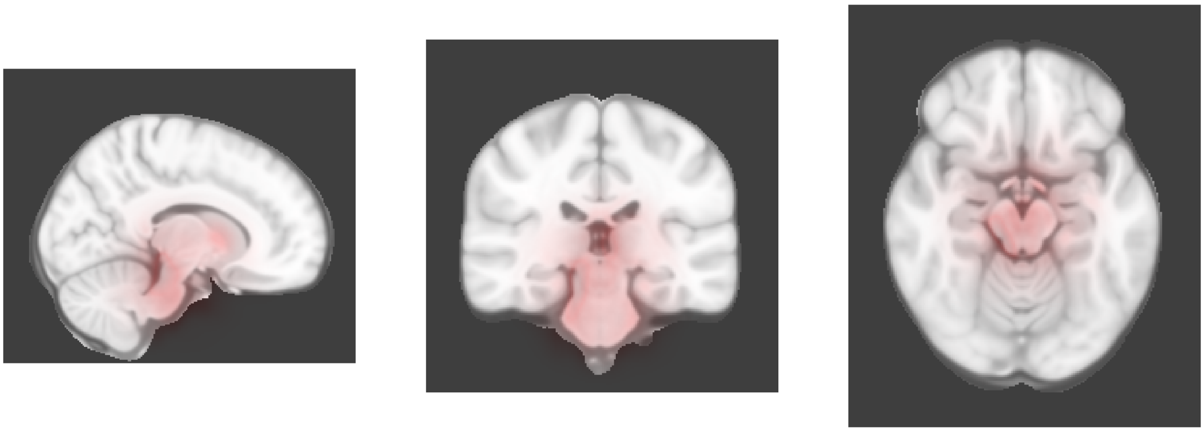

Dimension 12, 2-year follow-up

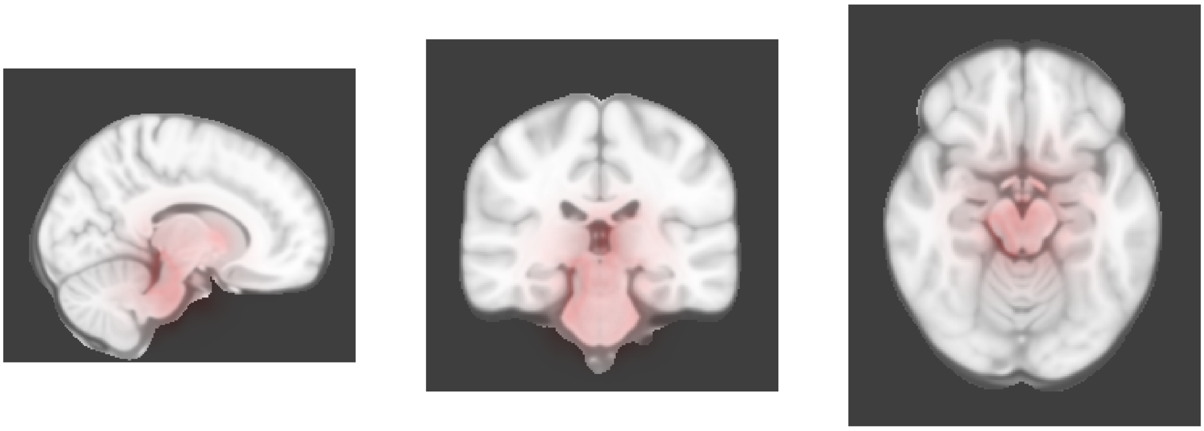

Dimension 13, Baseline

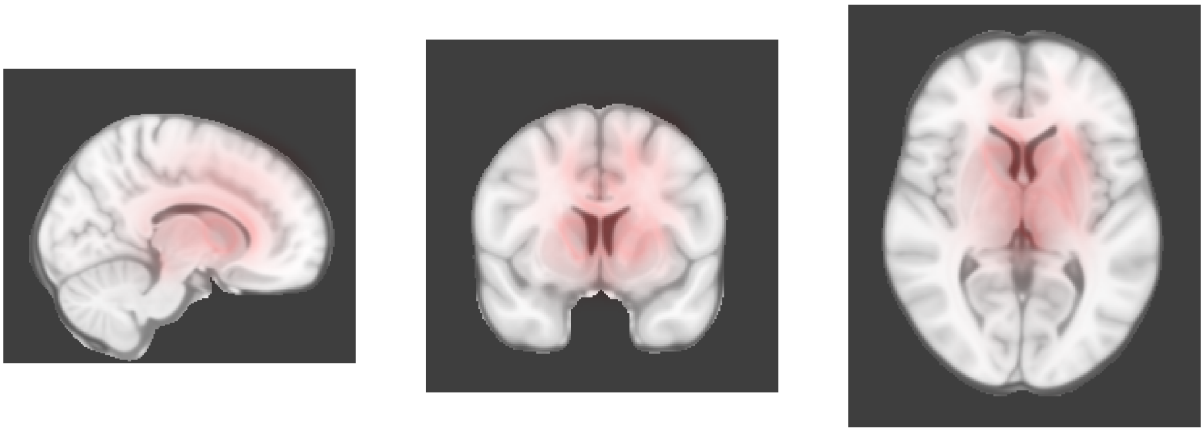

Dimension 13, 2-year follow-up

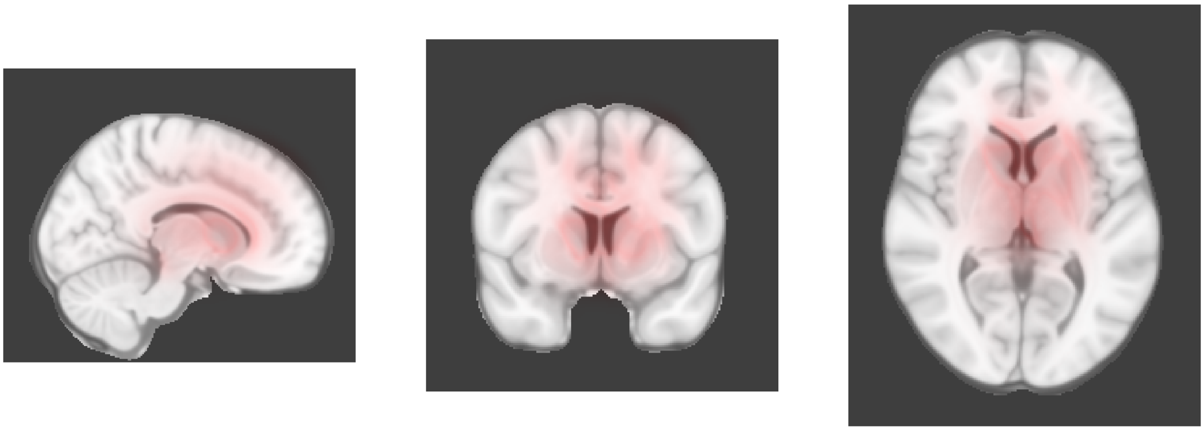

Dimension 14, Baseline

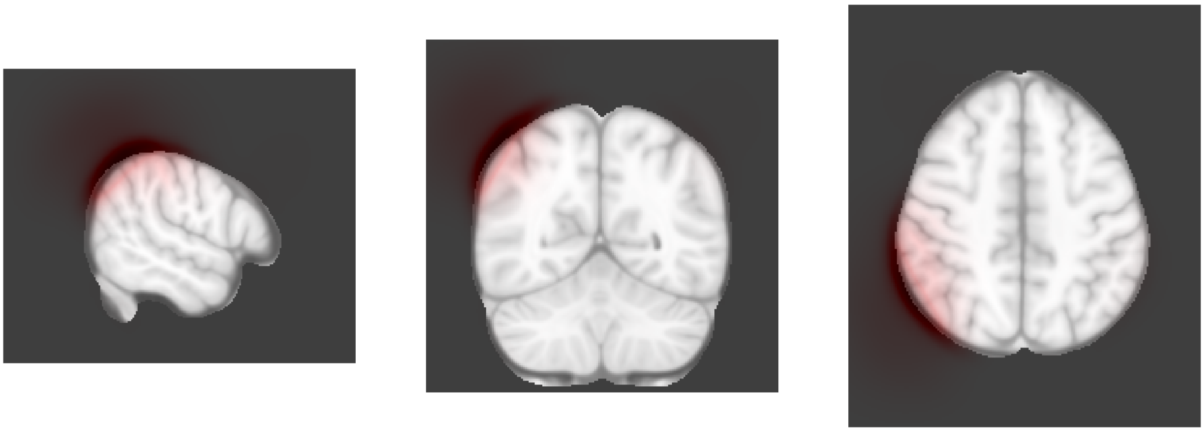

Dimension 14, 2-year follow-up

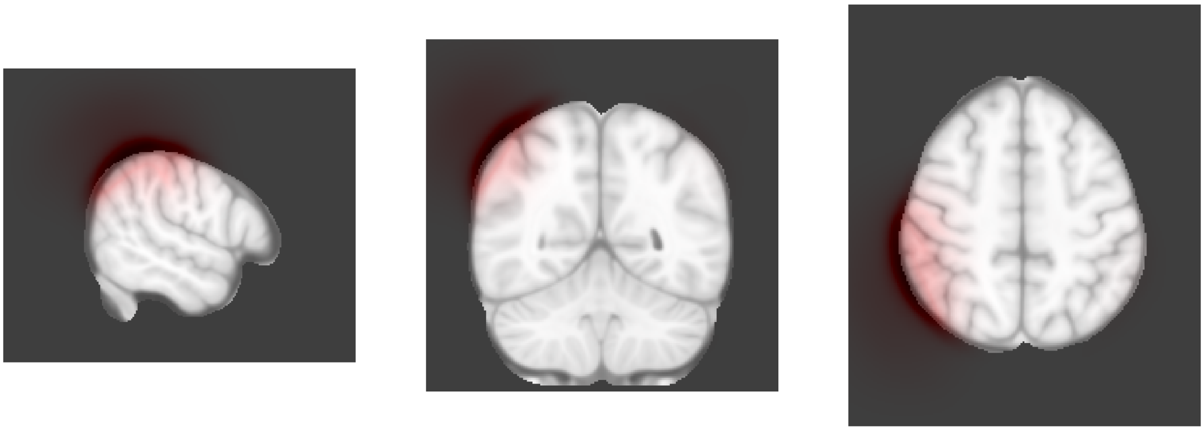

Dimension 15, Baseline

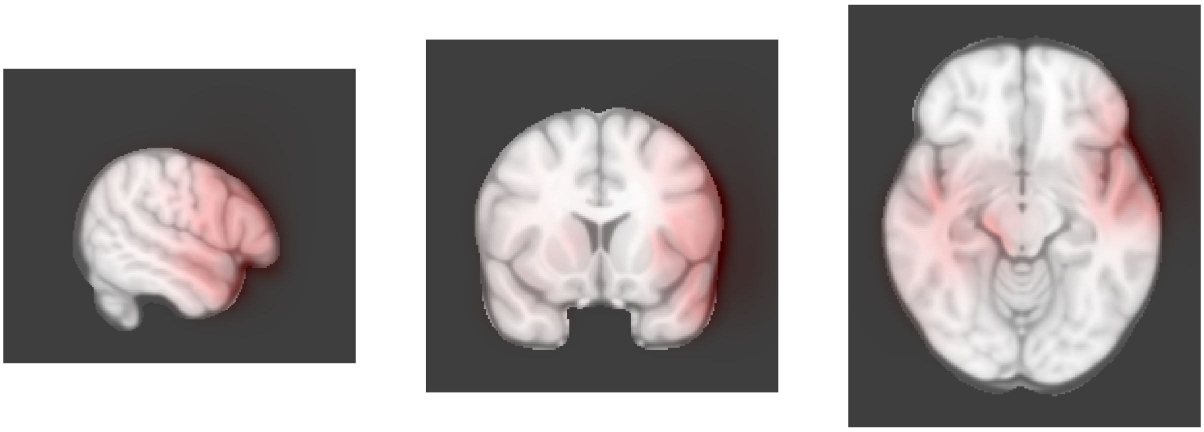

Dimension 15, 2-year follow-up

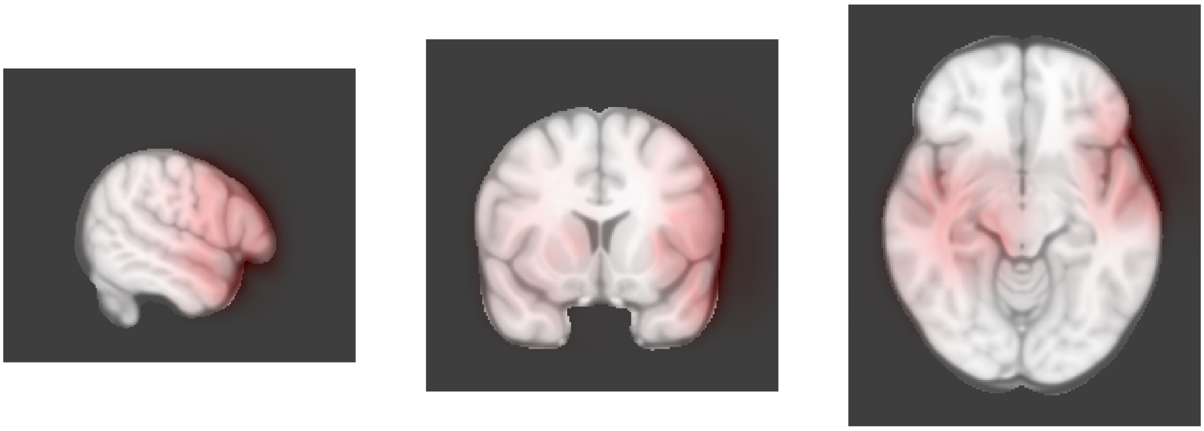

Dimension 16, Baseline

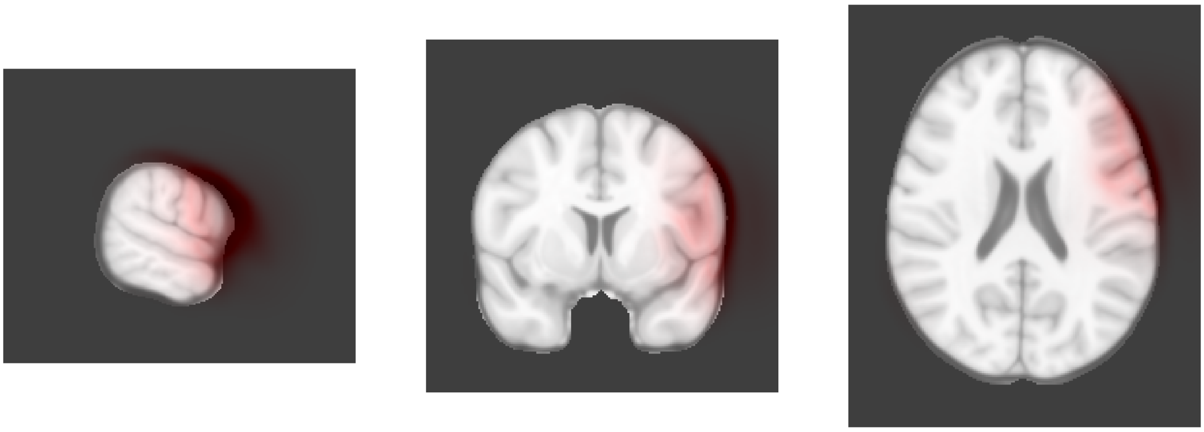

Dimension 16, 2-year follow-up

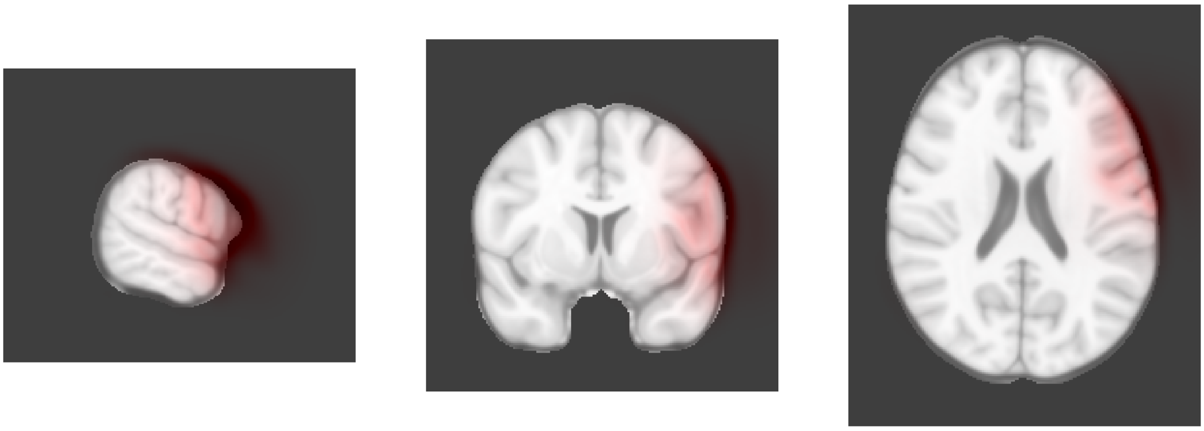

Dimension 17, Baseline

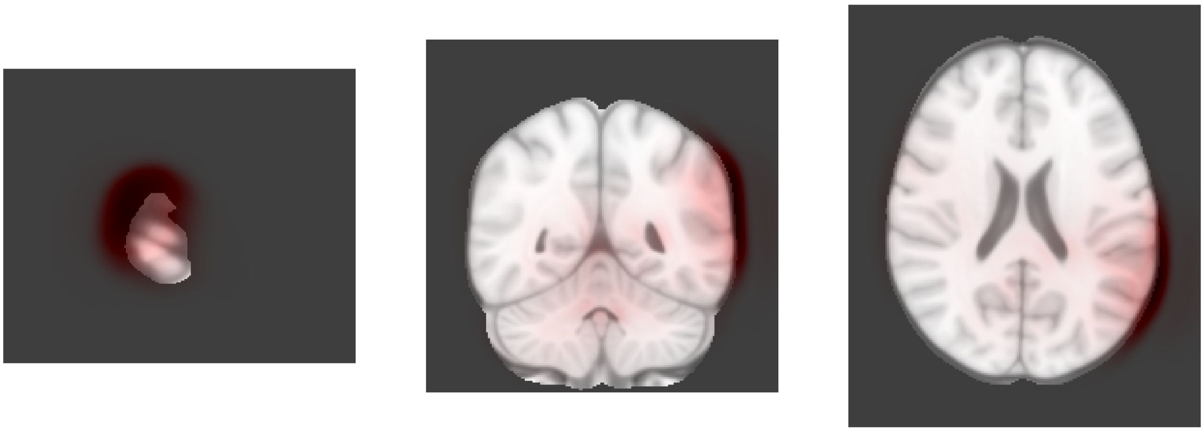

Dimension 17, 2-year follow-up

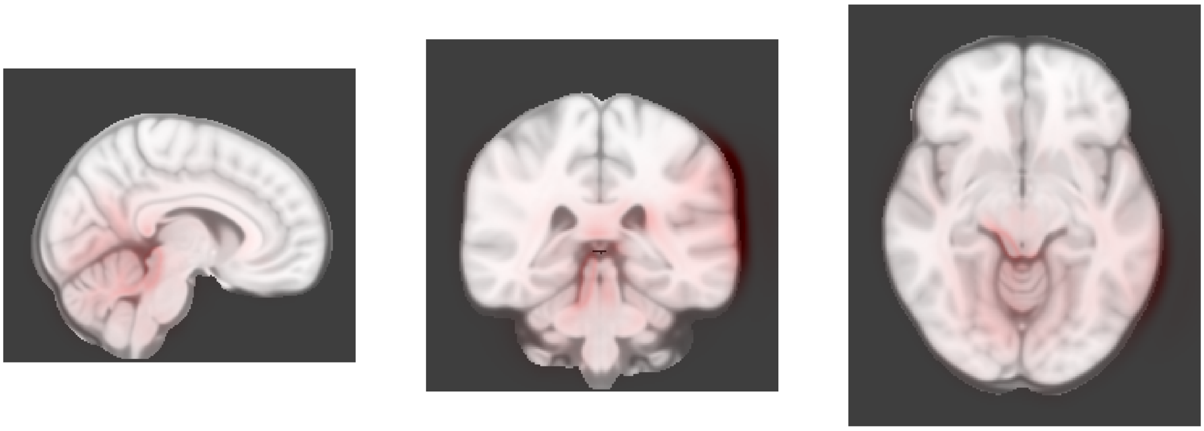

Dimension 18, Baseline

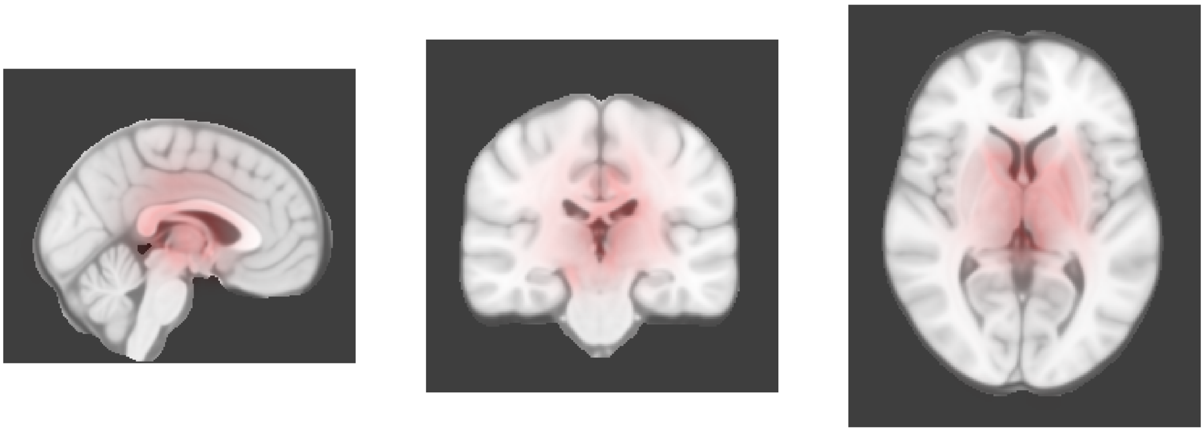

Dimension 18, 2-year follow-up

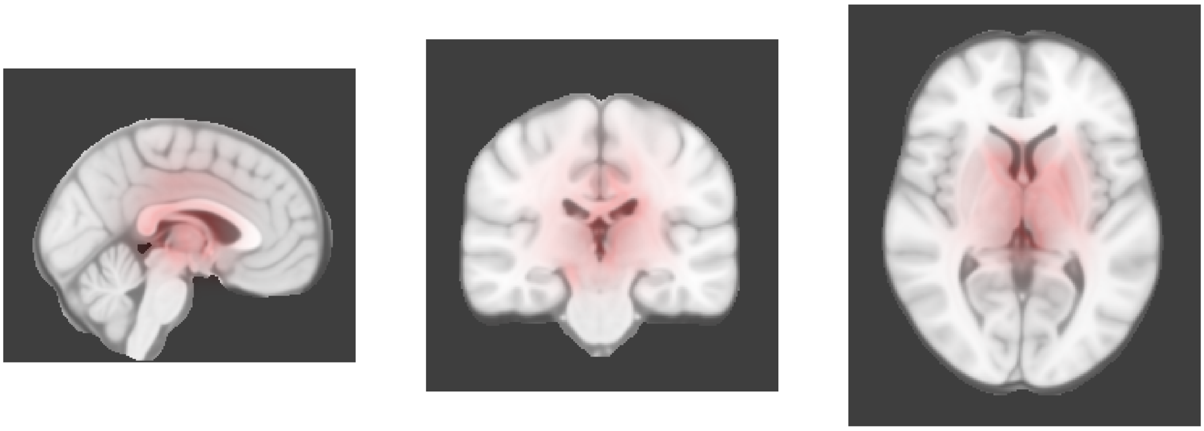

Dimension 19, Baseline

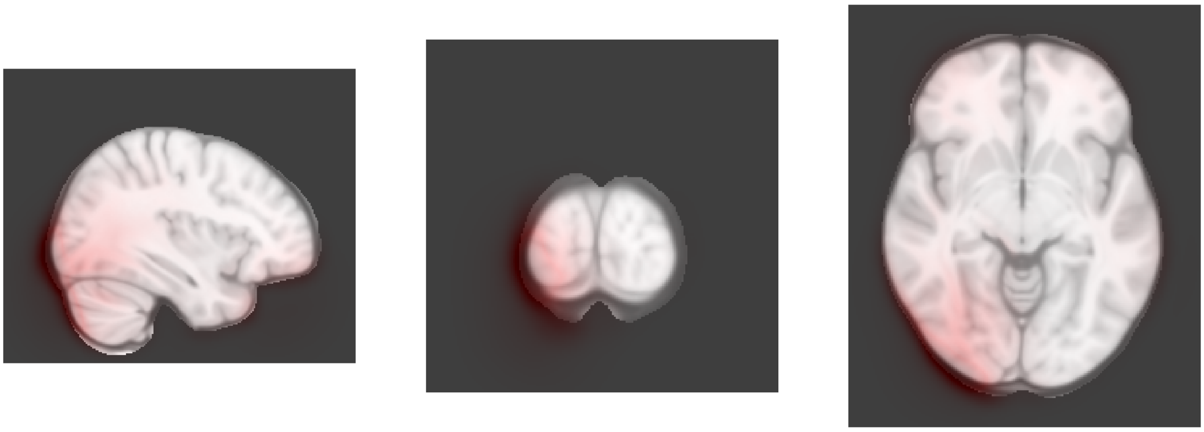

Dimension 19, 2-year follow-up

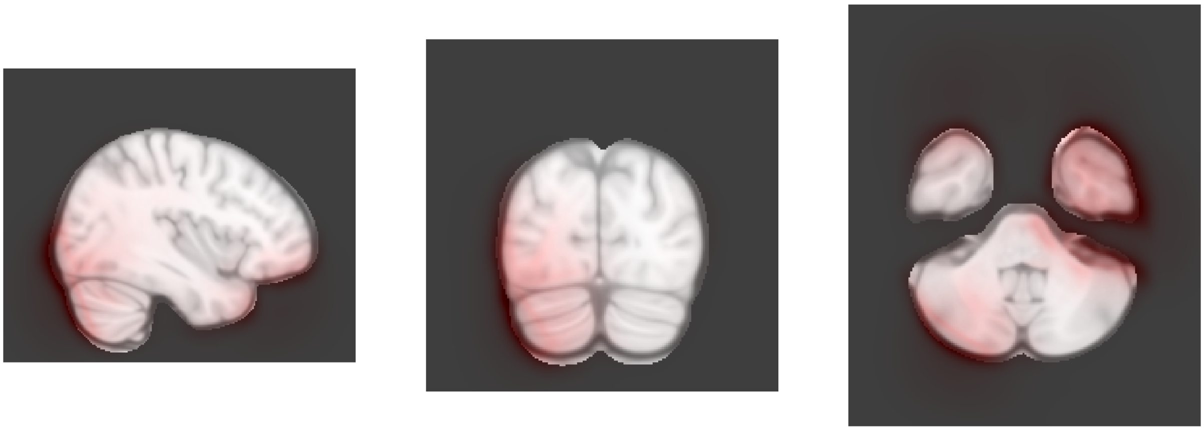

Dimension 20, Baseline

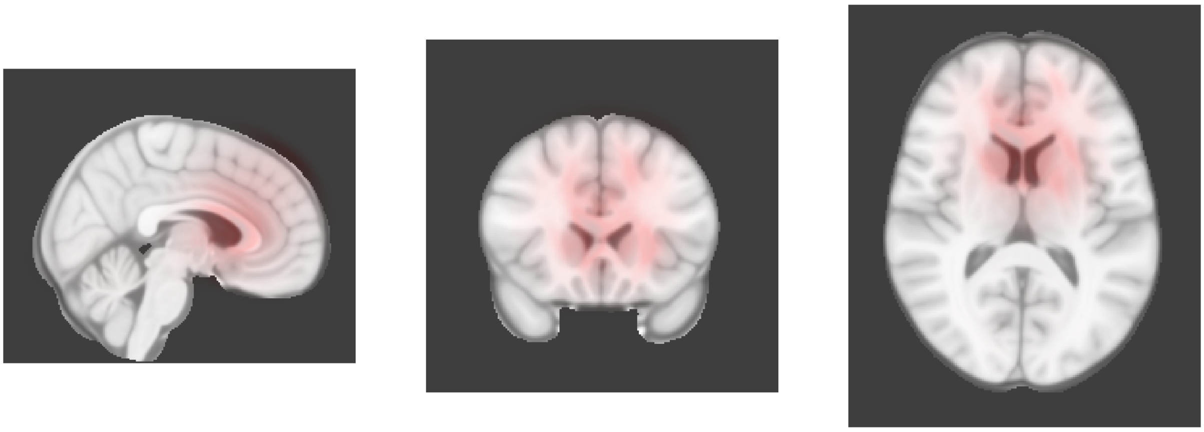

Dimension 20, 2-year follow-up

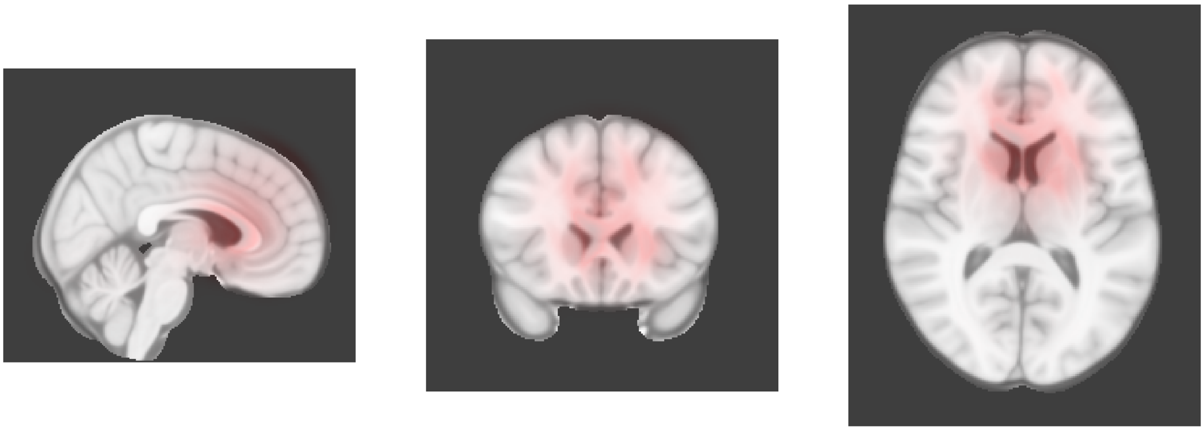

Dimension 21, Baseline

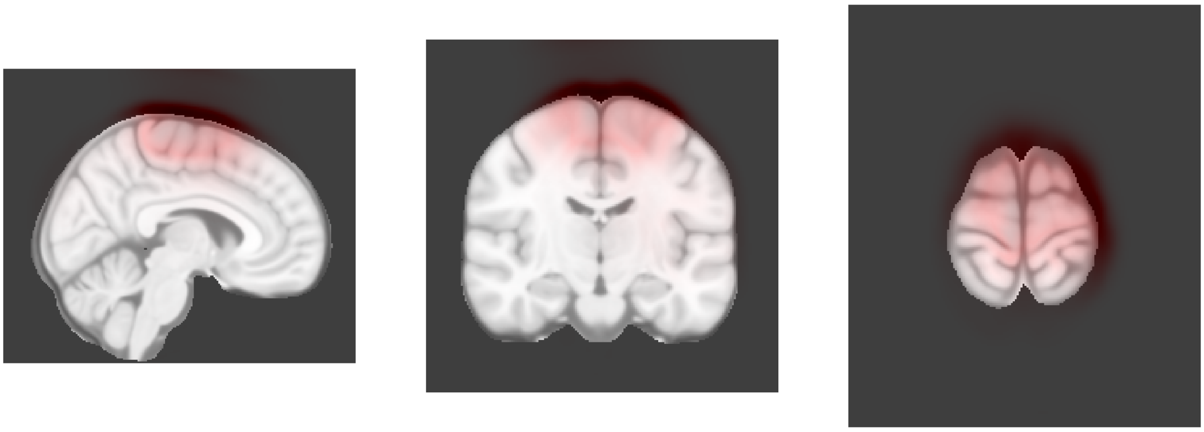

Dimension 21, 2-year follow-up

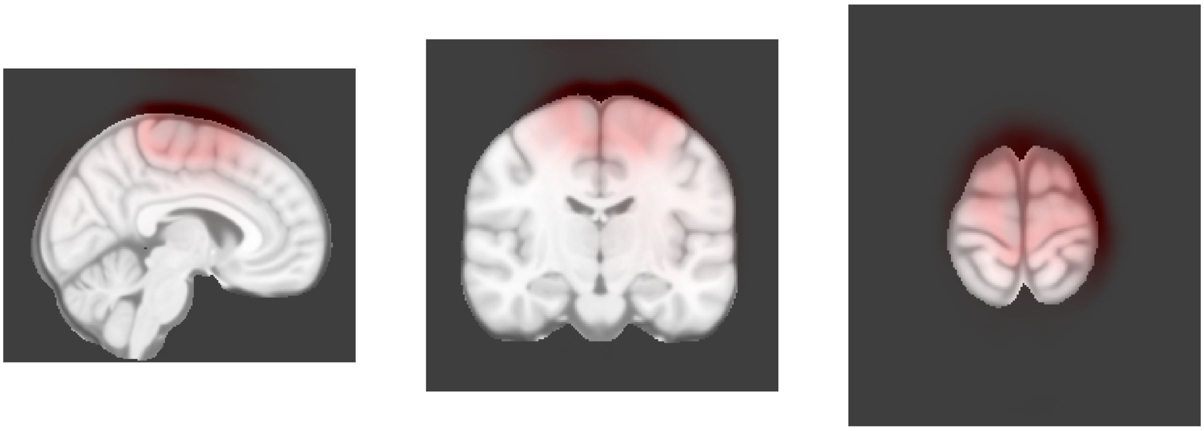

Dimension 22, Baseline

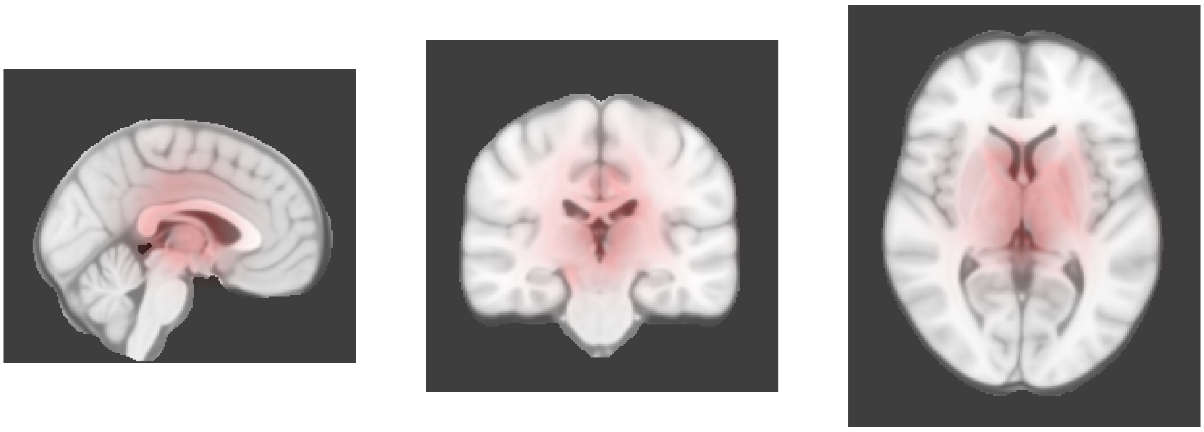

Dimension 22, 2-year follow-up

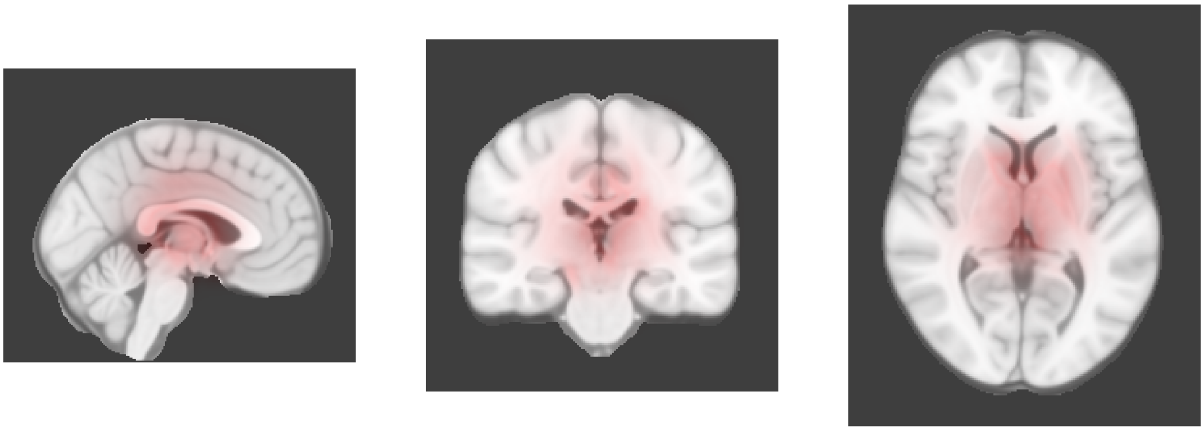

Dimension 23, Baseline

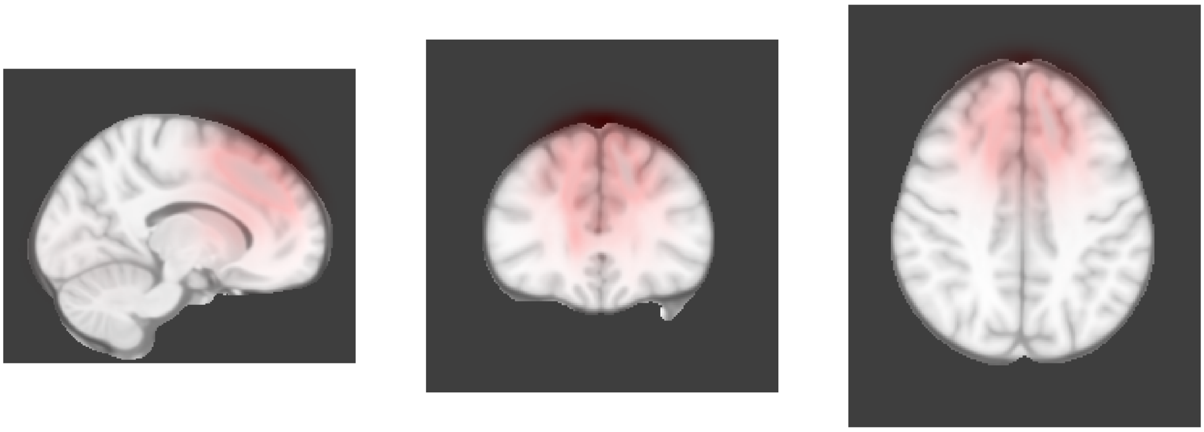

Dimension 23, 2-year follow-up

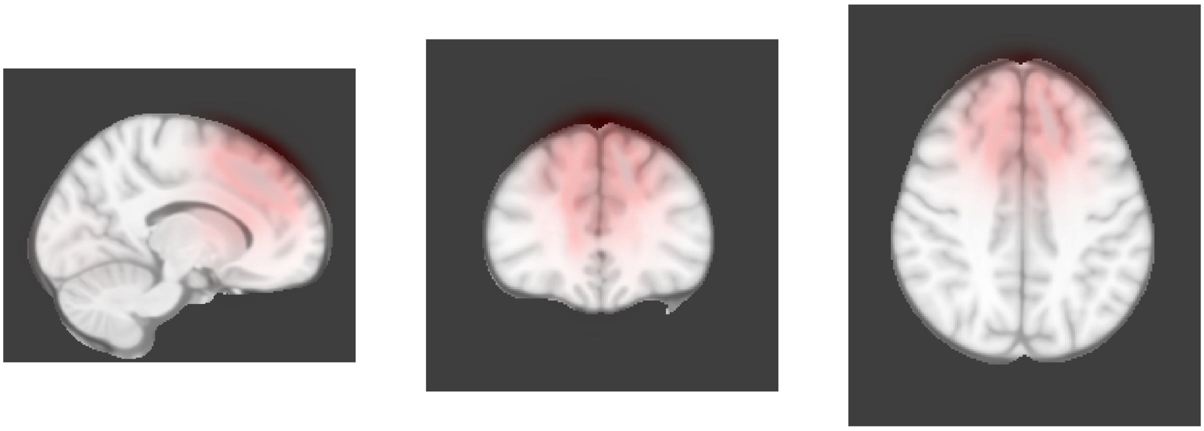

Dimension 24, Baseline

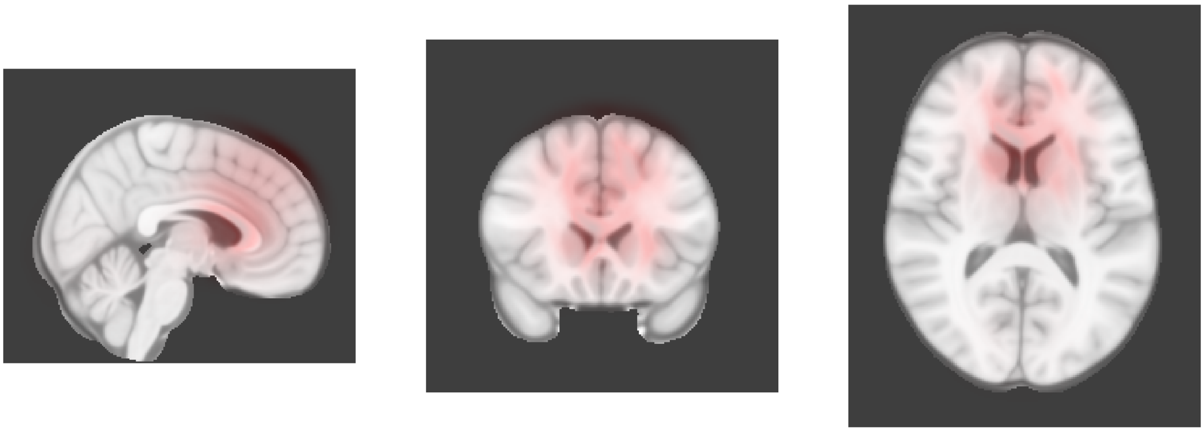

Dimension 24, 2-year follow-up

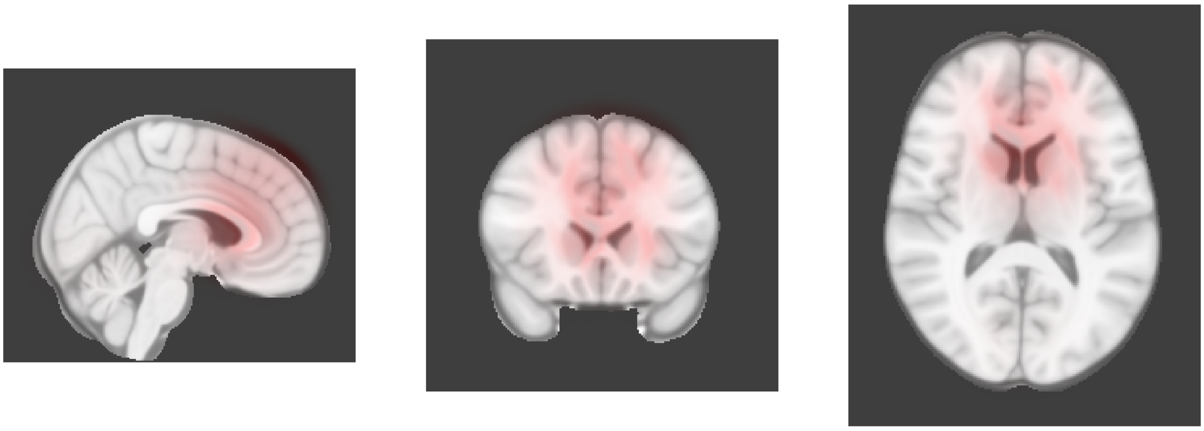

Dimension 25, Baseline

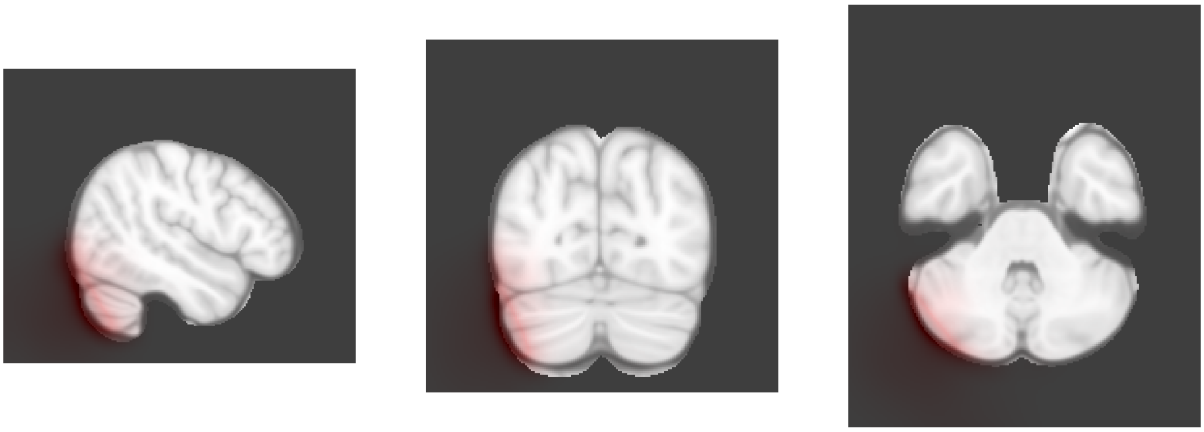

Dimension 25, 2-year follow-up

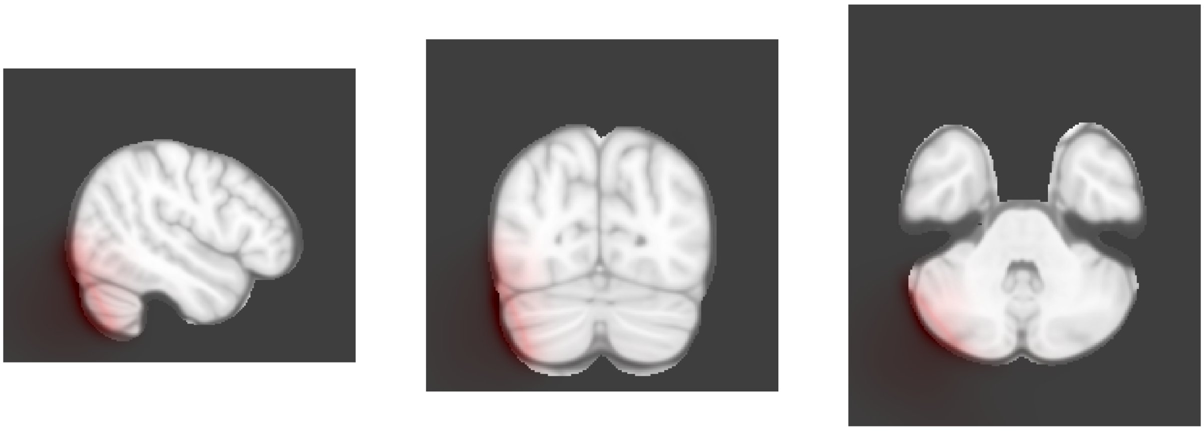

Dimension 26, Baseline

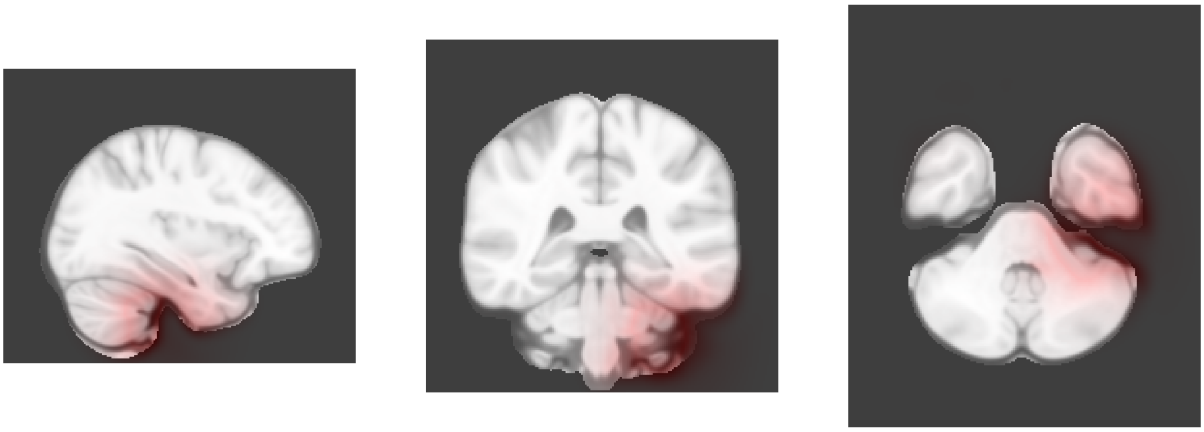

Dimension 26, 2-year follow-up

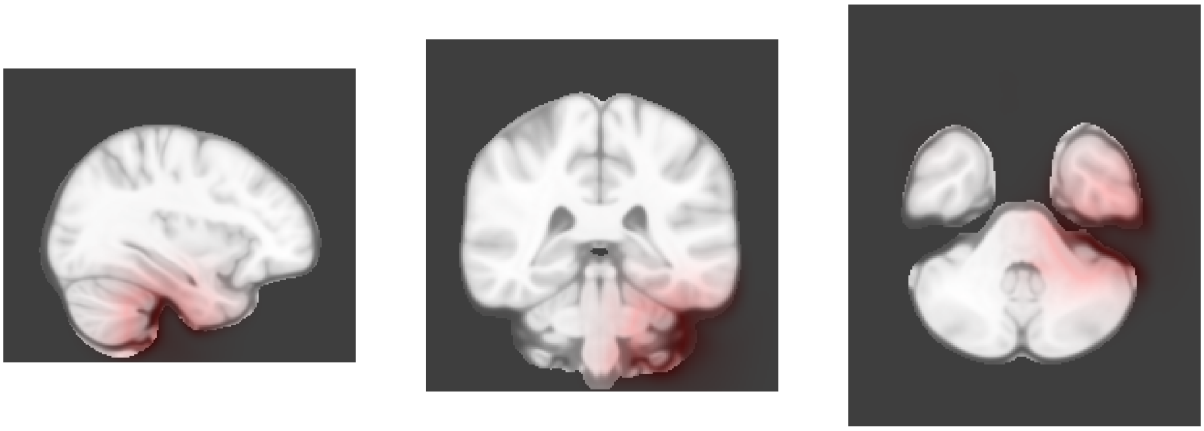

Dimension 27, Baseline

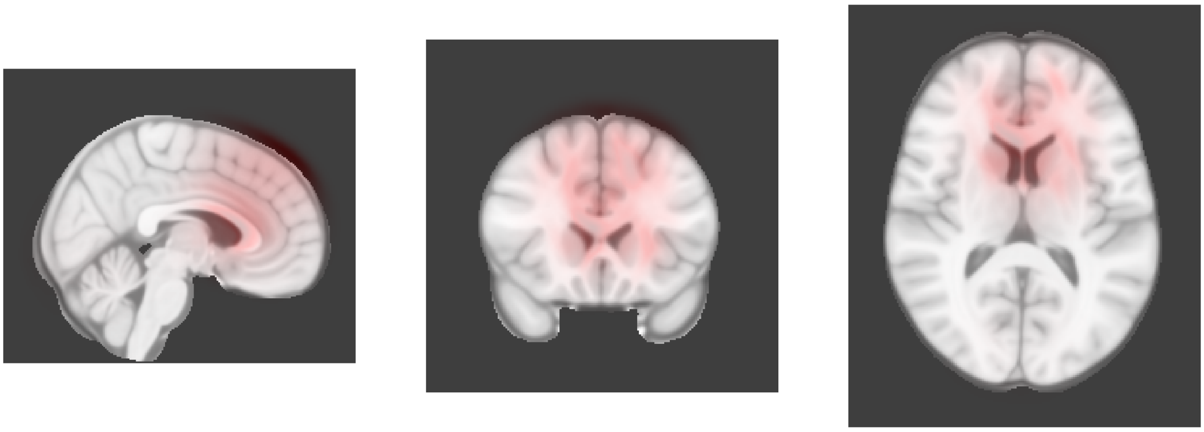

Dimension 27, 2-year follow-up

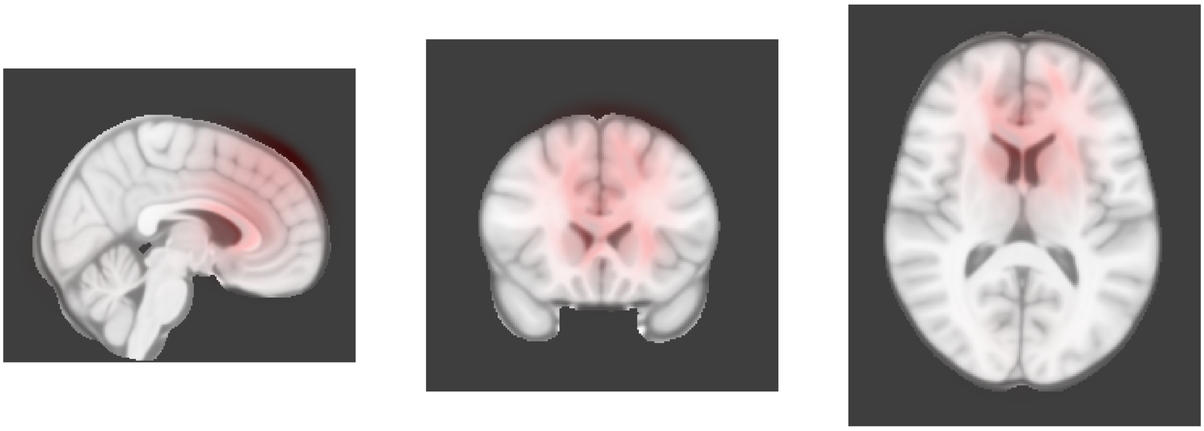

Dimension 29, Baseline

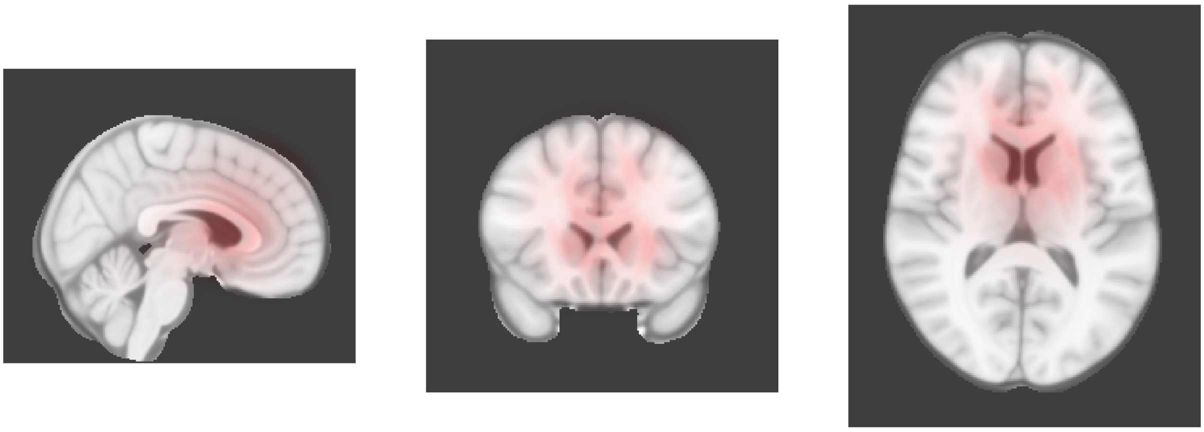

Dimension 29, 2-year follow-up

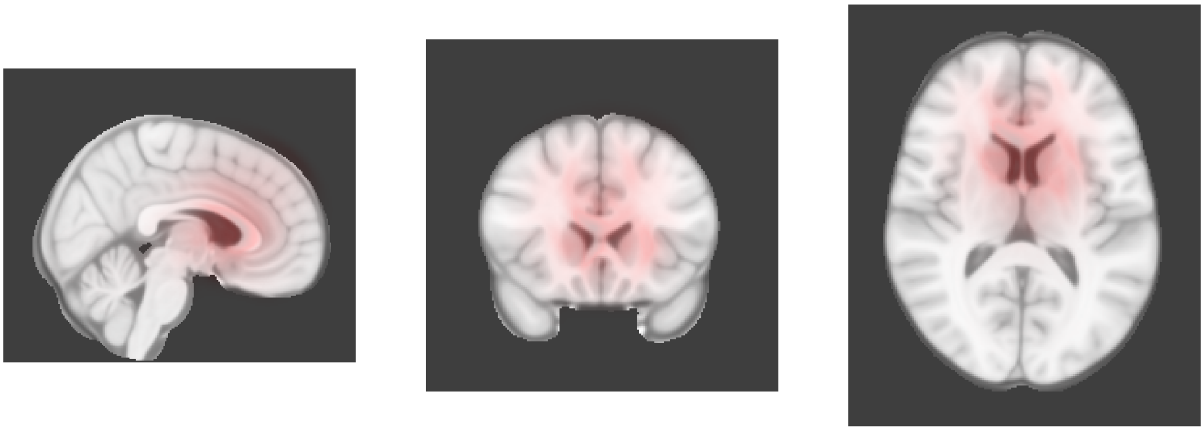

Dimension 30, Baseline

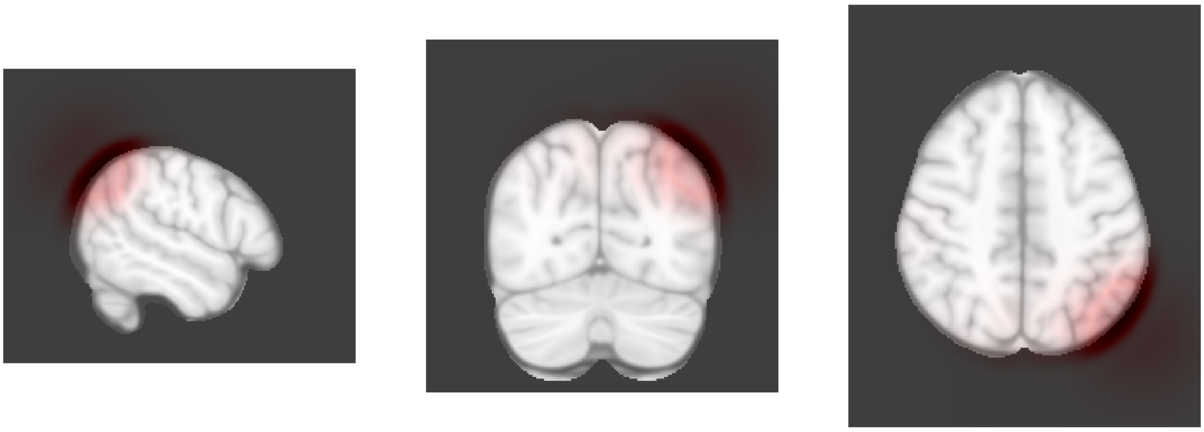

Dimension 30, 2-year follow-up

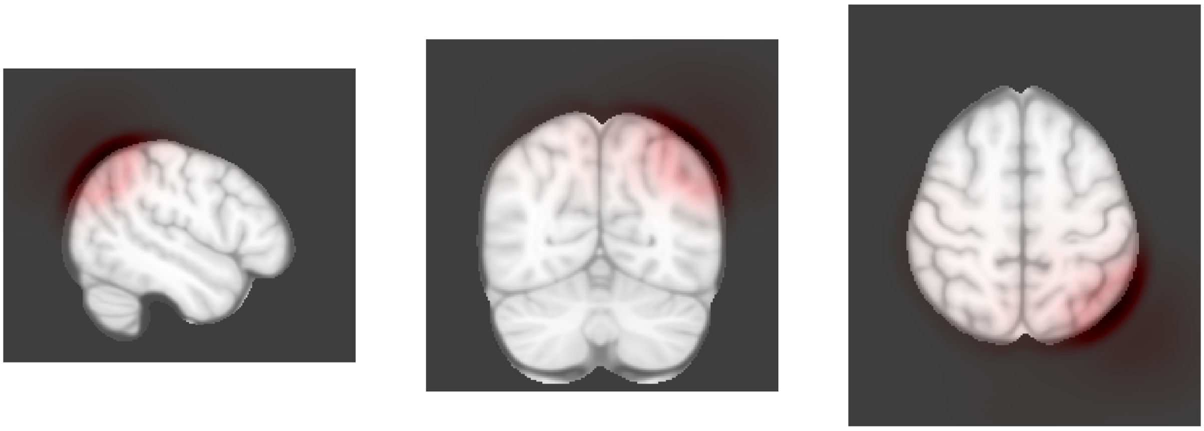

Dimension 31, Baseline

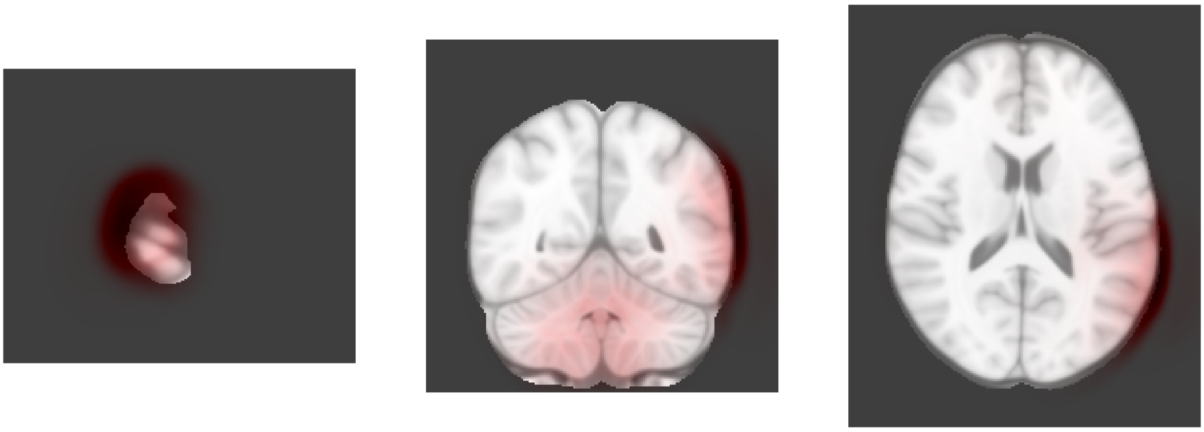

Dimension 31, 2-year follow-up

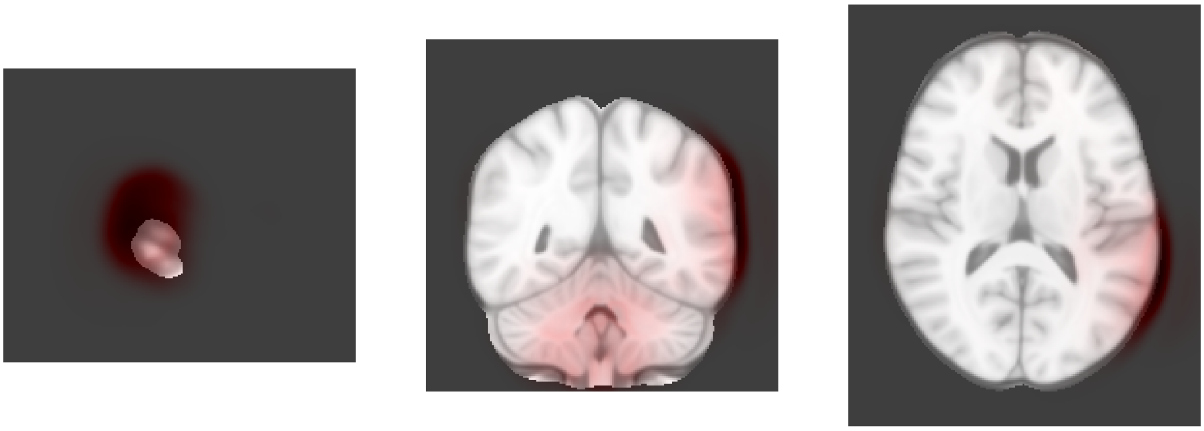

Dimension 32, Baseline

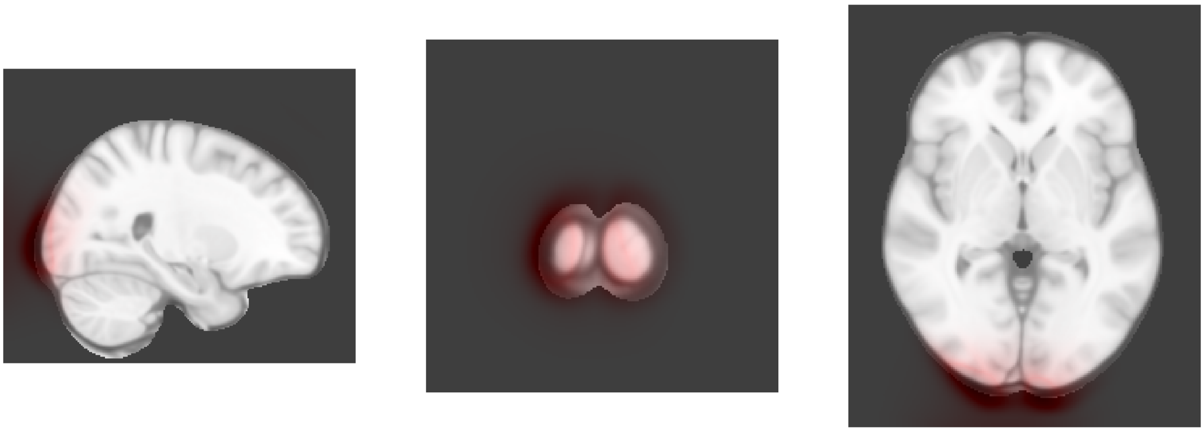

Dimension 32, 2-year follow-up

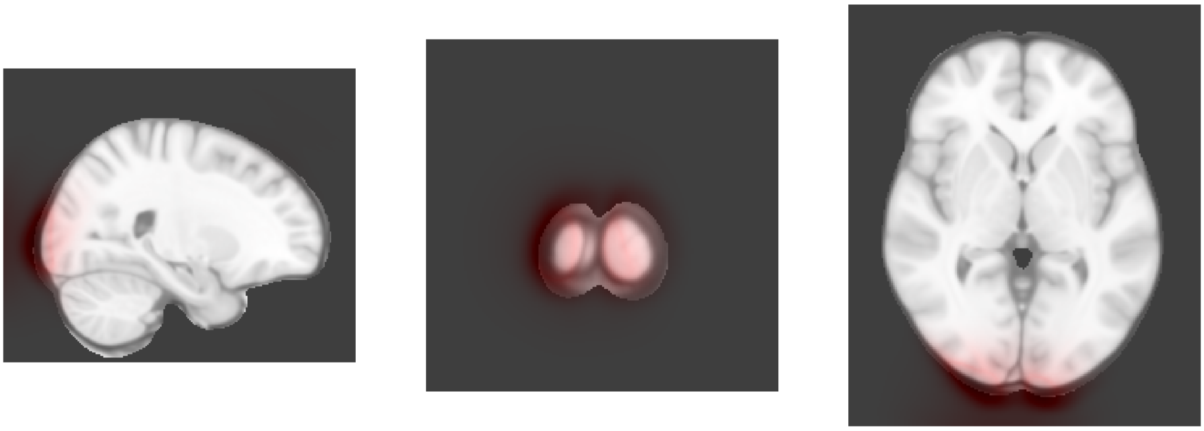

Dimension 33, Baseline

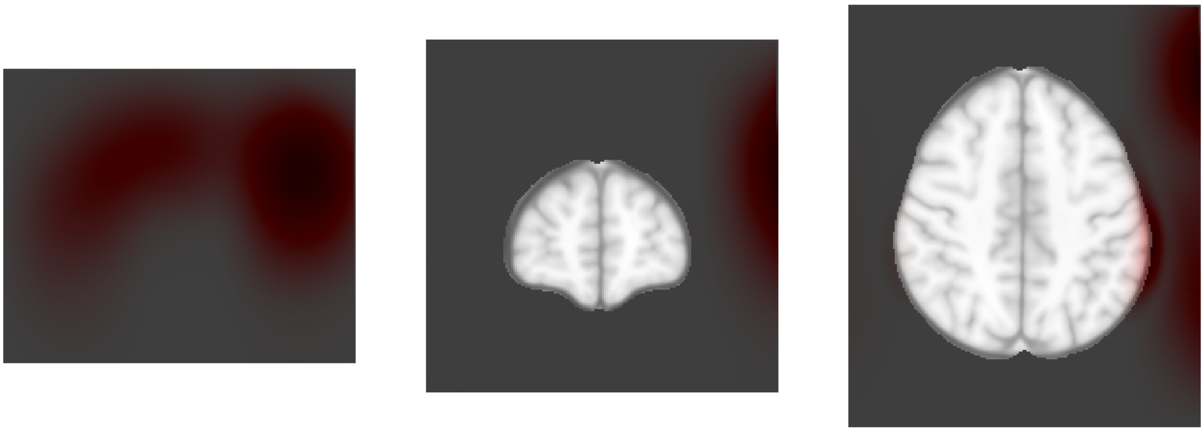

Dimension 33, 2-year follow-up

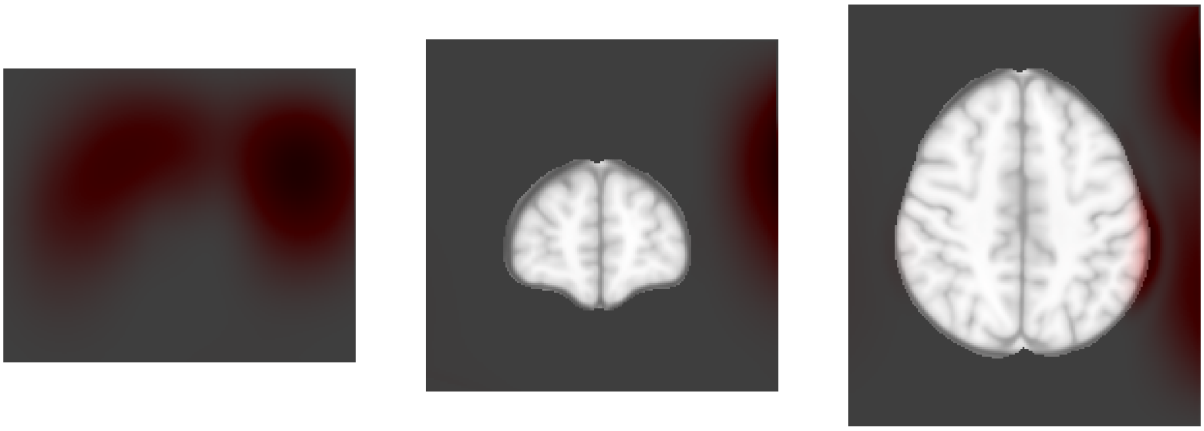

Dimension 34, Baseline

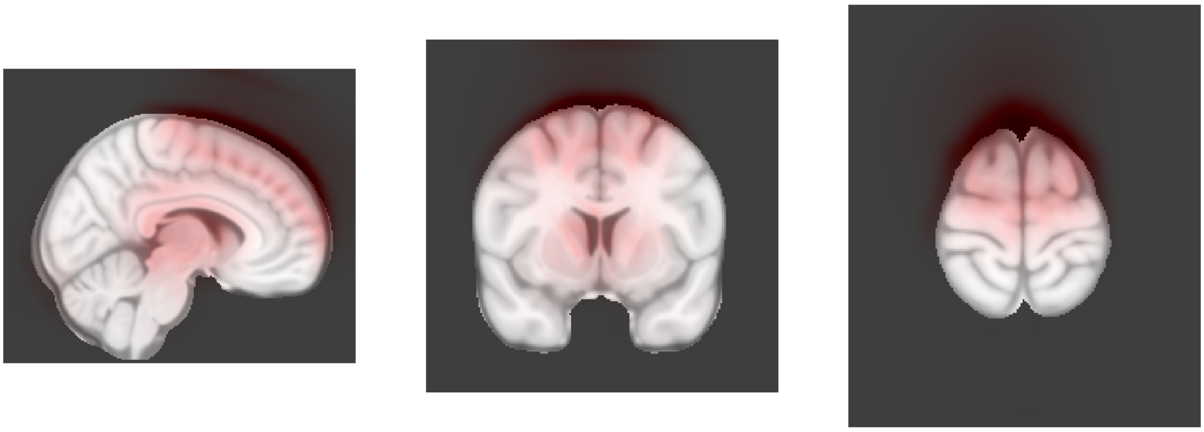

Dimension 34, 2-year follow-up

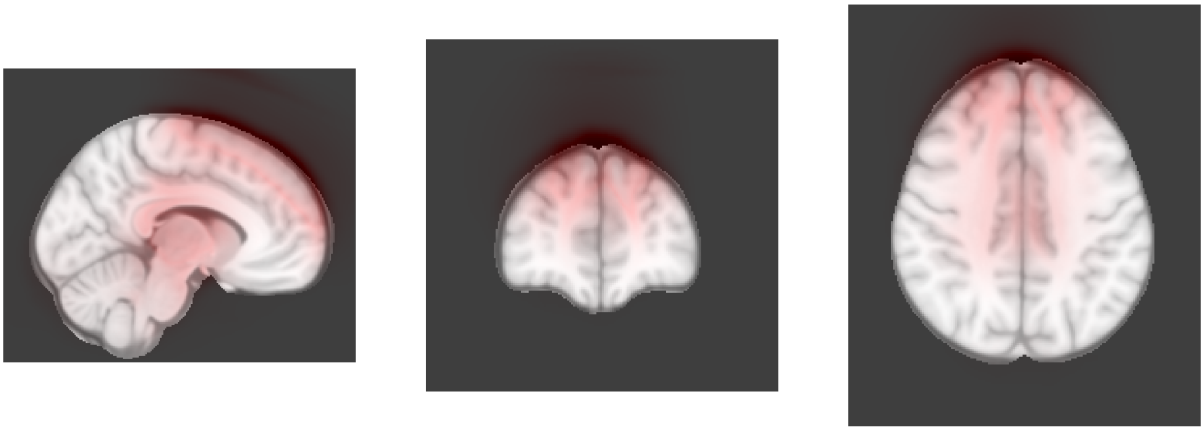

Dimension 35, Baseline

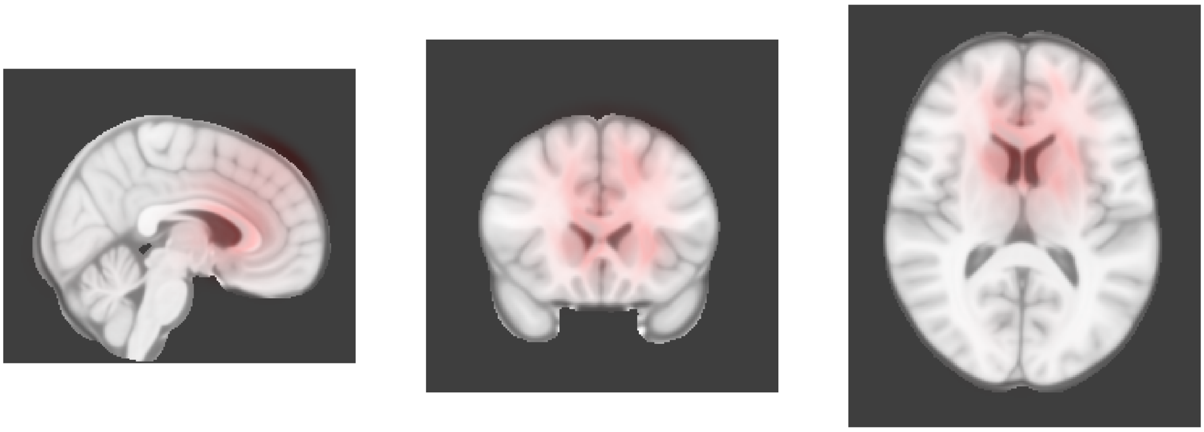

Dimension 35, 2-year follow-up

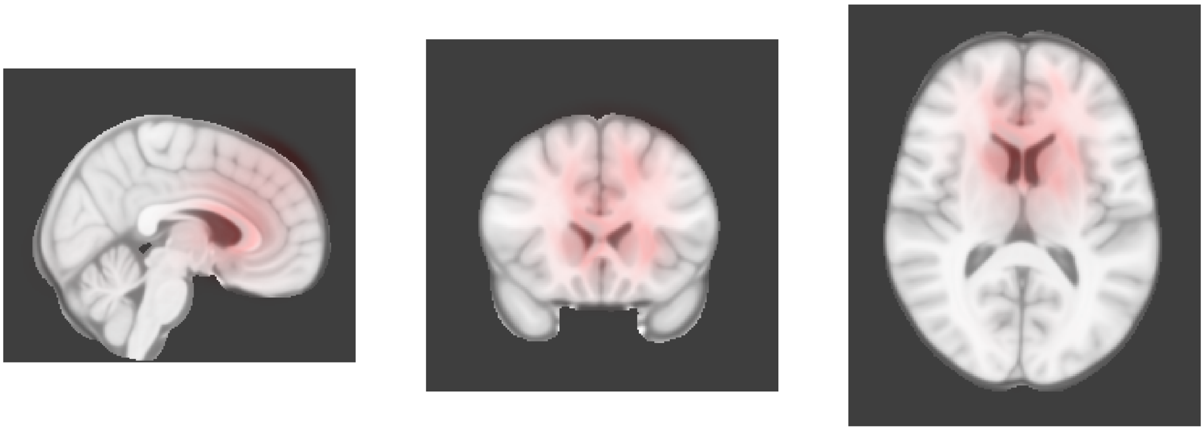

Dimension 36, Baseline

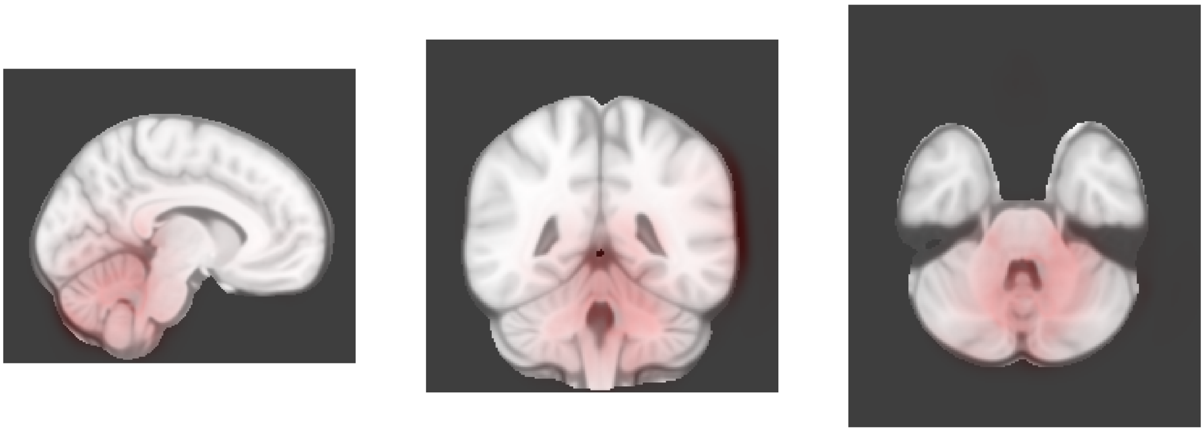

Dimension 36, 2-year follow-up

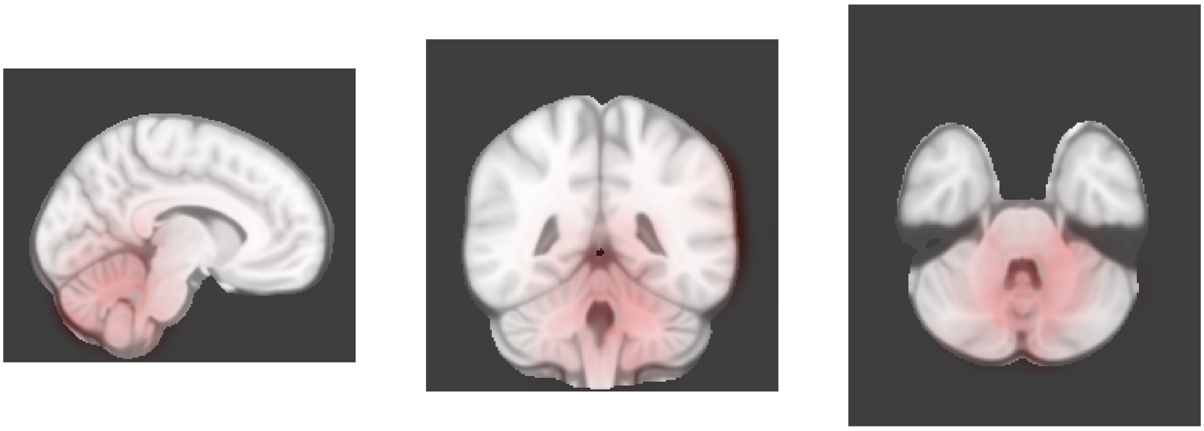

Dimension 37, Baseline

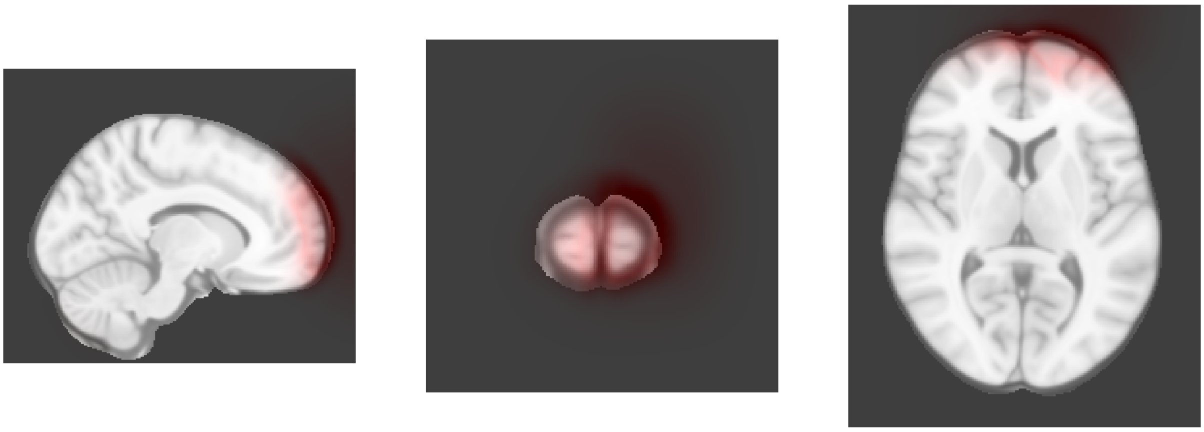

Dimension 37, 2-year follow-up

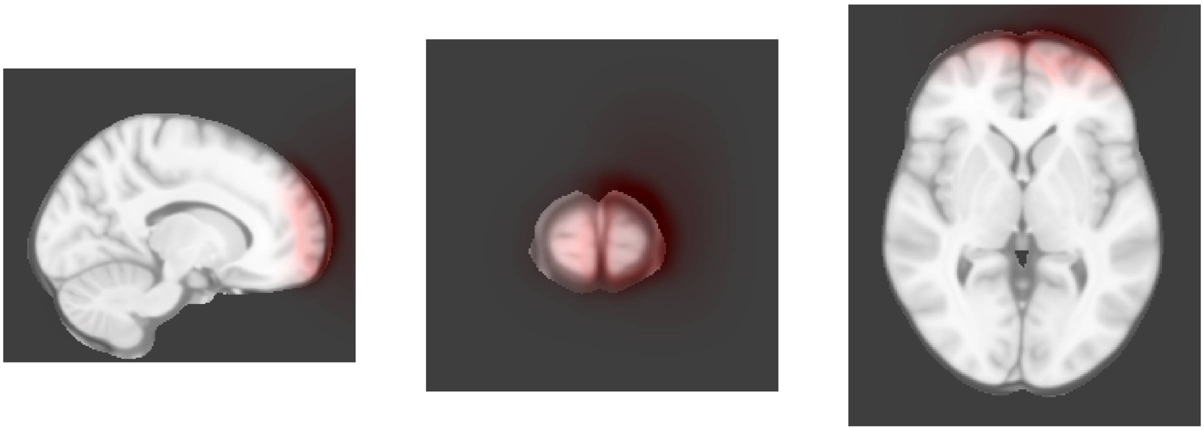

Dimension 38, Baseline

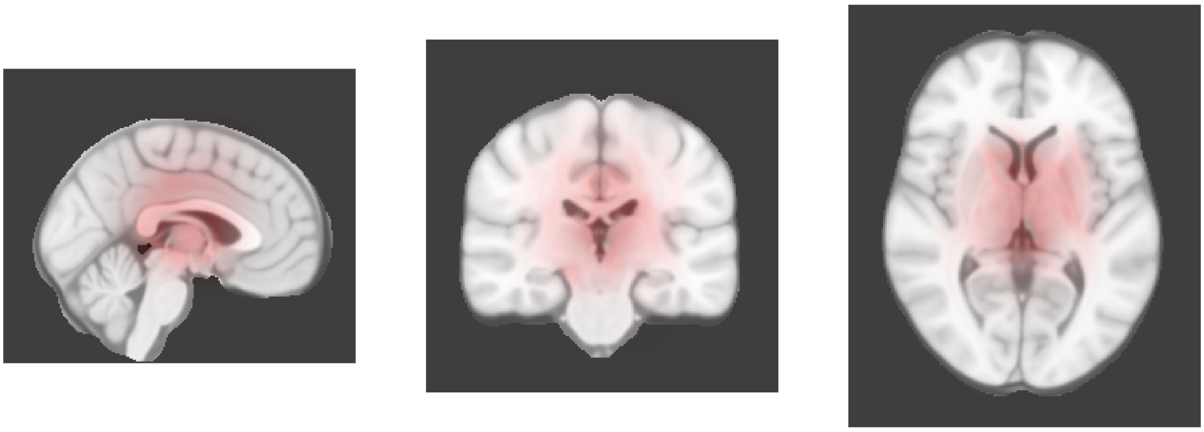

Dimension 38, 2-year follow-up

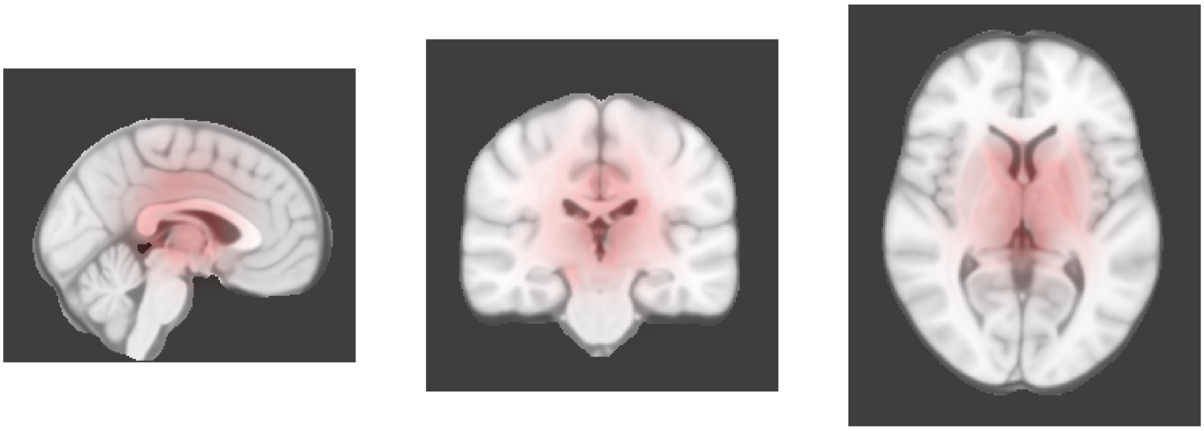

Dimension 39, Baseline

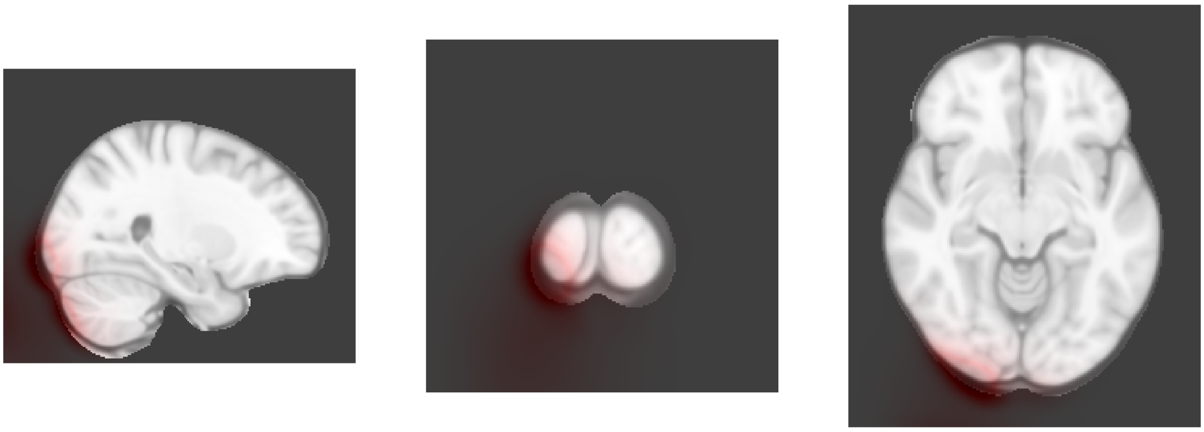

Dimension 39, 2-year follow-up

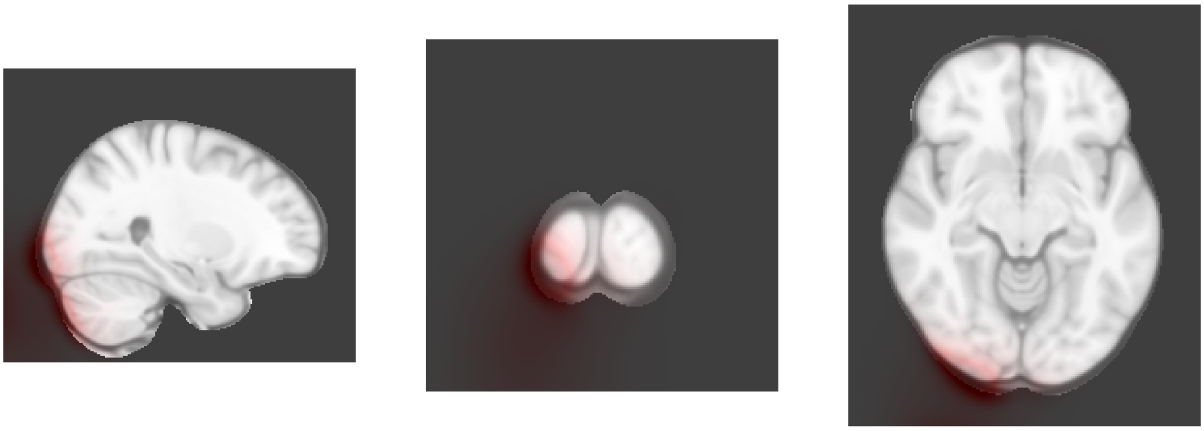

Dimension 40, Baseline

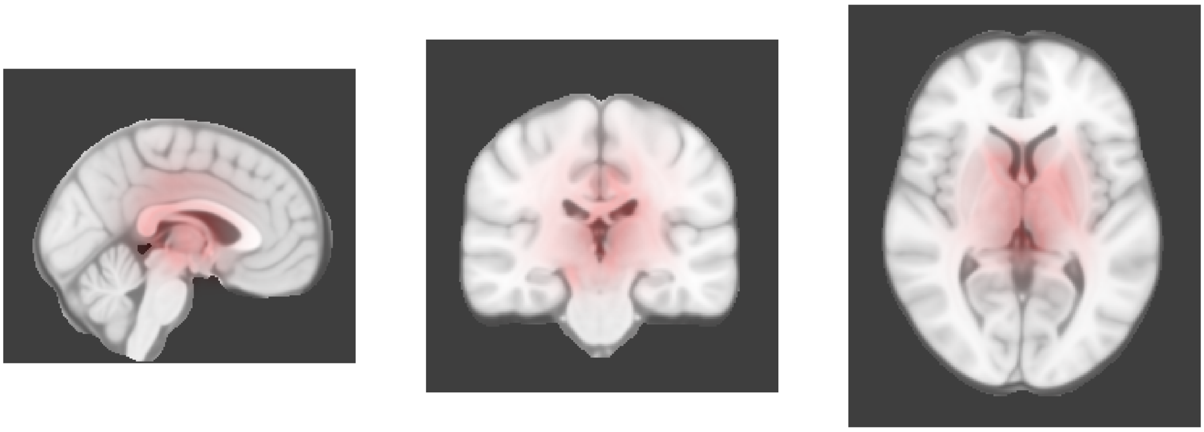

Dimension 40, 2-year follow-up

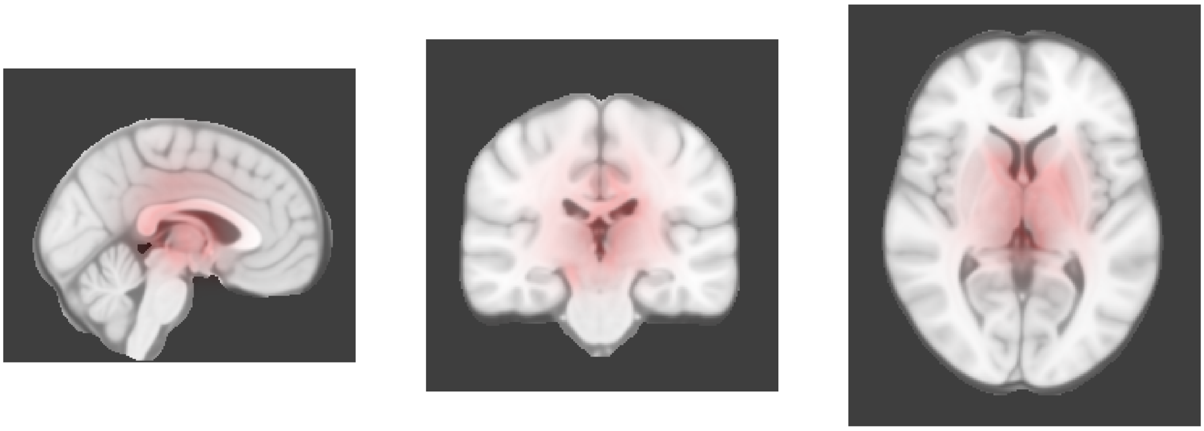

Dimension 41, Baseline

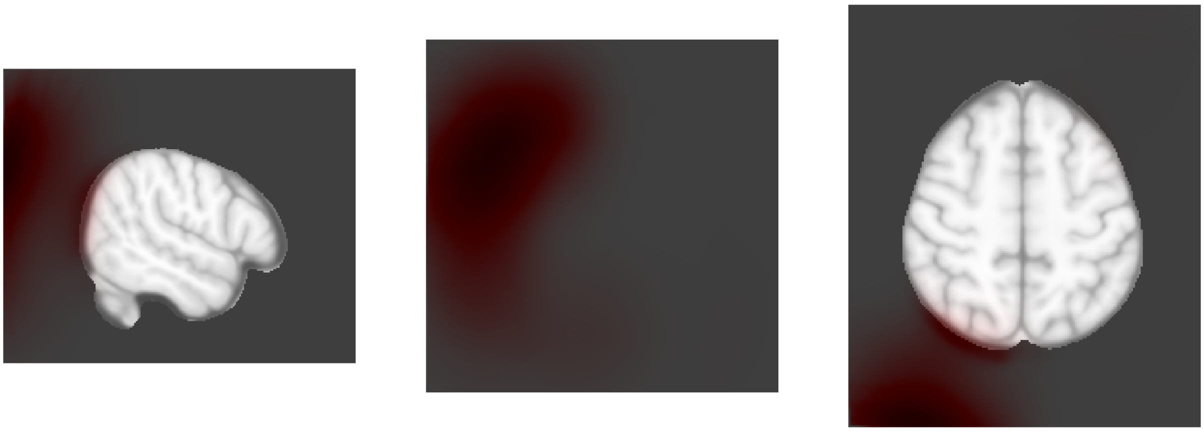

Dimension 41, 2-year follow-up

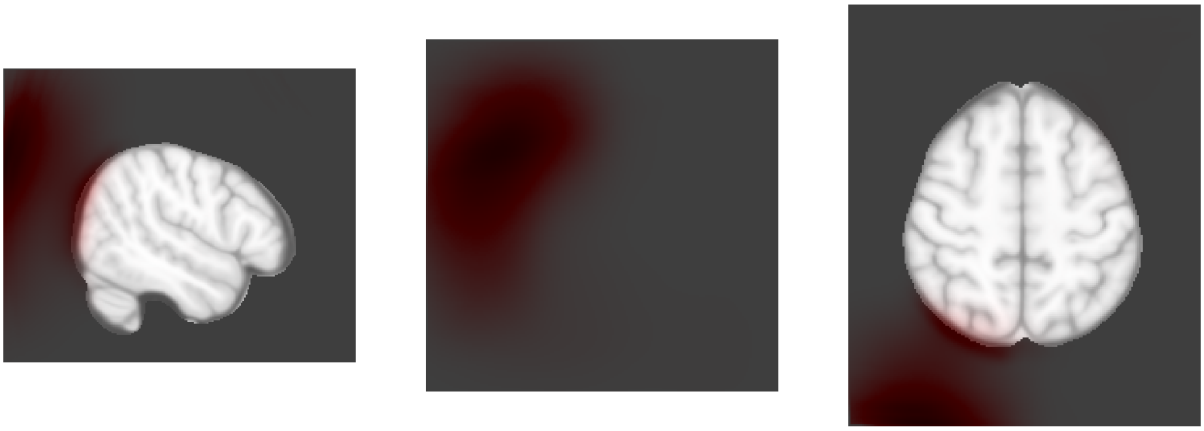

Dimension 42, Baseline

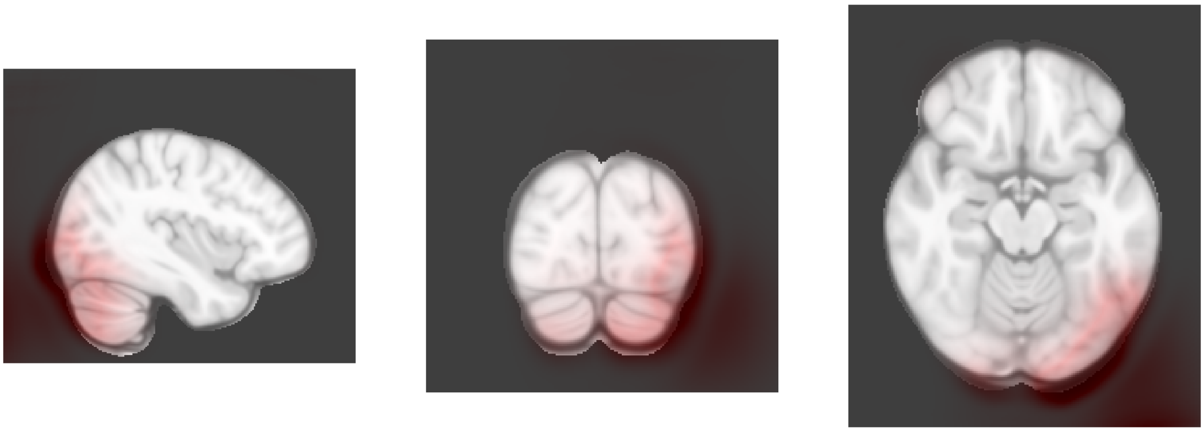

Dimension 42, 2-year follow-up

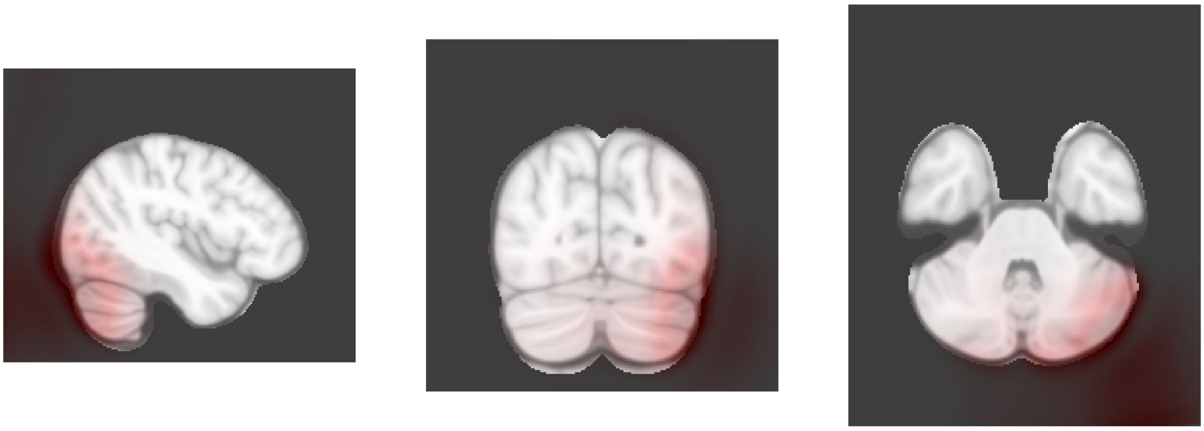

Dimension 43, Baseline

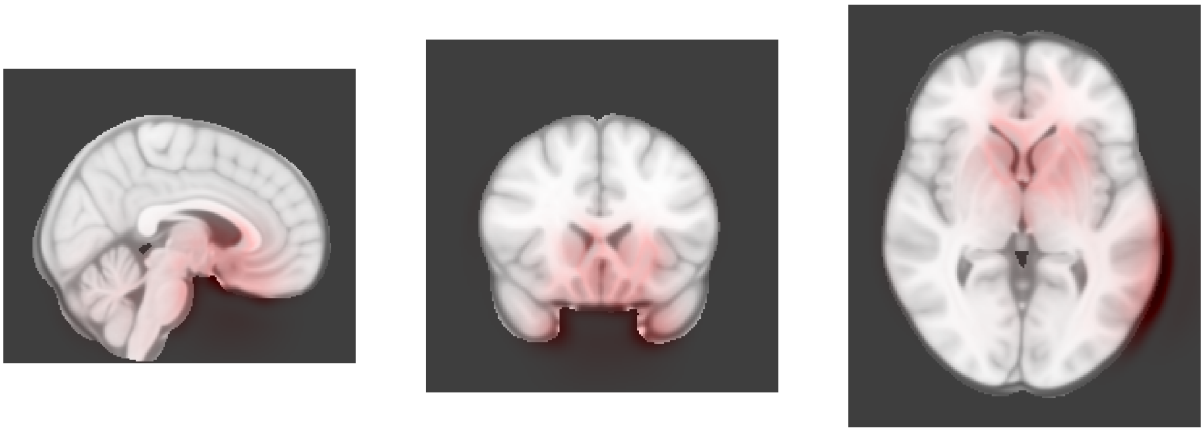

Dimension 43, 2-year follow-up

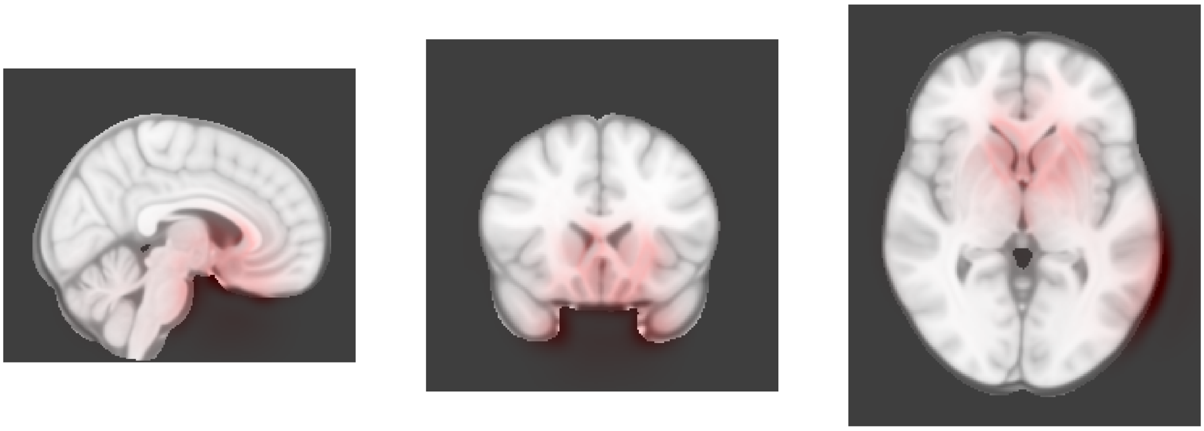

Dimension 44, Baseline

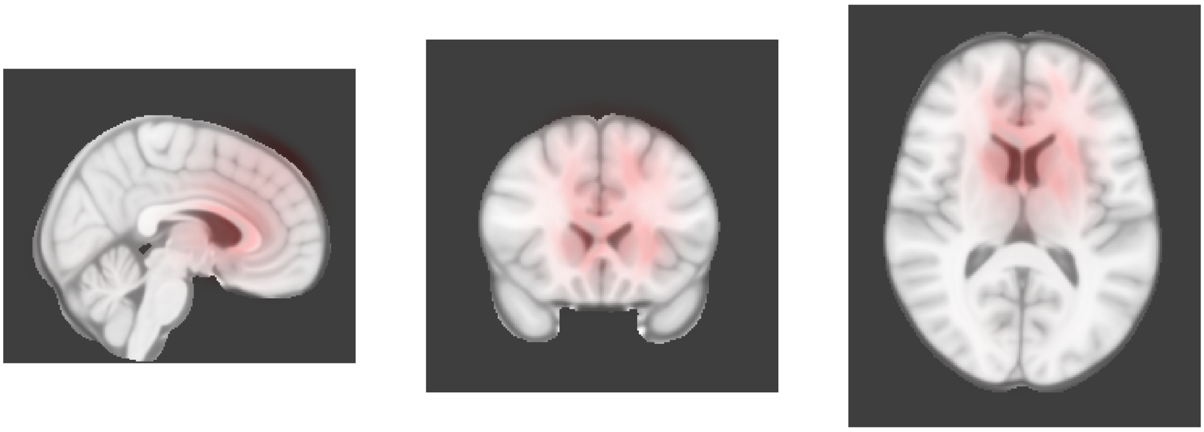

Dimension 44, 2-year follow-up

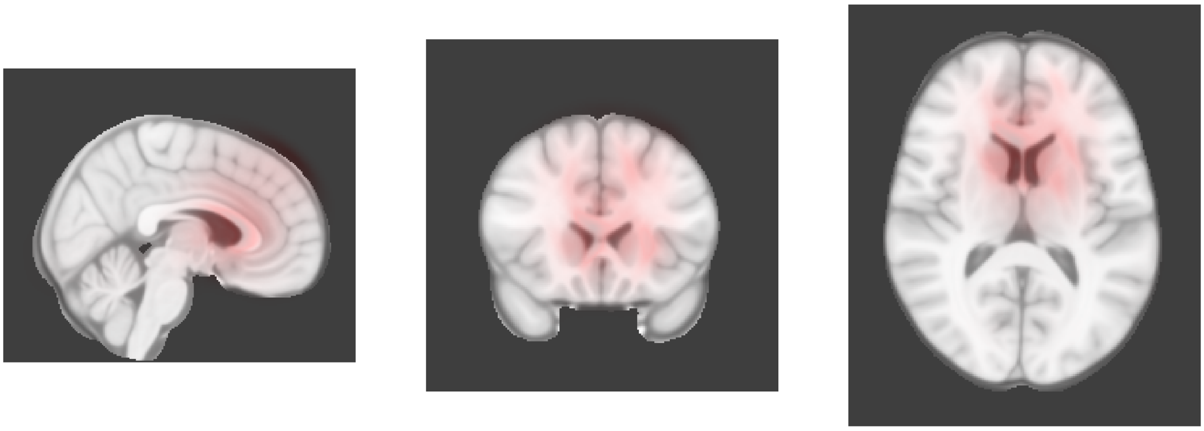

Dimension 45, Baseline

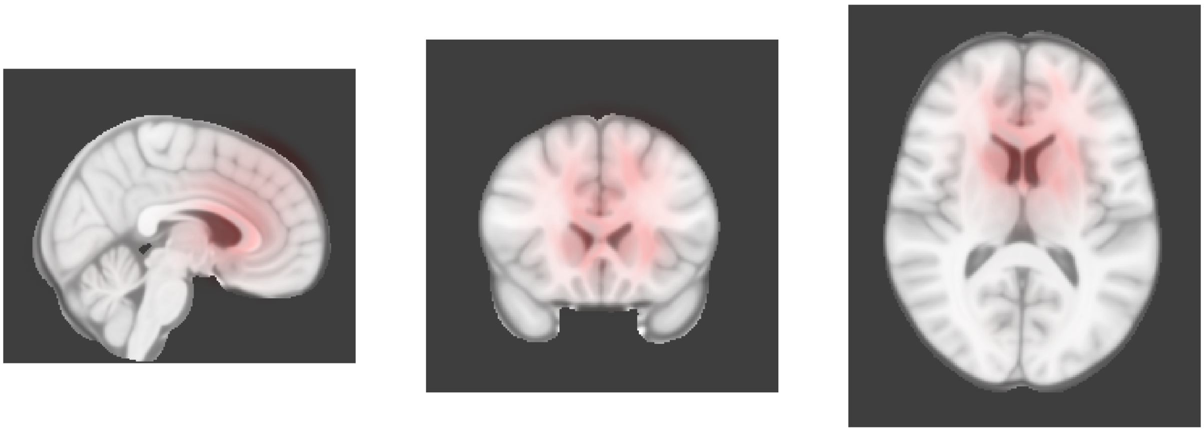

Dimension 45, 2-year follow-up

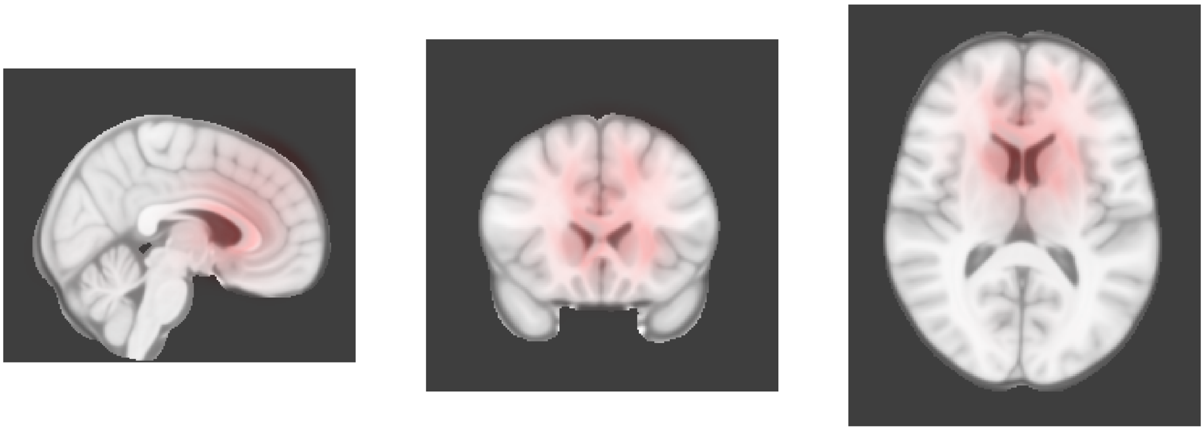

Dimension 46, Baseline

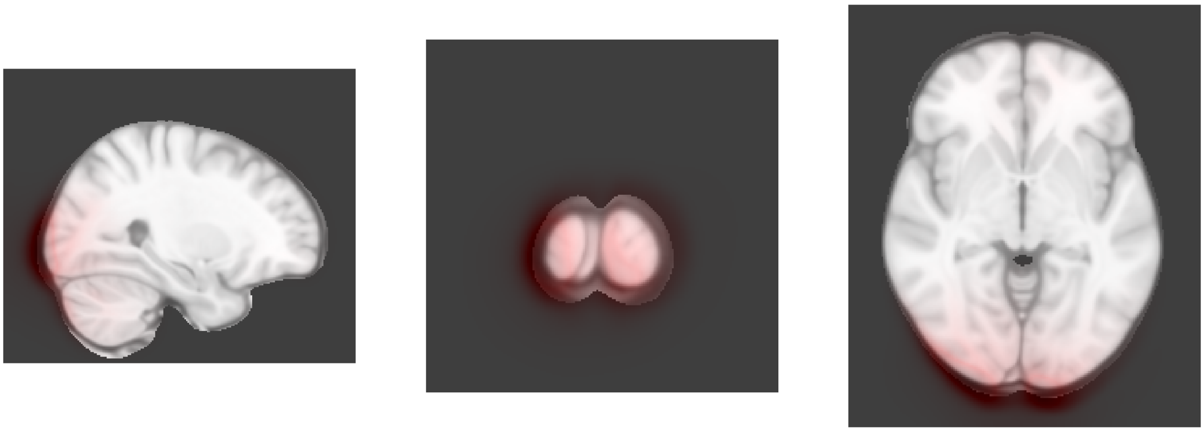

Dimension 46, 2-year follow-up

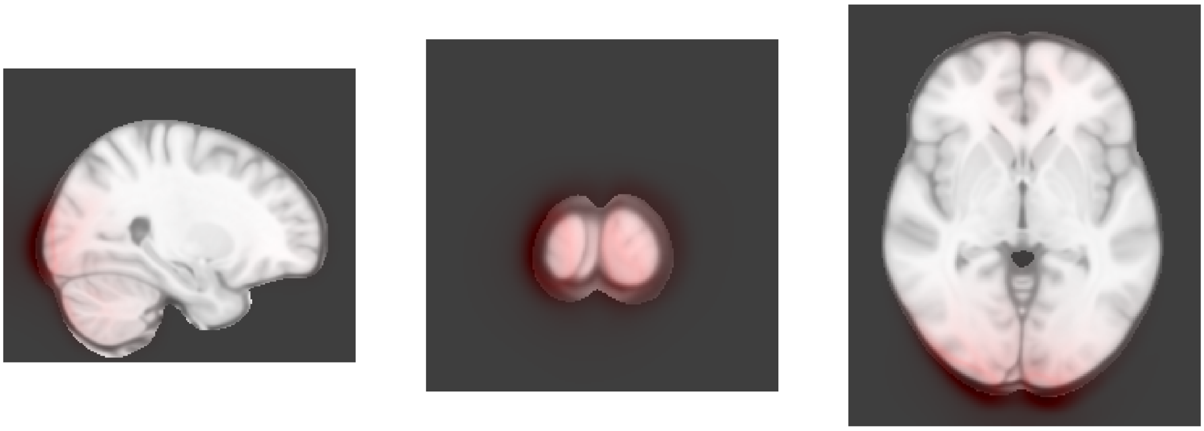

Dimension 47, Baseline

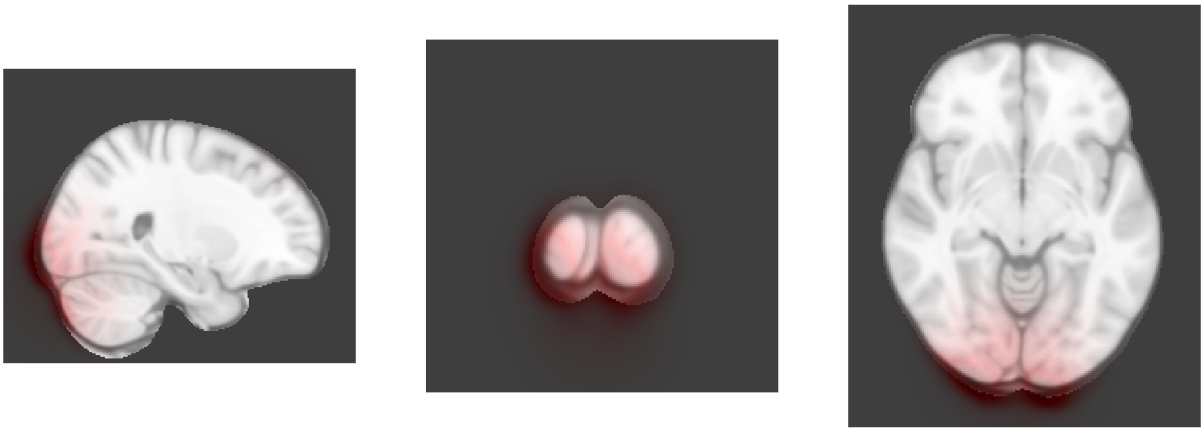

Dimension 47, 2-year follow-up

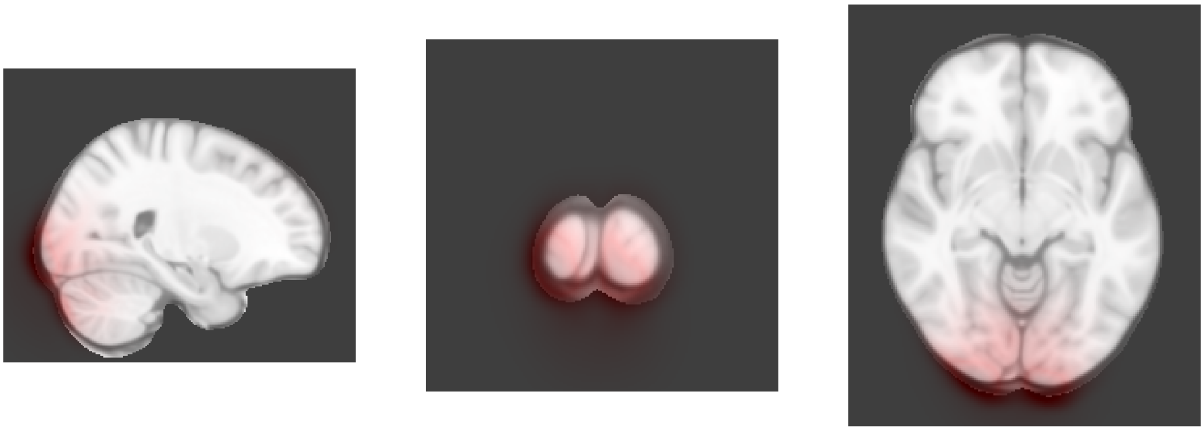

Dimension 48, Baseline

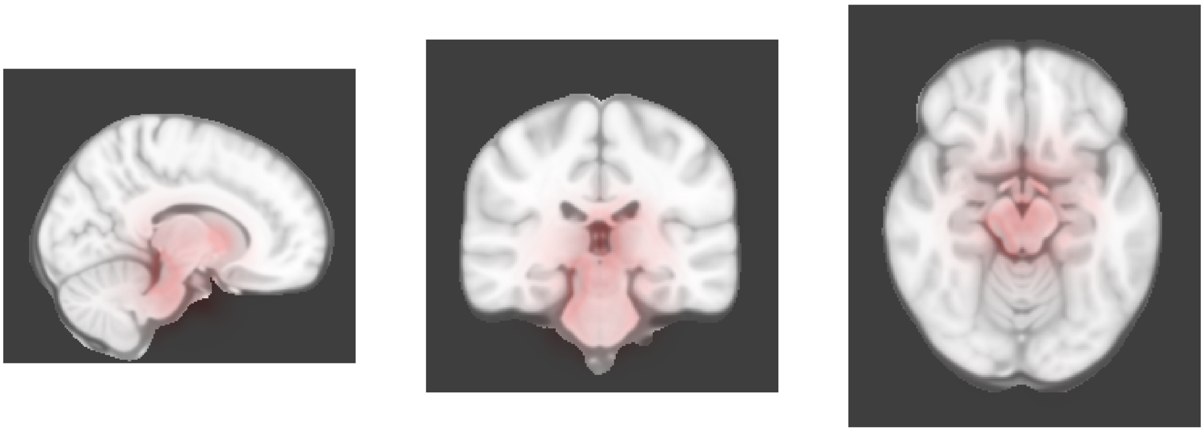

Dimension 48, 2-year follow-up

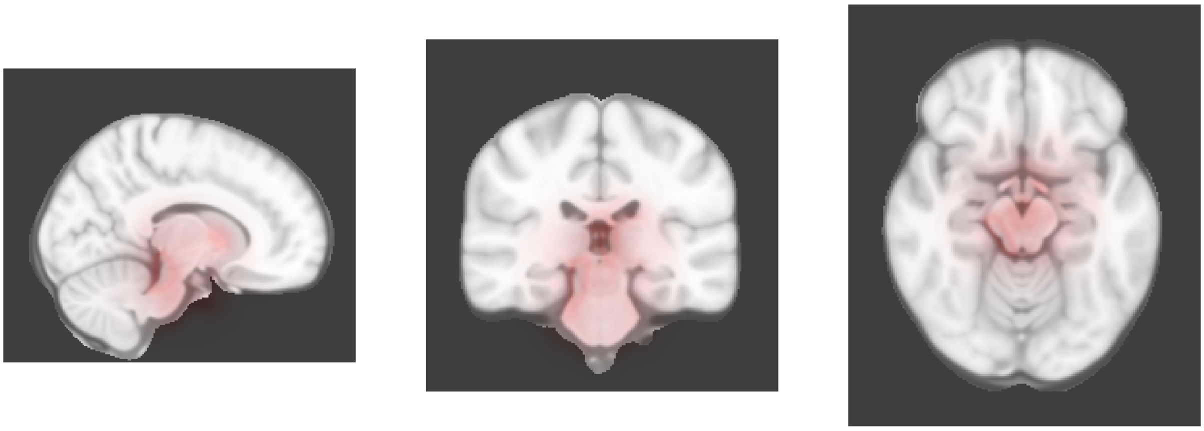

Dimension 49, Baseline

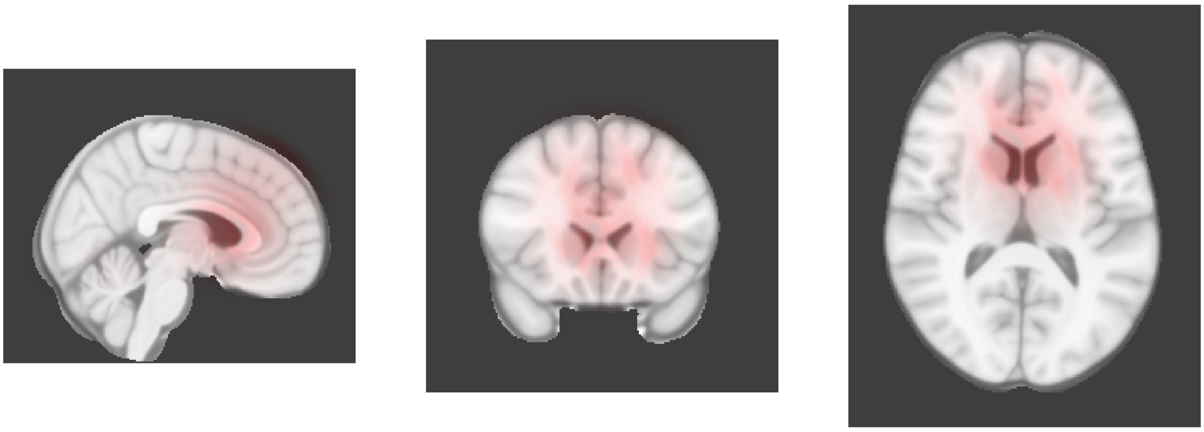

Dimension 49, 2-year follow-up

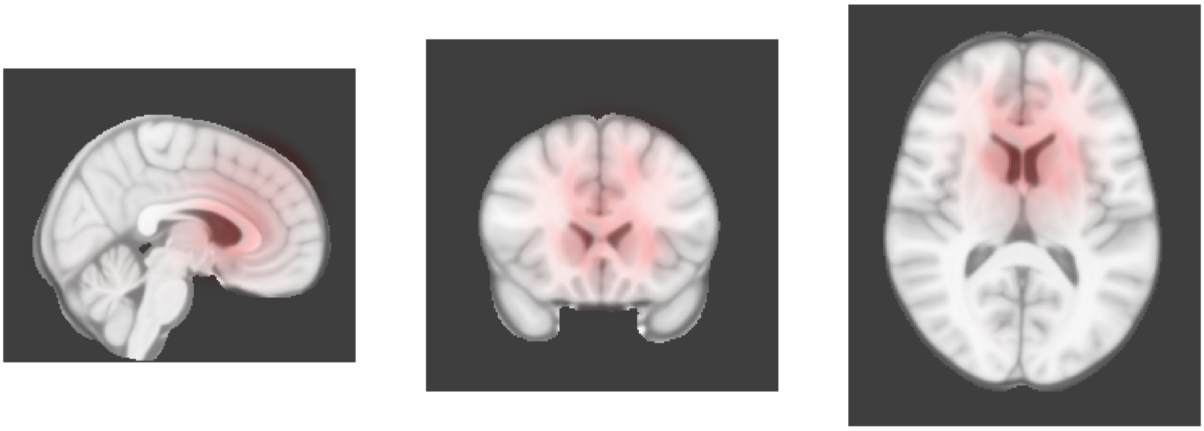

Dimension 50, Baseline

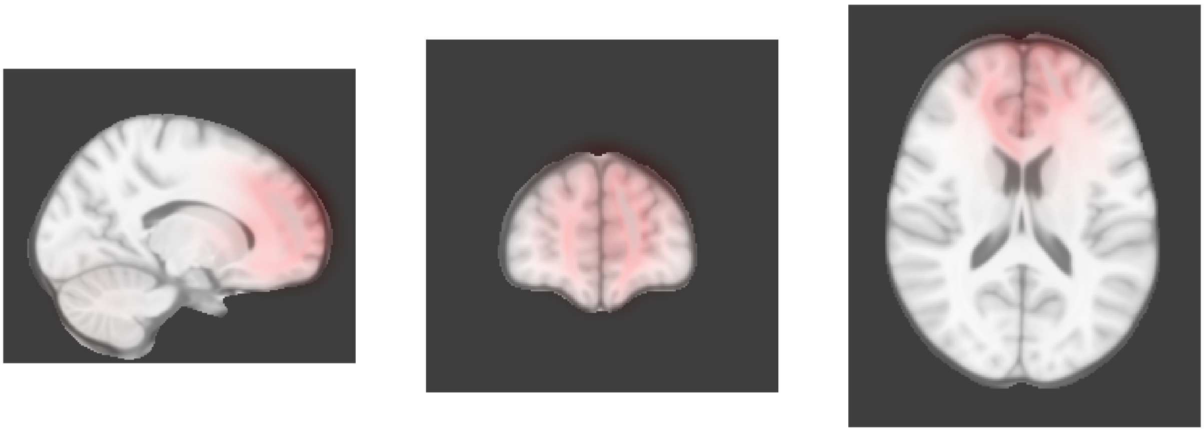

Dimension 50, 2-year follow-up

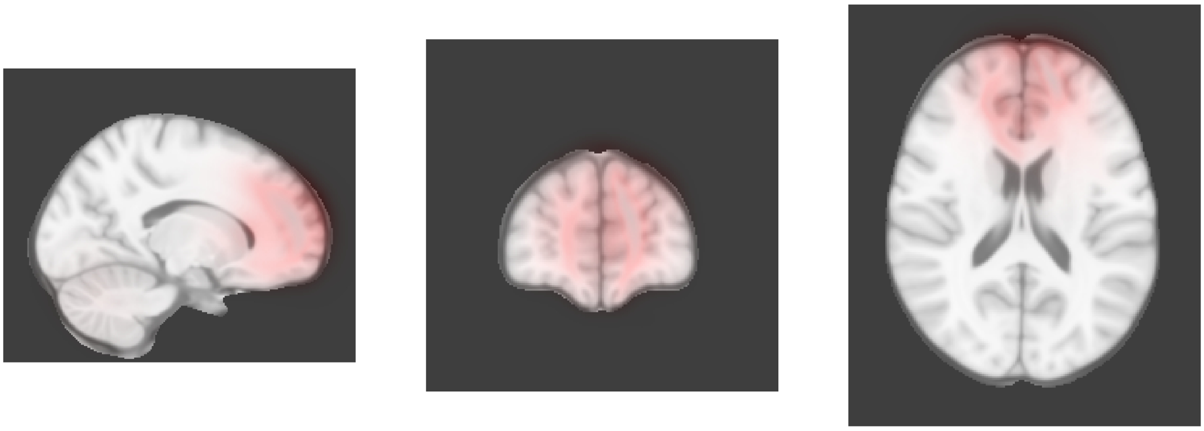

Dimension 52, Baseline

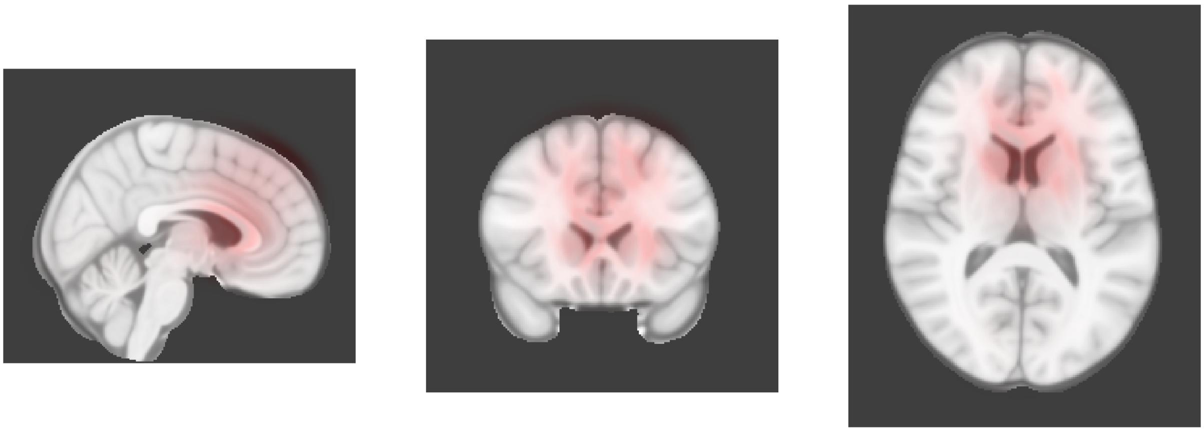

Dimension 52, 2-year follow-up

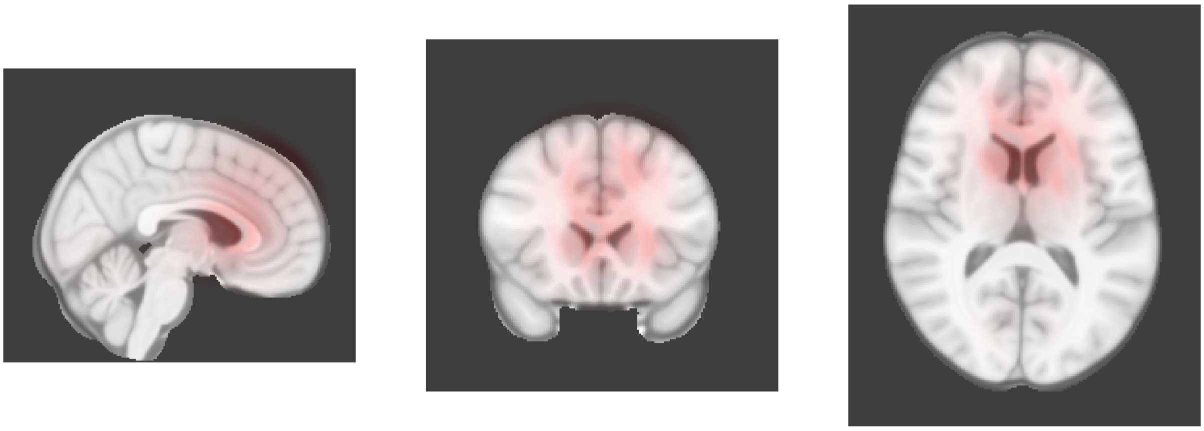

Dimension 53, Baseline

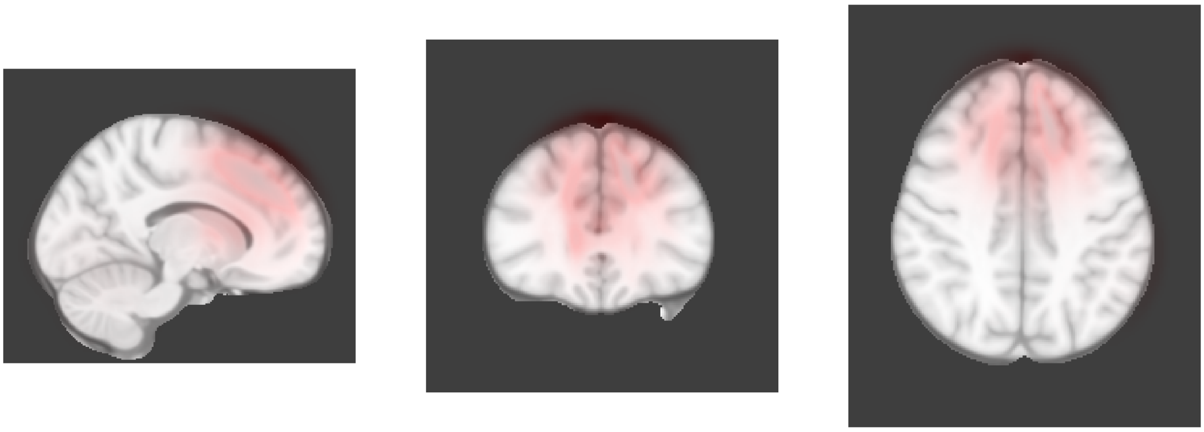

Dimension 53, 2-year follow-up

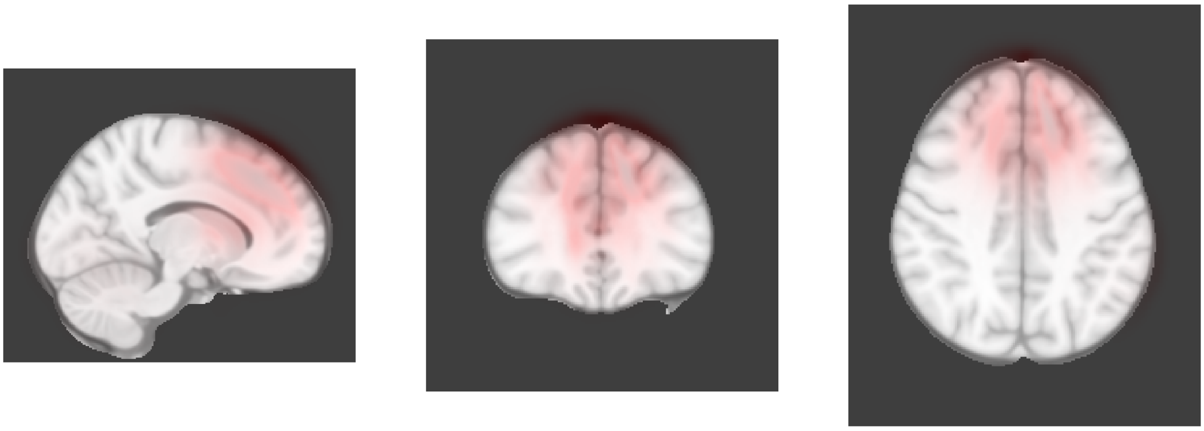

Dimension 54, Baseline

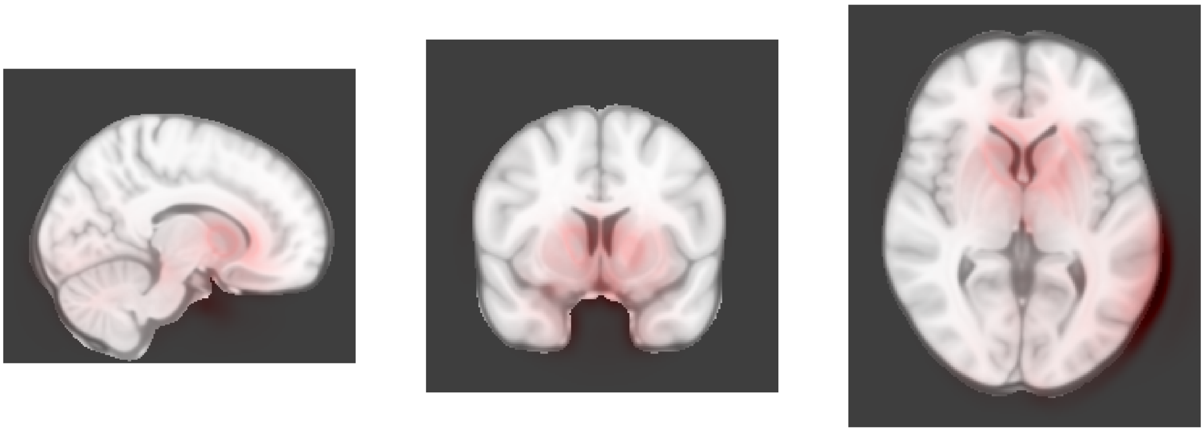

Dimension 54, 2-year follow-up

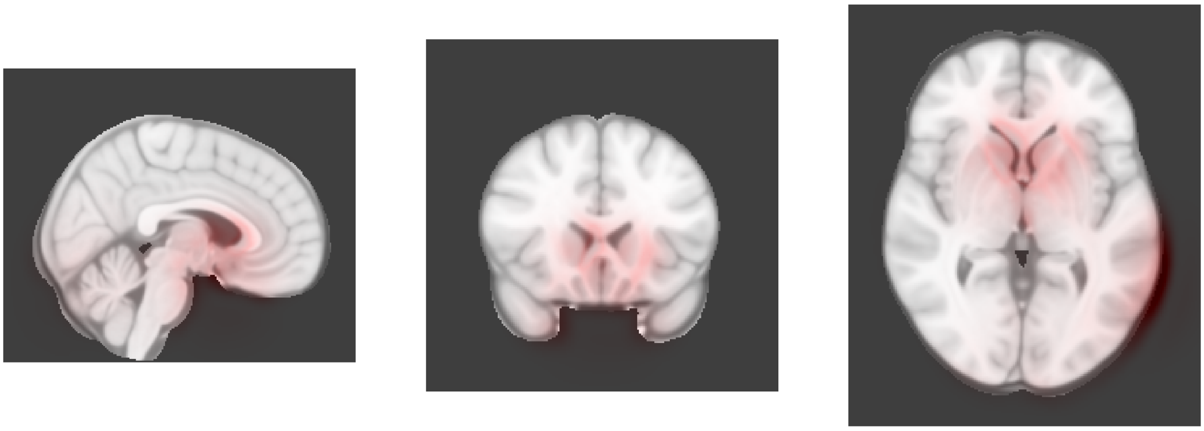

Dimension 55, Baseline

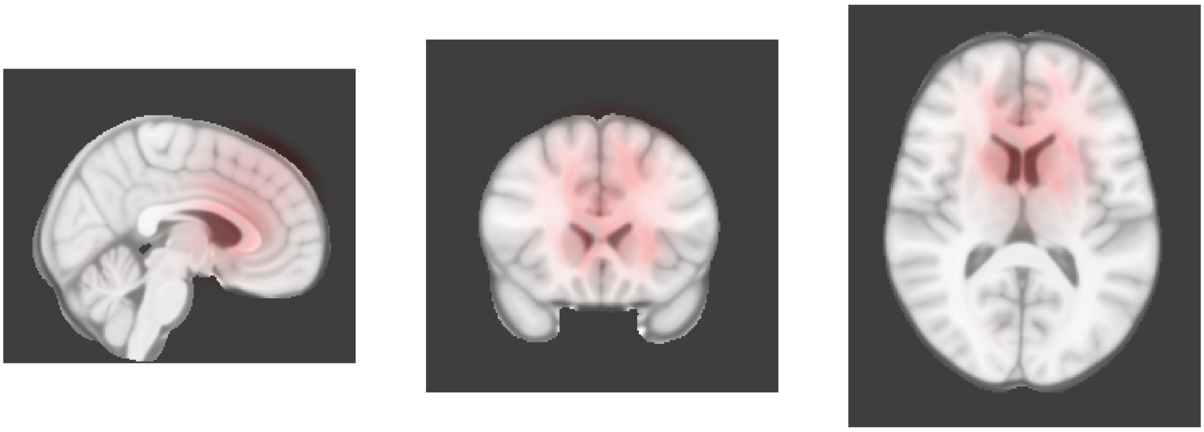

Dimension 55, 2-year follow-up

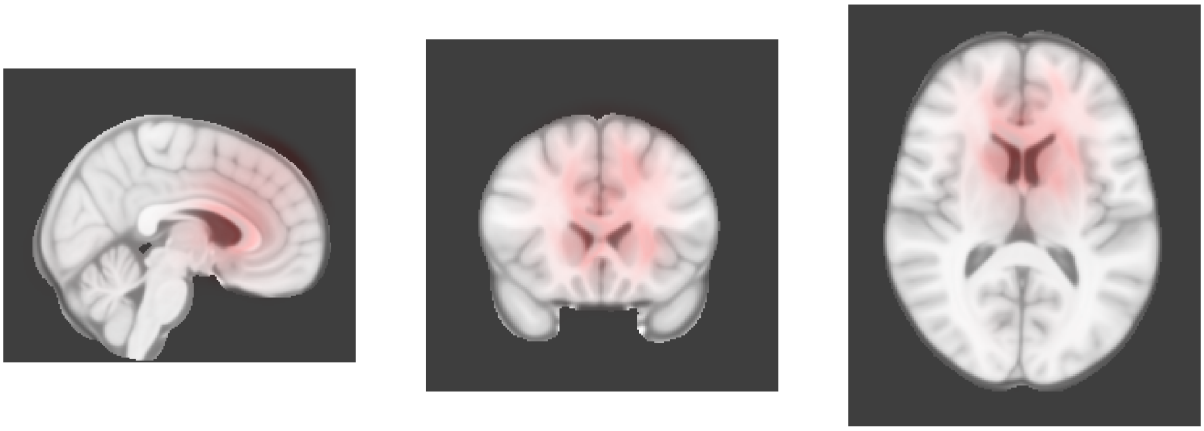

Dimension 56, Baseline

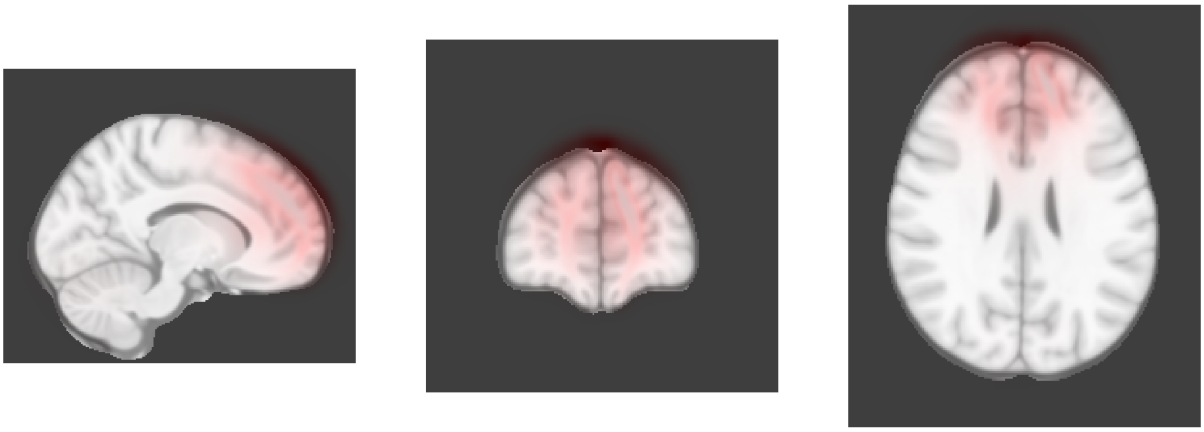

Dimension 56, 2-year follow-up

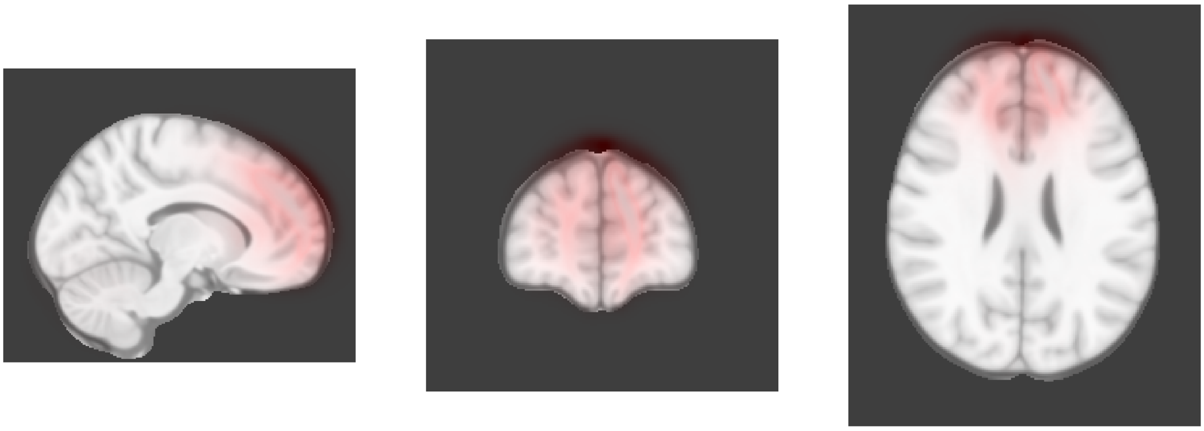

Dimension 58, Baseline

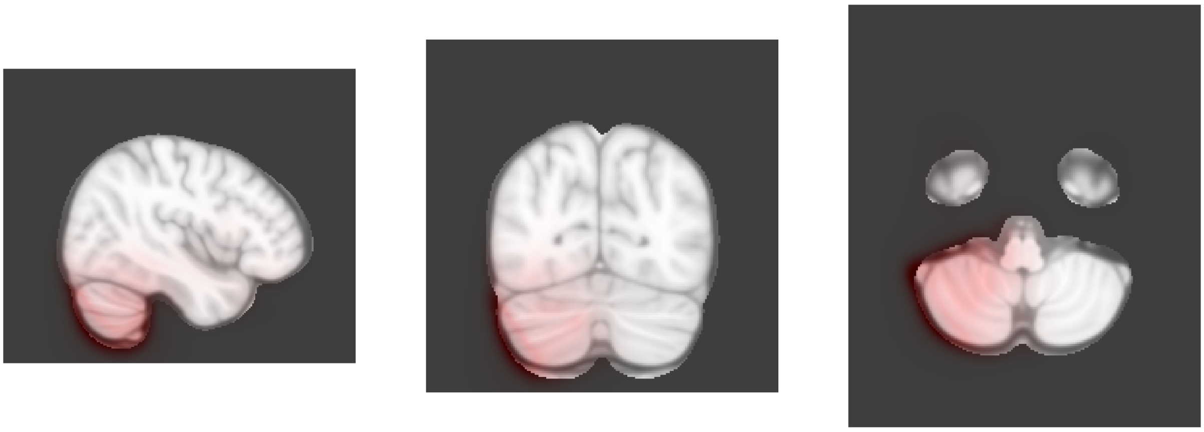

Dimension 58, 2-year follow-up

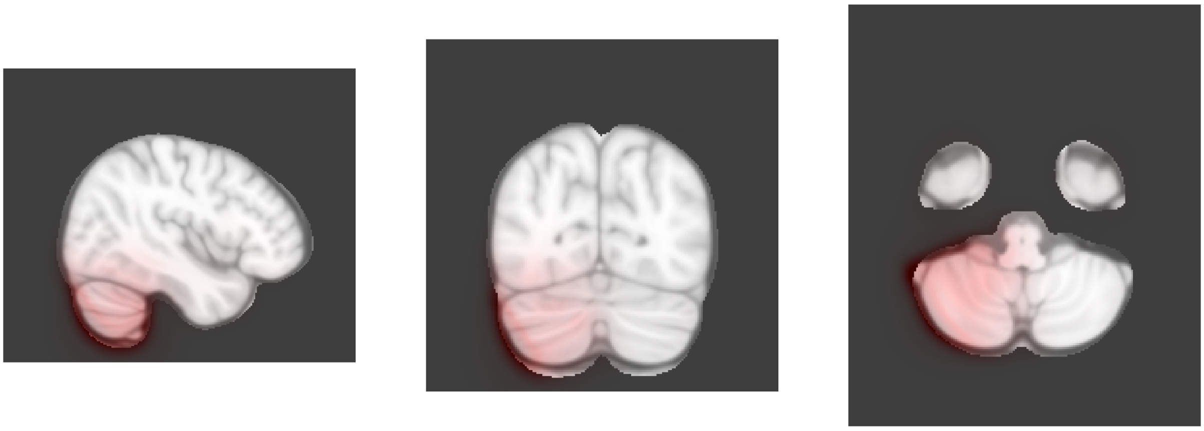

Dimension 60, Baseline

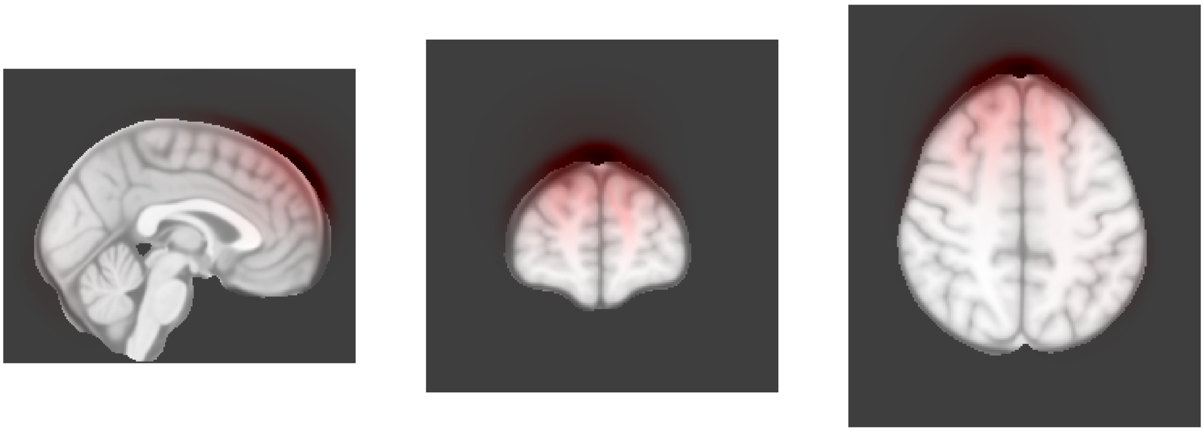

Dimension 60, 2-year follow-up

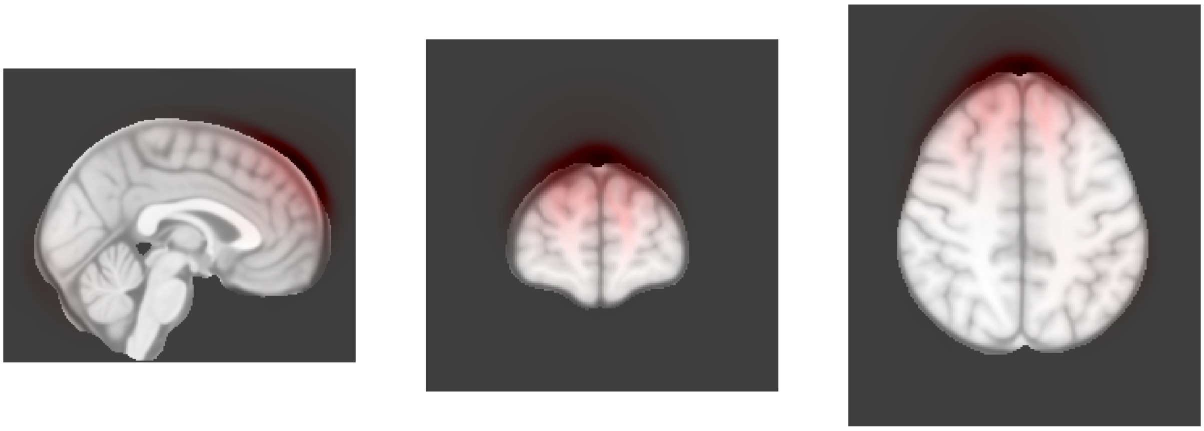

Dimension 61, Baseline

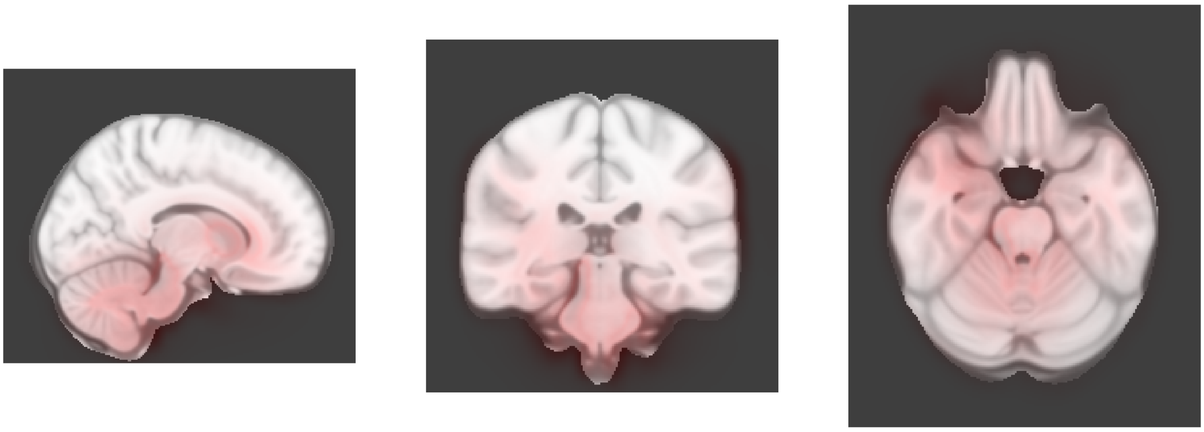

Dimension 61, 2-year follow-up

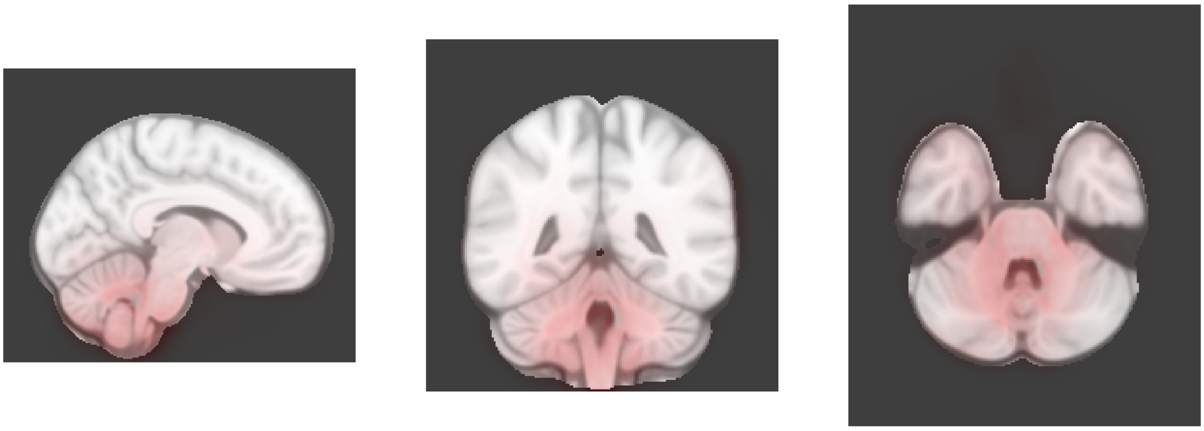

Dimension 62, Baseline

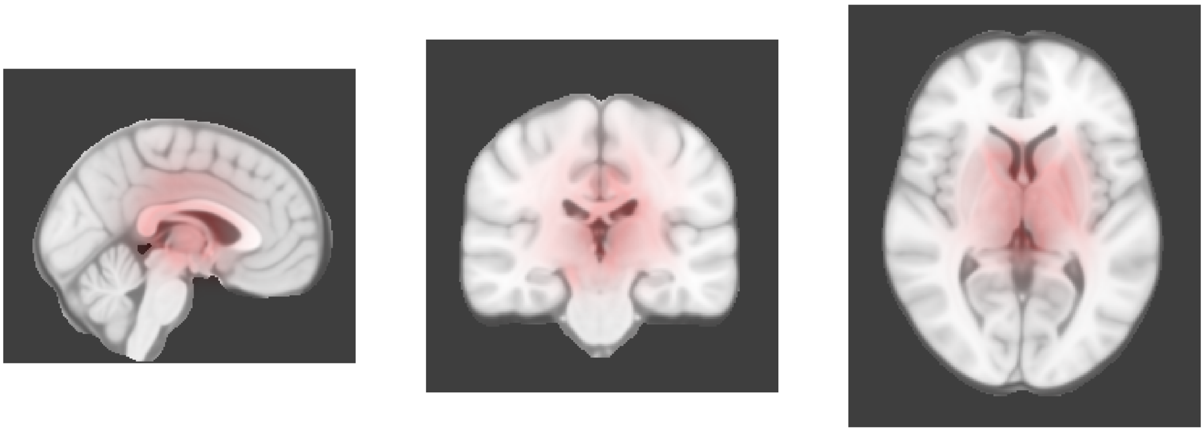

Dimension 62, 2-year follow-up

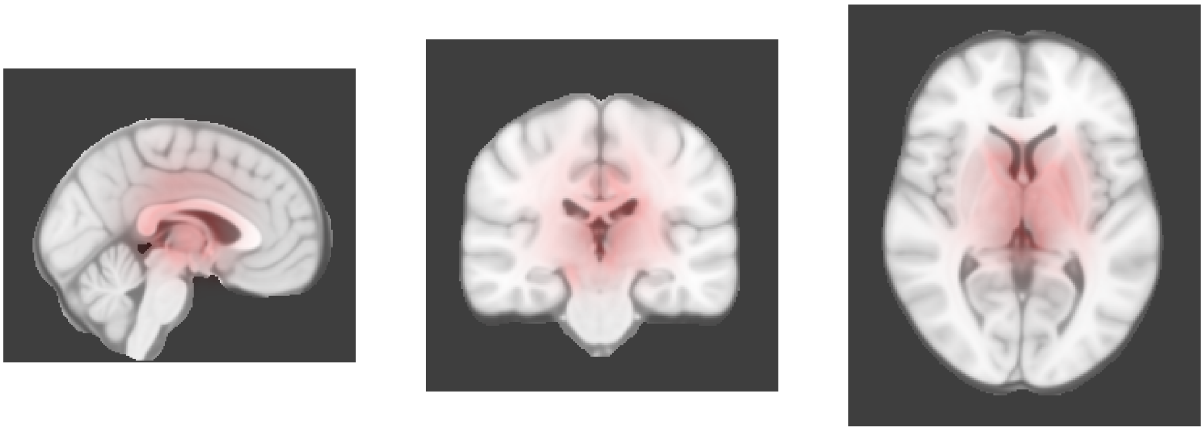

Dimension 63, Baseline

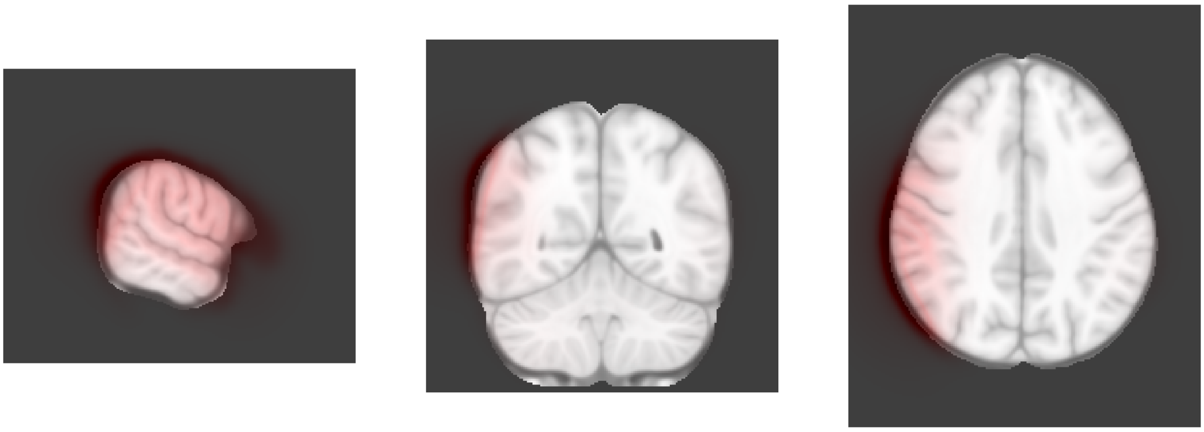

Dimension 63, 2-year follow-up

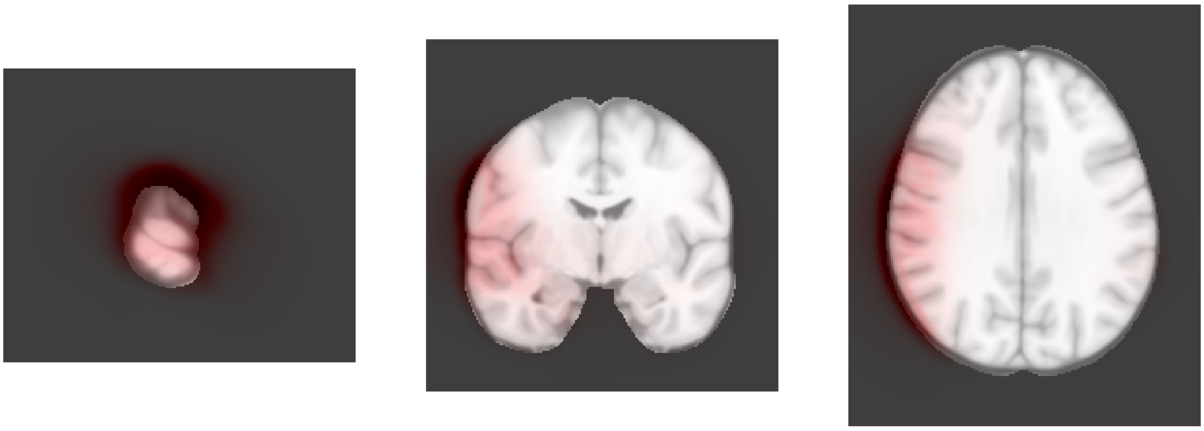
